# Identification of Cell-Type Specific Alternative Transcripts in the Multicellular *Alga Volvox carteri*

**DOI:** 10.1101/2023.01.10.523483

**Authors:** Ravi N. Balasubramanian, James Umen

## Abstract

Cell type specialization is a hallmark of complex multicellular organisms and is usually established through implementation of cell-type-specific gene expression programs. The multicellular green alga *Volvox carteri* has just two cell types, germ and soma, that have previously been shown to have very different transcriptome compositions which reflect differences in their respective forms and functions. Here we interrogated another potential mechanism for differentiation in *V. carteri*, cell type specific alternative transcript isoforms (CTSAI). We used pre-existing predictions of alternative transcripts and *de novo* transcript assembly to compile a list of 1978 loci with two or more transcript isoforms, 67 of which also showed cell type isoform expression biases. Manual curation identified 15 strong candidates for CTSAI, three of which were experimentally verified and provide insight into potential functional differentiation of encoded protein isoforms. Alternative transcript isoforms are also found in a unicellular relative of *V. carteri*, *Chlamydomonas reinhardtii*, but there was little overlap in orthologous gene pairs in the two species which both exhibited CTSAI, suggesting that CTSAI observed in *V. carteri* arose after the two lineages diverged. CTSAIs in metazoans are often generated through alternative pre-mRNA processing mediated by RNA binding proteins (RBPs). We interrogated cell type expression patterns of 126 *V. carteri* predicted RBP encoding genes and found 40 that showed either somatic or germ cell expression bias. These RBPs are potential mediators of CTSAI in *V. carteri* and suggest possible pre-adaptation for cell type specific RNA processing and a potential path for generating CTSAI in the early ancestors of metazoans and plants.

## INTRODUCTION

Cell type specialization is a key feature of multicellular organisms that enables development of complex forms and functions, as exemplified by plants and animals with hundreds of cell types forming complex tissues and organs [1, 2]. All multicellular eukaryotes evolved from single-celled ancestors with limited capacity for cell-type specialization, and one of the major questions in evolutionary biology is in understanding the origins of cell type specialization in multicellular lineages [1, 3, 4]. In the case of animals and plants, their complexity and ancient divergence from unicellular ancestors makes it challenging to infer how cell type specialization may have arisen.

A major contributor to cell type specialization is differential gene expression. Differential expression is most commonly achieved through combinatorial activities of different transcription factors that control morphogenesis and cell differentiation programs for each cell type in a multicellular individual. Modulation of gene expression through small RNAs (e.g. microRNAs) is another means of controlling gene expression that is widespread in plants and animals [5, 6]. Both of the above mechanisms modulate transcript abundance, but do not contribute directly to functional diversity of genes or gene products. Many multicellular eukaryotes employ a third mechanism for regulating gene expression that enables distinct cell-type-specific functions to be expressed from a single genetic locus using alternative transcript isoforms. Differential transcript isoform expression occurs via alternative splicing of pre-mRNAs and/or use of alternative transcription start/termination sites [7, 8]. The different transcript isoforms may specify production of structurally and functionally different proteins from the same locus, most dramatically exemplified by complex alternative splicing of some neuronal transcripts in animal brains [9]. This mechanism can promote cell type differentiation when alternative isoforms encode proteins with distinct functional properties. Another less common but well-documented phenomenon for controlling gene expression involves convergent overlapping anti-sense transcripts produced from adjacent loci, where the expression of one transcript antagonizes production of its antisense partner [10, 11]. It remains unclear to what extent each of these mechanisms may have been employed during the early transitions of life from single cells to multicellular individuals.

The volvocine algae are a related group of green algae within the Order Chlamydomonadales and are a model system for investigating the step-wise acquisition of multicellular organization and emergent behaviors [12, 13]. The unicellular volvocine species *Chlamydomonas reinhardtii* is an outgroup that is sister to different multicellular volvocine genera that are classified based on their cell number, organismal size and degree of cell type specialization [12, 14]. The estimated divergence time of volvocine algae from a unicellular ancestor is around 200-250 MYA, which is relatively recent compared to plants and animals that shared a common ancestor with unicellular relatives >500 and >600-800 MYA respectively [12, 15, 16]. Multicellular *Volvox carteri* belongs to the most developmentally complex volvocine genus and has evolved some hallmark features of more complex multicellular organisms including germ-soma cell type differentiation and morphogenetic patterning. Each vegetative phase *V. carteri* individual is a spheroid with a diameter of up to 500 μm and contains ~2000 cells of just two types embedded in a clear structured extracellular matrix (ECM). On the periphery of each spheroid are ~2000 bi-ciliated terminally differentiated somatic cells that provide motility and are specialized for secretion of ECM [17]. Within each spheroid are around 16 large non-ciliated reproductive cells termed gonidia. Upon reaching maturity, each gonidium undergoes embryogenesis which involves a programmed series of twelve or thirteen symmetric and asymmetric cleavage divisions and morphogenesis to produce a juvenile spheroid with ~2000 small somatic precursor cells and 12-16 large gonidial precursor cells. Cell type differentiation is controlled in an unknown manner by cell size at the end of embryogenesis, with cells below 8μm diameter adopting a somatic fate and cells above 8μm diameter adopting a gonidial fate [18]. Juveniles remain within their mother spheroid for around a day as they continue to grow, and eventually hatch and mature to begin the two-day vegetative life cycle again.

Early work on cell type differentiation in *Volvox carteri* (Volvox) focused on gene expression programs that are different between the two cell types [19] and light-dependent translational regulation [20]. The Volvox genome sequence [21] facilitated application of more complete transcriptome profiling in the two cell types. Matt & Umen [17] showed a high degree of transcriptome asymmetry between the two vegetative Volvox cell-types suggesting that, like in many other developmental systems, differential transcript abundance is probably the most important component of cell type differentiation. While transcriptional regulation is expected to be a major factor in cell type differentiation in Volvox, other regulatory mechanisms may also be involved including microRNAs, some of which show preferential cell type expression [22, 23]. *C. reinhardtii* (Chlamydomonas) also possesses miRNAs and siRNAs that resemble those in multicellular eukaryotes, implying that such complex RNA-silencing mechanisms evolved before multicellularity [24]. However, there exists little conservation between the Chlamydomonas and the Volvox miRNAs [22–24] suggesting rapid evolution of miRNA loci in the volvocine algae.

An additional potential mechanism for cell-type differentiation that has not been previously examined systematically for Volvox or other multicellular volvocine species is cell-type specific alternative transcript isoforms (CTSAI) that can be generated through alternative transcription initiation/termination sites or through alternative pre-mRNA processing. Here we made use of prior annotations and previously published cell-type-specific transcriptome data to identify candidate loci with transcripts exhibiting CTSAI. Using two independent prediction methods for alternative isoforms, we found less than 20% of expressed genes had more than one isoform. Among these, the number of genes showing cell type bias in expression of one or both isoforms was less than 100 total, including a high proportion of false positives. Nonetheless, some predicted cases of cell type specific transcript isoforms could be verified and provided potential insights into *V. carteri* cell-type specification. A possible mechanism for controlling cell type specific RNA processing in Volvox could come from RNA binding proteins (RBPs) which can mediate alternative splicing in plants or animals. We found several dozen predicted RBP genes in *Volvox* that are expressed in a cell type specific manner, and which could be mediators of CTSAS or CTSAI.

## METHODS

### HISAT2 RNA-seq protocol

We used HISAT2 RNA-seq software [25] (Supplemental Fig. S1) to analyze previously published *Volvox carteri* transcriptome data. First, RNA-seq reads obtained from Matt & Umen, 2018 [17] were passed through HISAT2 and aligned to a reference annotation of the Volvox genome available from Phytozome [26, 27]. We made annotation file following the methods of Matt and Umen, 2018 [17] by combining all 14,247 V. carteri version 2.1 gene models currently available on Phytozome [21, 26] with 1761 nonoverlapping (<20% sequence overlap) version 2.0 Phytozome gene models [21, 26] to generate a final set of 16,008 unique protein-coding gene models. Aligned reads were merged together into transcripts whose abundances were estimated using the StringTie software package. We then used StringTie to merge the transcripts across samples to get a full transcriptome for analysis, allowing us to restore transcripts with missing fragments in individual samples to their full length. Transcript abundances were then re-estimated by StringTie and normalized in Fragments per Kilobase per Million Reads (FPKM). Identification numbers were assigned to each transcript and gene, and when possible, gene names from the reference annotation in Phytozome [21, 26, 27] were associated with each locus. Previously unidentified transcripts were indexed by the Ballgown-generated ID number of the associated gene. We applied an expression filter of 1 FPKM to each transcript across all samples to remove low abundance transcripts prior to our analysis. Finally, we used the Ballgown software package [28] to identify candidate transcript isoforms with significant differences in expression between somatic and gonidial transcriptomes.

In addition to the analysis above that identified transcript isoforms de novo from mapped read data, we also directly queried loci with more than one transcript isoform predicted by Phytozome version 2.1 for Volvox using the same expression filtering criterion of >1 FPKM in at least one of the two cell types. These data we also used to validate predictions from Ballgown.

### RT-PCR

RT-PCR experiments were performed using the methods and RNA samples of Matt and Umen (2018). cDNA was prepared from two somatic and two gonidial samples described in Matt and Umen (2018) with 5 μg total RNA following the manufacturer’s protocol for Thermoscript (Invitrogen, Carlsbad, CA) using a 10:1 mixture of oligo dT and random hexamer for priming and the following cDNA synthesis reaction temperatures: 25°C 10’, 42°C 10’, 50°C 20’, 55°C 20’, 60°C 20’, 85°C 5’. After cDNA synthesis reactions were treated with RNaseH and diluted 1:10 with 10mM Tris pH 8.0, 1mM EDTA (TE) and stored at −20°C. PCR amplification reactions (20μL total per reaction) contained 1X Taq polymerase buffer, 0.2mM dNTPs, 2% DMSO, 1 μl purified recombinant Taq polymerase, 0.5 mM each primer (Supplemental Table 1) and 1 μl cDNA template. Samples were amplified in a peqSTAR thermocycler as follows 1) 94 °C for 2 min; 2) 35 cycles of 94 °C for 25 seconds, 55 °C for 30 sec, and 72 °C for 30 sec; 3) 72 °C for 10 min. After thermocycling, samples were separated on a 2% agarose gel containing ethidium bromide and imaged using a UV transilluminator and CCD camera.

## RESULTS

### Identifying Volvox genes with multiple transcripts

Raw data from a previously published study of purified Volvox gonidial and somatic cells was re-analyzed for cell-type-specific alternative transcript isoforms (CTSAI) using the Ballgown software package with additional filtering and curation as described below. We first verified that each set of replicates for somatic and gonidial samples showed correlated expression values and that differential cell type expression was comparable to that in the prior study of Matt and Umen, 2018 (Supplemental Figs. S2 and S3). CTSAI can only occur if two or more transcript isoforms are produced from the same gene. Out of the 11095 Volvox genes that had average expression across all samples >1 FPKM, 9860 had only one transcript isoform and were not analyzed further (Fig. 1). 1235 remaining genes had multiple predicted transcripts and were candidates for CTSAI. 974 of these genes (78%) had just two predicted isoforms and were analyzed further while those with three or more isoforms were not.

**Fig. 1:**
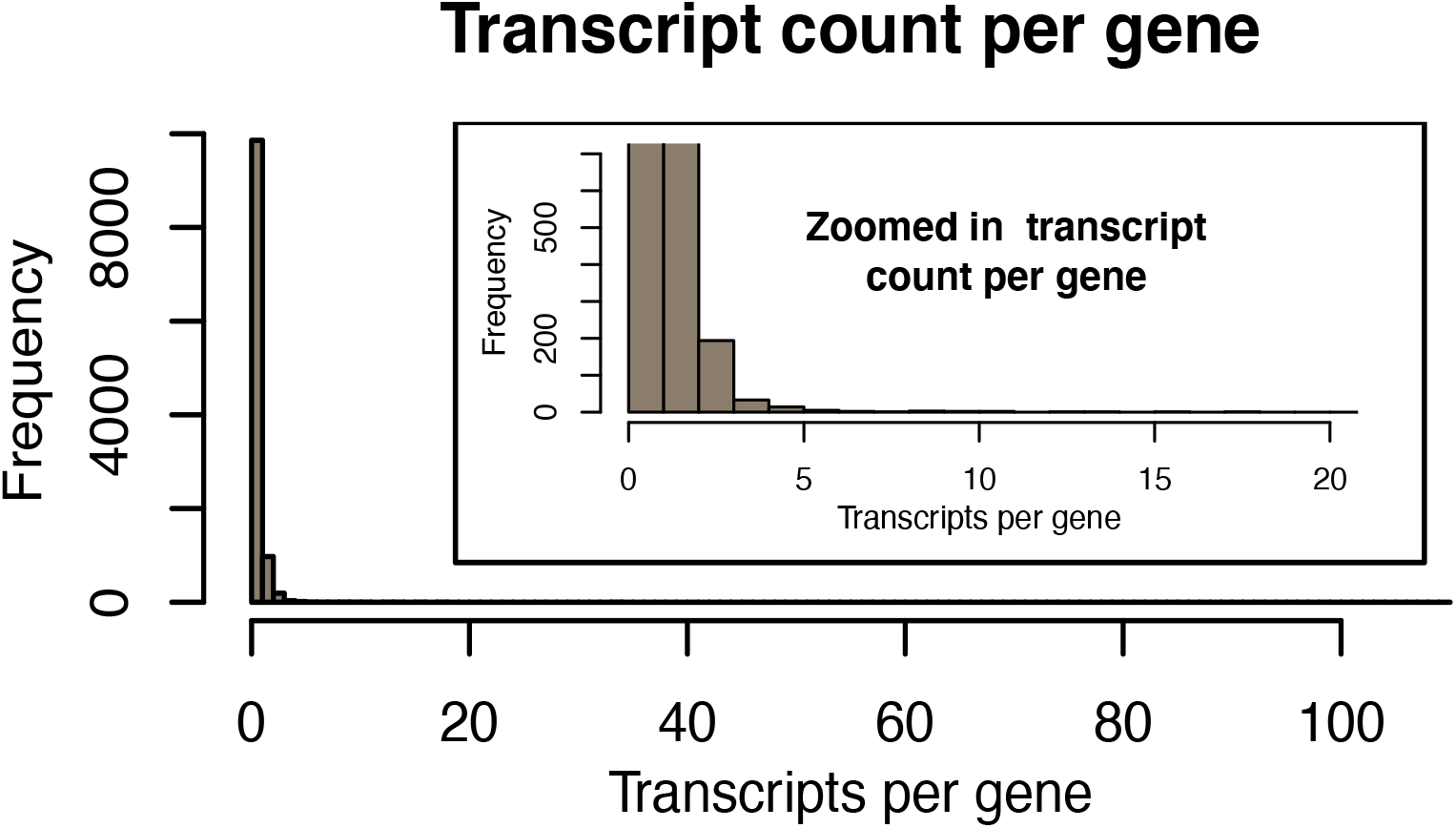
Histogram of transcript counts per gene. 9860 genes have one isoform, 974 have two isoforms, while the remaining 261 genes have three or more isoforms. The largest number of isoforms per gene is 34. The inset in the top right corner is the histogram zoomed up at lower numbers of isoforms per gene.

To quantify cell type expression bias of transcript isoforms we first computed the expression ratios between cell types for each isoform as (expression level in soma)/(expression level in gonidia). Large ratios (>>1) indicated bias towards soma, while small ratios (<<1) indicated bias towards gonidia. CTSAI are present when the computed cell type ratios of each transcript isoform significantly differ from each other.

We focused on genes whose transcript isoforms showed opposite cell type biases. We therefore considered cases where one transcript isoform was somatic-biased (s/g ratio>2) while the other isoform was gonidial-biased (s/g ratio <1/2) with p-values for differential expression < 0.1. Additionally, we required the overall cell type bias, defined as [isoform 1 (s/g)/ isoform 2 (s/g)], to be greater than 8. 60 Genes passed these two criteria and were assigned as candidates for CTSAI with an aggregate p-value < 0.01 since the individual p-values can be multiplied (Fig. 2a-c). Results using alternative filter cutoffs with higher or lower stringencies are in Supplemental Figures S4 and S5. Expression data for all 60 genes are in Supplemental Table 2.

**Fig. 2:**
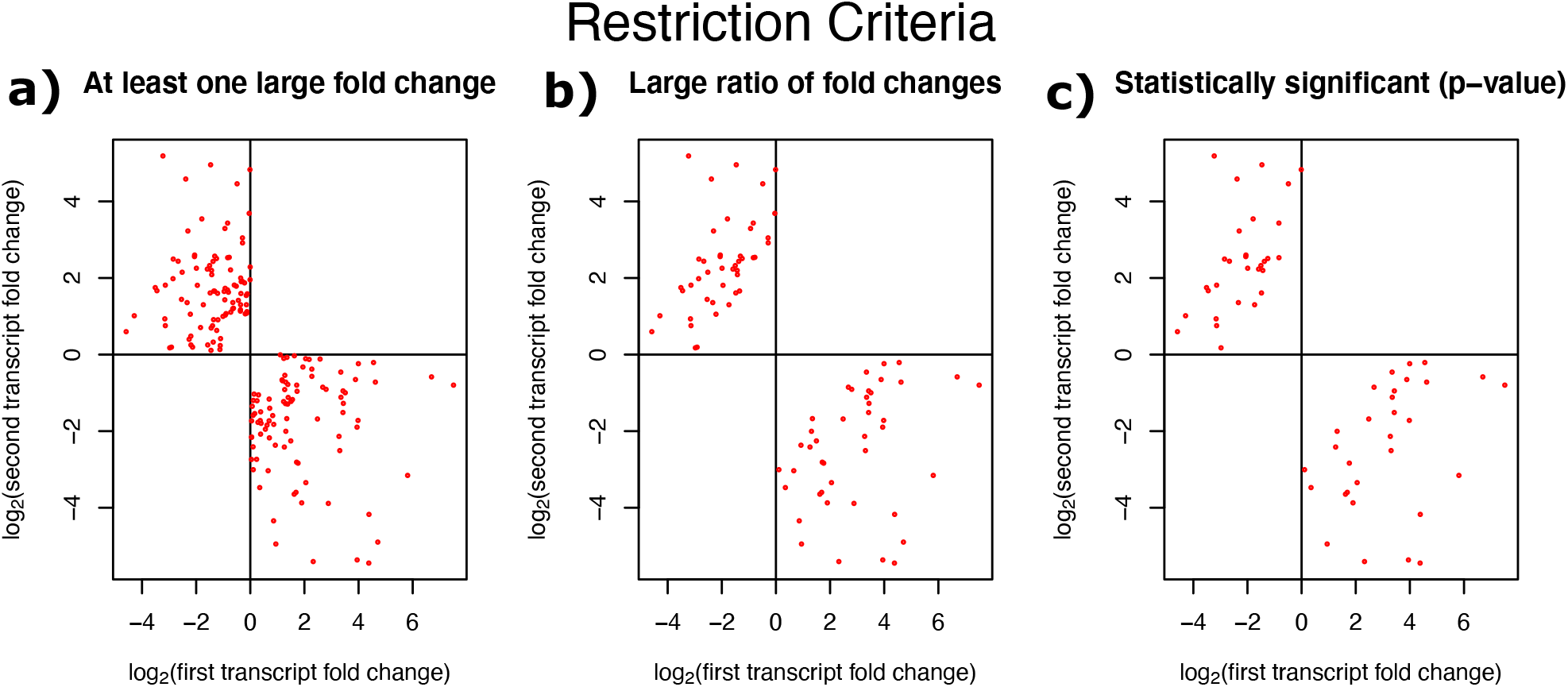
Criteria for identifying CTSAI. Log expression ratio (er = expression in soma over gonidia) for second vs. first transcript in genes with two transcripts and opposite cell-type bias.(a) Implemented restriction of at least one soma-gonidial expression ratio greater than 2, yielding 187 possible genes. (b) Implemented criterion that the ratio of expression ratios is greater than 8, yielding 86 possible genes. Genes far from the diagonal are candidates for CTSAS. (c) Implemented criterion for p-values (for each transcript) less than 0.1, yielding 60 candidate genes.

### Phytozome predicted alternative transcripts

To supplement our analysis using Ballgown software which relied on *de novo* transcript assembly, we repeated the same analysis above using alternative transcripts predicted from the version 2.1 genome assembly of *V. carteri* hosted on Phytozome [21, 26, 27]. Out of the 9750 genes that had average expression across all samples >1 FPKM, 9297 only had one isoform and were removed from the analysis. Of the remaining 453 genes, which had multiple isoforms, 415 had only two isoforms, and were analyzed further.

We proceeded to use the same filtering criteria as were used for Ballgown-predicted transcripts – opposite expression ratios between cell types for the two isoforms, at least one expression ratio in the isoform pair being greater than 2 or less than 0.5, an overall cell-type-bias of the two isoforms greater than 8, and p-values for differential expression of each individual isoform less than 0.1. This filtering resulted in identification of 20 candidate genes from Phytozome for CTSAI, 13 of which were predicted by Ballgown and 7 of which were unique.

### Manual curation

Each of the 67 combined total CTSAI candidate loci predicted by Ballgown and Phytozome were visualized on the Phytozome browser [26, 27] along with transcriptome coverage in each cell type replicate [17] to assess the results of automated alternative transcript predictions and the evidence for presence of CTSAI (Supplementary Table 3). We further filtered candidates following some general criteria: a) overall read mapping of both isoforms was sufficiently dense to support transcript structures [RD—read density]; b) reads mapping to differential portions of each transcript supported a significant difference in isoform abundance by cell type and not just overall transcript abundance between cell types [REL – relative expression level]. In doing so we identified 52 false positives that were excluded for one or more reasons as described above while retaining 15 genes as strong candidates for having CTSAI – 12 were in the Ballgown analysis and 4 were in the Phytozome analysis, with one gene in common between the two sets (Supplemental Table 3).

Of the 12 genes the genes that were identified as strong candidates for CTSAI in the Ballgown analysis, 10 were not in the set of candidates for CTSAI from the Phytozome analysis because Phytozome did not previously identify alternate isoforms for them, or predicted that those genes have more than two isoforms. Thus, we conclude from this that Ballgown performed well at identifying previously unknown alternative isoforms and determining which genes were strong CTSAI candidates.

### Convergent transcripts

Among our 60 candidate CTSAI loci, there were 4 clear cases where regions of overlapping convergent transcription from distinct adjacent loci were mis-identified by Ballgown as alternative isoforms (letter code in Supplemental Table 2 = CT). The Phytozome IDs that were assigned to these convergent genes by Ballgown were Vocar20012504m.g (version 2.0), Vocar.0004s0224, Vocar.0042s0098, and Vocar.0006s0317 (version 2.1). Although the available transcriptome reads used by Ballgown did not contain strand information, during manual curation the direction of transcription could be inferred for each transcript from a combination of Phytozome gene model predictions, annotations, ESTs, and/or polarity of consensus splice site sequences in each model. Notably, in most instances of convergent transcription that were originally mis-identified as CTSAI, the cell type specificity was correctly estimated, meaning that a different member of the convergent pair dominated in each of the two cell types. These convergently transcribed overlapping gene pairs might, therefore, have an antagonistic regulatory relationship that contributes to their cell type expression specificity (see Discussion). Examples of cell type specific convergent transcript pairs are shown in Fig. S7a and S7b.

### Experimental validation of alternative splicing

We selected three genes showing CTSAI for further validation: Vocar.0001s1758, Vocar.0011s0285, and Vocar.0018s0107 (Fig. 3). Vocar.0001s1758 encodes a predicted UDP-glucose pyrophosphorylase and is required for UDP-glucose synthesis, a critical substrate for protein glycosylation. Vocar.0011s0285 has a universal stress protein (USP) domain (PFAM 00582) and its Chlamydomonas ortholog, Cre03.g211521, is a cilia-associated protein, FAP165 [29]. Vocar.0018s0107 is a predicted flavin-dependent oxido-reductase with three flavodoxin binding domains (PFAM 00258) followed by a FAD domain (PFAM 00667). To confirm the bioinformatic results, we PCR amplified Volvox cDNA produced from one of the replicate RNA samples [17] with primers that were isoform specific. We created 3 primers for each tested gene, a shared primer which would bind to both transcript isoforms, and a specific primer for each isoform that could pair with the shared primer. We used isoform specific primer sets alone or combined in single PCR reactions from each cell type, and assessed relative band intensities. For all three genes we found a qualitative match to the cell type splicing patterns predicted using transcriptome data (Fig. 4).

**Fig. 3:**
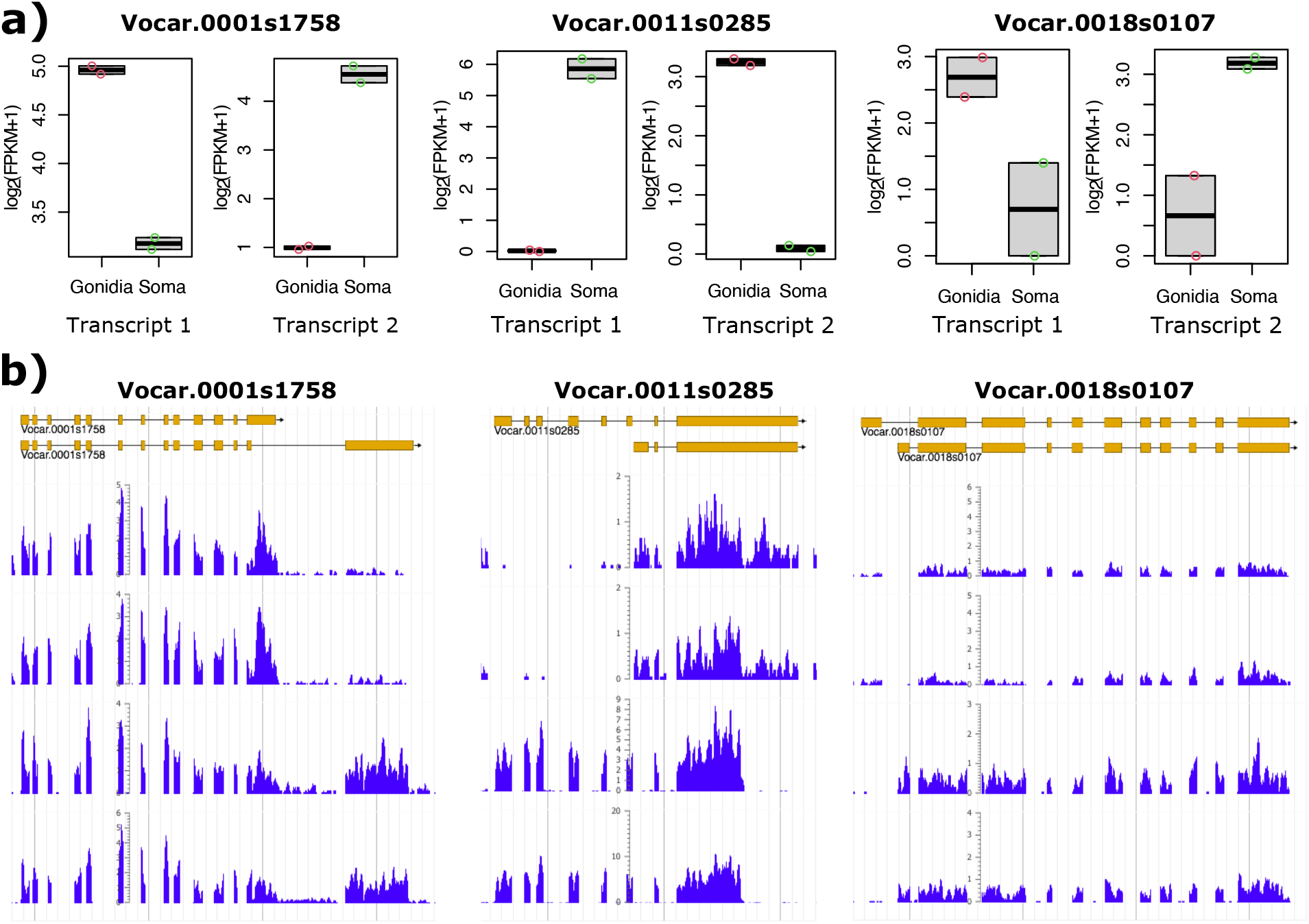
**a)** CTSAI expression estimates by cell type for Vocar.0001s1758, Vocar.0011s0285, and Vocar.0018s0107 genes. Colored circles show replicate measurements; dark black line is the mean. Transcript 1 and Transcript 2 for each locus are indicated in panel b [please label]. **b)** Alternative-splicing generates the CTSAI pattern for the Vocar.0001s1758, Vocar.0011s0285, and Vocar.0018s0107 genes. Browser screen shot from Phytozome with exons from alternative transcript isoforms shown as tan boxes at the top. Below the transcript models are replicate tracks showing RNA-seq read coverage from each cell type in blue.

**Fig. 4:**
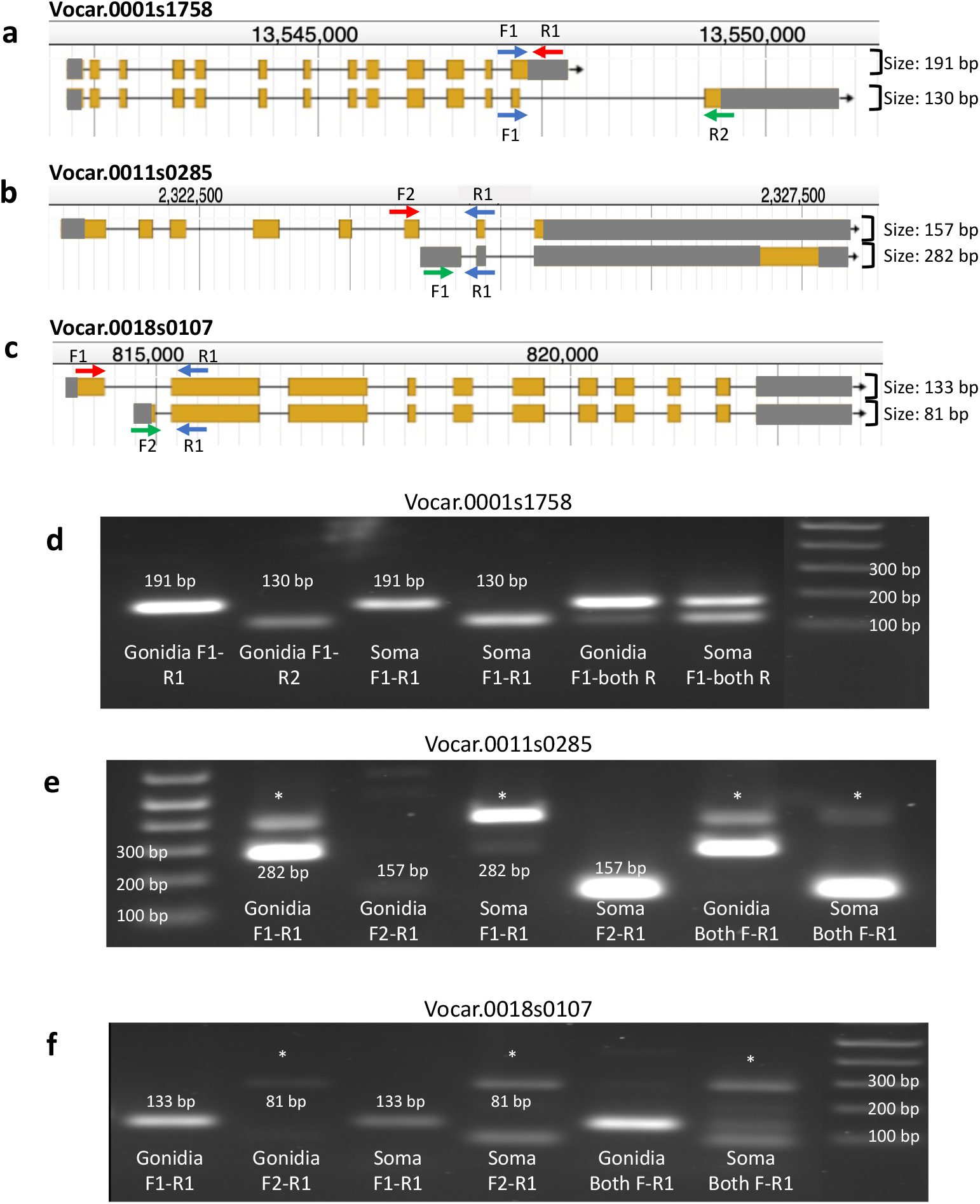
**a-c)** Graphic displaying binding sites and amplicon lengths for RT-PCR primer sets used for Vocar.0001s1758, Vocar.0011s0285 and Vocar.0018s0107, respectively. Arrows show primer direction with the common primer in blue and the isoform specific primers in red and green. Predicted product lengths are also indicated. **d-f)** Gel electrophoresis images from amplification reactions using primers in panels a-c. Lanes are marked with primer combinations and cell types. The end lanes have DNA size markers. Asterisks indicate non-specific amplification products which did not derive from the target locus, since the primers generated for these experiments were unique sequences found in each gene.

The gonidial form of Vocar.0001s1758 has 13 exons, while the somatic form has one extra exon at the 3’ end which is joined to an internal 5’ splice site within exon 13 of the gonidial isoform (Fig. 3). We compared transcript structures and the protein sequences encoded by Vocar.0001s1758 isoforms to that of the orthologous gene in *Chlamydomonas reinhardtii* (Chlamydomonas), Cre04.g229700, which has a single isoform that can be used as a potential proxy for the ancestral transcript structure and protein sequence (Supplemental Figure S8). Cre04.g229700 has an overall similar structure to that of the Volvox transcripts, but with two extra introns that are not present in either Volvox isoform. At its 3’ end the Chlamydomonas transcript matches the structure of the gonidial isoform in Volvox which is most likely the ancestral gene structure, with the somatic form being derived. Both predicted transcript isoforms retained the coding region for the catalytic domain, and differ in their C-terminal domains. UDP-glucose pyrophosphorylases show highest activity as monomers, but can also oligomerize to a form that is less active [30]. The solved crystal structure and additional conformational modeling of *Arabidopsis thaliana* UDP-glucose pyrophosphorylase paralog At3g03250 revealed a potential explanation for the impact of dimerization on activity: In dimeric form the C-terminal portion of each subunit partly occludes the active site in the N-terminal domain [31]. Thus, altering the C-terminal region in the two Volvox isoforms has the potential to affect dimerization efficiency and/or activity of the dimeric form, either of which could cause somatic and gonidial isoforms of Vocar.0001s1758 to have different activities and/or allosteric regulatory responses. While there is no direct evidence of differential activity of UDP-glucose pyrophosphorylase in *V. carteri* cell types, it is striking that nearly every enzyme in the starch-sugar nucleotide synthesis pathway in *V. carteri* has a transcript which is subject to cell type regulation with high expression in gonidia and low expression in soma—except for Vocar.0001s1758 which shows no difference in overall transcript abundance between cell types [17]. Our data showing that this gene produces cell type specific alternative isoforms of UDP-glucose pyrophosphorylase provides a rationale for why cell type expression level control is not observed for this member of the pathway, as regulation of activity is achieved post-transcriptionally by cell type specific alternative splicing.

The predicted somatic isoform of Vocar.0011s0285 encodes a protein that is orthologous to the Chlamydomonas gene Cre03.g211521 (Fig. S8), while the gonidial isoform uses an internal transcription start site and excludes the first six exons found in the somatic isoform (Fig 4b). Instead, the gonidial transcript begins with a unique alternative 5’ exon followed by a short penultimate exon and final exon, both of which are shared with the somatic isoform. The Cre03.g211521 predicted protein was found in the ciliary proteome and designated FAP165, but its function has not been characterized [29]. Its only domain (also shared by the somatic isoform of Vocar.0011s0285) is PFAM 00582/IPR006016 that is designated a universal stress protein domain [32], though its specific function in stress survival is unclear. Interestingly, several other chlorophyte algae, mostly within the Chlamydomonadales, have homologs of Cre03.g211521 with similarity extending outside of the conserved N-terminal PFAM 00582/IPR006016 domain. The closest matches of this protein outside of related green algae are in prokaryotes, and only in the conserved PFAM 00582/IPR006016 domain. It is thus possible that homologs of Cre03.g211521/Vocar.0011s0285 originated as a horizontal gene transfer event from prokaryotes into one or more subgroups of Chlorophyceaean algae. How the function of this protein domain was incorporated into cilia is an intriguing question. Our finding that only the somatic isoform of Vocar.0011s0285 can produce a FAP165-related protein supports the idea that this protein functions in the cilium of volvocine algae since most cilia-related proteins are not expressed at all in *V. carteri* gonidial cells [Matt and Umen 2017]. Indeed, the gonidial isoform of Vocar.0011s0285 appears to be a non-coding transcript produced from an internal initiation site whose only significant open reading frame would encode a short peptide that is unrelated to the somatic protein isoform. Whether the gonidial transcript has a function is unknown, as is the reason why this transcript came under cell type regulation by alternative transcript isoforms and not through cell type expression as is the case for most of the cilia-related genes in Volvox [17].

The two isoforms of Vocar.0018s0107 have different transcription start sites and first exons, but share all remaining exons in common and encode predicted proteins that differ in their N-termini (Figure 4c, Figure S8). The two predicted Volvox proteins are equally similar to the Chlamydomonas ortholog encoded at locus Cre16.g683550 which does not align with either Volvox protein in the N-terminal region. Interestingly, the predicted localizations for the two Volvox isoforms by the Predalgo algorithm are different, with the somatic isoform predicted to be imported to the chloroplast and the gonidial isoform predicted to be excreted [33]. The Chlamydomonas ortholog is also predicted to enter the secretory pathway, and this may be the ancestral localization for the protein. Vocar.0018s0107 is a predicted oxido-reductase with three flavodoxin domains (IPR008254) followed by a ferredoxin reductase-type FAD binding domain (IPR017927). These flavodoxin domains often serve as redox reaction activators [34]. Interestingly this domain architecture is shared with a family of predicted proteins in many other green algae and in the phytopthera clade of downy mildew plant pathogens, but to our knowledge has not been functionally characterized in any species. Its alternative splicing in Volvox may influence its function via differential subcellular localization in somatic versus gonidial cells, but its specific biochemical function(s) and substrates remain to be determined.

### RNA Binding Proteins as Possible Mechanism for CTSAI

The identification of cell type alternative splicing in Volvox implies that there is some cell type specificity to its RNA processing machinery. One broad class of regulatory proteins that can influence alternative RNA processing are those with RNA binding domains [35]. We therefore identified 126 predicted RNA binding domain proteins in Volvox based on their automated domain annotations, including RNA recognition motifs (PFAM 00076), KH domains (PFAM 00013), G patch domains (PFAM 01585), RNA helicase domains (PFAM 00270/1) and several others (Supplemental Table 4). Proteins with these domains whose expression is cell type specific are candidate mediators of cell type specific alternative splicing. We queried the transcripts for these 126 predicted RNA binding proteins for their cell type expression patterns as determined in our earlier study [17] and found that most were expressed at similar levels in both cell types (<2 fold cell type expression ratio) or showed modest cell type bias (2-5 fold cell type expression ratio). However, 40 predicted RNA binding protein coding genes had transcripts that were gonidial specific and four were somatic specific (>5 fold cell type expression ratio) (Supplemental Table T4). These 44 proteins are candidate regulators of cell type alternative splicing.

## DISCUSSION

Our analyses have for the first time characterized cell type specific transcript isoform usage in the multicellular alga *Volvox carteri*. Using both *de novo* transcript assembly with Ballgown software and previously predicted transcript isoforms from Phytozome, we identified candidate genes with cell-type alternative splicing. Based on bioinformatics predictions alone, the frequency of genes with two alternative isoforms was relatively high, with around 12-13% of all loci predicted to have two or more alternative isoforms. 19.7% of expressed genes in *Chlamydomonas reinhardtii* and about 42% of intron containing genes in *Arabidopsis thaliana* are alternatively spliced [36, 37], so the overall frequency of alternative transcript isoforms in *V. carteri* appears to be lower; but the differences in these estimates could also be due to the amount of data available and the methods used to identify alternative transcripts. Moreover, the sexual cycle in Volvox involves modified embryogenesis and differentiation of additional cell types including eggs, sperm and somatic sexual cells, and these might also partly rely on alternative splicing of genes only present in the sexual phase of the life cycle that was not queried here [21]. Only 67 of the 974 loci in Volvox that had two predicted alternative isoforms also showed cell type bias for expression of those isoforms. Moreover, of the 67 loci with high confidence and strong cell type isoform expression bias, most were false positives. In some cases, the transcripts that were predicted by Ballgown were not supported by our expression data, or the expression data was too sparse to adequately support the existence of an alternative transcript. A few false positives were due to adjacent genes being so close together that Ballgown fused them into erroneous transcript predictions. However, many false positives were also found among the Phytozome predictions, and these errors became apparent when visually inspecting transcriptome read mapping data (see Supplemental Figure S6c). For future work it would be helpful to use Iso-seq type methods [38] to capture full length transcripts and/or use sequencing methods that preserve transcript strand information to help reduce false positives.

Nonetheless, we did identify fifteen genes that passed manual inspection, and three of these were validated experimentally. In summary, cell type specific alternative transcript isoforms seem to represent only a minor component of cell differentiation in Volvox where biased cell type expression was found previously for thousands of genes and appears to be the predominant means of cell differentiation in this simple multicellular species. Overall, our data imply that CTSAI operates as a component of *V. carteri* cellular differentiation, and this simple multicellular organism could be a model for how alternative isoforms emerge and co-evolve with cell type specialization.

To determine the degree of conservation of alternative splicing in *V. carteri* and *C. reinhardtii*, we compared the frequency of alternative splicing in both species using a subset of 10447 predicted 1:1 orthologs between the two species identified based on mutual best hit similarity searching [17]. The merged Ballgown and Phytozome alternative transcript data set of genes with two isoforms (1978 genes total without duplicates) had 1520 in the 1:1 homolog list. Searching data from a recent study in Chlamydomonas that identified 2182 alternatively spliced genes Pandey et al. (2020), 1441 were found in the 1:1 homolog dataset 273 of which were shared with Volvox. We repeated the same analysis using the Chlamydomonas genes with multiple transcripts predicted by Phytozome v5.5, which contained 1531 genes, 790 of which were in the 1:1 homolog dataset, and 161 of which had a Volvox gene with alternative isoforms. However, among the set of CTSAI candidate genes, only one had alternative isoforms in Chlamydomonas. This suggests that new cases of CTSAI in Volvox mostly evolved from genes that previously did not have cell type specific alternative splicing patterns. However, we cannot rule that some *C. reinhardtii* genes whose *V. carteri* counterparts show CTSAI may have undetected alternative splicing, or that CTSAI was present ancestrally and lost in *C. reinhardtii*. Overall, there appears to be minimal conservation of alternative splicing in volvocine algae, suggesting that this feature of gene expression is labile, a property that could facilitate the evolution of cell-type specific alternative transcript functions.

During curation of automated prediction of CTSAS/CTSAI we also encountered cases of cell-type specific convergent transcription in which expression for a different member of the convergent pair dominates in each cell type. The overlapping 3’UTRs in these examples may be a mechanism for gene regulation that functions by antagonizing production, translation or mRNA stability of a convergent partner [39, 40]. A focused search for more examples of convergent transcripts with opposite cell type expression patterns could be used to determine how widespread is this phenomenon that was uncovered coincidentally in our study of CSTAS.

RNA binding proteins are potential mediators of CTSAS. Here we identified 126 predicted RNA binding proteins from Volvox, and a subset of 40 that showed cell type specific expression bias. These 40 may be potentiators for the evolution of CTSAS which could occur through direct binding to pre-mRNA sequences that influence splice site or transcription start/termination site usage. Moreover, there exist other possible means of regulating transcript processing such as riboswitches interacting with metabolites or small molecules [41], through small RNA (sRNA) or microRNA (miRNA) pathways [42], or by modulation of transcription start/termination sites via DNA binding transcription factors. The prevalence and relative importance of these different mechanisms is not known for Volvox or other organisms, and is an ongoing area of research [22, 23, 41].

## Notes

### Competing Interest Statement

The authors have declared no competing interest.

