## Supplemental Figures and Tables for "Identification of Cell-Type Specific Alternative Transcripts in the Multicellular *Alga Volvox carteri*"

**SUPPLEMENTAL MATERIAL**

*Supplemental Figures*

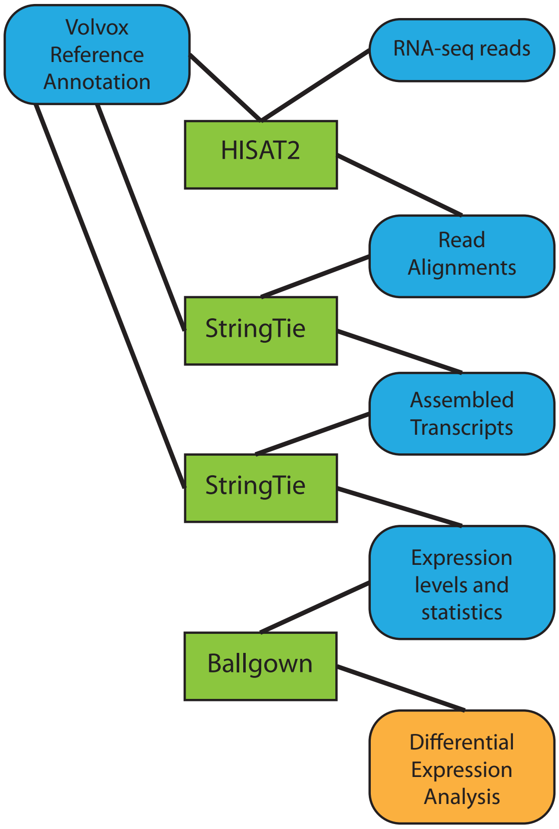


**Figure S1:** Schematic of the bioinformatic analysis using the HISAT2, String Tie, and Ballgown software packages [Pertea et al., 2016]. The workflow is from top to bottom.

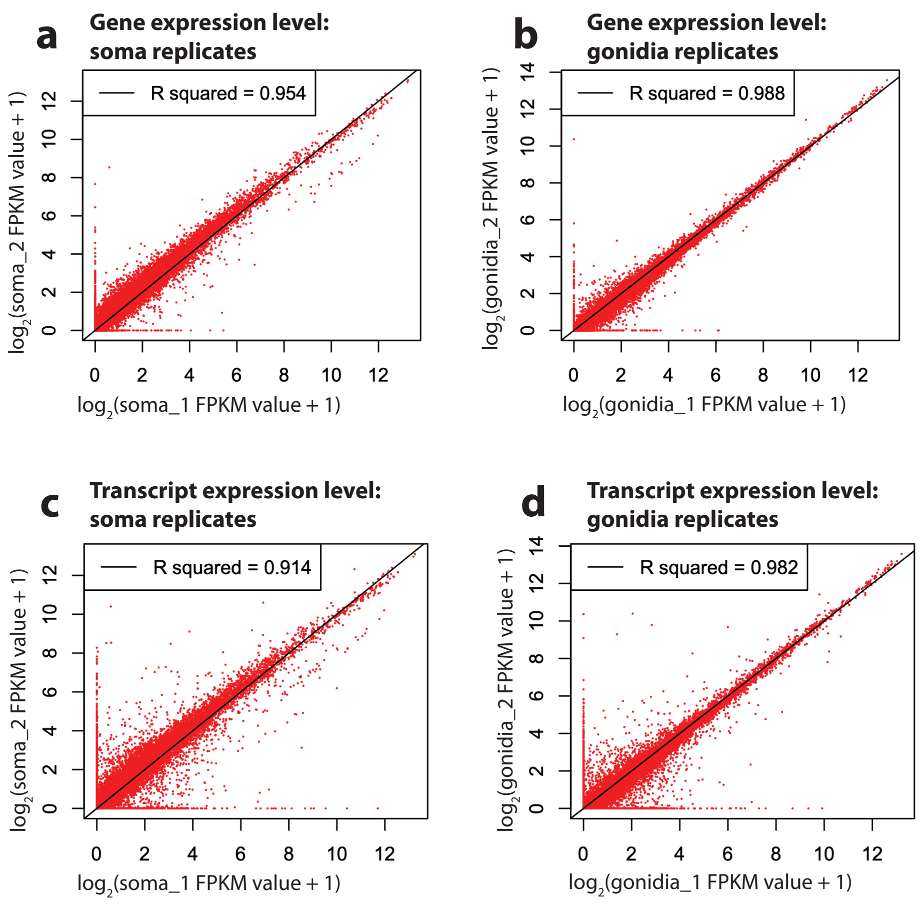


**Fig. S2**. (**a,b**) Comparison of gene expression levels for somatic and gonidial replicates. Red dots are individual genes. (**c,d**) Comparison of transcript expression levels for somatic and gonidial replicates. Red dots are individual transcripts. I added 1 to the FPKM values because the log is ill-defined when the FPKM is zero.

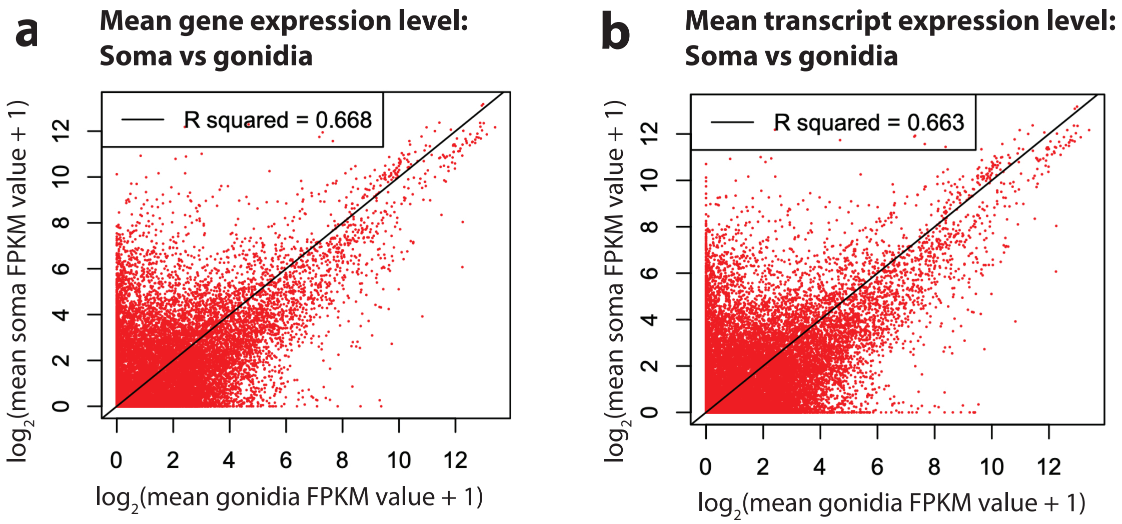


**Fig. S3:** (**a**) Gene expression levels (log FPKM values) show low correlation between somatic vs. gonidial cells. Red dots are individual genes. (**b**) Transcript expression levels show low correlation between somatic vs. gonidial cells. Red dots are individual transcripts.

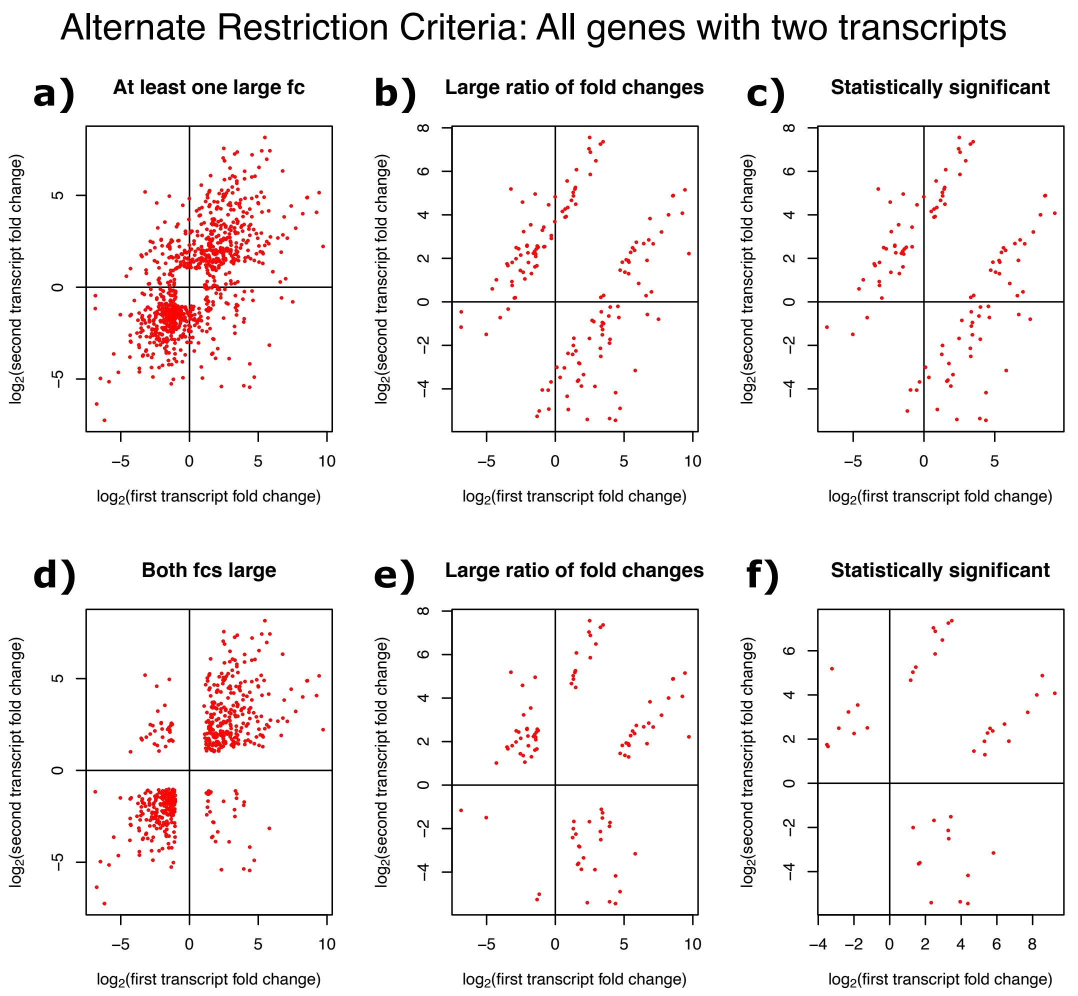


**Fig S4:** Log fold change (fc = expression in soma over gonidia) for second vs. first transcript in all genes with two transcripts. (**a**) Implemented restriction of at least one soma-gonidial expression ratio greater than 2 (or less than 1/2), yielding 863 possible genes. (**b**) Implemented criterion that the ratio of expression ratios is greater than 8, yielding 145 possible genes along with criterion (a). (**c**) Implemented criterion for p-values (for each transcript) less than 0.1, yielding 110 candidate genes along with criteria (a) and (b). (**e**) Implemented restriction of both soma-gonidial expression ratios greater than 2 (and/or less than 1/2), yielding 552 possible genes. (**f**) Implemented criterion that the ratio of expression ratios is greater than 8, yielding 96 possible genes along with criterion (e). (**g**) Implemented criterion for p-values (for each transcript) less than 0.1, yielding 41 candidate genes along with criteria (e) and (f). Genes far from the diagonal are candidates for CTSAS.

### Alternate Restriction Criteria: Opposite Fold Changes

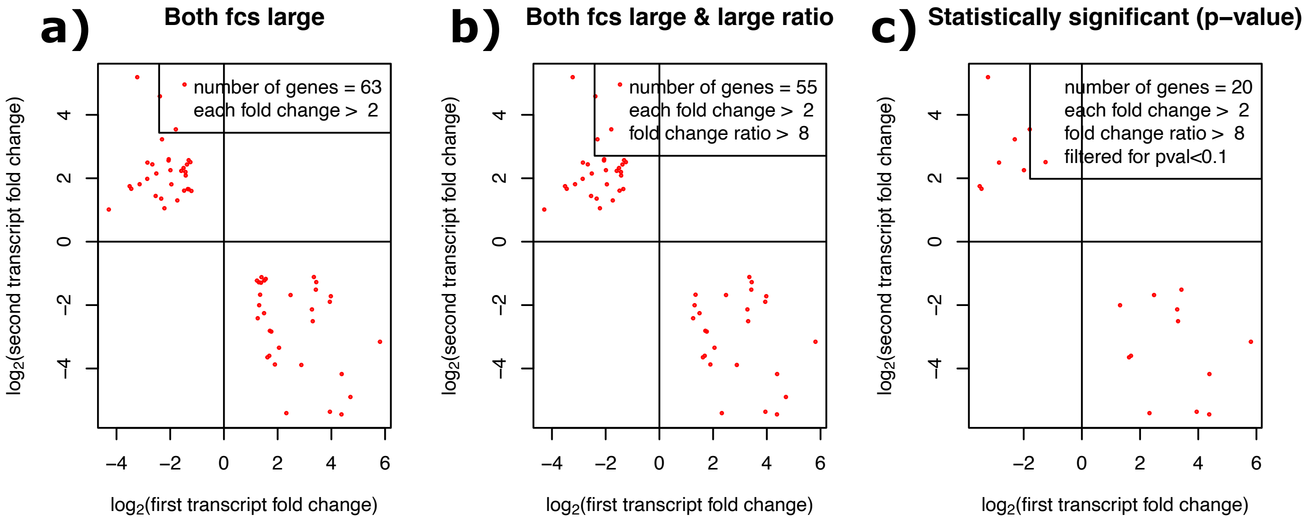


**Fig. S5**: Log fold change (fc = expression in soma over gonidia) for second vs. first transcript in genes with two transcripts and opposite cell-type bias. (**a**) Implemented restriction of both soma-gonidial expression ratio greater than 2 (or less than 1/2), yielding 63 possible genes. (**b**) Implemented criterion that the ratio of expression ratios is greater than 8, yielding 55 possible genes along with criterion (a). (**c**) Implemented criterion for p-values (for each transcript) less than 0.1, yielding 20 candidate genes along with criteria (a) and (b). Genes far from the diagonal are candidates for CTSAS.

### Manual Curation Examples

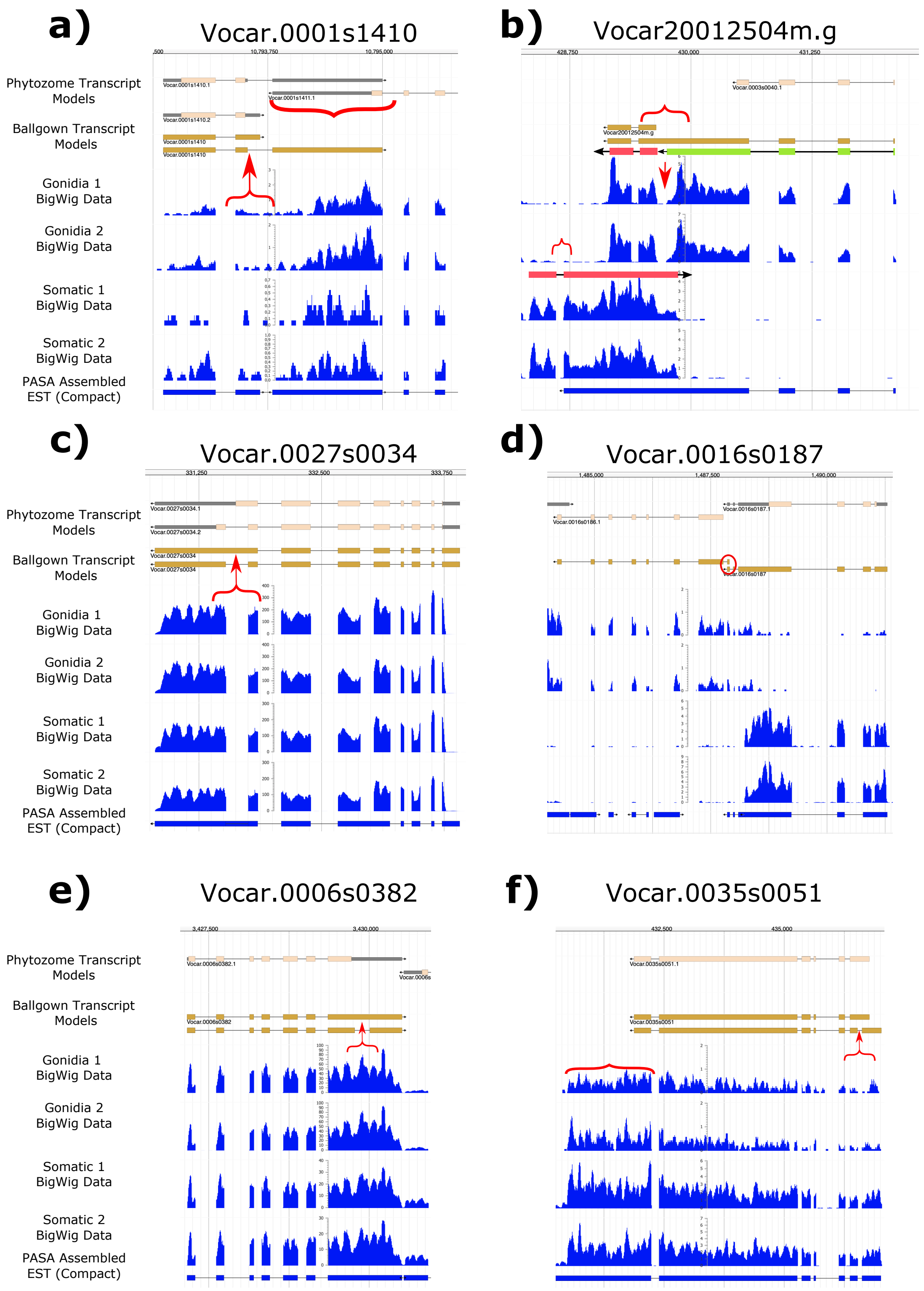

**Fig. S6:** Example genes that were removed from the analysis during manual curation, with their respective ballgown transcript models, Phytozome transcript models (if present), and read expression data. **(a)** Vocar.0001s1410 was removed due to low coverage in the alternatively-spliced eon marked by the red bracket and arrow. The domain marked with the large red bracket is an overlap region between two adjacent genes, so the high expression in the four samples in this region may not be due to the gene Vocar.0001s1410, but rather its neighbor Vocar.0001s1411. **(b)** Vocar.20012504m.g was removed due to little little evidence that the large, central exon of the central transcript was present due to the gap in expression data marked by the red arrow, the extended expression data to the left of the model implying that the model lacked some pieces, and the fact that the expression data, as presented, implied the existence of different transcript models. Looking carefully at the clear splice sites and the areas where the expression tapers, alternative transcript models were proposed: the green model is a separate gene, and the two red models are convergent transcripts over the same locus. **(c)** Vocar.0027s0034 was removed because of the clear lack of evidence of the existence of an alternative splicing region in the 3’ end exon, marked by the red bracket. **(d)** Vocar.0016s0187 was removed because ballgown added an additional exon to the top transcript (circled in red) and misidentified two neighboring genes as two transcripts of the same gene because of the created overlap created. **(e)** Vocar.0006s0382 was removed because of the clear lack of an alternative splicing site proposed by ballgown in the 3’ end exon marked by the red bracket and arrow. **(f)** Vocar.0035s0051 was removed because there the expression data in Gonidia was two weak in order to determine if the alternative splicing site at the 5’ end of the transcripts was present or not. These models also seem to be lacking an extension to the 3’ end, as evinced by the clear expression data that extends beyond the 3’ end.

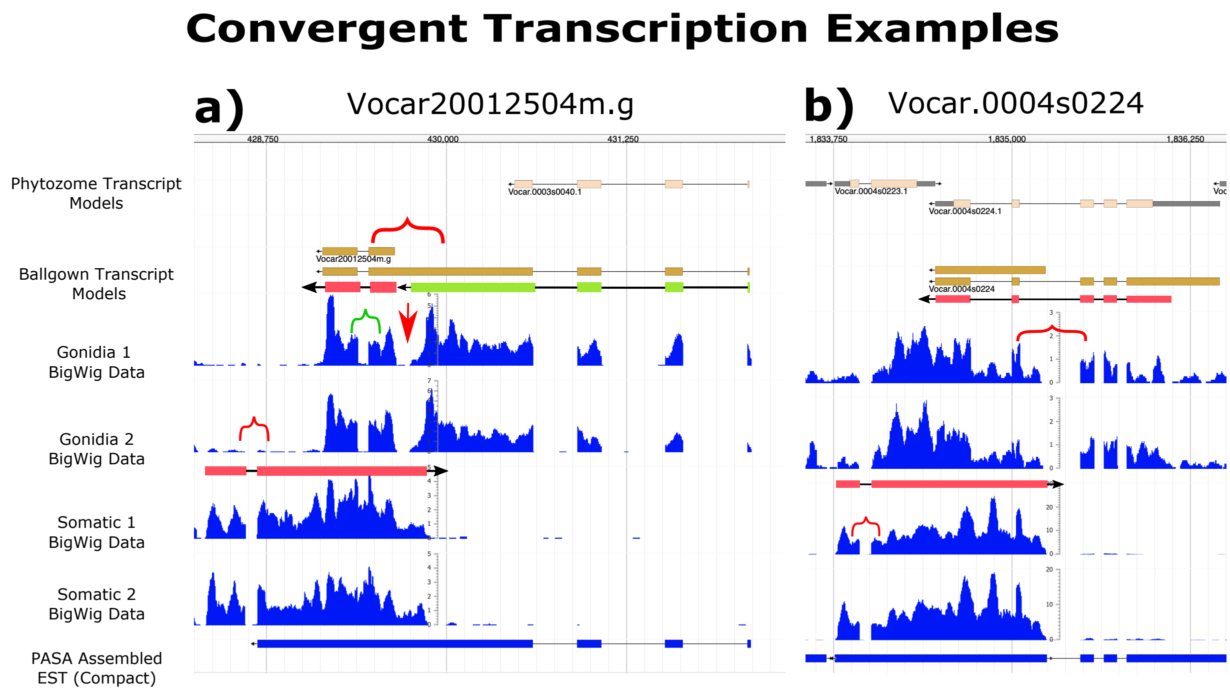

**Figure S7:** Examples of convergent transcription in *V. carteri*. In these cases, one gene has a transcript going in the forward-sense direction, while the other transcript goes in the anti-sense direction. Since both Vocar.20012504m.g and Vocar.0004s0224 had misconstructed gene and transcript models, new models were made in red. Since convergent transcription is not CTSAS by our definition, both these cases were removed from the analysis, although they are worth mentioning as an alternative mechanism of producing multiple cell-type specific transcripts. (a) Vocar20012504m.g displayed with its Ballgown models, its expression data, and the hand-made models based on the expression data. In the gonidial samples, a tapering of in gene expression can be seen on either side of the gap identified with the red arrow, which implies that the units of expression to the left and to the right of this gap are in fact separate transcripts, disagreeing with the models Ballgown proposed. Then, checking the steep drops in expression on either side of the gaps in expression in the gonidial samples, we found the GTA/GTG motif (5' splice site) on the right side of each gap, and the CAG/TAG motif (3' splice site) on the left of each gap, implying that the two transcripts in the gonidial samples go in the antisense direction. Meanwhile, the splice site in the somatic expression data (marked with the small red bracket) had the GTA/GTG motif (5' splice site) on the left side of the gap, and the CAG/TAG motif (3' splice site) on the right, implying that that transcript went in the sense direction. Overall, the two transcripts that overlapped were taken to be the alternative, convergent transcripts of the Vocar.20012504m.g gene (marked in red), while the third transcript (in green) was thought to be a separate gene. (b) Vocar.0004s0224 is another case of misidentified convergent transcription. Using the same method of identifying the motifs flanking each potential splice site, we determined that the gonidial transcript goes in the antisense direction, while the somatic transcript goes in the sense direction (in red). It is possible that this is a case where Phytozome misidentified two convergent transcripts as two different genes, as evinced by the two Phytozome gene models at the top of the panel which overlap the expression data.

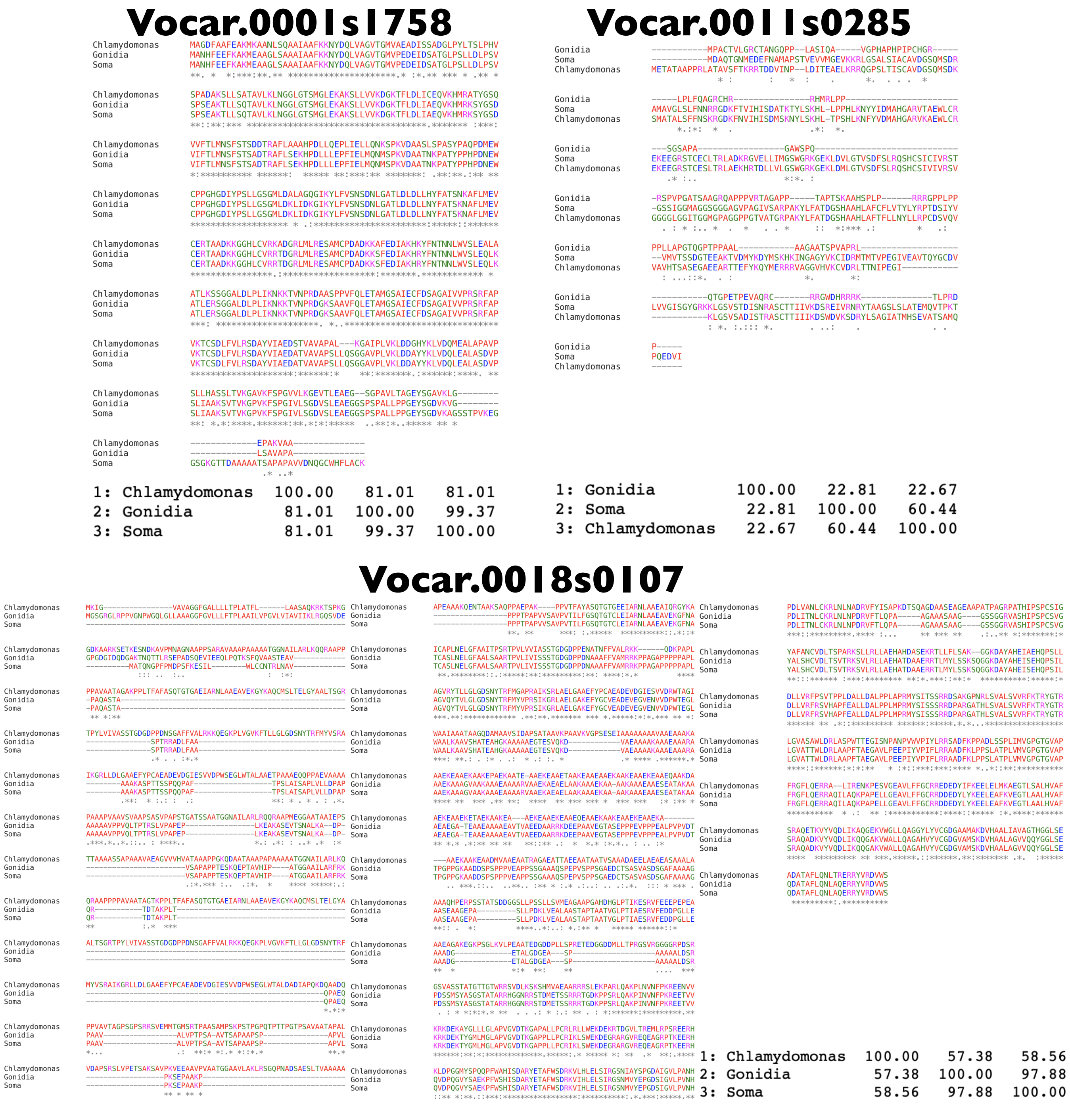


**Figure S8:** Three-Way alignment of the two isoforms (listed by the cell-type in which they are most expressed) and their Chlamydomonas ortholog. Vocar.0001s1758's isoforms are very similar to the original Chlamydomonas sequence and have similar lengths. The 3' end of the somatic isoform is different in length from that of the gonidial isoform and Chlamydomonas, while the gonidial and Chlamydomonas isoform are similar in length but differ by a few amino acids. Thus the gonidial isoform may be the more ancestral transcript. Vocar.0011s0285's isoforms are very different from the Chlamydomonas homolog (only ~22-23% similarity). The exon structure of the gonidial isoform is very different from that of the other two isoforms. It is probable that the Volvox sequence diverged from Chlamydomonas and then became two separate isoforms. Vocar.0018s0107's isoforms differ at the 5' end, but have mostly similar exon structure from that point forward. Meanwhile, there appear to be large regions where the Volvox isoforms sequences do not line up with the Chlamydomonas sequence. It is unclear which isoform is more ancestral, however the somatic isoform has a higher percent similarity than the gonidial isoform.

*Supplemental Tables*

**Supplemental Table 1**: PCR Primers

| Number | Forward Primer (Fw) | Sequence | Reverse Primer (Rev) | Sequence |
|---|---|---|---|---|
| Gonidia /Soma-1 | Vocar.0001s1758 Fw1 | GTGAGTCTGGA GGCTGAG | Vocar.0001s1758 Rev1 | GGTGTGAATGA TAGCGGTTT |
| Gonidia /Soma - 2 | Vocar.0001s1758 Fw1 | GTGAGTCTGGA GGCTGAG | Vocar.0001s1758 Rev2 | GGAAGGGCACA ACTGATG |
| Gonidia /Soma - 3 | Vocar.0011s0285 Fw1 | CAGGATCGCCA CACTTTAATA | Vocar.0011s0285 Rev1 | CTGCACGACGA TTATTGTCA |
| Gonidia /Soma - 4 | Vocar.0011s0285 Fw2 | CGTATGACCAT GACAGTACC | Vocar.0011s0285 Rev1 | CTGCACGACGA TTATTGTCA |
| Gonidia /Soma - 5 | Vocar.0027s0034 Fw1 | GCAGATTCACA GGAGGTTATC | Vocar.0027s0034 Rev1 | CTGATCTATTTG CCGCCG |
| Gonidia /Soma - 6 | Vocar.0027s0034 Fw2 | GTTGCAACACA CGACTAAATG | Vocar.0027s0034 Rev1 | CTGATCTATTTG CCGCCG |
| Gonidia /Soma - 7 | Vocar.0001s1758 Fw1 | GTGAGTCTGGA GGCTGAG | Vocar.0001s1758 Rev1 + Rev2 | GGTGTGAATGA TAGCGGTTT<br><br>GGAAGGGCACA ACTGATG |

| | | | | |
|---|---|---|---|---|
| Gonidia /Soma - 8 | Vocar.0011s0285 Fw1 + Fw2 | CAGGATCGCCA CACTTTAATA<br><br>CGTATGACCAT GACAGTACC | Vocar.0011s0285 Rev1 | CTGCACGACGA TTATTGTCA |
| Gonidia /Soma - 9 | Vocar.0027s0034 Fw1 + Fw2 | GCAGATTCACA GGAGGTTATC<br><br>GTTGCAACACA CGACTAAATG | Vocar.0027s0034 Rev1 | CTGATCTATTTG CCGCCG |

List of all reactions including sample number (G/S =Gonidia/Soma), labels for the forward (Fw) and reverse (Rev) primers used (gene name Fw/Rev Number), and the total volume of each primer type. Sequences for each primer are listed. Gonidia/Soma means that the reactions were repeated with the same primers for both gonidia and soma separately.

**Supplemental Table T2:** Data for all 60 genes isolated via the restriction criteria applied for CTSAS. In separate file.

**Supplemental Table T3**: Data for the 12 genes isolated via the restriction criteria applied for CTSAS and hand-curation. In separate file.

**Supplemental Table T4:** RNA binding proteins of *V. carteri* and their differential expression data for soma and gonidia (data from Matt and Umen, 2018). In separate file.

*Supplemental Documents*

**Document D1**: Exon loci for the 12 candidate genes for CTSAS identified by HISAT2. Exons for each transcript are indicated as bands in genomic position. Read counts across the transcript lengths are displayed below transcript models. These include new transcripts that Phytozome did not previously identify.
