## Supplemental Table 2 for "Identification of Cell-Type Specific Alternative Transcripts in the Multicellular *Alga Volvox carteri*"

OppositeOneLarge_fullsummary_PhytonameExpr1

| | gene | gene_id | transcript | scaffold | strand | start_base | end_base | num_exons | length | FPKM.gon_1 | FPKM.gon_2 | FPKM.som_1 | FPKM.som_2 | gon_average | som_average | Expr.ratio | p.value | CTSAS.index | GU_gon_average | GU_som_average | GU_Log2_expRatio | GU_expPattern |
|---|---|---|---|---|---|---|---|---|---|---|---|---|---|---|---|---|---|---|---|---|---|---|
| 1 | . | MSTRG.84 | MSTRG.84.1 | scaffold_1 | + | 833458 | 836860 | 7 | 1839 | 0 | 0.128125 | 10.688086 | 8.737857 | 0.0640625 | 9.7129715 | 10.14896197 | 0.00208184 | 3.801196752 | . | . | . | . |
| 2 | Vocar.0001s0104 | MSTRG.84 | PAC4GC:715259 | scaffold_1 | + | 833458 | 836904 | 7 | 1880 | 15.990289 | 15.257029 | 7.834883 | 16.562786 | 15.623659 | 12.1988345 | 0.728026828 | 0.387534084 | 3.801196752 | 18.96 | 25.55 | 0.43 | Constitutive |
| 3 | Vocar.0001s0311 | MSTRG.245 | PAC4GC:715031 | scaffold_1 | - | 2341677 | 2344887 | 4 | 1962 | 534.229187 | 486.64325 | 182.402527 | 214.225616 | 510.4362185 | 198.3140715 | 0.385495106 | 0.000677417 | -3.809471156 | 614.88 | 253.7 | -1.28 | Low confidence |
| 4 | . | MSTRG.245 | MSTRG.245.2 | scaffold_1 | - | 2341682 | 2344887 | 5 | 1912 | 0 | 0 | 21.650108 | 0 | 0 | 10.825054 | 5.404853847 | 0.374870811 | -3.809471156 | . | . | . | . |
| 5 | Vocar.0001s0639 | MSTRG.474 | PAC4GC:714966 | scaffold_1 | + | 5010816 | 5014276 | 5 | 2468 | 0.211506 | 0.082021 | 1.232916 | 2.802331 | 0.1467635 | 2.0176235 | 2.483530144 | 0.08578817 | 3.31675259 | 3.6 | 2.54 | -0.5 | Constitutive |
| 6 | Vocar.0001s0639 | MSTRG.474 | PAC4GC:714967 | scaffold_1 | + | 5010816 | 5014276 | 6 | 2296 | 3.236624 | 2.778548 | 0 | 0 | 3.007586 | 0 | 0.249245553 | 0.031944538 | 3.31675259 | 3.6 | 2.54 | -0.5 | Constitutive |
| 7 | Vocar.0001s1138 | MSTRG.888 | PAC4GC:715283 | scaffold_1 | + | 8817789 | 8821971 | 7 | 1622 | 0 | 0 | 191.282242 | 169.5495 | 0 | 180.415871 | 181.9778674 | 0.005299291 | 8.305970956 | 3.51 | 218.61 | 5.96 | Somatic specific |
| 8 | . | MSTRG.888 | MSTRG.888.1 | scaffold_1 | + | 8817789 | 8823248 | 9 | 2730 | 6.801142 | 6.428853 | 3.335359 | 3.438738 | 6.6149975 | 3.3870485 | 0.575005727 | 0.024497622 | 8.305970956 | . | . | . | . |
| 9 | Vocar.0001s1410 | MSTRG.1087 | PAC4GC:714720 | scaffold_1 | + | 10792613 | 10793668 | 2 | 840 | 4.318156 | 3.758701 | 0 | 0 | 4.0384285 | 0 | 0.198248292 | 0.026647466 | -3.692143682 | 1.91 | 1.11 | -0.77 | Constitutive |
| 10 | Vocar.0001s1410 | MSTRG.1087 | PAC4GC:714719 | scaffold_1 | + | 10792613 | 10795000 | 3 | 1902 | 0 | 0 | 0.303724 | 4.678529 | 0 | 2.4911265 | 2.562450304 | 0.327307768 | -3.692143682 | 1.91 | 1.11 | -0.77 | Constitutive |
| 11 | Vocar.0001s1758 | MSTRG.1368 | PAC4GC:714674 | scaffold_1 | + | 13542209 | 13547795 | 13 | 2113 | 31.071108 | 29.248623 | 7.670852 | 8.436695 | 30.1598655 | 8.0537735 | 0.289013906 | 0.002617838 | -5.333992498 | 22.47 | 28.87 | 0.36 | Constitutive |
| 12 | Vocar.0001s1758 | MSTRG.1368 | PAC4GC:714673 | scaffold_1 | + | 13542209 | 13550813 | 14 | 3069 | 1.038219 | 0.943377 | 19.807402 | 25.4422 | 0.990798 | 22.624801 | 11.65763566 | 0.014902566 | -5.333992498 | 22.47 | 28.87 | 0.36 | Constitutive |
| 13 | Vocar.0011s0285 | MSTRG.1661 | PAC4GC:724672 | scaffold_11 | + | 2321358 | 2327887 | 8 | 3741 | 0.029278 | 0 | 45.558735 | 71.373711 | 0.014639 | 58.466223 | 56.1579082 | 0.022389733 | 8.967946247 | 6.1 | 68.88 | 3.5 | Somatic specific |
| 14 | . | MSTRG.1661 | MSTRG.1661.2 | scaffold_11 | + | 2324356 | 2327884 | 3 | 2992 | 8.849433 | 8.11259 | 0.030358 | 0.106083 | 8.4810115 | 0.0682205 | 0.112147631 | 0.006342485 | 8.967946247 | . | . | . | . |
| 15 | Vocar.0011s0296 | MSTRG.1667 | PAC4GC:724679 | scaffold_11 | - | 2424661 | 2435531 | 16 | 4845 | 13.111536 | 8.043317 | 0.162231 | 0.43301 | 10.5774265 | 0.2976205 | 0.112058388 | 0.056764161 | -4.09083049 | 9.96 | 2.05 | -2.28 | Gonidial specific |
| 16 | Vocar.0011s0296 | MSTRG.1667 | PAC4GC:724680 | scaffold_11 | - | 2424661 | 2435555 | 16 | 4860 | 0.016547 | 1.457866 | 1.427534 | 2.72362 | 0.7372065 | 2.075577 | 1.909444566 | 0.521872404 | -4.09083049 | 9.96 | 2.05 | -2.28 | Gonidial specific |
| 17 | Vocar.0012s0202 | MSTRG.1883 | PAC4GC:724624 | scaffold_12 | - | 1928511 | 1934238 | 10 | 3176 | 2.273755 | 1.936586 | 1.596247 | 0.088755 | 2.1051705 | 0.842501 | 0.560327098 | 0.369738486 | -3.366154514 | 2.78 | 9.01 | 1.7 | Low confidence |
| 18 | Vocar.0012s0202 | MSTRG.1883 | PAC4GC:724625 | scaffold_12 | - | 1928511 | 1934238 | 10 | 3090 | 0.182942 | 0.088479 | 4.175347 | 7.699463 | 0.1357105 | 5.937405 | 5.777701552 | 0.049562904 | -3.366154514 | 2.78 | 9.01 | 1.7 | Low confidence |
| 19 | Vocar20007194m.g | MSTRG.2173 | Vocar20007194m.2.0 | scaffold_13 | - | 2995092 | 2996757 | 1 | 1666 | 30.825924 | 19.747421 | 1.405373 | 1.134222 | 25.2866725 | 1.2697975 | 0.087693765 | 0.079127689 | -5.260936789 | 67.77 | 149.9 | 1.15 | Low confidence |
| 20 | . | MSTRG.2173 | MSTRG.2173.1 | scaffold_13 | - | 2995092 | 2997985 | 4 | 2099 | 34.429821 | 37.993633 | 130.762024 | 115.854897 | 36.211727 | 123.3084605 | 3.362548002 | 0.007914645 | -5.260936789 | . | . | . | . |
| 21 | . | MSTRG.2320 | MSTRG.2320.1 | scaffold_14 | + | 1525733 | 1528132 | 1 | 2400 | 0.131461 | 0.149254 | 12.428675 | 9.998949 | 0.1403575 | 11.213812 | 10.74885521 | 0.016770363 | 4.375456734 | . | . | . | . |
| 22 | . | MSTRG.2320 | MSTRG.2320.2 | scaffold_14 | + | 1525788 | 1528132 | 2 | 2125 | 4.306681 | 4.440683 | 0 | 8.27395 | 4.373682 | 4.136975 | 0.517867302 | 0.5693379 | 4.375456734 | . | . | . | . |

| | | | | | | | | | | | | | | | | | | | | | |
|---|---|---|---|---|---|---|---|---|---|---|---|---|---|---|---|---|---|---|---|---|---|
| 23 | . | MSTRG.2584 | MSTRG.2584.1 | scaffold_15 | + | 1687436 | 1699379 | 20 | 6286 | 2.555383 | 2.376377 | 17.000534 | 20.118183 | 2.46588 | 18.5593585 | 5.583828717 | 0.009694248 | 4.163834609 | . | . | . | . |
| 24 | Vocar.0015s0224 | MSTRG.2584 | PAC4GC:728492 | scaffold_15 | + | 1691767 | 1699379 | 12 | 4419 | 35.47438 | 33.936588 | 9.208444 | 11.327595 | 34.705484 | 10.2680195 | 0.311525049 | 0.022840299 | 4.163834609 | 25.1 | 32.15 | 0.36 | Constitutive |
| 25 | . | MSTRG.2855 | MSTRG.2855.1 | scaffold_16 | - | 1484204 | 1487907 | 8 | 1114 | 6.817277 | 4.997365 | 0.534875 | 0.10581 | 5.907321 | 0.3203425 | 0.191599692 | 0.106868607 | -6.968639386 | . | . | . | . |
| 26 | Vocar.0016s0187 | MSTRG.2855 | PAC4GC:727485 | scaffold_16 | - | 1487858 | 1491301 | 6 | 1841 | 1.490429 | 0.882817 | 43.188118 | 62.825443 | 1.186623 | 53.0067805 | 23.9974055 | 0.007559364 | -6.968639386 | 1.69 | 64.29 | 5.25 | Somatic specific |
| 27 | Vocar.0016s0251 | MSTRG.2912 | PAC4GC:727760 | scaffold_16 | - | 1882653 | 1890980 | 9 | 4870 | 3.525218 | 3.041289 | 1.002062 | 0.149408 | 3.2832535 | 0.575735 | 0.361856248 | 0.164922958 | -6.4231455 | 4.2 | 25.49 | 2.6 | Somatic specific |
| 28 | Vocar.0016s0251 | MSTRG.2912 | PAC4GC:727761 | scaffold_16 | - | 1885088 | 1890980 | 8 | 2527 | 0 | 0 | 24.126991 | 38.845455 | 0 | 31.486223 | 31.05242612 | 0.030832364 | -6.4231455 | 4.2 | 25.49 | 2.6 | Somatic specific |
| 29 | Vocar.0017s0204 | MSTRG.3116 | PAC4GC:721054 | scaffold_17 | - | 2276465 | 2285770 | 4 | 8316 | 0 | 0 | 6.763828 | 4.083592 | 0 | 5.42371 | 6.391774801 | 0.052369435 | 3.529005474 | 3.37 | 8.79 | 1.39 | Low confidence |
| 30 | Vocar.0017s0204 | MSTRG.3116 | PAC4GC:721052 | scaffold_17 | - | 2276465 | 2285996 | 5 | 8419 | 3.075252 | 2.522674 | 0 | 4.059011 | 2.798963 | 2.0295055 | 0.55371331 | 0.444871972 | 3.529005474 | 3.37 | 8.79 | 1.39 | Low confidence |
| 31 | Vocar.0018s0107 | MSTRG.3211 | PAC4GC:728217 | scaffold_18 | + | 813935 | 823398 | 11 | 5148 | 6.928167 | 4.247898 | 0 | 1.637437 | 5.5880325 | 0.8187185 | 0.239613965 | 0.043904248 | -4.620748714 | 7.5 | 10.31 | 0.46 | Constitutive |
| 32 | Vocar.0018s0107 | MSTRG.3211 | PAC4GC:728218 | scaffold_18 | + | 814746 | 823398 | 11 | 4950 | 0 | 1.505609 | 8.698296 | 7.496864 | 0.7528045 | 8.09758 | 5.895166699 | 0.178966568 | -4.620748714 | 7.5 | 10.31 | 0.46 | Constitutive |
| 33 | Vocar.0002s0039 | MSTRG.3600 | PAC4GC:720640 | scaffold_2 | - | 272487 | 276178 | 13 | 1599 | 32.3382 | 28.5474 | 8.247209 | 11.118618 | 30.4428 | 9.6829135 | 0.332618801 | 0.013916718 | -3.820594731 | 35.48 | 12.01 | -1.56 | Gonidial biased |
| 34 | . | MSTRG.3600 | MSTRG.3600.2 | scaffold_2 | - | 274074 | 277473 | 8 | 1670 | 0 | 0 | 0.389163 | 18.740366 | 0 | 9.5647645 | 4.699594734 | 0.353055618 | -3.820594731 | . | . | . | . |

| | | | | | | | | | | | | | | | | | | | | | | |
|---|---|---|---|---|---|---|---|---|---|---|---|---|---|---|---|---|---|---|---|---|---|---|
| 35 | Vocar.0002s0180 | MSTRG.3719 | PAC4GC:720493 | scaffold_2 | + | 1505693 | 1513365 | 11 | 3616 | 9.90454 | 1.53404 | 10.358089 | 8.520172 | 5.71929 | 9.4391305 | 1.923463261 | 0.617241853 | 5.891773301 | 13 | 10.75 | -0.27 | Constitutive |
| 36 | . | MSTRG.3719 | MSTRG.3719.2 | scaffold_2 | + | 1509859 | 1513367 | 5 | 2456 | 27.236959 | 33.244682 | 0.004976 | 0 | 30.2408205 | 0.002488 | 0.032395404 | 0.021491624 | 5.891773301 | . | . | . | . |
| 37 | . | MSTRG.3844 | MSTRG.3844.1 | scaffold_2 | + | 2651846 | 2654434 | 7 | 959 | 0 | 0 | 25.911076 | 21.124086 | 0 | 23.517581 | 24.59611377 | 0.0140603 | 5.339775347 | . | . | . | . |
| 38 | Vocar.0002s0335 | MSTRG.3844 | PAC4GC:720613 | scaffold_2 | + | 2651846 | 2654434 | 6 | 1191 | 269.335693 | 293.131104 | 142.052414 | 209.750748 | 281.2333985 | 175.901581 | 0.607342879 | 0.23329247 | 5.339775347 | 328.71 | 228.5 | -0.52 | Constitutive |
| 39 | Vocar.0021s0174 | MSTRG.4448 | PAC4GC:725625 | scaffold_21 | - | 1873620 | 1878433 | 6 | 3006 | 0 | 0 | 14.775574 | 15.087212 | 0 | 14.931393 | 15.91793017 | 0.001625273 | 4.231022514 | 2.32 | 23.89 | 3.36 | Somatic specific |
| 40 | . | MSTRG.4448 | MSTRG.4448.1 | scaffold_21 | - | 1873620 | 1880345 | 7 | 4446 | 3.307399 | 3.50919 | 0.879138 | 7.370333 | 3.4082945 | 4.1247355 | 0.847660422 | 0.819575016 | 4.231022514 | . | . | . | . |
| 41 | Vocar.0022s0092 | MSTRG.4553 | PAC4GC:721507 | scaffold_22 | + | 929738 | 942676 | 28 | 5247 | 0.014392 | 0 | 3.077606 | 5.484785 | 0.007196 | 4.2811955 | 5.008209566 | 0.062568705 | 7.731051727 | 49.15 | 5.16 | -3.25 | Gonidial specific |
| 42 | Vocar.0022s0092 | MSTRG.4553 | PAC4GC:721508 | scaffold_22 | + | 929738 | 942676 | 27 | 5166 | 45.735771 | 37.130627 | 0 | 0 | 41.433199 | 0 | 0.023572412 | 0.021069823 | 7.731051727 | 49.15 | 5.16 | -3.25 | Gonidial specific |
| 43 | . | MSTRG.4810 | MSTRG.4810.2 | scaffold_23 | + | 1730458 | 1740386 | 16 | 3249 | 1.407032 | 0.356848 | 5.165819 | 5.293861 | 0.88194 | 5.22984 | 3.396809923 | 0.173663343 | 4.600175004 | . | . | . | . |
| 44 | Vocar.0023s0175 | MSTRG.4810 | PAC4GC:712921 | scaffold_23 | + | 1731460 | 1740382 | 15 | 2484 | 6.660413 | 5.608265 | 0 | 0 | 6.134339 | 0 | 0.140049184 | 0.029154921 | 4.600175004 | 8.88 | 6.45 | -0.46 | Constitutive |
| 45 | . | MSTRG.5198 | MSTRG.5198.1 | scaffold_25 | - | 1863730 | 1864125 | 2 | 226 | 5.31846 | 6.800029 | 0.612907 | 0.943028 | 6.0592445 | 0.7779675 | 0.251537442 | 0.089671442 | -4.246053047 | . | . | . | . |
| 46 | . | MSTRG.5198 | MSTRG.5198.2 | scaffold_25 | - | 1863740 | 1868521 | 8 | 1232 | 0 | 0.025698 | 3.343902 | 4.475035 | 0.012849 | 3.9094685 | 4.773005899 | 0.040292984 | -4.246053047 | . | . | . | . |
| 47 | Vocar.0027s0034 | MSTRG.5391 | PAC4GC:716687 | scaffold_27 | - | 330803 | 333910 | 8 | 2066 | 2.772401 | 4.502927 | 69.698517 | 61.393063 | 3.637664 | 65.54579 | 14.78565024 | 0.043675021 | 4.53837693 | 3246.03 | 2159.96 | -0.59 | Constitutive |
| 48 | Vocar.0027s0034 | MSTRG.5391 | PAC4GC:716688 | scaffold_27 | - | 330803 | 333910 | 9 | 1839 | 3118.518555 | 3046.334473 | 2070.433594 | 1841.400024 | 3082.426514 | 1955.916809 | 0.636286693 | 0.079279571 | 4.53837693 | 3246.03 | 2159.96 | -0.59 | Constitutive |
| 49 | Vocar.0028s0067 | MSTRG.5584 | PAC4GC:725240 | scaffold_28 | + | 781234 | 784660 | 7 | 2516 | 61.839245 | 46.46307 | 3.631404 | 10.338736 | 54.1511575 | 6.98507 | 0.127071366 | 0.046903161 | -3.152601316 | 66.76 | 11.55 | -2.53 | Gonidial specific |
| 50 | Vocar.0028s0067 | MSTRG.5584 | PAC4GC:725241 | scaffold_28 | + | 781234 | 784660 | 6 | 2654 | 0.527484 | 7.804008 | 2.521322 | 3.577738 | 4.165746 | 3.04953 | 1.129991714 | 0.933335741 | -3.152601316 | 66.76 | 11.55 | -2.53 | Gonidial specific |
| 51 | Vocar.0028s0134 | MSTRG.5637 | PAC4GC:725300 | scaffold_28 | - | 1274674 | 1282727 | 12 | 2149 | 4.379229 | 3.520309 | 0.003063 | 0 | 3.949769 | 0.0015315 | 0.202278432 | 0.042632218 | -5.533763732 | 6.24 | 20.91 | 1.75 | Low confidence |

| | | | | | | | | | | | | | | | | | | | | | | |
|---|---|---|---|---|---|---|---|---|---|---|---|---|---|---|---|---|---|---|---|---|---|---|
| 52 | Vocar.0028s0134 | MSTRG.5637 | PAC4GC:725299 | scaffold_28 | - | 1274677 | 1282727 | 11 | 3015 | 1.168971 | 0.715355 | 14.957023 | 20.192558 | 0.942163 | 17.5747905 | 9.370838448 | 0.011545505 | -5.533763732 | 6.24 | 20.91 | 1.75 | Low confidence |
| 53 | . | MSTRG.5815 | MSTRG.5815.9 | scaffold_3 | - | 163497 | 170671 | 16 | 4003 | 6.923425 | 8.896676 | 0 | 0 | 7.9100505 | 0 | 0.113535375 | 0.039625297 | -3.894231493 | . | . | . | . |
| 54 | . | MSTRG.5815 | MSTRG.5815.10 | scaffold_3 | - | 163517 | 171685 | 17 | 4804 | 4.145289 | 0.64624 | 3.731758 | 4.452962 | 2.3957645 | 4.09236 | 1.688152546 | 0.565215963 | -3.894231493 | . | . | . | . |
| 55 | Vocar20012504m.g | MSTRG.5840 | Vocar20012504m.2.0 | scaffold_3 | - | 429142 | 429643 | 2 | 426 | 3.524976 | 6.428251 | 24.545616 | 20.812023 | 4.9766135 | 22.6788195 | 4.147165532 | 0.108662447 | 5.39722148 | 15.67 | 13.42 | -0.22 | Constitutive |
| 56 | . | MSTRG.5840 | MSTRG.5840.1 | scaffold_3 | - | 429142 | 432105 | 5 | 1689 | 33.768219 | 35.122581 | 2.363889 | 2.631449 | 34.4454 | 2.497669 | 0.098406959 | 0.013985069 | 5.39722148 | . | . | . | . |
| 57 | Vocar.0030s0060 | MSTRG.6247 | PAC4GC:724162 | scaffold_30 | + | 559876 | 565992 | 8 | 3837 | 0.182271 | 0.069037 | 9.07756 | 11.550079 | 0.125654 | 10.3138195 | 9.890195672 | 0.004961037 | 5.814105153 | 0.41 | 14.38 | 5.15 | Somatic specific |
| 58 | . | MSTRG.6247 | MSTRG.6247.2 | scaffold_30 | + | 562744 | 568645 | 9 | 2905 | 10.268705 | 10.022028 | 0.7424 | 1.251655 | 10.1453665 | 0.9970275 | 0.175786223 | 0.028949322 | 5.814105153 | . | . | . | . |
| 59 | Vocar.0030s0109 | MSTRG.6279 | PAC4GC:724073 | scaffold_30 | - | 941473 | 947658 | 4 | 4656 | 0.481395 | 0 | 4.072567 | 3.06493 | 0.2406975 | 3.5687485 | 3.729103755 | 0.151655222 | 5.77180038 | 15.27 | 4.3 | -1.83 | Gonidial biased |
| 60 | . | MSTRG.6279 | MSTRG.6279.2 | scaffold_30 | - | 941879 | 947572 | 3 | 4200 | 13.275834 | 14.076639 | 0 | 0 | 13.6762365 | 0 | 0.068252635 | 0.007891549 | 5.77180038 | . | . | . | . |
| 61 | . | MSTRG.6393 | MSTRG.6393.1 | scaffold_31 | - | 791923 | 800242 | 2 | 1729 | 3.38133 | 1.692265 | 6.861724 | 7.873623 | 2.5367975 | 7.3676735 | 2.391641468 | 0.179911015 | 3.673117101 | . | . | . | . |
| 62 | Vocar20013296m.g | MSTRG.6393 | Vocar20013296m.2.0 | scaffold_31 | - | 792397 | 795286 | 4 | 519 | 4.055518 | 4.657697 | 0 | 0 | 4.3566075 | 0 | 0.187489802 | 0.026080845 | 3.673117101 | 10.54 | 4.83 | -1.13 | Low confidence |
| 63 | . | MSTRG.6517 | MSTRG.6517.1 | scaffold_32 | - | 465143 | 472488 | 11 | 4172 | 0.839622 | 0.700687 | 23.699373 | 29.783619 | 0.7701545 | 26.741496 | 15.42067947 | 0.007364754 | 9.310966984 | . | . | . | . |
| 64 | Vocar.0032s0063 | MSTRG.6517 | PAC4GC:723368 | scaffold_32 | - | 469248 | 472488 | 7 | 1082 | 41.817276 | 40.79491 | 0.031704 | 0.02291 | 41.306093 | 0.027307 | 0.024278573 | 0.003052422 | 9.310966984 | 48.45 | 8.86 | -2.45 | Gonidial specific |
| 65 | Vocar.0033s0137 | MSTRG.6695 | PAC4GC:715939 | scaffold_33 | - | 1063256 | 1068822 | 9 | 2222 | 2.26689 | 1.719803 | 0 | 2.073272 | 1.9933465 | 1.036636 | 0.559321865 | 0.297996416 | -4.272089808 | 2.83 | 13.85 | 2.29 | Somatic biased |
| 66 | Vocar.0033s0137 | MSTRG.6695 | PAC4GC:715940 | scaffold_33 | - | 1063256 | 1068822 | 10 | 2132 | 0.123636 | 0.1186 | 11.860853 | 10.295535 | 0.121118 | 11.078194 | 10.80659743 | 0.013267858 | -4.272089808 | 2.83 | 13.85 | 2.29 | Somatic biased |

| | | | | | | | | | | | | | | | | | | | | | | |
|---|---|---|---|---|---|---|---|---|---|---|---|---|---|---|---|---|---|---|---|---|---|---|
| 67 | Vocar.0035s0051 | MSTRG.6861 | PAC4GC:726466 | scaffold_35 | - | 431881 | 436778 | 6 | 3993 | 5.189751 | 5.796627 | 4.477898 | 2.773444 | 5.493189 | 3.625671 | 0.71330227 | 0.143713243 | -4.947990551 | 7.43 | 48.89 | 2.72 | Somatic specific |
| 68 | . | MSTRG.6861 | MSTRG.6861.1 | scaffold_35 | - | 431881 | 437027 | 7 | 4152 | 1.557412 | 0.142355 | 34.132244 | 41.573456 | 0.8498835 | 37.85285 | 22.01745819 | 0.086361102 | -4.947990551 | . | . | . | . |
| 69 | Vocar.0038s0054 | MSTRG.7243 | PAC4GC:717261 | scaffold_38 | - | 945934 | 972835 | 20 | 9938 | 1.892656 | 1.760809 | 0.009139 | 0.44682 | 1.8267325 | 0.2279795 | 0.420869601 | 0.074451294 | -3.757707152 | 1.21 | 8.6 | 2.82 | Somatic specific |
| 70 | . | MSTRG.7243 | MSTRG.7243.3 | scaffold_38 | - | 946026 | 972774 | 19 | 9845 | 0.131278 | 0 | 3.501766 | 7.640172 | 0.065639 | 5.570969 | 5.692855043 | 0.058494889 | -3.757707152 | . | . | . | . |
| 71 | Vocar.0039s0085 | MSTRG.7312 | PAC4GC:727415 | scaffold_39 | - | 832092 | 836377 | 6 | 3217 | 2.737362 | 1.441622 | 7.037776 | 11.479494 | 2.089492 | 9.258635 | 3.223217708 | 0.054635076 | 5.287297041 | 3.82 | 6.2 | 0.7 | Constitutive |
| 72 | . | MSTRG.7312 | MSTRG.7312.2 | scaffold_39 | - | 832129 | 834593 | 4 | 1944 | 8.834282 | 14.245603 | 0 | 0 | 11.5399425 | 0 | 0.082538132 | 0.06796487 | 5.287297041 | . | . | . | . |
| 73 | Vocar.0004s0120 | MSTRG.7416 | PAC4GC:719327 | scaffold_4 | - | 1007444 | 1018654 | 17 | 2652 | 0 | 0 | 6.690708 | 11.733656 | 0 | 9.212182 | 9.694629271 | 0.050930182 | 5.413874662 | 0.26 | 9.99 | 5.25 | Somatic specific |

| | | | | | | | | | | | | | | | | | | | | | | |
|---|---|---|---|---|---|---|---|---|---|---|---|---|---|---|---|---|---|---|---|---|---|---|
| 74 | Vocar.0004s0121 | MSTRG.7416 | PAC4GC:719376 | scaffold_4 | - | 1015550 | 1027717 | 20 | 5435 | 9.155589 | 7.197547 | 1.313049 | 0.846292 | 8.176568 | 1.0796705 | 0.227401089 | 0.090515351 | 5.413874662 | 10.09 | 0.61 | -4.05 | Gonidial specific |
| 75 | . | MSTRG.7492 | MSTRG.7492.1 | scaffold_4 | - | 1834464 | 1835237 | 1 | 774 | 11.205629 | 9.095021 | 180.38945 | 168.751068 | 10.150325 | 174.570259 | 15.77834764 | 0.032215017 | 5.700038115 | . | . | . | . |
| 76 | Vocar.0004s0224 | MSTRG.7492 | PAC4GC:719129 | scaffold_4 | - | 1834464 | 1836457 | 5 | 1143 | 8.716306 | 11.233939 | 1.773707 | 3.024516 | 9.9751225 | 2.3991115 | 0.30351424 | 0.144271089 | 5.700038115 | 16.57 | 85.26 | 2.36 | Somatic biased |
| 77 | Vocar.0004s0303 | MSTRG.7556 | PAC4GC:719267 | scaffold_4 | - | 2368850 | 2380393 | 6 | 3654 | 7.42391 | 0.263286 | 2.73383 | 4.202516 | 3.843598 | 3.468173 | 1.273217426 | 0.854326498 | 3.822649103 | 16.39 | 4.13 | -1.99 | Gonidial biased |
| 78 | . | MSTRG.7556 | MSTRG.7556.2 | scaffold_4 | - | 2369079 | 2374108 | 5 | 3346 | 8.217759 | 15.995314 | 0.09978 | 0.121639 | 12.1065365 | 0.1107095 | 0.089985085 | 0.099677985 | 3.822649103 | . | . | . | . |
| 79 | Vocar.0004s0325 | MSTRG.7577 | PAC4GC:718989 | scaffold_4 | - | 2649704 | 2653136 | 4 | 1194 | 0 | 0.117853 | 2.586477 | 1.882073 | 0.0589265 | 2.234275 | 3.07634117 | 0.006255207 | 5.268071601 | 10.57 | 2.92 | -1.86 | Gonidial biased |
| 80 | . | MSTRG.7577 | MSTRG.7577.2 | scaffold_4 | - | 2649796 | 2651207 | 2 | 995 | 11.874612 | 11.152957 | 0 | 0 | 11.5137845 | 0 | 0.079833823 | 0.008857754 | 5.268071601 | . | . | . | . |
| 81 | . | MSTRG.7781 | MSTRG.7781.1 | scaffold_40 | + | 709305 | 717033 | 8 | 5307 | 0.905582 | 1.188853 | 35.111279 | 49.975292 | 1.0472175 | 42.5432855 | 20.78397045 | 0.043785568 | 9.824044179 | . | . | . | . |
| 82 | Vocar20010095m.g | MSTRG.7781 | Vocar20010095m.2.0 | scaffold_40 | + | 716260 | 717033 | 1 | 774 | 42.76453 | 42.734123 | 0 | 0.006919 | 42.7493265 | 0.0034595 | 0.022929603 | 0.000348476 | 9.824044179 | 55.5 | 0.11 | -8.99 | Gonidial specific |
| 83 | Vocar.0041s0075 | MSTRG.7856 | PAC4GC:718318 | scaffold_41 | + | 708457 | 710773 | 1 | 2317 | 4.058009 | 3.399011 | 4.0191 | 3.333557 | 3.72851 | 3.6763285 | 0.991298031 | 0.959780815 | -4.842952069 | 4.09 | 1.66 | -1.3 | Low confidence |
| 84 | Vocar.0041s0076 | MSTRG.7856 | PAC4GC:718329 | scaffold_41 | + | 710122 | 738660 | 38 | 10551 | 0.110651 | 0.071142 | 29.279177 | 31.00033 | 0.0908965 | 30.1397535 | 28.44972572 | 0.000402725 | -4.842952069 | 0.27 | 37.07 | 7.12 | Somatic specific |
| 85 | . | MSTRG.7890 | MSTRG.7890.1 | scaffold_42 | + | 162960 | 167181 | 5 | 3236 | 8.861084 | 9.309844 | 0.035464 | 0.124469 | 9.085464 | 0.0799665 | 0.106772519 | 0.016194407 | -8.415231254 | . | . | . | . |
| 86 | Vocar.0042s0026 | MSTRG.7890 | PAC4GC:721774 | scaffold_42 | + | 164598 | 167181 | 3 | 2189 | 0 | 0 | 27.189533 | 48.331371 | 0 | 37.760452 | 36.4499148 | 0.035744732 | -8.415231254 | 2.99 | 35.29 | 3.56 | Somatic specific |
| 87 | Vocar.0042s0098 | MSTRG.7941 | PAC4GC:721832 | scaffold_42 | + | 695781 | 707608 | 22 | 4200 | 28.447895 | 24.168289 | 1.551953 | 2.905361 | 26.308092 | 2.228657 | 0.113535508 | 0.016944701 | -4.950921258 | 31.82 | 4.72 | -2.75 | Gonidial specific |
| 88 | . | MSTRG.7941 | MSTRG.7941.2 | scaffold_42 | + | 705360 | 706145 | 1 | 786 | 2.139013 | 3.650192 | 11.574377 | 13.184559 | 2.8946025 | 12.379468 | 3.511619982 | 0.141150995 | -4.950921258 | . | . | . | . |
| 89 | . | MSTRG.8537 | MSTRG.8537.1 | scaffold_5 | - | 1875206 | 1891288 | 15 | 9287 | 3.994195 | 3.730544 | 1.019645 | 0.46346 | 3.8623695 | 0.7415525 | 0.357910603 | 0.09193053 | -3.091655235 | . | . | . | . |

| | | | | | | | | | | | | | | | | | | | | | | |
|---|---|---|---|---|---|---|---|---|---|---|---|---|---|---|---|---|---|---|---|---|---|---|
| 90 | . | MSTRG.8537 | MSTRG.8537.3 | scaffold_5 | - | 1882159 | 1888972 | 8 | 5356 | 5.918167 | 0 | 6.255115 | 9.004852 | 2.9590835 | 7.6299835 | 3.051093569 | 0.484746836 | -3.091655235 | . | . | . | . |
| 91 | Vocar.0005s0310 | MSTRG.8581 | PAC4GC:713334 | scaffold_5 | - | 2368195 | 2369336 | 1 | 1142 | 359.184631 | 357.766907 | 132.913864 | 131.515839 | 358.475769 | 132.2148515 | 0.370699801 | 0.003974265 | -3.629539361 | 416.15 | 166.56 | -1.32 | Low confidence |
| 92 | . | MSTRG.8581 | MSTRG.8581.2 | scaffold_5 | - | 2368198 | 2369261 | 2 | 960 | 0 | 0 | 0 | 26.588469 | 0 | 13.2942345 | 4.58799114 | 0.424270329 | -3.629539361 | . | . | . | . |
| 93 | Vocar.0005s0373 | MSTRG.8625 | PAC4GC:713389 | scaffold_5 | + | 2880859 | 2884486 | 8 | 1353 | 16.620239 | 20.830643 | 0 | 0 | 18.725441 | 0 | 0.051251684 | 0.02797726 | -5.299771988 | 21.79 | 4.98 | -2.13 | Gonidial biased |
| 94 | . | MSTRG.8625 | MSTRG.8625.1 | scaffold_5 | + | 2880859 | 2885583 | 10 | 1691 | 1.077276 | 0.867182 | 1.999248 | 4.572366 | 0.972229 | 3.285807 | 2.01882409 | 0.141453461 | -5.299771988 | . | . | . | . |
| 95 | Vocar.0005s0510 | MSTRG.8727 | PAC4GC:713309 | scaffold_5 | - | 4119325 | 4125981 | 6 | 4784 | 14.747833 | 11.489532 | 0 | 13.287538 | 13.1186825 | 6.643769 | 0.240477286 | 0.332110235 | -4.655280539 | 15.25 | 19.85 | 0.38 | Constitutive |
| 96 | . | MSTRG.8727 | MSTRG.8727.1 | scaffold_5 | - | 4119325 | 4132221 | 16 | 7887 | 1.245678 | 0.795705 | 13.276483 | 9.2013 | 1.0206915 | 11.2388915 | 6.059728169 | 0.090457744 | -4.655280539 | . | . | . | . |
| 97 | . | MSTRG.8947 | MSTRG.8947.1 | scaffold_54 | + | 310263 | 313915 | 2 | 501 | 24.554502 | 21.458118 | 0 | 0.000606 | 23.00631 | 0.000303 | 0.041624185 | 0.015724244 | -5.184042764 | . | . | . | . |
| 98 | Vocar.0054s0037 | MSTRG.8947 | PAC4GC:724180 | scaffold_54 | + | 311364 | 313915 | 3 | 2408 | 4.810687 | 5.285655 | 7.851524 | 8.464064 | 5.048171 | 8.157794 | 1.51320621 | 0.10967362 | -5.184042764 | 3.18 | 9.05 | 1.51 | Low confidence |
| 99 | . | MSTRG.8980 | MSTRG.8980.2 | scaffold_55 | - | 124618 | 128867 | 3 | 3680 | 6.08406 | 6.319328 | 0 | 0 | 6.201694 | 0 | 0.138984365 | 0.006426473 | -5.341012826 | . | . | . | . |
| 100 | Vocar.0055s0013 | MSTRG.8980 | PAC4GC:721171 | scaffold_55 | - | 124618 | 128881 | 2 | 3903 | 3.737332 | 2.479733 | 19.232286 | 25.858477 | 3.1085325 | 22.5453815 | 5.633405531 | 0.03157065 | -5.341012826 | 9.3 | 23.46 | 1.34 | Low confidence |
| 101 | . | MSTRG.9394 | MSTRG.9394.1 | scaffold_6 | - | 2628570 | 2636864 | 14 | 4072 | 0.166103 | 0.378826 | 21.778744 | 30.361937 | 0.2724645 | 26.0703405 | 20.89000263 | 0.045515362 | 8.559655455 | . | . | . | . |
| 102 | Vocar.0006s0263 | MSTRG.9394 | PAC4GC:723710 | scaffold_6 | - | 2628597 | 2635765 | 12 | 3195 | 18.472054 | 15.627407 | 0 | 0 | 17.0497305 | 0 | 0.055363744 | 0.021176788 | 8.559655455 | 19.65 | 16.81 | -0.22 | Constitutive |
| 103 | Vocar.0006s0317 | MSTRG.9450 | PAC4GC:723448 | scaffold_6 | - | 2990966 | 2993343 | 5 | 1575 | 0.563458 | 0.110562 | 131.485458 | 143.008408 | 0.33701 | 137.246933 | 103.609458 | 0.024466349 | 7.277860545 | 4.35 | 115.48 | 4.73 | Somatic specific |
| 104 | . | MSTRG.9450 | MSTRG.9450.2 | scaffold_6 | - | 2990999 | 2992178 | 1 | 1180 | 23.520716 | 26.410393 | 13.209172 | 20.70916 | 24.9655545 | 16.959166 | 0.667644189 | 0.318679874 | 7.277860545 | . | . | . | . |
| 105 | . | MSTRG.9452 | MSTRG.9452.1 | scaffold_6 | + | 3027613 | 3030234 | 5 | 2134 | 10.65753 | 9.179134 | 0.00091 | 0 | 9.918332 | 0.000455 | 0.091544569 | 0.022096045 | -5.117092105 | . | . | . | . |
| 106 | Vocar.0006s0322 | MSTRG.9452 | PAC4GC:723697 | scaffold_6 | + | 3029056 | 3033311 | 9 | 2691 | 0.722356 | 1.113538 | 4.714942 | 5.427638 | 0.917947 | 5.07129 | 3.177099364 | 0.091470902 | -5.117092105 | 2.39 | 4.6 | 0.95 | Constitutive |
| 107 | Vocar.0006s0382 | MSTRG.9496 | PAC4GC:723709 | scaffold_6 | + | 3427162 | 3430512 | 7 | 1967 | 780.335938 | 831.460388 | 305.476013 | 258.920471 | 805.898163 | 282.198242 | 0.35285805 | 0.012747857 | -3.826652487 | 990.06 | 380.31 | -1.38 | Low confidence |
| 108 | . | MSTRG.9496 | MSTRG.9496.2 | scaffold_6 | + | 3427163 | 3430512 | 8 | 1737 | 52.457832 | 0 | 42.37521 | 35.951492 | 26.228916 | 39.163351 | 5.006534484 | 0.633767278 | -3.826652487 | . | . | . | . |
| 109 | . | MSTRG.9702 | MSTRG.9702.2 | scaffold_64 | + | 244375 | 249361 | 6 | 3874 | 6.695939 | 7.242332 | 1.993226 | 0.830645 | 6.9691355 | 1.4119355 | 0.300359664 | 0.071553774 | -3.036052978 | . | . | . | . |
| 110 | Vocar.0064s0022 | MSTRG.9702 | PAC4GC:727291 | scaffold_64 | + | 246875 | 249361 | 5 | 1870 | 3.348914 | 2.286651 | 8.725819 | 7.983385 | 2.8177825 | 8.354602 | 2.463681852 | 0.138588071 | -3.036052978 | 9.37 | 6.43 | -0.54 | Constitutive |
| 111 | . | MSTRG.9907 | MSTRG.9907.1 | scaffold_7 | - | 931938 | 940203 | 7 | 5571 | 0 | 0 | 7.097149 | 13.930489 | 0 | 10.513819 | 10.7242832 | 0.059130586 | 4.936610925 | . | . | . | . |
| 112 | Vocar.0007s0099 | MSTRG.9907 | PAC4GC:717737 | scaffold_7 | - | 931938 | 940205 | 6 | 6005 | 14.862828 | 13.818495 | 3.718909 | 5.272663 | 14.3406615 | 4.495786 | 0.350187233 | 0.037132993 | 4.936610925 | 6.34 | 1.72 | -1.88 | Gonidial biased |

| | | | | | | | | | | | | | | | | | | | | | | |
|---|---|---|---|---|---|---|---|---|---|---|---|---|---|---|---|---|---|---|---|---|---|---|
| 113 | Vocar.0008s0063 | MSTRG.10419 | PAC4GC:726579 | scaffold_8 | + | 589616 | 592792 | 5 | 1534 | 12.976501 | 18.62113 | 4.804866 | 0 | 15.7988155 | 2.402433 | 0.157591974 | 0.145020819 | -5.103817954 | 10.69 | 4.63 | -1.21 | Low confidence |
| 114 | . | MSTRG.10419 | MSTRG.10419.2 | scaffold_8 | + | 589876 | 592551 | 5 | 1106 | 0 | 0 | 3.263095 | 6.188732 | 0 | 4.7259135 | 5.419215121 | 0.070799258 | -5.103817954 | . | . | . | . |
| 115 | Vocar.0008s0322 | MSTRG.10614 | PAC4GC:726528 | scaffold_8 | - | 2745174 | 2747769 | 5 | 1861 | 0 | 0 | 10.051441 | 8.261657 | 0 | 9.156549 | 10.19018901 | 0.017495005 | 4.464220331 | 3.84 | 12.62 | 1.72 | Low confidence |
| 116 | Vocar.0008s0322 | MSTRG.10614 | PAC4GC:726529 | scaffold_8 | - | 2745174 | 2747769 | 5 | 1865 | 3.281902 | 2.721073 | 0 | 2.813169 | 3.0014875 | 1.4065845 | 0.461655499 | 0.302600554 | 4.464220331 | 3.84 | 12.62 | 1.72 | Low confidence |
| 117 | Vocar.0009s0279 | MSTRG.11003 | PAC4GC:723068 | scaffold_9 | + | 2748631 | 2758128 | 21 | 3287 | 2.10411 | 1.512163 | 1.391351 | 2.993706 | 1.8081365 | 2.1925285 | 1.078243508 | 0.408140125 | 3.114986017 | 5.82 | 2.87 | -1.02 | Low confidence |
| 118 | . | MSTRG.11003 | MSTRG.11003.2 | scaffold_9 | + | 2754533 | 2758873 | 9 | 1951 | 7.316256 | 6.736248 | 0 | 0 | 7.026252 | 0 | 0.124455079 | 0.013465476 | 3.114986017 | . | . | . | . |
| 119 | Vocar20014470m.g | MSTRG.11131 | Vocar20014470m.2.0 | scaffold_97 | - | 840 | 2813 | 1 | 1974 | 1.462812 | 3.674678 | 87.366196 | 67.117584 | 2.568745 | 77.24189 | 23.46989074 | 0.059692935 | 4.761135599 | 29.8 | 93.88 | 1.66 | Low confidence |
| 120 | . | MSTRG.11131 | MSTRG.11131.1 | scaffold_97 | - | 840 | 4761 | 3 | 2846 | 23.219858 | 21.501585 | 20.590895 | 17.759081 | 22.3607215 | 19.174988 | 0.865498738 | 0.372344162 | 4.761135599 | . | . | . | . |

| sequence | ProteinSeq | StartStop | Chlamy Protein Homolog | Score |
|---|---|---|---|---|
| CCGTAGGTTGTGCGCTTATGCTTGACGGCAGTACAGCAAGGCTTGTTTAGCAGTCTCTGTGCGTAGCTTGGCCCCCACACGGACAAGAATGAGCAACTCGGGTGGGGGCTTCCTCTCAAAATTGAAAGCACTAGATGCATATCCCAAA | >lcl\|ORF2<br>MSNSGGGF<br>LRLYMTTQS<br>HLDLVDHDV<br>GSCYGAEDI<br>EQTGEGCHI<br>RHTVNKLSF<br>DNKTLATNC<br>FLSFLTSVC | Start 89 Stop 1270 | >Cre13.g579850.t1.2<br>MSGGGFLGKLKALDAYPKI<br>PCAWLSLDAMDISGELHLD<br>SCYGAEDKQGDCCNTCDE<br>HVIHKLSFGEPYPGMKNPL<br>SFLSFLTSVCAIVGGVFTVS | 680.6 |
| CCGTAGGTTGTGCGCTTATGCTTGACGGCAGTACAGCAAGGCTTGTTTAGCAGTCTCTGTGCGTAGCTTGGCCCCCACACGGACAAGAATGAGCAACTCGGGTGGGGGCTTCCTCTCAAAATTGAAAGCACTAGATGCATATCCCAAA | >lcl\|ORF2<br>MSNSGGGF<br>LRLYMTTQS<br>HLDLDHDVY<br>SCYGAEDR<br>QTGEGCHM<br>HTVNKLSFG<br>NKTLATNQF<br>LSFLTSVCAI | Start 89 Stop 1267 | >Cre13.g579850.t2.1<br>MSGGGFLGKLKALDAYPKI<br>PCAWLSLDAMDISGELHLD<br>CYGAEDKQGDCCNTCDEV<br>VIHKLSFGEPYPGMKNPLD<br>FLSFLTSVCAIVGGVFTVSGI | 679.1 |
| TCAAAAATTTGGGAGATCGTCTATTGCCAAGTTGAATAAGTTGTCAATTTCGCACGATCCTCTTACTTTTTAAGCGCGTCATTTGCGCAATCAGTACTTCTTCAAGATGGCTCCTCTCGCCCTGAACCGTGCTGCTTCTCGCGTCAGCGCT | >lcl\|ORF3<br>MAPLALNR<br>FVEKDTAGC<br>GLGLTALINT<br>AEVVVAPPV | Start 107 Stop 649 | >Cre13.g570851.t1.1<br>MYPTVMKPFEAGSASDTV<br>VAAPVAEPVAVEAPAVASE* | 62.4 |
| TCAAAAATTTGGGAGATCGTCTATTGCCAAGTTGAATAAGTTGTCAATTTCGCACGATCCTCTTACTTTTTAAGCGCGTCATTTGCGCAATCAGTACTTCTTCAAGATGGCTCCTCTCGCCCTGAACCGTGCTGCTTCTCGCGTCAGCGCT | >lcl\|ORF3<br>MAPLALNR<br>FVEKDTAGC<br>GLGLTALINT<br>AEVVVAPPV | Start 107 Stop 604 | >Cre13.g570851.t1.1<br>MYPTVMKPFEAGSASDTV<br>VAAPVAEPVAVEAPAVASE* | 62.4 |
| AAAGAGTTGTAAGGCTTTGCAAAGGACGACAGATGTCAGCGACTGCTAGGGGCATGTGGAAAAAGCGCGGCAATTGAATTCCTTAAGTTGCCTCGGTTCCAACTGGAACGCTACACTGGGGGATGCTCCACCTGCACGCTCCTCAATT | >lcl\|ORF8<br>MWYFPCED<br>QKKAGHPS<br>KAVQMRAL<br>DDEAGEGG<br>KNSYCLNPL<br>VWLVARWV<br>SVPSPQLQS | Start 279 Stop 1283 | >Cre01.g044650.t1.2<br>MASSELKVVEVRSPPAEPL<br>GSIASTVNAPASSEISEAAM<br>ASLRIQESLKQLPSVASDAA<br>LSFLFYLNKKRKAEVNELLS<br>TFGAAPFRVESIWVDETETE<br>SMAPAAVTPVVNYFMKEIIT<br>TKKVTLDSKKNSTSSNANA<br>NGRQLMGRQLIKLSEACKD<br>TSTANKKEWRSSTRYWSR<br>VKVMEASLLSSAECLGRLD<br>DEED* | 114 |
| AAAGAGTTGTAAGGCTTTGCAAAGGACGACAGATGTCAGCGACTGCTAGGGGCATGTGGAAAAAGCGCGGCAATTGAATTCCTTAAGTTGCCTCGGTTCCAACTGGAACGCTACACTGGGGGATGCTCCACCTGCACGCTCCTCAATT | >lcl\|ORF9<br>MWYFPCED<br>QKKAGHPS<br>KAVQMRAL<br>DDEAGEGG<br>KNSYCLNPL<br>VWLVARWV | Start 279 Stop 1151 | >Cre01.g044650.t1.2<br>MASSELKVVEVRSPPAEPL<br>GSIASTVNAPASSEISEAAM<br>ASLRIQESLKQLPSVASDAA<br>LSFLFYLNKKRKAEVNELLS<br>TFGAAPFRVESIWVDETETE<br>SMAPAAVTPVVNYFMKEIIT<br>TKKVTLDSKKNSTSSNANA<br>NGRQLMGRQLIKLSEACKD<br>TSTANKKEWRSSTRYWSR<br>VKVMEASLLSSAECLGRLD<br>DEED* | 115.2 |
| AAACTACCATTGACACGATACATGCTTCAGGTCTCCCGATAGTCGATGTCAAAGTAAAGCTGCTGCAACTAGTCGATCCCCGCCTTTCCTGAGAGAACTTCGTTTCGCTTGTAACTTCAAATACCATCATGTCCTCTATGCTTAGATATTGCT | >lcl\|ORF4<br>MSSMLRYC<br>KYEELRSSL<br>DAYKHMGK<br>DFMSSGRIC<br>SYNACHGS<br>GAGLVVDMI<br>LTSGPVLAL<br>TIKNGVHCS | Start 129 Stop 1268 | >Cre12.g558700.t1.2<br>MAANLRYCFISEWLDPASG<br>TRNRMESQSERTLAMIKPD<br>QARAEAPGSLRAQFGTDKT<br>EVYKGVLPAGDFNSMVEQL | 632.5 |
| AAACTACCATTGACACGATACATGCTTCAGGTCTCCCGATAGTCGATGTCAAAGTAAAGCTGCTGCAACTAGTCGATCCCCGCCTTTCCTGAGAGAACTTCGTTTCGCTTGTAACTTCAAATACCATCATGTCCTCTATGCTTAGATATTGCT | >lcl\|ORF9<br>MSSMLRYC<br>KYEELRSSL<br>DAYKHMGK<br>DFMSSGRIC<br>SYNACHGS<br>GAGLVVDMI<br>LTSGPVLAL<br>TIKNGVHCS | Start 129 Stop 1268 | >Cre12.g558700.t1.2<br>MAANLRYCFISEWLDPASG<br>TRNRMESQSERTLAMIKPD<br>QARAEAPGSLRAQFGTDKT<br>EVYKGVLPAGDFNSMVEQL | 632.5 |
| TTTGTATATGTTTCAGTGCTTTACTGGCCGCCATAGTTTCGCACACTGCAAGCTGACAGTTAGAATTGACGCCTTGGTCCGAGCGTCCTTGCAAACTAATCTTAGATAAGGTCCTGAAACATACGATGCACACCAGATAATGCACATCTCAG | >lcl\|ORF2<br>MAATTLVAV<br>LDHATGTHF<br>GGDKGWSH<br>VAEQPAAPA | Start 197 Stop 679 | >Cre06.g288750.t1.2<br>MATTLVAVGGKGTNPKTTLY<br>LFGRVLKVNYAQPMKIKGG | 236.1 |
| TTTGTATATGTTTCAGTGCTTTACTGGCCGCCATAGTTTCGCACACTGCAAGCTGACAGTTAGAATTGACGCCTTGGTCCGAGCGTCCTTGCAAACTAATCTTAGATAAGGTCCTGAAACATACGATGCACACCAGATAATGCACATCTCAG | >lcl\|ORF4<br>MAATTLVAV<br>LDHATGTHF<br>GGDKGWSH<br>VAEQPAAPA | Start 197 Stop 679 | >Cre06.g288750.t1.2<br>MATTLVAVGGKGTNPKTTLY<br>LFGRVLKVNYAQPMKIKGG | 236.1 |
| GTGCAATCTGCAAAATCTTGCTCCTTTCATTTAGTGACAGCTGCCCAACACAACAGCGAAGTCTCCGGGCAACAGAACTCCGATCAGTTCGTTGTAAACTCACTTCCAGTACTAACGCTGGCCCTGTAGATAAACCTGTCCACGTAAGAG | >lcl\|ORF1<br>MANHFEEFK<br>LPSLLDLPS<br>TFLDLIAEQV<br>FIELMQNMS<br>LIDKGIKYLF<br>GHLCVRRTD<br>ATLERSGGA<br>IVVPRSRFAF<br>LDDAYYKLV<br>EGGSPSPAL | Start 172 Stop 1611 | >Cre04.g229700.t1.2<br>MAGDFAAFEAKMKAANLS<br>TSMGLEKAKSLLVVKDGKT<br>PGHGDIYPSLLGSGMLDAL<br>KHKYFNTNNLWVSLEALAA<br>VAVAPALKGAIPLVKLDDGH | 720.3 |
| GTGCAATCTGCAAAATCTTGCTCCTTTCATTTAGTGACAGCTGCCCAACACAACAGCGAAGTCTCCGGGCAACAGAACTCCGATCAGTTCGTTGTAAACTCACTTCCAGTACTAACGCTGGCCCTGTAGATAAACCTGTCCACGTAAGAG | >lcl\|ORF1<br>MANHFEEFK<br>LPSLLDLPS<br>TFLDLIAEQV<br>FIELMQNMS<br>LIDKGIKYLF<br>GHLCVRRTD<br>ATLERSGGA<br>IVVPRSRFAF<br>LDDAYYKLV<br>EGGSPSPAL<br>VVDNQGCW | Start 172 Stop 1716 | >Cre04.g229700.t1.2<br>MAGDFAAFEAKMKAANLS<br>TSMGLEKAKSLLVVKDGKT<br>PGHGDIYPSLLGSGMLDAL<br>KHKYFNTNNLWVSLEALAA<br>VAVAPALKGAIPLVKLDDGH | 717.6 |
| TATAAACGTTATTGCTTAGTATTTCTTTTCACCGGTGGCTTGAATTACCAATTCCTGTTTGCGTCCTTGCGCTCGTTCTATAGTGCTAGGCTTGAGTGGTATCACGCTTGTCATTTTGAAAGAGGCTTGATGCATTAATTATATCGACGATATGTG | >lcl\|ORF9<br>MDAQTGNM<br>MAVGLSLFN<br>VTAEWLCRE<br>SDFSLRQSH<br>TDGSHAAHL<br>KHKINGAGY<br>VSTDISNRA<br>I | Start 188 Stop 1243 | >Cre03.g211521.t1.1<br>METATAAPPRLATAVSFTKR<br>MSKNYLSKHLTPSHLKNFY<br>GGITGGMGPAGGPPGTVAT<br>PEGIKLGSVSADISTRASCT | 378.6 |
| CGACACATGCCGCTCTTGTGCATGTGCATGTGCATTTTGCCGTCCATCTTGTGAGAACGGATAGGAAGGAAAACAGTTTAAAAGGGGAATGAAGGCGAACAGGTTTCTCAGGATCGCCACACTTTAATAGGAGCTATTGCCGCATTGCG | >lcl\|ORF3<br>MPACTVLGF<br>MRLPPSGS<br>AHSPLPRRF<br>TPEVAQRCF | Start 2227 Stop 2751 | N/A | N/A |
| CATAGAGTCTCGAGCCACTTGCACAGGGAGGACGGGAACCGTATGAGGGTGATACAGCGCCGGGGCTGGCTCACAGGCGTGGGCCAGGCTTTCCGTAAGCCAGATCGCCATTTCGGTGTAAATGTAAATGCTGCTGAACTTGAGGTT<br>GACCGAGCTTGCGCGAAAGGCCCGTCTTTGGGGCCGTAGCGGTACCCGCCATCACCACCATCCGCGGAGGAGGGCGAGGACACGGACGAGGACGAGGATGCCGCCACCTCTGCGGTCGACTGGGTTTGGATTTCTCCCCTCTTC | >lcl\|ORF1<br>MRVIQRRGV<br>LQLLSQSFG<br>AYPQASVQF<br>LLALLLQTD<br>QQGRRVGA<br>QQPHQQLQ<br>GAVRAAAAT<br>QQLAEQTAV<br>GEAFGQRVS<br>TANAAADTC<br>SSSGSDGK<br>DVLHAWEV<br>AQAAASFRF<br>LLTPLDWKE<br>RALAPVLAA<br>SASESSAAA<br>DRLAASEPG<br>SALMAAAYF<br>ATQAVAGAA | Start 43 Stop 2808 | >Cre16.g650700.t1.1<br>MTLSTQRRALLAGFGKARG<br>VLAQLPAAAELMERAAGTP<br>AELPAAASPSASAASAASTA<br>ASSAAKSLGRRGGRLPAEL<br>TISGLSRHLAKQAAQAKAA<br>ITALSASLRRPTSSPTSSAS<br>PALLQPLLQPAGSGAAARN<br>PLLQRAADRHRALLAGGDA<br>FLERLLATPALPGSGRRIRG<br>V* | 135.6 |
| ACTTCTTACACATGATGAGGTTGACATAGAGTCTCGAGCCACTTGCACAGGGAGGACGGGAACCGTATGAGGGTGATACAGCGCCGGGGCTGGCTCACAGGCGTGGGCCAGGCTTTCCGTAAGCCAGATCGCCATTTCGGTGTAAAT<br>CGCACCAGGCGGGACCGAGCTTGCGCGAAAGGCCCGTCTTTGGGGCCGTAGCGGTACCCGCCATCACCACCATCCGCGGAGGAGGGCGAGGACACGGACGAGGACGAGGATGCCGCCACCTCTGCGGTCGACTGGGTTTGGA | >lcl\|ORF1<br>MRVIQRRGV<br>LQLLSQSFG<br>AYPQASVQF<br>LLALLLQTD<br>QQGRRVGA<br>HQQLQQGE<br>RAAAATDGD<br>AEQTAVMRD<br>FGQRVSYGK<br>AAADTQLLG<br>GSDGKASL<br>HAWEVAEQ<br>AASFRPEVL<br>PLDWKEAE<br>APVLAAAIA<br>ESSAAALPS<br>AASEPGLG<br>MAAAYPVD<br>AVAGAAPAA | Start 67 Stop 2823 | >Cre16.g650700.t1.1<br>MTLSTQRRALLAGFGKARG<br>VLAQLPAAAELMERAAGTP<br>AELPAAASPSASAASAASTA<br>ASSAAKSLGRRGGRLPAEL<br>TISGLSRHLAKQAAQAKAA<br>ITALSASLRRPTSSPTSSAS<br>PALLQPLLQPAGSGAAARN<br>PLLQRAADRHRALLAGGDA<br>FLERLLATPALPGSGRRIRG<br>V* | 135.6 |
| CCTTTGCGTCGATGGTGCCGCTTCCATCCGTATCAAAACTGTCAAACACAGACAAGCAGAGTGATGTGTCAAGGCCCAAACACCATCTCGAGTTCCGGGGCGCGCTTCTCCGAACCGCATTGCCTGCCAAGGGCAAGTGTGCCCCC | >lcl\|ORF3<br>MVPVLAHAF<br>PCDHAEAAF<br>CLHALPPTF<br>LIAANVAGL<br>SMIQVGQQ<br>LAEAEAAKA<br>KPAKELTVIT<br>LLWTPRNES | Start 611 Stop 1774 | >Cre11.g468400.t1.1<br>MSLNASASRCRASARQCG<br>LVALAAAAAAALVLGPADEA<br>AGSYAGAARSSGNSGGRV<br>VVVAILRNMQDESEPVSVVK<br>REESVVRHIGETQSDDEAE<br>ESDYLTIPQMEADYPDLVPL | 274.2 |
| CCTTTGCGTCGATGGTGCCGCTTCCATCCGTATCAAAACTGTCAAACACAGACAAGCAGAGTGATGTGTCAAGGCCCAAACACCATCTCGAGTTCCGGGGCGCGCTTCTCCGAACCGCATTGCCTGCCAAGGGCAAGTGTGCCCCC | >lcl\|ORF7<br>MVPVLAHAF<br>PCDHAEAAF<br>CLHALPPTF<br>LIAANVAGL<br>SMIQVGQQ<br>LAEAEAAKA<br>KPAKELTVIT<br>LLWTPRNES | Start 525 Stop 1688 | >Cre11.g468400.t1.1<br>MSLNASASRCRASARQCG<br>LVALAAAAAAALVLGPADEA<br>AGSYAGAARSSGNSGGRV<br>VVVAILRNMQDESEPVSVVK<br>REESVVRHIGETQSDDEAE<br>ESDYLTIPQMEADYPDLVPL | 274.2 |
| CTCCCCCAGGCTGTGGCAGCTGAGGTTAAAGTCGGCCCGCTCAGCACTCCACCACACCATTACCAACAACACCAACATCACACCGTCAACATTGTCAATACCAATCGAGCCCCACAACATGCACTGAACGGGCATCATCACGTAGTAC | >lcl\|ORF5<br>MCTFVCKW<br>VSGLCRLPF<br>RCARRSGG | Start 1073 Stop 1492 | N/A | N/A |
| CAAATTTTGAAGAATCGTAAAATATCGTAGAACATACTTCATTGAAGACAAGCCTTACGCGGTGCAGACAACGCGAGGACGCCCAAGGGGGCTTTCCTTAACATAGTTTTGCTTTTCGCTGAACATAGGCGTGTCACTGGTCCTCCATTATA | >lcl\|ORF13<br>MCTFVCKW<br>VSGLCRLPF<br>RCARRSGG | Start 1506 Stop 1925 | N/A | N/A |
| GCCGAGCTCCTTACTTGGGTCCCACAGGCGCATTGGTTCTTTAACTGCCGTTGTTTTGACTATTACCAAGAGGCGTGTCCCGCTGCTGGGACAGTGTCCTCTAGTTTTGTGTTGCACCGTCTTTTTCTAGCGTTACCAGAATCGAATGTC | >lcl\|ORF7<br>MWPFGPGS<br>FRWPFGPG<br>LFMWPFGP<br>IA | Start 1373 Stop 1832 | N/A | N/A |
| TTGACTATTACCAAGAGGCGTGTCCCGCTGCTGGGACAGTGTCCTCTAGTTTTGTGTTGCACCGTCTTTTTCTAGCGTTACCAGAATCGAATGTCGGCTATGGAAATTGGCGCTGCGGGCGCAGAAAGATTAGTACTCACTCCCGAAATG | >lcl\|ORF9<br>MWPFGPGS<br>FRWPFGPG<br>LFMWPFGP<br>IA | Start 1098 Stop 1556 | N/A | N/A |

| | | | | |
|---|---|---|---|---|
| CTCCCATCCAAAGGGTTTGAGTGAATGTAGATGAGGTGACACTTGCCATGTGTCTCCATTCTTTAATCTTTGAAACCGCGTCTCTGCCCAGCGTCGAGTTGCAAAATGACTGTCACAGTAAAATCCGGCCGAAGTGAGCTTTTGCCCAGC<br>GAGACACGATGGTCGGCAAACTAACTACCGTTCGCGAGAGAAGATTATCCTGTTGAACACATTTTCTATTATATTTGCAAGTATAACTTGTCCCTAATCGCATATGGACATATGCGCGTCGATAAGTTCGTTGCATGCACTTGTGGGCCGATA | >lcl\|ORF22<br>MQPSLPTV<br>AQEEQRGD<br>VAAPDVPAP<br>VPKPSLVTA<br>LEKPPPLAG<br>VVVPRYENY<br>HNRGKNIY<br>DWHTALLP<br>DEYRERMR<br>LHEVLKVNN<br>ACKMALQK<br>VMLGSGRQ<br>RFEPCGLN<br>ANKLRESIN<br>KYQW | Start 2076 Stop 4190 | >Cre03.g185250.t1.2<br>MQTCMSPPGTSRSAALPQ<br>AAETAARTAATPASAAPPAP<br>EAPEAEEEEPPAVEKPPPLA<br>GVDYVFVDHPTFHGRGKN<br>GPFVESEHLELNEEYRERF<br>HYDVDTLAEGKAKCKAAL<br>ITAAADILLMPSRFEPCGLN<br>AASIYEEVLVAAKYQW* | 968.8 |
| ACATTCTGAGGACTCTTTTCCGCTGACGTGGCTATTACAGCGAGGCCATAACTAACTGTGCTTGTGCGGATATTTTATAATATTCTTATTGTTTGCAATAAGGTGCTCATTCTCATCTAGACTGCCGTCTTGACTTTTAAGAATTTTTCCACTAC | >lcl\|ORF6<br>MQPSLPTV<br>AQEEQRGD<br>VAAPDVPAP<br>VPKPSLVTA<br>LEKPPPLAG<br>VVVPRYENY<br>HNRGKNIY<br>DWHTALLP<br>DEYRERMR<br>LHEVLKVNN<br>ACKMALQK<br>VMLGSGRQ<br>RFEPCGLN<br>ANKLRESIN<br>KYQW | Start 209 Stop 2323 | >Cre03.g185250.t1.2<br>MQTCMSPPGTSRSAALPQ<br>AAETAARTAATPASAAPPAP<br>EAPEAEEEEPPAVEKPPPLA<br>GVDYVFVDHPTFHGRGKN<br>GPFVESEHLELNEEYRERF<br>HYDVDTLAEGKAKCKAAL<br>ITAAADILLMPSRFEPCGLN<br>AASIYEEVLVAAKYQW* | 968.8 |
| GCTGCGTAGGATGTTCCTCACAGCAGCTGTTCCTCAAGGATACATTCAGACCTGGGCGGAAGTGCGCGGGCGGCAAGTCGAACCCTTCAGCATCGTGCGGCAGGAGGTCGAAATCGTATCTGAGCGCCTCCGCCACAGCGTCTCAA | >lcl\|ORF1<br>MFLTAAVPQ<br>LKTAAEYFF<br>APELRRRQ<br>ILAGDFLLA<br>DLDHYLAKT<br>VVDDLLDLT<br>KQEGDVAL<br>AREALIQITH | Start 11 Stop 1114 | >Cre12.g511700.t1.2<br>MSLKHALRQAGGLISAVAS<br>LRHSVTTGIPALKTAAEYFF<br>SLNTSVGNKTAILAGDFLLA<br>SYGRHLGIAFQVVDDLLDLT<br>LPTAQSDHAEIAREALIQITH | 615.9 |
| AACAAGTGCTCCCCCCAATAAACAAATATGACAATATAGCAAACTATCTGGCAGATCGGCCTAAATAACGTAAGACTGCATCTAAGTCTGTGCGTCCTCTATCCCAGTTCCTCCATCAGCTTCTAGAATCCCATCAGCATATTTGAGCGATAA | >lcl\|ORF8<br>MSGVKACL<br>LGSRGLAAV<br>GDDTESTTY<br>EVRYRPAGA<br>YRGSGRAG<br>PHQRRDEG | Start 231 Stop 1091 | >Cre12.g511750.t1.2<br>MSGVRACLQPNEGPACPIG<br>AEPSVMGGHASYYGRRAV<br>QQYQFESSLGRKIRVGQDS<br>VPDDD* | 350.9 |
| CTGAGTACACGGCAACCTCTCAACACTTTCGCCTTGCGATTTATGACAATTCCAGCGCTTGAACGTCAATATCTGAACGTAACGCCAGGTGAAAACAAATTGTGACGTAAACTCCAATAAAATTGCACCTACAATTTCACTTAACAAGCTAA<br>AAAGTCGTCCCGTCGTAAATACGTTCCCAGATTCTTCATTCTGACAGCAGCCTGGATTGCAGCAACGGAAAACAGAAAAGGACTGACAACTAACAACCAATAACCGTTAATGGACAGATGCGCGAGTAAGCCCGTCTCCTCCCCCTTCC | >lcl\|ORF9<br>MSSGLLEM<br>YSAAEPPQS<br>NTVLSPLVL<br>SAVRSWSFT<br>QPEAAVYS<br>GIDICLVGDS<br>FGCYETDA<br>MGHVGLTP<br>PGPIAAAITR<br>FCKQYARV<br>RKRGLEEA<br>GGASSKAP | Start 182 Stop 1861 | >Cre12.g508550.t1.1<br>MSAVCSQALGAAAVRVCA<br>STASASTSSTYLWQQRQER<br>GFQAALGAANRLRQPSAII<br>IRKKYEQGEQLTMVTAYDY<br>GMDAVKLEGGSPARVEAA<br>GECSGQVLVYHDLLGMMS<br>KAQLAAAAAAAGGGGPAG | 550.4 |
| CTGAGTACACGGCAACCTCTCAACACTTTCGCCTTGCGATTTATGACAATTCCAGCGCTTGAACGTCAATATCTGAACGTAACGCCAGGTGAAAACAAATTGTGACGTAAACTCCAATAAAATTGCACCTACAATTTCACTTAACAAGCTAA | >lcl\|ORF5<br>MSSGLLEM<br>YSAAEPPQS<br>NTVLSPLVL<br>SAVRSWSFT<br>QPEAAVYS<br>GIDICLVGDS<br>FGCYETDA<br>MGHVGLTP<br>PGPIAAAITR<br>FCKQYARV<br>RKRGLEEA<br>GGASSKAP | Start 182 Stop 1861 | >Cre12.g508550.t1.1<br>MSAVCSQALGAAAVRVCA<br>STASASTSSTYLWQQRQER<br>GFQAALGAANRLRQPSAII<br>IRKKYEQGEQLTMVTAYDY<br>GMDAVKLEGGSPARVEAA<br>GECSGQVLVYHDLLGMMS<br>KAQLAAAAAAAGGGGPAG | 550.4 |
| TGTAATGTTACTACCTAAGTCAATCTAGGATGACGCGCTTCCAAAACATACGTCCACATCATATACATGTTTATGAATCCTTGCTCAGGAAAGTTATCAAAAATTCGGGAGCTTTCATTTAAATGGAGAAGTGGCGCCAGGACGGTGGCGATT<br>ACATCGGCAGCATCGGCCGGTTTTGCGGCAGATTTCGGCCTCGTGATTACTGCGGCGGGTCTTTGTGCGCCCCGGCTTGAAGGGGGAGTGGTTTGGAGAAATGGTGGCTGTCAGGGATGATGGGTTCCCAACCCCACACACACGC | >lcl\|ORF25<br>MEKWRQD<br>SHGTAGSA<br>GRTGKSCR<br>PGRTDNMV<br>HPQLQQPQ<br>TGSGATTSS<br>PPATSAAM<br>PLPAGHTG<br>SSFTSAGGT<br>TAALSTFGL<br>AGLPMIPHV<br>PLQQQQPQ<br>VAAAAPMV<br>LFHQTAAVE<br>PQIPELPWT<br>PMYGTLSM<br>VSTVAASFP<br>SDGPGARR<br>EAVAAPAGP<br>DMFILGGSS<br>SVEALMAM<br>LTPSRAQQ<br>CGRREDMK<br>HHPTQGHH<br>GFLVDNPM<br>LDGIMDKA<br>NGGGIGSF | Start 123 Stop 4055 | >Cre02.g108350.t1.2<br>MDKDYDREASPTLSTTDTG<br>TEIAASLPGRTGKSCRLRW<br>SPAAANAAAAAAAAAAAAA<br>APATPTVTASGGASGGTISS<br>AAAATPWGPGAARPAPSLA<br>SAGPAGAQKAAAPPSAEQ<br>VPPRPLSELPTAAQSAAAS<br>RDRDRDRDRDRDRDRDRP<br>VIAAGSSPSRTTADGFHMS<br>HSSGSGGSGGLAQSNSM<br>QPMQVSMQAMPGSNPAA<br>MYVSQLMAQQPQPLQQH<br>PSAATATPLHALAAAVPLSQ<br>GPDGDFTSRHMPHHAYTH | 234.6 |
| CATTGCATTCTGGACTGCATCCTGTGCACTCGTATCGTATGTAATCTAAATCAGCGCATTTTGTGCAAGGTCCAGAAGGGTGAGAGTCGGCGCGACGGCGGGACACTCATACACTGTATCGTCCTCCACAGAGACATAATGAGAATATCC<br>CCCAGTGGGAATGCGTCCGGTCTCATCACCCGCGGGTCGAAGTCTGTCGGGAAGGTTCTATAAGGGTTTGCGGAGGACCGCCTGGTGGTCCTGTGGTGGAAACATCGGCAGCATCGGCCGGTTTTGCGGCAGATTTCGGCCTCGT | >lcl\|ORF1<br>MEKWRQD<br>SHGTAGSA<br>GRTGKSCR<br>PGRTDNMV<br>HPQLQQPQ<br>TGSGATTSS<br>PPATSAAM<br>PLPAGHTG<br>SSFTSAGGT<br>TAALSTFGL<br>AGLPMIPHV<br>PLQQQQPQ<br>VAAAAPMV<br>LFHQTAAVE<br>PQIPELPWT<br>PMYGTLSM<br>VSTVAASFP<br>SDGPGARR<br>EAVAAPAGP<br>DMFILGGSS<br>SVEALMAM<br>LTPSRAQQ<br>CGRREDMK<br>HHPTQGHH<br>GFLVDNPM<br>LDGIMDKA<br>NGGGIGSF | Start 226 Stop 4158 | >Cre02.g108350.t1.2<br>MDKDYDREASPTLSTTDTG<br>TEIAASLPGRTGKSCRLRW<br>SPAAANAAAAAAAAAAAAA<br>APATPTVTASGGASGGTISS<br>AAAATPWGPGAARPAPSLA<br>SAGPAGAQKAAAPPSAEQ<br>VPPRPLSELPTAAQSAAAS<br>RDRDRDRDRDRDRDRDRP<br>VIAAGSSPSRTTADGFHMS<br>HSSGSGGSGGLAQSNSM<br>QPMQVSMQAMPGSNPAA<br>MYVSQLMAQQPQPLQQH<br>PSAATATPLHALAAAVPLSQ<br>GPDGDFTSRHMPHHAYTH | 234.6 |
| GTAAAGTTTGCCTAACTCCGACCAGGCTGCGCCGCTAGGCGATGCTTCACCGTTAGACCATCGTTTTCCTCAACGAGCCCTGATTGCGGTGACTTGTTTCTATTACAAGACAGCTATACAGGTGCAACGGCGGACATGGGCAGCGGAC<br>ATAATTGTAGCTAGGGCTGATTTAACAGCGTGAGCGGACATTGTATGGTTGAATAGTATAGGATTGTGTTTGGTAGGACGGATTAGATTTAAGAGTGGGAAGGGGTGTGAGTGGGAGCCTGTTGCCATTAGCATTGGTGCCCGCTGGAGTG | >lcl\|ORF1<br>MGSGRGLR<br>IKLRGQSVD<br>VAASTEAVP<br>APLVLLDPA<br>PVSAPAPPT<br>AAVALVPTP<br>ILFGSQTGT<br>VISSSTGDG<br>SNYTRFMY<br>ALKAAVSHA<br>AAAGVAAK<br>TAKAAAEAE<br>EVPPPEALP<br>PSGAEDCT<br>PTAATVGLP<br>PDSSMSYA<br>NFPKREETV<br>GRARGVRE<br>RKVIHLELSI<br>LQPAAGAA<br>LLAEHATDA<br>HAPFEALLD<br>RLGVATTWL<br>VMVGPGTG<br>YKEELEAFK<br>HVYVCGDG<br>VWS | Start 136 Stop 4047 | >Cre16.g683550.t1.2<br>MKIGVAVAGGFGALLLLTPL<br>GNAILARLKQQRAAPPPPA<br>VALRKKQEGKPLVGVKFTLL<br>PVAAVSVAAPSASVPAPSTG<br>AAAATGGNAILARLKQQRA<br>NSGAFFVALRKKQEGKPLV<br>VTAGPSGPSRRSVEMMTG<br>PNADSAESLTVAAAAAAPE<br>GDGDPPENATNFFVALRKK<br>MAAVSIDAPSATAAVKPAAV<br>AAKDAAEKEAAEKETAEKA<br>LAEAEASAAALAAAAQHPE<br>LLSPRETEDGGDDMLLTPR<br>GLLLGLAPVGVDTKGAPAL<br>PGDAIGVLPANHPDLVANL<br>SEKRTLLFLSAKGGKDAYA<br>LASPWTTEGISNPANPVWV<br>LSALHVAFSRAQETKVYVQ | 766.5 |
| CGTCTATACATCCAGGCATTTACCTGACGGAGAAGCGTATCCGAATGGCAATAAACGAGTGTGCAGACTTCCTGCACAAGAAATGCAAACACATACCATGGTGCTAGAATTGTTTACTGTGTTAGCCCATATATACTCATCGCTTCATCGTTG<br>GCCTTGCCTTGCCATTGTGCGCTTTGCCATTGTGCTTTGTTATGGCCTGCGCCCAGGAAGCTGACTAATACATTCTTGGCCTCAGTATGAAAGCTGGGATGTCAGCCTGAGCCCTTGGCGTGGGACTAGACTTCGCTGGCGCAGAGTAT | >lcl\|ORF1<br>MATQNGPF<br>AKASPTTSS<br>PAPEPLKEA<br>LARFRKQRT<br>SEPAAKPPP<br>TCASLNELG<br>GAPPPPPA<br>VEADEVEGV<br>DVAEAAAAK<br>AKAAAEKA<br>DAARRKDE<br>PPPVEAPPS<br>AAGEPASLL<br>DGETALGD<br>TSSRRRTGD<br>VDTKGAPPL<br>SAEKPFWS<br>NHPDLITNL<br>SVGYALSH<br>YAHEISEHQ<br>ARGATHLSV<br>YVPIFLRRA<br>APELLGEAV<br>VYVQDLIKQ<br>LSEQDATAF | Start 172 Stop 3849 | >Cre16.g683550.t1.2<br>MKIGVAVAGGFGALLLLTPL<br>GNAILARLKQQRAAPPPPA<br>VALRKKQEGKPLVGVKFTLL<br>PVAAVSVAAPSASVPAPSTG<br>AAAATGGNAILARLKQQRA<br>NSGAFFVALRKKQEGKPLV<br>VTAGPSGPSRRSVEMMTG<br>PNADSAESLTVAAAAAAPE<br>GDGDPPENATNFFVALRKK<br>MAAVSIDAPSATAAVKPAAV<br>AAKDAAEKEAAEKETAEKA<br>LAEAEASAAALAAAAQHPE<br>LLSPRETEDGGDDMLLTPR<br>GLLLGLAPVGVDTKGAPAL<br>PGDAIGVLPANHPDLVANL<br>SEKRTLLFLSAKGGKDAYA<br>LASPWTTEGISNPANPVWV<br>LSALHVAFSRAQETKVYVQ | 765.8 |
| CCTTACAGATCGAGTACAGTAATATTCCTGCAGAAAATCCTTGCGTGAACAGTATTTCGATATCGGAGAAATGGACACTGAGTTGGCAAGCCTCTCCAAGGAGGAGTTGATAGCAAAACACAAAAAGCTCAAGGATCAGGCTGACTACGA | >lcl\|ORF2<br>MDTELASLS<br>QIEQEEEYIT<br>KEKVDLENF<br>EYILNRLQK<br>NMLEMEEE<br>NDWPRVYS<br>AITGSRRER | Start 71 Stop 1045 | >Cre06.g282900.t1.2 CDS<br>ATGGAGGACCTTGCGGCC<br>CAGAGGATCTCAGGCTCG<br>CCAACTCAAACGGGAGAA<br>GAGAAGGTGGACCTGGAG<br>TGATGAGCGAGAAGGTGG<br>GCTGTCGCTGCAGAAGGA<br>ATGCTGGAGATGGAGGAG<br>AGGGTATCACGCTCAGAG<br>GTTCGACCGCGTGCTCAG<br>CACGGCGCGTCACCGCC<br>CCAAGTGA | 428.3 |
| ATTCATAATCTTCCAATACCAAACGGTATTTATCAATACCACCTTTATCGTTGGATAATTTCCTTTTTCGCAACTGCTTGCCAACTGCAAGCTTTGTTACAACTTTGGTATCTTAGACCAACCGGTCAAAAGACCTCGGAGTCGAGTACTCCAA | >lcl\|ORF1<br>MASPRNPQ<br>RGITPPSRM<br>NAAQDADN<br>LREERERND<br>RIIARTDDM<br>AGPPPGGP | Start 206 Stop 1009 | N/A | N/A |

| | | | | |
|---|---|---|---|---|
| CATTCGGTAATCTTGCAGGAATTGCTCTTCGCACTTTCACCTTAAAGTCCTTACTTAGCAGAGCTGCATCCCTCGGCTGCTATAAGCTGTGCAACATTAAAATTATAAGCAAAAAATATGAATATAATTTTCATGCTACAGTGCGGGAGTTTGTT | >lcl\|ORF1<br>MADILFGMV<br>GNNGDLGE<br>TQSLFILTKY<br>LRGAFIADRI<br>QMAENFNVS<br>TLDYVYKEIS<br>VDPPDRGDA<br>L | Start 277 Stop 1332 | >Cre06.g267550.t1.2<br>MADLFGARPEPNLCWQDR<br>ITTVNVRAATSRIKGSTQSLY<br>QGNLGTFFISNVRVVWYAN<br>LAGAASGILPSEQDVEIVEP | 525.8 |
| AGCAAGACGTAGAAATTGTGGACCCGCCTGACCGCGGCGATGCGCTGGCCCTGTATTACGCTGATGCAACGAAAGGCGTCGACCGCGATCCCGTCTACTGTGCCGAGCTGGGGCTTGCCATTGAACAACTAAAAGAGGGGCTCACC | >lcl\|ORF4<br>MALSCLNAR<br>LLLVGNIPEL<br>DAFDWCPG<br>STTSETLVET<br>NGIPIPAVLA<br>VVEVNGADK<br>NRRDLLMRI | Start 479 Stop 1444 | >Cre06.g269650.t1.2<br>MLTKRLPAAAGLRSSLRRN<br>SAEYKYIIKRKDSLDWCPG<br>HEISAEAVERAEHRAVAVST<br>TLKATAPVAAKKALDVLTVSE | 214.9 |
| CTCCCTTTTCATATTCGCACCTTACACCACGCGCCTGAACGTGCTGCAACTAGCTGACACTGCAGAAGCGTCTCACCTCACAAACAATATGCAGACGTGCAAGATGGCCCTCCGGGGTCAGCGCAGCCAGTTTGCGGGCAGCCCCAA | >lcl\|ORF1<br>MQTCKMAL<br>KEKLDKSIIV<br>TQGWETLAE<br>NQEVKPPTE<br>KLATGIKMVI<br>QNR | Start 89 Stop 850 | >Cre06.g272850.t1.2<br>MQSSTMTLRGQRSQFGAK<br>SVYVTKNSLMKVAVSQTKG<br>DQLYATIARLAKQPAQKLAV | 359.4 |
| CTCCCTTTTCATATTCGCACCTTACACCACGCGCCTGAACGTGCTGCAACTAGCTGACACTGCAGAAGCGTCTCACCTCACAAACAATATGCAGACGTGCAAGATGGCCCTCCGGGGTCAGCGCAGCCAGTTTGCGGGCAGCCCCAA | >lcl\|ORF1<br>MQTCKMAL<br>KEKLDKSIIV<br>TQGWETLAE<br>NQEVKPPTE<br>KLATGIKMVI | Start 89 Stop 802 | >Cre06.g272850.t1.2<br>MQSSTMTLRGQRSQFGAK<br>SVYVTKNSLMKVAVSQTKG<br>DQLYATIARLAKQPAQKLAV | 362.5 |
| AACGGTGTGAAGTTCGTGAAGAGGAGGACGGCTCCGCTGCAAGCGGGCGTCCCGTTTATCCTGGACTGCTGGTGTGGACCGTACTCGATAGTCGGTTGCGGCCAAAGGCTAGTTTTTGCGCAATGCTCGTCCCATTAGTGTTTAGTTC | >lcl\|ORF11<br>MALDKHEPS<br>GGQIITFFSD<br>GNVVDRGKI<br>GESPRRAEC<br>ETERPGVGS<br>SSVSGLGGS<br>RLEDYGSPE<br>TAPSDTLIVL<br>IYKVDASEGI<br>AALTRGRKC | Start 459 Stop 1928 | >Cre07.g322550.t1.2<br>MEPTEGIIHYPSGNIAVVACI<br>PRTSQEVISLDVTECISVRFT<br>LRKALRNLDPASDLSATVSE<br>ADKIRHKARRAKLPWMRLK<br>MLLLFYEGRLVAATNNIRTEA | 535.8 |
| CATTTATCTGTCCTGCACTGGCCTAAAGGGCTTTACCCTTTTACATCGCGTGTAGAGCGGGCGCAACACGGTCAAGCTTCACGTTTAAGCTCGCTCGCCTATCGTCAAGGCTACCGCCTATTCGGCTGTCATGTTTGGTGTAGTTTCAACA | >lcl\|ORF15<br>MALDKHEPS<br>GGQIITFFSD<br>GNVVDRGKI<br>GESPRRAEC<br>ETERPGVGS<br>SSVSGLGGS<br>RLEDYGSPE<br>TAPSDTLIVL<br>IYKVDASEGI<br>AALTRGRKC | Start 1899 Stop 3368 | >Cre07.g322550.t1.2<br>MEPTEGIIHYPSGNIAVVACI<br>PRTSQEVISLDVTECISVRFT<br>LRKALRNLDPASDLSATVSE<br>ADKIRHKARRAKLPWMRLK<br>MLLLFYEGRLVAATNNIRTEA | 535.8 |
| CCACAGATTGCCTTCAAAGTACCGGGTTAGCACCGGGCTTTGCTTGTGAAAGGGGGGCAGTGTTCCTTGATCATTGACCGTCGTGCCCTTTCGTAATTTGGTGCTGATCCCTTGCCACTGTTCAATAGTGAGGCACGACTAGCCCGCGC<br>CAGCATGGTAGAGAGCGGGTTTTCCGGGGCCTGCTGCTTCTAAATAGCGGAGAGGATGAGTAGTGCGCTTTGCGTGTGAAGGTACTGTCGAGACTGTAACTAATGTGAGACCCTCCTCACGTAATTTGCGGAATACGCCATCATATGGG | >lcl\|ORF5<br>MDPAAAVAA<br>TCLVQTLRT<br>VTKQALLEA<br>SSNQQAQIC<br>AGASAATSF<br>MFIALAETSA<br>EAREQSPGN<br>GELYDPGQE<br>AQIAEGCSK<br>GPELQAKHI<br>YLDTLILKLL<br>MHILTSANA<br>AGGVDADD<br>QVVDEEDLN<br>LKEGFWQY<br>SFGAIYYTM<br>WTPLLEALR<br>VLEKADKRF<br>LKKFGDDVL<br>GMAKHFNN<br>VAEIMAGIIQ<br>TLWLQSLPL<br>GGTTYVAGF<br>GRVPE | Start 155 Stop 3622 | >Cre14.g633750.t1.1<br>MDPAAAVAAIAANPAAFTEL<br>SDLKAPVLWDKCSPAVQTG<br>FVNMLVTMLGSGVKDVTVG<br>SLEPQTRQLAVEFLVSLCEA<br>TWITDAQWQKRAAVFICLAC<br>CAAIVNFSEGVEADVLPPYL<br>DRFRDHARQVLAYMQQVQ<br>ILEEKATAVNMLSCYAEELKE<br>KEPEGEIQAVQLDSIGEIVEV<br>LPLVESLLMTRYGAMLTDKS<br>IRHPEAKSEDNDMATDNAVA<br>PVGVRMAQLLHRLQGAVPA | 1738.8 |
| CCACAGATTGCCTTCAAAGTACCGGGTTAGCACCGGGCTTTGCTTGTGAAAGGGGGGCAGTGTTCCTTGATCATTGACCGTCGTGCCCTTTCGTAATTTGGTGCTGATCCCTTGCCACTGTTCAATAGTGAGGCACGACTAGCCCGCGC<br>GTACTGTCGAGACTGTAACTAATGTGAGACCCTCCTCACGTAATTTGCGGAATACGCCATCATATGGGCCAGCCGTGTCATGTTAACAAGGGCCTTGTCCAAAAAGACCAGGGACATGGCGGGGAACAGCGGACCCGAGATCTCGGCT | >lcl\|ORF5<br>MDPAAAVAA<br>TCLVQTLRT<br>VTKQALLEA<br>SSNQQAQIC<br>AGASAATSF<br>MFIALAETSA<br>EAREQSPGN<br>GELYDPGQE<br>AQIAEGCSK<br>GPELQAKHI<br>YLDTLILKLL<br>MHILTSANA<br>AGGVDADD<br>QVVDEEDLN<br>LKEGFWQY<br>IAAAERGVQ<br>VEMVDKSLL<br>AIEAENELEE<br>SRSAEERRIA<br>AVYGLGVMA<br>AALGRVLTH<br>EAQDPRIVP<br>EAISSVLSTF | Start 155 Stop 3541 | >Cre14.g633750.t1.1<br>MDPAAAVAAIAANPAAFTEL<br>SDLKAPVLWDKCSPAVQTG<br>FVNMLVTMLGSGVKDVTVG<br>SLEPQTRQLAVEFLVSLCEA<br>TWITDAQWQKRAAVFICLAC<br>CAAIVNFSEGVEADVLPPYL<br>DRFRDHARQVLAYMQQVQ<br>ILEEKATAVNMLSCYAEELKE<br>KEPEGEIQAVQLDSIGEIVEV<br>LPLVESLLMTRYGAMLTDKS<br>IRHPEAKSEDNDMATDNAVA<br>PVGVRMAQLLHRLQGAVPA | 1749.2 |
| ATATTGCGGATCCGCCCAACCGAAAATACATGGTGTTTCTTGGAGGCGCGGTGCTGGCGGATATTATGAAGGACCATACGGACTTCTGGATCACACGGGAACAGTACCAAGAGGACCCAGGCAATGCTCTCAAGAAGTGCGGAGTTGC | >lcl\|ORF7<br>MAMSLRCG<br>ASCVKLGSC<br>TFLSATVLVI<br>VLFLLFEMG<br>GTVFLEQIAI<br>TRFGSATLG<br>GPTALQTVA<br>LITQRLGLSD<br>SSINLAFLQS<br>LSQGGEFAF<br>ANKLDELYP<br>GFSPAGQM<br>GDATRSVVL<br>SDLRHAAEL<br>EETIEARTQA<br>AAAERDAKN<br>AAEPSLPTG<br>EPAGGLMDS<br>TTPAAEVAM | Start 492 Stop 3245 | >Cre16.g687450.t1.2<br>MLVPLGLDFLTFLSATVLVIP<br>YAFGIGSLQVLLCTGIFTAFA<br>VLLPLITTHGTDLEGASPVSI<br>YRTQVEADIRPFKGLLLGLF<br>NQLLIIVVVLSMALTPFLAES<br>VANMLESPLATPGGGGEPF<br>ELEEAGASRVIIRSVEAGLAL<br>QVAAALSAEAAQPSPRQPV<br>EAGPVAVVDADSQARVRA* | 866.7 |
| ATGCTCCTACCGCTCGGGCTGGACTTCCTCACATTCCTATCAGCCACCGTCCTTGTTATTCCGCTCTTTAAATCACTAAAGCTTTCGCCTGTCTTGGGCTTCTTGTTCAGCGGCGTGGTGCTGAAACAGCTTGGCCTCTTTCAGGACCTC | >lcl\|ORF1<br>MLLPLGLDF<br>QDTERLGEL<br>AFALPVGHG<br>QLLRERGEL<br>ASPVSLLTLL<br>LCLLTVAGTS<br>LLGLFFVTTC<br>RSESIRTGFV<br>PFLAETGKV<br>TGVSDPVVIC<br>GRKAGFKVL<br>FPSVPIYACA<br>IDLNFLKRGI<br>ARSAAAAAA<br>AAANGSVAA<br>VSAPPPPPF<br>AEAVTATTTT | Start 1 Stop 2484 | >Cre16.g687450.t1.2<br>MLVPLGLDFLTFLSATVLVIP<br>YAFGIGSLQVLLCTGIFTAFA<br>VLLPLITTHGTDLEGASPVSI<br>YRTQVEADIRPFKGLLLGLF<br>NQLLIIVVVLSMALTPFLAES<br>VANMLESPLATPGGGGEPF<br>ELEEAGASRVIIRSVEAGLAL<br>QVAAALSAEAAQPSPRQPV<br>EAGPVAVVDADSQARVRA* | 867.5 |
| TTCACCCGTTCCTAGCGGCAGTCAGTGCGATGGAGACTAGGCGTTGGTTCGCGCAAGCGACAGCACATCGGCAGGAAATCACGCGTCAGGCCCAAGCTATGGACCGGATCATCCGTGCATTTATGTTCTTGGAACACATATGAACCTTA | >lcl\|ORF3<br>MQYPVVQSI | Start 30 Stop 143 | N/A | N/A |
| CTCTAAAACGTGATTAGGAAAGCGGAACACCTAAAACTTTAACCAGTGCAAGACAAAAGGATGTAAATTAGCGTGCTTTAGAGGCAAAGGTTTGGTCCCCTGCATGGGGTATGACCTGCTGCGCCTTATTTGTTGGCTTGCGCAAAGTTCC | >lcl\|ORF1<br>MIPKAATQA<br>ESEEQKLLR<br>GSLASQFSA<br>CVIPVIQVAL<br>MQRMHATE<br>NSVAVETVD<br>IPAKQRIFPA | Start 196 Stop End | >Cre04.g215550.t1.2<br>MTRLHAVPGTNSRVAVDNV<br>GTQAADFAVARGVDLSRLA<br>ERMHATEGGYNLTRLGGLH<br>QDGGAECSPVAIAEIPAVGR<br>VCERLAGHARPFTHSQSEL | 307 |
| TTCTATGCCAAATAAATACACAGCTTAATGCATCGCTCTGCACAATTTGTGGTGCCGTCCTGGGCCTGATCTTATCCGCTCCAGAACCGCCCGAGCGGCCTGGTTATTGTGCGCACAGCAGGCTAGTAACTTCGCTCTTCCCTTGCGGCA | >lcl\|ORF16<br>MPRPGIRPS<br>AALAVLLLTF<br>LVARHVLHE<br>SAGGSIAQV<br>NGVQTGAVF<br>RTGDGLNRA<br>RHRAEHAAA<br>QAARAVLER | Start 175 Stop 1242 | N/A | N/A |
| TTCTATGCCAAATAAATACACAGCTTAATGCATCGCTCTGCACAATTTGTGGTGCCGTCCTGGGCCTGATCTTATCCGCTCCAGAACCGCCCGAGCGGCCTGGTTATTGTGCGCACAGCAGGCTAGTAACTTCGCTCTTCCCTTGCGGCA | >lcl\|ORF1<br>MQTMRAPA<br>EMPLNTYSN<br>YGVIPPGTKI<br>DPETGKEDF<br>VATGTGIAPF<br>ALAKAYPSQ<br>MYFCGLKGI | Start 175 Stop 1215 | >Cre11.g476750.t1.2<br>MQTVRAPAASGVATRVAGR<br>CHIIIETEGKIPFWEGQSYGV<br>PTGKVLLLPADANAPLICVAT<br>KVEEYADEIFDLLDNGAHMY | 655.6 |
| AACCCTACGCGAGCGCAGCAATGAAGCGCCCAATCATCAAGTATTGCCCTGTCTATCTTCAGGTTTTACTCGATAGTATGGCTCTAAGGAATCTGCGCCGCCCTCCCACGGGGCGATCACAAGCGTTTGGCCCATCGATTCGACCTGCA | >lcl\|ORF8<br>MKRPIIKYCF<br>GRRELLQLG<br>LEFQLVPPAS<br>VRNAQSLRF<br>QTALGPVEL<br>WEQYGSELI | Start 21 Stop 845 | >Cre13.g564050.t1.1<br>MALNTLERRPACSAFSRRR<br>SIPGMASQPYLDVDFQLLPI<br>AGSVLQVALPPRQTALGPVI | 277.3 |
| AACCCTACGCGAGCGCAGCAATGAAGCGCCCAATCATCAAGTATTGCCCTGTCTATCTTCAGGTTTTACTCGATAGTATGGCTCTAAGGAATCTGCGCCGCCCTCCCACGGGGCGATCACAAGCGTTTGGCCCATCGATTCGACCTGCA | >lcl\|ORF8<br>MKRPIIKYCF<br>GRRELLQLG<br>LEFQLVPPAS<br>VRNAQSLRF<br>QTALGPVEL | Start 21 Stop 698 | >Cre13.g564050.t1.1<br>MALNTLERRPACSAFSRRR<br>SIPGMASQPYLDVDFQLLPI<br>AGSVLQVALPPRQTALGPVI | 203.4 |
| ACCAATCGAATTAACTTGCCCACAGGTTCCCATACTGTAGTGCTGCGTACGTTCGCCTCGTAATTGCATTTGCGAACAGTAATCTGAAAAGCAAGTAAAGGCATTAAAAGCGGTGACGACCATGTATCCTGCGTCACAGCCCTTGCGAGAT | >lcl\|ORF1<br>MYPASQPLF<br>DRHNHPARI<br>TAFEARIWQ<br>GGHQHHPS<br>KAPEGYSVE<br>TYWLSYDRA<br>LTATPFPSGC<br>DNPQFTFSA<br>ILSKKDKEFC<br>IRVLHGSISV<br>VTWMDANW<br>SLHLFTPDTI | Start 122 Stop 1876 | N/A | N/A |

| | | | | |
|---|---|---|---|---|
| ACCAATCGAATTAACTTGCCCACAGGTTCCCATACTGTAGTGCTGCGTACGTTCGCCTCGTAATTGCATTTGCGAACAGTAATCTGAAAAGCAAGTAAAGGCATTAAAAGCGGTGACGACCATGTATCCTGCGTCACAGCCCTTGCGAGA | >lcl\|ORF4<br>MYPASQPL<br>DRHNHPAR<br>TAFEARIWQ<br>GGHQHHPS<br>KAPEGYSVE<br>TYWLSYDR<br>LTATPFPSG<br>DNPQFTFSA<br>ILSKKDKEF<br>IRVLHGSISV<br>VTWMDANW<br>SLHLFTPDT | Start 122 Stop 1876 | N/A | N/A |
| CCAAGCCTGGCTTCGTATTCACCACTTGCGAGGGAGCCACTGGATTTTACAAGGATAAGCCACACCTTTCATCTGCACCGACTTCCGGAGCGCCTCTTTTGCCTGCGGCGCCTCGCGAAAGGAAGGTGGGCCCAGCCGCCGTGCCC | >lcl\|ORF9<br>MAAKMEMA<br>RSNKRAASL<br>LEPALDTLR<br>VLYKTLASA<br>WCQGGGSS<br>TPKSTDGRV<br>GKVRVTSAL<br>KLTRLLSIEA<br>VPAGAVRPA<br>GAGSSAPTA<br>PSARPTDSG<br>QQQQQQAE<br>SAPDKAAAS<br>TTSGSSQA<br>ASAASPAAN<br>GLFGGLSFG<br>AGTSLFGAF<br>SAIGGTSSA<br>SSPRVTVAP<br>PSTFGSPAP<br>GVAAPSGG<br>QGIAFGTTG | Start 342 Stop 3608 | >Cre07.g313500.t1.2<br>MELQPAVGALDESEDVGFY<br>KYARPQPADVCAWHAGLR<br>AVVVAPSGVYAADWCPSDA<br>GSGVAPSGAELLTAPPGFVE<br>LDTTVPLDPIPNPLNAEKPP<br>DLEESEGDAEGGTGGRAM<br>SDGDSGSHAVARPGGRRR<br>ASTAFGTSLFGAAAPAAPAL<br>PVAAASSGASGGGFGFGF<br>FGSSSTAPSGGLFGAPAAG<br>STGAPSAAPTPAATKPAAFG<br>ELEDAEDVGFYFKFYGGIDF<br>ADQAVLAAVQGGEGRVSFF<br>GRRLALVSVAADLSVCAVA<br>LSGPYLHAVCLPSWGAIVR<br>LLVATSDGVLRVYGFGHEEL<br>DSESGEEVTSPGITGKAPPA<br>TGLFGAATSAPSTSGGFGF<br>PPQPAPLSALPPGYVEVQP<br>KLASESKEGAAAEASFLREL<br>LFSGGLDGGGGLGGGGRL<br>AARLEEAGALPVGAMRDPH<br>GAGATPPSPTQPQPPVASP<br>PPKPAAAAPPAFNFSSSGF<br>AAAAGGQPPVPSMALFAQ<br>GATGTSAGTAAAAGSAAAS<br>APAAAGSGGGGLFGSFGA<br>AASASGAASAPAAAAPTTT<br>SAPTAPATGAASSPFGSFAS<br>QQGVFSSFGAAPAFGSPAA<br>GAAAPSNPFTGFAQQAAAA<br>AGGFGTPSASSGAPGGTN | 204.5 |
| GCAGTCGTCACGTAATTCATGTATCAGTCTGGACACGCCGCTTTCTGTTCAACTCGCCCATCATGGTGCAGGCGTTGTGGGCAGCCTTGCTACCCACGCGTCCAACAAGCAGGCGGCCTTGCCGATGGGCAGCATTGCCTTTCCAGC<br>ACTGAGCACCTGGTGCCAGTGCAACTACAGACGTGCTGAAAACATAATCTTAACTGGCGTCCGGTGGTCGCTGGCAGCAATTAAGATGCGGGTTCACGCGAGTTAGATTAGTGCTTGAGAC | >lcl\|ORF4<br>MGSIAFPAL<br>GFAGFGTGF<br>RATGGLFGS<br>TGTVGGAQ<br>GPAGSGGL<br>KDKPHLSSA<br>KAPAAAPVP<br>KMEMAMKD<br>KRAASLFDG<br>ALDTLRRDI<br>TLASAGSVL<br>GGSSTAGD<br>TDGRWSPL<br>VTSALRLQT<br>LLSIEAAAA<br>AVRPASVAQ<br>SAPTAATRT<br>PTDSGSSA<br>QQAEHQPE<br>KAAASAAPA<br>SSQAQSSS<br>SPAANIALA<br>GLSFGSTSS<br>LFGAFASSA<br>GTSSAPSTA<br>VTVAPSFGS<br>GSPAPAAPA<br>PSGGGFGV<br>FGTTGPSGF | Start 125 Stop 4429 | >Cre07.g313500.t1.2<br>MELQPAVGALDESEDVGFY<br>KYARPQPADVCAWHAGLR<br>AVVVAPSGVYAADWCPSDA<br>GSGVAPSGAELLTAPPGFVE<br>LDTTVPLDPIPNPLNAEKPP<br>DLEESEGDAEGGTGGRAM<br>SDGDSGSHAVARPGGRRR<br>ASTAFGTSLFGAAAPAAPAL<br>PVAAASSGASGGGFGFGF<br>FGSSSTAPSGGLFGAPAAG<br>STGAPSAAPTPAATKPAAFG<br>ELEDAEDVGFYFKFYGGIDF<br>ADQAVLAAVQGGEGRVSFF<br>GRRLALVSVAADLSVCAVA<br>LSGPYLHAVCLPSWGAIVR<br>LLVATSDGVLRVYGFGHEEL<br>DSESGEEVTSPGITGKAPPA<br>TGLFGAATSAPSTSGGFGF<br>PPQPAPLSALPPGYVEVQP<br>KLASESKEGAAAEASFLREL<br>LFSGGLDGGGGLGGGGRL<br>AARLEEAGALPVGAMRDPH<br>GAGATPPSPTQPQPPVASP<br>PPKPAAAAPPAFNFSSSGF<br>AAAAGGQPPVPSMALFAQ<br>GATGTSAGTAAAAGSAAAS<br>APAAAGSGGGGLFGSFGA<br>AASASGAASAPAAAAPTTT<br>SAPTAPATGAASSPFGSFAS<br>QQGVFSSFGAAPAFGSPAA<br>GAAAPSNPFTGFAQQAAAA<br>AGGFGTPSASSGAPGGTN | 220.7 |
| CATATTACTCTGTATGCGCAACAAGTATGTTGCCAATGCAGAACCGCTACGCTTATCAATAGCTGGCAACCCCATTCAGGCGAGATGGGCCGGCTTGCGGACATTTGGAAAAAGGCTCAGCCGTGGCAGCGGCGCAACCTCATCTTAAG | >lcl\|ORF1<br>MGRLADIW<br>LDEEVVDK | Start 85 Stop 261 | N/A | N/A |
| CTCAAGTTCGTGGAGGTTCTGGAGCTGATCGGTCACAACAACACCACGGGGAAGCGCGCGGGTCTGGCCCCCGAGCAAGACCGCCACGGCCGCGGAATGAGCGACACGCAGCAGCAGCAGGTGCTGAACCTGTTCAAGGAGAC | >lcl\|ORF2<br>MSLSVSHF<br>TLINSWQPH<br>LKNDPVIRK | Start 1162 Stop 1524 | N/A | N/A |
| ATGGAAAAAGGAAATGTGGATGGCATTGTCGACGCTCGTGGTGAAGCCGGTCGTGTTTCTCCTGAGTATAAAGAGCCCTACAACAGAAGACTTGTAGATGGATTGATTCTAACTGAAAGCGGCTCTGGCAGCAAGGGTAACAGTTTCCT | >lcl\|ORF1<br>MEKGNVDG<br>PAPVREASD<br>HIPALGSEV<br>PEASGLVPV<br>VSPEPAEEQ<br>DDADGPAD<br>DGEGGSAP<br>ADVGAGTTD<br>DGAVMGDG<br>GAGAAAPS<br>GAAAAVQQ<br>ASGGGSDG<br>TAQQRSSTT<br>GQPSWRPS<br>RPSGSVEDT<br>GGSSGGLA<br>QASQSSAA<br>AANGGGGA<br>LGMPLSVG<br>TGPQEFVV<br>AGPFYPSG<br>VASFHQPSA<br>VFYVLWPPV<br>GEGEGEGE<br>SSSEDEEEE<br>LGYGGQPY | Start Begin Stop 3837 | >Cre06.g308650.t1.1<br>MDPEVAPPPTLEGADPTEG<br>EPEAAEPEAAEPEPEAAEP<br>EGGGEEEVPEHSGKILTYIS<br>ASAAAAPPAVSESAPGPGS<br>ASGAPSGAASRATSFVGAG<br>PSSSLATAVAAAPPPPLPPL<br>STANICGIRSLPNALNPDFR<br>LKHKLSQTTPSEFVVRAPPA<br>EDSNGSVVVSFDKSRAVSE<br>GGGGGGGSGGAYDGEEG | 354.8 |
| AGCACATCGCGACCGTACATTTACTGCACCCAGTCATAAACCTCAGTTGTCGTACGGTCTGGAGGGTGGCCCGCTGGGTATGCCCCTCAGTGTGGGCCAGAAGCAGGTGGCTCGAATCAACCCCCTGGCCGGAATGTTCACGCAATT | >lcl\|ORF2<br>MPLSVGQK<br>PQEFVVRAP<br>PFYPSGRV<br>SFHQPSAV<br>YVLWPPWV<br>GFATAGAAG<br>AASGSQRR | Start 80 Stop 1018 | >Cre06.g308650.t1.1<br>MDPEVAPPPTLEGADPTEG<br>EPEAAEPEAAEPEPEAAEP<br>EGGGEEEVPEHSGKILTYIS<br>ASAAAAPPAVSESAPGPGS<br>ASGAPSGAASRATSFVGAG<br>PSSSLATAVAAAPPPPLPPL<br>STANICGIRSLPNALNPDFR<br>LKHKLSQTTPSEFVVRAPPA<br>EDSNGSVVVSFDKSRAVSE<br>GGGGGGGSGGAYDGEEG | 350.1 |
| GTTCCTGTGCCCCAGATGGGTTGCGCCGATGGGATCCGCATTGCACAGCTCAAGAACAAGGACATACACGGCAACGAGGTCGCGTCGCAGGCCTATGGGGGCTTTGACGACGACGATGATGGTGGTGACGATGATGATGACGGCG | >lcl\|ORF3<br>MHPKTDDK<br>PPPSPPTPP<br>SRTRRDPH<br>LEKHATAPL<br>IVDTRAGKL<br>PRLTAHSGG<br>AAVVGAVM<br>SGGESENC<br>VGVMCNSL<br>MSYFASSG | Start 1981 Stop 3441 | >Cre06.g282100.t1.2<br>MLAASLRYKPPGRPPLQQH<br>GYLFEVHVWATYNFDAVEK<br>GEGEGEGEGEGEGEGWYT | 48.5 |
| GCAGGCCTATGGGGGCTTTGACGACGACGATGATGGTGGTGACGATGATGATGACGGCGGCGGGGGCTACGGCAATGGCAATGGCAATGAGGATGAGGAACAGGAGCATGACGGCGGCGGCGGTGACAGAGGTACGGGAGGGGG | >lcl\|ORF7<br>MHPKTDDK<br>PPPSPPTPP<br>SRTRRDPH<br>LEKHATAPL<br>IVDTRAGKL<br>PRLTAHSGG<br>AAVVGAVM<br>SGGESENC<br>VGVMCNSL<br>MSYFASSG | Start 1931 Stop 3391 | >Cre06.g282100.t1.2<br>MLAASLRYKPPGRPPLQQH<br>GYLFEVHVWATYNFDAVEK<br>GEGEGEGEGEGEGEGWYT | 48.5 |
| GACAGCCCCTGTGCGGGAATGAACATTAACAGACTCGAGTTTATCGTCAACAGCACTTGTGTCAACGAGGAGCCCAAGGCGCTACGCGCTGCCACCGTCAATGGTACTTCAGTTTACCCGCTTTTCAGCCCCAAGACCTGGAAGGGC | >lcl\|ORF5<br>MFGDLVGA<br>DYYCVCRG<br>QAGGMLSG<br>HENR | Start 1268 Stop End | >Cre01.g028777.t1.2<br>MLGCVGSITHVGIAQPSIDF<br>DDPRSFPEAEGRRHLA* | 52.4 |
| ATGTATAACTTCTACATTTGCTGGCTTTATGTTAATTACCTTAAAAGCCATGTTGTTATGGGCCTCATGTACAAATGTGGTACCAGAATATATGTTGACAGCACTTGTGTCAACGAGGAGCCCAAGGCGCTACGCGCTGCCACCGTCAATGGT | >lcl\|ORF1<br>MYNFYICW<br>GTSVYPLFS<br>NFRMERDS<br>HNLENLFAI | Start Begin Stop 519 | >Cre12.g551552.t1.1<br>MVLTTVACCALLRSAAGIAL<br>APPSPAPPSPEPPTPPSPP<br>TVPLPDLEGPPCHEPTSLP<br>CPCQPPSPSPPPPAPPPC<br>GPNCHEPYDDSLLTAATVP<br>PPPSPPPPPPPSPPPPPGS<br>TATAEPSPPTGVCAGMDLF<br>GSACTYSIFNKKQNCCPTS | 43.5 |
| GGCATGTTGAATTGCACTTGAATAAATTGCAAGAAAAAGGCAAGCTGTATCACTGTCGGAGTAGTTGTCGATTTAGCTTTCAGTTGGTGCCAGTGTTTGCAAGTTTAGGATGGCAACCACATTGTACCGGTCTACAATTAACCTACACAGAC | >lcl\|ORF5<br>MATTLYRST<br>NPSPGIGEL<br>AKKALEQAI<br>RGIKVQVQS<br>DARGKTEEV<br>LSPEDGQW | Start 110 Stop 949 | >Cre16.g650500.t1.2<br>MAATPMRTALNLHRPAAAS<br>KDLELELKLAVKGENYGEA<br>KSSPSQGRYMFAYHITLHN<br>KIGKFMLDAKGPQAV* | 331.3 |
| GGCATGTTGAATTGCACTTGAATAAATTGCAAGAAAAAGGCAAGCTGTATCACTGTCGGAGTAGTTGTCGATTTAGCTTTCAGTTGGTGCCAGTGTTTGCAAGTTTAGGATGGCAACCACATTGTACCGGTCTACAATTAACCTACACAGAC | >lcl\|ORF1<br>MATTLYRST<br>NPSPGIGEL<br>AKKALEQAI<br>RGIKVQVQS<br>DARGKTEEV<br>LSPEDGQW | Start 110 Stop 949 | >Cre16.g650500.t1.2<br>MAATPMRTALNLHRPAAAS<br>KDLELELKLAVKGENYGEA<br>KSSPSQGRYMFAYHITLHN<br>KIGKFMLDAKGPQAV* | 331.3 |
| CCTCTTACTGTAGGTAGATTCATTCGCACCATACTTCAATTCAGGAGCTTCCCAGCCATGGAGCAGCGAACCCAACAACCATGGCCTAGTGAGCTTAATCCTGCCCGCCAGGGCTATGGACTAGAGTTGAAGGACAAGTTTACAGTGCG | >lcl\|ORF1<br>MEQRTQQP<br>VPSNCTVYY<br>RTLVASWAG<br>GDTVGAGIH<br>NFGQKPFM<br>ETLRAFDLA<br>DDGTALGLF<br>CQKYIELVR<br>PEASPLAHL<br>GAVLQLRGV<br>GALHDANG | Start 58 Stop 1638 | >Cre09.g387801.t1.1<br>MGSLEPLPSALNSARQGH<br>TRQPGWEVGSYGYHGDD<br>LAEERAAQRAAVERIPVSPG<br>DSGRFGDVHFQLACQKYIE<br>AADASTGEAAPGAQDAGA<br>REHMLAPGAV* | 478 |
| CCTCTTACTGTAGGTAGATTCATTCGCACCATACTTCAATTCAGGAGCTTCCCAGCCATGGAGCAGCGAACCCAACAACCATGGCCTAGTGAGCTTAATCCTGCCCGCCAGGGCTATGGACTAGAGTTGAAGGACAAGTTTACAGTGCG | >lcl\|ORF1<br>MEQRTQQP<br>VPSNCTVYY<br>DGRKFIETG<br>VRAGLYPTI<br>VSPGITHKL<br>ADSNAAAVA<br>QCPVVLADA<br>SAASLESSL<br>SDIGVGGAA<br>LLSQQRSQ | Start 58 Stop 1548 | >Cre09.g387801.t1.1<br>MGSLEPLPSALNSARQGH<br>TRQPGWEVGSYGYHGDD<br>LAEERAAQRAAVERIPVSPG<br>DSGRFGDVHFQLACQKYIE<br>AADASTGEAAPGAQDAGA<br>REHMLAPGAV* | 493.4 |

| | | | | |
|---|---|---|---|---|
| ATGGCTGACGTCTTCAGCACATCCTTTGCCGCGGAGTCGCATACAGCTTGGGCTGCAAGCCAGTACGCATGGGACCCATCGCAACTGCAAGCTTTGCCGCTGGCTGCAAGCCCAGCTTCGGGATCTACTGGAGCTGCTTCAGAGCAT | >lcl\|ORF1<br>MADVFSTSF<br>SEGEIFWTG<br>VGWQESTR/<br>YQRYRVCEF<br>QHNARRRKI<br>MNDKVMTN<br>NQFQAMLQ<br>DYLEDPRAA<br>PQLGMMSL<br>ADLAAALNK<br>QQQQQQQ(<br>ASSSVLPMD<br>TANMGNTVS<br>LVAPPANAA<br>DQLPVDLYQ<br>DGFGDDGT(<br>APMGAKPH<br>TPGVELCAR<br>ALERAGRK(<br>AAELSLCAG<br>TEDRLACVD<br>DFSMARVST<br>ALLAWIAFS(<br>SARSPLDGA<br>SGAITAIENT<br>AIMFVVSVL(<br>LLGLAIQFLF | Start Begin Stop 3993 | >Cre06.g300600.t1.1<br>MDVFNSGFQHTAGNLQQQ<br>LPGSGGREQRARLKGPLVC<br>NASSGRRQQRRMVLALPN<br>AGGGWPQQQRPLGPLPM(<br>AFESLALPLLAPRQQGSYQ<br>PQYPQQQRQLQPGGGPG\<br>LLGTDNGCGGEVHVDDPRI<br>QSYSQQRGSPNTNSQLHL(<br>PFDSNTNNHVVSGAAMAG(<br>KIANCTPQDLPPDLYDRLCN<br>WPAAAPPGCEPPNLGSVSE<br>SREVGSSPQLAAAAAAASR<br>LGRLLDLRSYVYEPPTAAAG<br>VSALRSDYRQDMQSVLEAF<br>RRDGATPLHLQRMMLATLP<br>STSCSCSGAAPATPYSSSA/<br>PHFAHPHLETEYAEHRRASI<br>CSSPVALYGASLALASLLLP | 391.3 |
| CGGAAACCCTATTATATGTTAGAGGAGATTGCACTCATGCAAGGCCGAGGTCCAGCGTTTACTTTGCTTATTGACTACTCGGTCTCGTAACTTCGCATATCCTTCGAACTGCTTTATCTTCTGTCCCAGGGTTCACGCGGCTCTGGCCCATT | >lcl\|ORF1<br>MADVFSTSF<br>SEGPCSSA(<br>GPLVCQVEG<br>RFQELTEFD(<br>WSDSSNDA<br>GGMTTWNV<br>GTDVGDALD<br>GLASGPVSG<br>LPTSGTWQ(<br>LQQQQQLQ<br>QQLLQQQQ<br>VLGQLNSNN<br>MNGIASGMS<br>QYQSEDLLV<br>CTHLVLDVQ<br>VSRSALLVQ<br>AAYIDGAGR<br>AAAADPKM/<br>QTGPLLGNV<br>RVLDKRSYT<br>FSEATLALV(<br>ADGLAIVHR(<br>LVPATWPVL/<br>LVDAFAPKPI<br>SNGDKTRGI<br>GAASAQRIF/ | Start 250 Stop 4152 | >Cre06.g300600.t1.1<br>MDVFNSGFQHTAGNLQQQ<br>LPGSGGREQRARLKGPLVC<br>NASSGRRQQRRMVLALPN<br>AGGGWPQQQRPLGPLPM(<br>AFESLALPLLAPRQQGSYQ<br>PQYPQQQRQLQPGGGPG\<br>LLGTDNGCGGEVHVDDPRI<br>QSYSQQRGSPNTNSQLHL(<br>PFDSNTNNHVVSGAAMAG(<br>KIANCTPQDLPPDLYDRLCN<br>WPAAAPPGCEPPNLGSVSE<br>SREVGSSPQLAAAAAAASR<br>LGRLLDLRSYVYEPPTAAAG<br>VSALRSDYRQDMQSVLEAF<br>RRDGATPLHLQRMMLATLP<br>STSCSCSGAAPATPYSSSA/<br>PHFAHPHLETEYAEHRRASI<br>CSSPVALYGASLALASLLLP | 392.5 |
| GTATGCACTACATGGCGACCTTTATCTTTGTTGAAGCCGGTTTGTCTACAGTCCAAGTATCTTTAGACTGAGTGAATTGCCGATCGTACTTTCAAGTGCATCCTGCATTTGTTTATCTTAAGTGCTTGTCCCAATGTCGATGTCACTAGATGAC<br>GATTGGCACAACCTTCAGGGCTGCCGGATGACAGAGAGGCTGATGCCTATACCGTTGCACTGCTGCAAGGTGGCGCCAAGAAGTTATCAATGTCAAATGCATATGGATGCATGCTTCCTTGGGCAGATGCAGTAGATGTTGCAGCCCTG<br>TGGTGCATGCACTGATGATGGCCAGGCTTTTGGCAGACAGCAGTGGAGCAGCCTGCAGGGGCCGCTTTTGGTGAGATGTGCTCATGTGTGTCAAGTGGGTAATGGGAGGCCCGATGTGGTGCTGGTGAACATGGGCATGGGCTATAC | >lcl\|ORF14<br>MSMSLDDS<br>DDSDDSNCI<br>LDSDVFSDA<br>ECHTDRCIP<br>ASRDSARCF<br>ETDAGISME<br>SVAAMEATF<br>CRAWRRYH<br>AMEGKTER\<br>TACAEAAVW<br>LLAKLAAASI<br>LAAVIRTCLS<br>VALKRAVAG<br>VDAATCGILF<br>LGIMCGAFE<br>MHLWRTFLI<br>GLFTRGVKF<br>FGLHRCTHL<br>CHENYLTAH<br>ALSASPVTK<br>TWLCNLSRL<br>LQDLSLRQN<br>GLSGCSGLE<br>LLTSLRVLD\<br>TGNMIVDVP<br>ILQQNNISLL<br>NNPVVATMA<br>ILDPFHRLE/<br>HWHQGLTG<br>FSIAAGCSIH<br>RMEHALPAT<br>AYTVALLQG<br>ARTSCTGPA<br>ARKLQWLS(<br>SAIQASWRG<br>LFPLRGEQ(<br>KEDDQFVDS<br>ADPVYSLHH<br>PSLQPRGMI<br>TSVAAVSPG<br>RLPENFLVS/<br>REERLQELV<br>LLAHNNQA/<br>YGNDAPVGI<br>PHRLPTL | Start 134 Stop 6757 | >Cre06.g258363.t1.2<br>MSALDDDPDFDVEAAFEAE<br>AAMQKREHHLEERGKQLTS<br>KGAAGLAAAATAAAAAGAA<br>AAARAAAEAAAAEERMRRN<br>RHMAAWRIGRAWRRYRRG<br>AKLTSAASGGSCMLFRTAL/<br>AAAPAFDAPGFTALVERGQL<br>SAFTALLRQSRALGMPLAM<br>PAAGRELADVVAALGPALGI<br>SSSSKSGSSGAGVGLQQR\<br>PRDYWGVPLLPSWRQAHH<br>ASAWRSFINPAPVAPAPALS<br>ASLHPALDRVSRLDLGLERL<br>ALRRLALDSNQLSGRLMGL<br>AARAAGLHGGVGVGVAGA(<br>SSSSGSAAPVLLPALQALNI<br>GNAEHAAAAAAYRDRLLTL\<br>SAKSGGAAATAAVAPLLHN/<br>ALAELHRQQLAVAAALAPS(<br>GGGRASLAAAVPAELLAAEI<br>LRSEQHQARCRDAATLIQA\<br>LMDSGELDDMMAFLQPLD<br>VQALAHPLGGSGSPLVPVP<br>NRRRTSGAGDYGAAGSEY(<br>QVVEQRPALRPQDVARLPQ<br>NRSNSHGSAASAPLALAAA<br>LPALANNWVTSPAQTPPGH | 169.5 |
| TTTAGACTGAGTGAATTGCCGATCGTACTTTCAAGTGCATCCTGCATTTGTTTATCTTAAGTGCTTGTCCCAATGTCGATGTCACTAGATGACTCCGATGCCGAAGCTGCCTTCGAGGAAGAGCTATCAAGGTTGAAGCTTCAGGCCTTGA(<br>TTCCTTGGGCAGATGCAGTAGATGTTGCAGCCCTGCGCTGCTTTCGACCCACGTTTCGATCCTTTCAGTACTTGGCCTTTGCTCGTACTTCGTGCACTGGTCCAGCATCCCTTGCCGTGAATACCCACTGTTTGCCAGAAGCTGCCTGC<br>GGTGCATGCACTGATGATGGCCAGGCTTTTGGCAGACAGCAGTGGAGCAGCCTGCAGGGGCCGCTTTTGGTGAGATGTGCTCATGTGTGTCAAGTGGGTAATGGGAGGCCCGATGTGGTGCTGGTGAACATGGGCATGGGCTATAC( | >lcl\|ORF1<br>MSMSLDDS<br>DDSDDSNCI<br>LDSDVFSDA<br>ECHTDRCIP<br>ASRDSARCF<br>ETDAGISME<br>SVAAMEATF<br>CRAWRRYH<br>AMEGKTER\<br>TACAEAAVW<br>LLAKLAAASI<br>LAAVIRTCLS<br>VALKRAVAG<br>VDAATCGILF<br>LGIMCGAFE<br>MHLWRTFLI<br>GLFTRGVKF<br>FGLHRCTHL<br>CHENYLTAH<br>ALSASPVTK<br>TWLCNLSRL<br>LQDLSLRQN<br>GLSGCSGLE<br>LLTSLRVLD\<br>YLDLGENMI<br>LTLLDLSFNI<br>RSGMDVFS\<br>PVWAISGAR<br>RGTNRQSEI<br>VKLGPMAA(<br>WQTTAPFET<br>AYGCMLPW<br>PEAACSEAF<br>PRVSLKNEF<br>HLAAWKAQS<br>AATIIQAHFR<br>FLSPLQEVF<br>LQALGRNEL<br>IPYPGTVQH<br>ISVMNSKRR<br>SSSFEIASGS<br>TVETYYRRL(<br>PRMQLNNP(<br>SQGCHEGFI | Start 73 Stop 6642 | >Cre06.g258363.t1.2<br>MSALDDDPDFDVEAAFEAE<br>AAMQKREHHLEERGKQLTS<br>KGAAGLAAAATAAAAAGAA<br>AAARAAAEAAAAEERMRRN<br>RHMAAWRIGRAWRRYRRG<br>AKLTSAASGGSCMLFRTAL/<br>AAAPAFDAPGFTALVERGQL<br>SAFTALLRQSRALGMPLAM<br>PAAGRELADVVAALGPALGI<br>SSSSKSGSSGAGVGLQQR\<br>PRDYWGVPLLPSWRQAHH<br>ASAWRSFINPAPVAPAPALS<br>ASLHPALDRVSRLDLGLERL<br>ALRRLALDSNQLSGRLMGL<br>AARAAGLHGGVGVGVAGA(<br>SSSSGSAAPVLLPALQALNI<br>GNAEHAAAAAAYRDRLLTL\<br>SAKSGGAAATAAVAPLLHN/<br>ALAELHRQQLAVAAALAPS(<br>GGGRASLAAAVPAELLAAEI<br>LRSEQHQARCRDAATLIQA\<br>LMDSGELDDMMAFLQPLD<br>VQALAHPLGGSGSPLVPVP<br>NRRRTSGAGDYGAAGSEY(<br>QVVEQRPALRPQDVARLPQ<br>NRSNSHGSAASAPLALAAA<br>LPALANNWVTSPAQTPPGH | 179.9 |
| CTTGAAGTTAATCACCTTTGTTGCTCCATTTTTTTACAGTTTCAAATAAAGTTTGTGTTTGTAGGACTCCAACCATGCCGCTGGGCGCTGCCCTTGAATCTCGCGAGGTTCTCAGCTCGTGCCTCAAGGAAGCTCTCGAAGATCCAGATC | >lcl\|ORF1<br>MPLGAALES<br>VDQLKDKLY<br>SVALRVNVQ<br>TPHTSEKKIL<br>ATAPKPVITK<br>SPNPAATVA:<br>KTTKPGVHL<br>FDASASTSL<br>AKSKETPPA<br>MELAAAREK<br>LHNKKMEEI<br>TRAEAPVFA:<br>VEEYRKTLEI<br>SRDDKDGD(<br>SALRQSAST<br>SVTELPEEEI | Start 76 Stop 2382 | >Cre17.g747847.t1.1<br>MACASPDSRAFLVGCLRSA<br>ANGPVSLRMLGTSSGRVSL<br>RLPTVAEEAVEQVGGVVVVS<br>AAHKLRTSMAESPLSPSAL/<br>SELRPKSCVLSREDLDLIEA<br>TAPASPPKPAPSPVTKGAAF<br>MPDFSAPFRPDPALAKPVT:<br>RSPNGAAASPNAANGGLR\ | 275.8 |
| TCCATGCATCTGGAGACCTTGCTGAGTAGTCCGCAAGGCCAGCGTGTTACGAAGCCACAGCCGTTTACTTTACGCTCGGAGATGCGAGTCAAGAGCACCGTGATGTCTACAGAAGAGATGGAGCTGGCGGCCGCGCGCGAGAAGGC | >lcl\|ORF2<br>MEEKRKKAI<br>PVFASEARIS<br>KTLEFKARP<br>DGDGIGKG\<br>SASTAVVGP<br>PEEEEACSP | Start 355 Stop 1146 | >Cre17.g747847.t1.1<br>MACASPDSRAFLVGCLRSA<br>ANGPVSLRMLGTSSGRVSL<br>RLPTVAEEAVEQVGGVVVVS<br>AAHKLRTSMAESPLSPSAL/<br>SELRPKSCVLSREDLDLIEA<br>TAPASPPKPAPSPVTKGAAF<br>MPDFSAPFRPDPALAKPVT:<br>RSPNGAAASPNAANGGLR\ | 94.7 |
| ATGGCAGGGCGAGGCCCTGCAGTGGGTGGGCCTTCGAAGGCACAACAGGATCAATGGAACGAAAAAGCAATGCAATTGATGAAGCGGCACGACCAAAATGGCGACGGCTTGCTCGATCCCCGTGAAATGGCGTCCGCAATCAACGA | >lcl\|ORF1<br>MAGRGPAV(<br>CGTHGFSAI<br>GGGGGGAG<br>TGAAAKMM<br>RTSGLSLRG<br>WAAATHVIS/<br>ASGGSGFAI<br>GSGLTRVSP<br>TARAAEEQV<br>RISGTEGQL<br>MVADSAMP\<br>WGEGTVCIS<br>LSEVKRVIMS<br>VVNRLVAMG<br>NMEKLVTQF<br>TAASAAATA/<br>FNLKSALGA<br>MLAADPPPA | Start Begin Stop 2652 | >Cre17.g718900.t1.2<br>MSGRGPGGPSKAQQDKW<br>DMYRRAMLMADTGDRGG(<br>DLGNIFSTASQRSSGLTLKQ<br>GGGGGGGGGGGGGGGG(<br>SGGLRSGSASGQKSLDLPS<br>SPGADAPRLVAEVRDNLDLI<br>NIEALDGQLRKAMRDGKVN<br>SEEDIRLQIKLLCEAMENVK<br>GMARLADVKDTENYLVDRG<br>ATAAQTASEARAAVERERAD<br>NEMTGKVIMKMARQITEMC<br>GARTSINGSGSSGGGLLQG | 441 |

| | | | | |
|---|---|---|---|---|
| CTTGAGTCCACATAAACTATGTCAAGGCCGTAAAAAGGCTATGTTTAAGATGCAGCAGCCACCTTTGTCTACGGGGGCTTTGTTTCGCGTTGCAACCACCCCATGCGGACCTGATGCCAAAGCTCCCATAAGTGGAGTGTTCCAAGTGC<br>GCCTCCCTGAAGCAGTGCTACAACAACTACTGCCGGCTGACGACCGGGGCCGCCGCCAAGATGATGATGGACGGCGACTTGAAGCTGTCCTCCGCGCAGTGGCACCAGCTGTGCGCGGATGTGGGGCTGGCGCAGCCAGTGGG | >lcl\|ORF5<br>MFKMQQPP<br>PNNKPRFRL<br>VAAKRIFIAL<br>TGQQQQHG<br>PPGSMYGG<br>GGGPLPQQ<br>QGAIARDEA<br>FSFDLLDLA<br>PTSHDYEIR<br>DIIGLVETCD<br>GDQIEQMV<br>RLRSWYDG<br>PDWITVSAV<br>WNCERCNY<br>RQLEAENT<br>TDFVKEGNL<br>GGYNPQGG<br>GGGPPRTP<br>SGSGSGAG<br>QQQNNTNN | Start 41 Stop 2947 | >Cre17.g718850.t1.1<br>MNPPLSTGALLHVAATSSG<br>AIVQITDFIINQVNDRKMFIA<br>VPPGAAGGMYGGPPPPM<br>AIARDDAPPRFIPISALNPYT<br>NHQYEIKLDRNSMVERVAE<br>QIEQMVRGGGRPVLAAKAL<br>VNVSAVLDMIKTDNQGGAS<br>VDRPQEFDRLAESIRFTPFF<br>GAGPSHYGGAAGGGGGY<br>PQQPGGGGYNQQGGGGY | 790 |
| TAAGAGTTTCGCTTACAGTATGTAAACCTCAGTCGCCGTGTGTCGAAATTCGCACGTACACTCGCATATATTAATACGTTACCCATGCACACTCCCTGACACAAGCACACACACACACACATGTACACGCGGTATCCTCCAGACAACAGCA | >lcl\|ORF1<br>MCSFDIINS<br>VEAACVIEL | Start 398 Stop 646 | >Cre17.g704850.t1.2<br>MADVEAKKEIIKNAIRGIPD<br>VPLRKPGKLPGDKISEEYK | 50.4 |
| GTACTTGTCCGTACTTTAACCATGGTCGAGTTGGCAGGGGTCCGGTAGGTTGGTTTACGTGACCGTGGTTACGGACAAGGTTCTGAGCTCAAGGTAGCGCTTATAACCTCAGTTCTGTCCATATCTAGGGCTGAGTCGTGCTACTTCAGA | >lcl\|ORF1<br>MAEDDAKK<br>RYKDKQIDV<br>KTEYSTDKI<br>EAACVIELP | Start 470 Stop 1015 | >Cre17.g704850.t1.2<br>MADVEAKKEIIKNAIRGIPD<br>VPLRKPGKLPGDKISEEYK | 280.8 |
| GTGTTTCCGAGCTACCTAGTGATCTTAAACAGTAGTTGGACCTCGGAGCTCAGCAGCCCTTGCATCGGTTGCGCTACTTTACCGTGTCGCCTAGCGCGCCACAAGCGCAAGCCATGGACCAGGAGGCCCAGGATGCCAAGCTGATTG | >lcl\|ORF10<br>MDQEAQDA<br>QLKLVDVLI<br>LNLIMRHCP<br>PLFPNPAEG<br>ALMESKPAS<br>KMVPLPAAA<br>DFGSTVRTG<br>WKELLIKGG | Start 114 Stop 1316 | >Cre16.g694950.t1.1<br>MADPINRRLKSLDAEVKDR<br>AHIDRPNEAILQLIVRRLSS<br>SGEASNSAATPNCAAGGP<br>KGFVHMDVKADNIFVDAAG<br>LEHLRQHSKVDEALLASLS | 255.4 |
| TCGGCTTTCAAGTTTTTATTCGGCAGTCATACCTGCATATCGTTTGCTTGGTAGCTGCTTAAAAAAAACAAAAAGAAAGAAGAAAGACAATTAAAAAAGATAAAGATAAACACCAATACAAAGGATAGCTCATGGTGACGTGGGGAGAAAGC | >lcl\|ORF5<br>MRHCPARM<br>NPAEGSASL<br>SKPASVPLV<br>LPAAAVLAG<br>TVRTGSPVT<br>LIKGGCSPP | Start 347 Stop 1237 | >Cre16.g694950.t1.1<br>MADPINRRLKSLDAEVKDR<br>AHIDRPNEAILQLIVRRLSS<br>SGEASNSAATPNCAAGGP<br>KGFVHMDVKADNIFVDAAG<br>LEHLRQHSKVDEALLASLS | 231.9 |
| GTTGCCGGAGTGTTGCTGTTCACCGCCGACCCCTTCAGCCCCTCGCTGCTCTTCATCCTGCCCAAGGACCCGATTCATGTGTTTTTCTGCCGTGTGCTGGGTGCAGCCACTCTTGCCAATGCGGTGTTCATGTTCACTGTGAAGCAG | >lcl\|ORF6<br>MVLSQHATI<br>TLLLLYGVE | Start 794 Stop 1042 | N/A | N/A |
| GAATCTCATATCTCTTTCGCCTTTTACTAAAGAGATCAGCGAGCACGGCTCTCTCAAGCGCATATCAACAATCTTTGAAAGGCTTTTGCCTTCACCCAAAACATTAATTGAGAAAGAATGTTATTTCTACGTTCGTGTGCATGAGACCGGTTG | >lcl\|ORF6<br>MVLSQHATI<br>TLLLLYGVE | Start 687 Stop 935 | N/A | N/A |
| GCTCAAGGCAGTGTGGCAAATGGTTGACCATTCTGTACTGTAACTAAAGGAAATGCATACCGAATTCTCTCTTCGAGGACAGATGTTAGGCCATAGAGCACCGTGAAAAATTAGCATGTAAACTACTGTTATCAGAGAATTATGGACGCATC<br>ACGAACTCGCGGCTTCCTGCGGTCCTTTTCATCATCAGCCAACCAGCCAGCCAGCCAACTGACGCTAACTGACCGATCAACCAACCAACCACCAAATCCTCGAAAGCCAGTCCACTTGGAATGCGGAGACCTCTATAAAACAGATTAG | >lcl\|ORF1<br>MDASTASLT<br>GFTIRDLPK<br>EFKMLNHIT<br>ELSQQQAE<br>ALVAPITGTL<br>ITWNVVENA<br>PGPGSNSL<br>VSRGNMGA<br>SYNGFGEP<br>EAAASADE<br>APDFRAAAA<br>R | Start 142 Stop 1797 | >Cre03.g168200.t1.2<br>MTQSMGGAAAGATVELEK<br>LVGTFSKPFIRRTATDEFKM<br>ALARSALADREPPTSPSAR<br>LPSLEALAAGAGLPVPAGG<br>SGLSGIAEGSARGDGASSD<br>YGSEPPPPPPDAPVVQALA<br>LRLLVWAALVQGCLRPEVL<br>RHLHRLFSMVPETRDALGS<br>LGALEKLRGLDPLLPAPYAV<br>WMEEDSLRGITGPDGMLA<br>RQGEVWRAIQAQFAESGVK<br>KSSMLKNSLQSFQFQHGE | 295 |
| CCTCGTCCACGTCGCCAATTCGCAAAAGGGAAAACCCTCGCTCATTTCCTTCGACTGTAAAAGGACTTTTTTACAGCCCAGCTTGCGCCTCCGCAACCTCTCTCCTTTGATACCGCTCTCCATGGTCTCAAGCTCCATGGGGATACGAA | >lcl\|ORF2<br>MACYPLQH<br>RRVPPVLLIT | Start 350 Stop 559 | N/A | N/A |
| GCCCGTCAATGCGAACATCCGCGATATCATGCAAGTGGATGATCAGCGTGGTCTTGCAAAGGCAATTTTTGAGATGTTTCGCTCCCGGCATGAGCACCTAACTACCTAACTTTCTCGGCGGTAACAGGCAATTGTGTCTCAGAGTCAAAT | >lcl\|ORF6<br>MECLQDRL<br>VIGNTKALW<br>ATHIRWGFN<br>AVVVVDAST<br>WADWADLR | Start 542 Stop 1282 | >Cre16.g654750.t1.2<br>MAATADLAQQFALAGLTIYQ<br>RKTISSIVDGVYLPPTDLEP<br>AYAEVVAAGGLQNWAASW | 164.1 |
| ATGTCGTCGACTGGTCCAGTTGTGTCTGTTACCGACGGCAAGTCCTCGCCAACAGCACAGATGCAGCAGGCAGCAATGCAGGGGAATACTGCCCTTTTATCTCAGCTGATATCACAGCATCCTAATCACGTTGACGGTGCCTCGGACT<br>GTGTGCGCCATCAACGCCACCATACAGCGCCTCGTGTCGTTATGGATCGGCGCGGCGTCGTGCGAGCCCGTCGCGCCGTACCGCGCTGTCGTACATCGCGTACCCATGCACGGCGATTCGGTGTTGCTTGCGGTGGGTCCAGCAC<br>GTGCAGCCACCGAGGAGGCGGAAGCCACAGCAGCGGCGGCGGTGGCGGCGGCGGCCGCGGCGCCGCCGCCGTTGCTGGTTCCTGCGCCGCCGACCAAGCCCCAGTCTTCGGACGGTCACGGGCCATCCCCACGGATCACC | >lcl\|ORF1<br>MSSTGPVV<br>GGQAPLLVA<br>LRVLLGRGA<br>ELRTGKGET<br>STDANGGG<br>TAAAVRRGG<br>SPQQSDIAG<br>QPDWSLHE<br>EADLPLLES<br>SELRELVLL<br>HHRQAHTA<br>GGVGVGGG<br>VAELAITAAP<br>SKARDRKET<br>RLKGPVRVA<br>HMLVGHIRR<br>GRGAAYDV<br>DGEEKAPKL<br>AHRGLLVAP<br>WLRDLHNP<br>QAMSGSGT<br>PHAGRQEA<br>LNPAQLHLQ<br>WYDEDVLL<br>DDGSTLKC<br>YVCEHPQA<br>GGAAMQVD<br>PGQPVTVFT<br>PIRLGAKGV<br>ESGDLSPEV<br>PPASPRDPA<br>TIQRLVSLWI<br>SSGGGGGG<br>APHLDLEAQ<br>NFQGRWILT<br>QLAAVGPC<br>SMPLSHLQL<br>GNAAPIGPL<br>EDGAAAVA<br>DDGTRRGQ<br>VAARFNDAL<br>GITFMGQDA<br>ETGEADGA<br>LGRKGLTAT<br>VVDLATRLA<br>LLQQRVDAA<br>LEIANECAS<br>AEVQVEAVR<br>AYESVAVQA<br>TRAAQSLAL<br>SKAKLSVLT<br>ADISRLLDE<br>EAATAALEK<br>SGNTSLGG<br>GRSLVGRPV<br>RSRAVQSP<br>VQEQRQAA<br>GGGGGGRS<br>EPGRRAAPA<br>VSEAELLAP<br>QAGRCSGV<br>NAELVRQTL<br>APPTKPQSS<br>AVAAPARER<br>TLSEELAEG<br>GGGGSIPG | Start Begin Stop 9888 | >Cre16.g654800.t1.1<br>MLAAQKGHVDVIRSLLFRG<br>VRVLLLSGARRQAVQDAAD<br>PPAAAGPRTPACNSRPAGG<br>SAPPPALSQPPVRRSPRAD<br>SFPALVGRLQGGVPRWLG<br>TQLRTALLAALKPQGGRAH<br>ELGSWESGRLTRLLRDLPA<br>VVVAEPPSSKAGLAGGAAA<br>PPVTPLSAAAGVLVTRGHA<br>GGGGGGGSADAEAGPRW<br>SSAGLEVTWMRDPHNPRT<br>SWQTGPEPGGPGTLLAPL<br>LKPAGAATAGGGGGGGGR<br>VRRCYDDGSSLKTSSKEQ<br>AVVGGSSLQVDESSKLRLL<br>SGSMSNGNGNGGSGSSTA<br>GGGGTEVDRGTVMWEVSW<br>AVAAGRLPGLWLAAAAADC<br>PHLDLEAQRAAERAAAAEA<br>GGSSASSGGGGGCSRHVA<br>APTADPQHQQHQQHQQAM<br>LVAVAGGVSVGDLEGLVEA<br>QQAAATAEGLEVAAANLAM<br>QDAAAAAAVMAAARWVAA<br>ALNARCGALQRQLAAAEAA<br>DNAKESAAAAGGGQRDLD<br>AARLLRSLATSHAALAPGG<br>VRDAQSDIAKALDDSDREM<br>SHNLAALGVMGLATTSMHS<br>AAAAALGAASSAVCGSLSS<br>LETAASDDEPHGGSGGGG<br>LEALIEAANLELAAAAVALR<br>QLSALLTPVPVPYHGRRAA<br>SLRRIDSSSAGPEPTLARLR<br>DGEIDFGLGD* | 358.2 |
| AAAGTCGTCATCTGATGCAATCGCTTAGGAGCCATACCTGTCGTCGGACAGCTTTGTCGAGGAGTCCAAAGCTTCAAGCGATGGGCTTATTCGGCTTCAATAACGTTTTTAGCTTTTTTGGTAAACCCGAAGGATGCAAGCTTGCTGCAA | >lcl\|ORF5<br>MSTSEAKPL<br>NEPIVVLFS<br>KALRQAIAE<br>NATQLMAD<br>EDMENPNE<br>DVDLQRTLL<br>TGISKTLPN<br>RCLTAKYCS<br>PSIKKR | Start 1391 Stop 2611 | >Cre08.g380350.t1.2<br>MAGGAKPFVWFGRACDSL<br>GTELSATETARLRSLRRAVE<br>QALRPLLTWMAEEGEVEAA<br>PGAAPTGGAHALRASLDM<br>QWVVVLVRRYSRLLAGLAA<br>GGGAAAAAGGQGPADGGT | 197.6 |
| CTTGACTTTTTAATGCATCGGCAGCCAATCCCTCACCTTATATATTCAACTTTACGTGTTCGAAGGGTCGGCATGACACAGGCCGTAACTGTGTAACCTTCATCCCCTAGTTCTTTTCAGATTAGTAAGAATATTTCTTTCTGTACATGTGTTAC | >lcl\|ORF4<br>MSTSEAKPL<br>NEPIVVLFS<br>KALRQAIAE<br>NATQLMAD<br>EDMENPNE<br>DVDLQRTLL<br>TGISKTLPN<br>RCLTAKYCS<br>PSIKKR | Start 344 Stop 1564 | >Cre08.g380350.t1.2<br>MAGGAKPFVWFGRACDSL<br>GTELSATETARLRSLRRAVE<br>QALRPLLTWMAEEGEVEAA<br>PGAAPTGGAHALRASLDM<br>QWVVVLVRRYSRLLAGLAA<br>GGGAAAAAGGQGPADGGT | 197.6 |
| AAACTTCAACGTGGGCATAGTAAGTTCTCTGCCTACCCCTCCGTTTCGTGGCAATCGCTATCTCTAAGTCAAGTATATTTTTAAATTGAATTTAGTAGCACAAAATACGCCCGCTGGGTGTAAATCCCCACGTGTCAAGCTTGGTTAGGCGAT | >lcl\|ORF9<br>MEAELQQL<br>IDDSIRQSA<br>TLSTPPRVS<br>LQIANNVFK<br>STAGGPAAN<br>HKMLTYEN<br>FAASVWQV<br>CESVIVPNL<br>TERFPAEVS<br>AAHGATSTT<br>LLPKEACLA<br>PVDLGPNLS<br>VCLQRLAA<br>ALFPAFELV<br>MFWERSGN<br>HEGFYIVNA<br>ALFVCKYGA<br>ASVKLITELP<br>QDFEEMQG | Start 201 Stop 3041 | >Cre16.g685901.t1.1<br>MDPELQQLATVFQATLSPD<br>QPLPDAEKAQIKQLLLSVAL<br>LRPLHETHTRLAGLFPAASA<br>LLEIDEDTFKPYLQRFAASV<br>SDTRRRAAADLVRSLTDRF<br>LLKADAVKFVTTFRSMLPKE<br>RVIVFVGPAIAPVSAVCLQR<br>LPQLYLAIFPPLLSPVFWER<br>SSKTPKFCRFFVVFLALFIC<br>TGGGSQGGSGQEDEEPDF | 1520.8 |
| CCTGGCGCTGTTCGTGTGCAAGTACGGTGCAGCGGTGGCCGCTGAGCAGCTCGACAAGGTGCAGCCCGGCATCATCTACATGCTGCTGCAGCAGGTGCGTGGTGTAAGCGCTGGCTGTGGCTGGTGCATATGATTAAGTAGTGCAA | >lcl\|ORF2<br>MWWWNGS | Start 587 Stop 682 | N/A | N/A |
| TTGAAAGGGCATAGGCTCCGTTTCTCGTTGGGATGCCACGATGATTACAAAGAGGGCCGAGCGACTTAAGCCAGCTAACAACCTACAAGATGTAACAATGATGACATAACAAACTAAACAACAACAACGACTTAAATAGTTATCCAGAGCC<br>TTTTTATTCTTGATCGATGTTCTAAATACATCACCGATGTGTATGTATATGCAGACTGGGCATTGAAATTTATGTCCCTTTTCCATGCTTGGACGACCTCGTTTCGTTGACGGGTATCCTGACAAAAGCGCTTTAGCGCTGCGTGCAATTCAGC | >lcl\|ORF6<br>MGGFGAPD<br>PPALPPTSG<br>DAKLEALVR<br>ASCVGVLAG<br>AERISEFRC<br>CDSTERGE<br>LPYVLEALA<br>AAAAAATTA<br>KMKSVGLD<br>QNKRRSCR<br>HGDNGGGD<br>DGRSAVMP<br>RVSYACRG<br>SLCNMHFL<br>GATGVKCA<br>AGDGSGDN<br>KQLVDSVM | Start 4924 Stop 7419 | >Cre10.g429601.t1.1<br>MGGFGAPDRGGSRESKNS<br>LQRAGNCGVGPILTYTTDD<br>AAIALPLVLYEVRASEALVER<br>PQNIDDVFMREGRNEPSNL<br>PAGAKTPAGKGGKAAAAAT<br>AKKGKGKAGAGKAANAFA<br>RPAPRNFVAEYEAAAAAKG<br>LGEGRLDCPTCAAKLGKW<br>ANFSSFRNKNFGFRLKDQK | 478.8 |

| | | | | |
|---|---|---|---|---|
| CGATGACGAAGATGACCAGCGGAGCGAAGACGAAAGCGATGACGACGAAGACGATGAAGACGAAGGCGGAGCGAAGCGTATAGGCCGGATCGTCTAGATCGATACCAACTGAAAGGGCATAGGCTCCGTTTCTCGTTGGGATGCCA<br>GAACTGGAGAGGATGAAAGGGGCCAAGGGGCGGCTGTAGAATGCGCGCGGATTGGGCGATTGGCCGTGACGGTCATTGTTGCACAAACATGTACGGCGTGAGAAATGTTTTTTATTCTTGATCGATGTTCTAAATACATCACCGATGTGT | >lcl\|ORF10<br>MRRHGLQT<br>VLVVAIVTLD<br>DRCVEGFDH<br>SFTEKDEMF<br>VVLISKGLTT<br>VRVGLNPCA<br>AATDLTTSPF<br>TGSVQELVA<br>GGPVFGGP<br>QQQGGPNL<br>VASAAPAVA<br>AAAEPSAAA<br>KSKSPRGTV | Start 2624 Stop 4492 | >Cre10.g429600.t1.2<br>MRRHGLQSPLDPHQLASW<br>CGLCQASVARSSKHCRACI<br>MAMLVAAVVMLGELFFFHV<br>AGAGGAAASAAGGGAAAG<br>GAGGLREPALLGAAAPSGT<br>QLTLGVSDTPPGSSGPTPAI<br>VPASPPPASAAQRSLMFPAI | 459.1 |
| ATGCGTTCATCGAGGGATACCTTCCCGCGTCGTGCGGCGACCGTTCGATCTTCTGCCCATAAATTGCAGCCGAACGCCATATCCTACTTATACGTAGTTGAACTGAAGGATCCTTAGTGCTGTGAAAGAATTAAGCCGCTTGGTCTTCAAG | >lcl\|ORF5<br>MSWDEYITS<br>KVMAAMAD<br>MGFYEDPP | Start 201 Stop 593 | >Cre10.g427250.t1.2<br>MAWEAYITSNLMCPVDSEG<br>RGRKDKSGFVARKGAQCIII | 154.5 |
| GCCATATCCTACTTATACGTAGTTGAACTGAAGGATCCTTAGTGCTGTGAAAGAATTAAGCCGCTTGGTCTTCAAGTTCAACTTCGATTGCAGATCATAGCCCTCCTACATTTTGTGTACGCAAGATGAGCTGGGACGAATACATCACAAGCA | >lcl\|ORF1<br>MALSGRNL<br>RVVEALGDY | Start 217 Stop 414 | >Cre10.g427250.t1.2<br>MAWEAYITSNLMCPVDSEG<br>RGRKDKSGFVARKGAQCIII | 82 |
| GCTTGTTCCATGATCACACACAATTGTCGATCTTATGGTCGTTGGCCGGGCCATGCAAGCAGTGTAGCCGGCAAGCCGAAGGTCGTTCGGCTCCTTGGCATTCGTGAAGCATCAAGTTCGTGGCGCGATGCCGGTCCGGCTGACCCA | >lcl\|ORF1<br>MITHNCRSY<br>RDIDSQPIN<br>AQTQEATRA<br>MGPCKVND<br>FAKDMGLPL<br>RHLQSACG | Start 10 Stop 798 | >Cre10.g426350.t1.1<br>MLAQGVTKEAGHGIRRAPH<br>ALGSDSKLSGYDAIIDLTRKI<br>LVERCRYLEQAGCASVCINS | 343.6 |
| GCTTGTTCCATGATCACACACAATTGTCGATCTTATGGTCGTTGGCCGGGCCATGCAAGCAGTGTAGCCGGCAAGCCGAAGGTCGTTCGGCTCCTTGGCATTCGTGAAGCATCAAGTTCGTGGCGCGATGCCGGTCCGGCTGACCCA | >lcl\|ORF1<br>MITHNCRSY<br>RDIDSQPIN<br>AQTQEATRA<br>MGPCKVND<br>FAKDMGLPL<br>RHLQSACG | Start 10 Stop 798 | >Cre10.g426350.t1.1<br>MLAQGVTKEAGHGIRRAPH<br>ALGSDSKLSGYDAIIDLTRKI<br>LVERCRYLEQAGCASVCINS | 343.6 |
| GGGCGGGCGGGGAGTTGCAGAACGTGGGAGATTGCTTCGCTTCAGGATGAGGGGGCGCTCTGGGAGGCAAGGCACTGTTGAGGCGGTTGAGAGGTAGTCTGTCGGTGGTGGCTGTTAGTTATTATTGCAGGTGGGGAATATGAATG<br>ACGGGGAACGATACATATACGTATCCCCCAAAGCCTTAGGCCACAGAACA | >lcl\|ORF10<br>MDALGPALT<br>PTAGGVIFRA<br>LAEHLADQA<br>GPRIDFFLDI<br>DSAAKKGGT | Start 3286 Stop 3951 | >Cre17.g696850.t1.2<br>MAEQVAELASGFLSFANQVA<br>KAIESKDTADDVQWLTYWTI<br>IEAAVALGLQAWKSLLAAVK | 236.5 |
| CGCCAACACCCTGCCTTCTAAGAGCCGCAGTCCACAATGCTTGCCATCCGTAGTTATTCCCTTACACTCCAAAGGCGGCATCAAAGGTATCCATCCAAGGAAAGAAAGTGACGCGAACGACTCTTATCGTAACCCAGGGATTGGCATTAT<br>GGGGGCAGACAGACAGACAGATAGACCTGCAGGTACACGCACGGGCCGGATGGCGGGCAGGCAGGCAGATAGGCAAGCAGGAAGCTACGTACCCGCCATAAGCGGGCTGACAGCTGCTTGGCGGAGGGGTTTCGGCACATGGCA | >lcl\|ORF11<br>MNSGSRCW<br>LAPACTSLE<br>LAALSRMPS<br>AAAVAAAAP<br>SVGAALAAV<br>NSFLASPMS<br>KTPPATATAF<br>SFTRAISRP<br>LHAGRTAPC<br>SGGLSKLQY<br>SAVTCLLLNI<br>RLRRLELAC<br>SVRSLVLSD<br>YLSIAHNRLS | Start 275 Stop 2317 | >Cre17.g697000.t1.1<br>MEEQTYSLDQFAPHQQPPA<br>WRHTSCCWLQSLSPACAS<br>HSDGVPAAGPAPGGEGQG<br>ANTHAAAATAAAAAGADAST<br>PAPATPGPATPQQVPAASPA<br>ASLREVAVRGRQRPASGLP<br>MTALEHLDLQRNKLAALPAI<br>AGALTHLDLAANNLVPADVL<br>VESE* | 449.1 |
| GGCCTGCGCAGTGATGAGTGGAAGGTATCAAGGGCAGCACGCGGGGCTCGTGAGGAAGGATTTGCATTTGCGGGAGAGGTAGTAAGGACGAAGAAACTTAGCGCTCGAAGGTTGTGCTAGAATGAAAAACGATGTCGAAGATTTGAG | >lcl\|ORF1<br>MMAVYVTCA | Start 297 Stop 446 | N/A | N/A |
| CTGCAACCAATTCGCAAGACACCCTAACCTCCTTTTATGAAGGCTCCCACTCCACAGGGCAAAGGTCGCCATCGCGTTCGCTGACAAGATTCCAAGTCTTGAGGCGAAGTTCGGCGAGCATCCGTTCACTATCGACCTGTCCCTGAAC | >lcl\|ORF2<br>MEKIKTVSD<br>ARSAGLAMI<br>TRDVQDFWI<br>FVGTVHVQF<br>SYLQAEEDE | Start 371 Stop 1003 | N/A | N/A |
| TTTCCTGAAAATGCGTTTGTCAAATGTTGAGGGTTGCACAGAAAGCCTTGTAACTAGCGGAGACTCTTCCAGTCGAACCCCCTGACGTCCTGCCGGCCTGCCTAACCCCGTTACTATAGATTGTTCATAAACGAATTGCCAAACTAAGGC | >lcl\|ORF1<br>MQPAVVDTL<br>RVPESTYLL<br>RDNRGYSVA<br>FHDTIRLLLV<br>SYQDVTGST<br>WLTATQLCP<br>VLLYKVTTSD<br>AVVPASSGA<br>SPALWAGN<br>GNRHLFLAF<br>IDVFVGLSVA<br>SNGCVENCI | Start 328 Stop 2127 | >Cre10.g446700.t1.2<br>MAQRDAIDTLSKACAYGDF<br>RGSLPVIETLLRHNADWEAF<br>KGNGEACTVLLQGGSACVL<br>ALRAWSWSVVISAGIALALL<br>NIKMLDSPALWAGNWQQL<br>MIVFEVLDVFVGLSVAALAIA<br>SHK* | 828.6 |
| GTAAAATATCGTATTTTCCTGAAAATGCGTTTGTCAAATGTTGAGGGTTGCACAGAAAGCCTTGTAACTAGCGGAGACTCTTCCAGTCGAACCCCCTGACGTCCTGCCGGCCTGCCTAACCCCGTTACTATAGATTGTTCATAAACGAATTG | >lcl\|ORF13<br>MQPAVVDTL<br>RVPESTYLL<br>RDNRGYSVA<br>FHDTIRLLLV<br>SYQDVTGST<br>WLTATQLCP<br>VLLYKVTTSD<br>AVVPASSGA<br>SPALWAGN<br>GNRHLFLAF<br>IDVFVGLSVA<br>SNGCVENCI | Start 342 Stop 2141 | >Cre10.g446700.t1.2<br>MAQRDAIDTLSKACAYGDF<br>RGSLPVIETLLRHNADWEAF<br>KGNGEACTVLLQGGSACVL<br>ALRAWSWSVVISAGIALALL<br>NIKMLDSPALWAGNWQQL<br>MIVFEVLDVFVGLSVAALAIA<br>SHK* | 828.6 |
| ATGTCGTCCGTCGACCGTGTAGCTTACATGGACAACATTAGAAGTAGTCAGATAGCTGAGCAGATCAATCGTGAAAAGGACATCCAGCGAAAATACTTCCAAGAGCAGGAAGCAGCAGGGCGCATAGTTCCAAAGCCGCCCTCTCCAG | >lcl\|ORF5<br>MAAKRKLAE<br>FDCGIHPAFI<br>FKGRIFMTH<br>DFYQTIEVS<br>PGADTPPVT<br>PIVALGRAQI<br>SLNEDIKKV<br>ASRDFFEAW<br>IAVHNISFSA<br>KAAGLAREV<br>RQAGGFGQ<br>EGVEGAGT<br>VLEWEGGR<br>AAAELQLIA<br>VGLRARIEKS | Start 1229 Stop 3415 | >Cre12.g498050.t1.2<br>MAATAKRKIDEGPADGEPDT<br>HFHLDHCAAVPYLLRKTRFI<br>GCRCLYTGDYSRLPDRHLF<br>QASSMMSKALGVYQTYVES<br>LGGPSSITTREGRRVPLRIAV<br>RLGEKPPREGAAVRGVLVR<br>LVTVRYVPADDASGVVSHV<br>VDGVEAAVDYRAGKVVCGE | 1139.8 |
| TTGTACTTACGGAACTTTTTCGGCATTGCTCCATGCCATAAATTGTGTGGATTCTGGACTTCTGGATGTGGGGACACGGACAGACACAAATGCTTTTCGCTTACATTTTTGGTGCAGTGCGGATACGACATGGCGTACAAGCAATCGGGCA | >lcl\|ORF1<br>MAAKRKLAE<br>FDCGIHPAFI<br>FKGRIFMTH<br>DFYQTIEVS<br>PGADTPPVT<br>PIVALGRAQI<br>SLNEDIKKV<br>ASRDFFEAW<br>IAVHNISFSA<br>KAAGLAREV<br>RQAGGFGQ<br>EGVEGAGT<br>VLEWEGGR<br>AAAELQLIA<br>VGLRARIEKS | Start 379 Stop 2565 | >Cre12.g498050.t1.2<br>MAATAKRKIDEGPADGEPDT<br>HFHLDHCAAVPYLLRKTRFI<br>GCRCLYTGDYSRLPDRHLF<br>QASSMMSKALGVYQTYVES<br>LGGPSSITTREGRRVPLRIAV<br>RLGEKPPREGAAVRGVLVR<br>LVTVRYVPADDASGVVSHV<br>VDGVEAAVDYRAGKVVCGE | 1139.8 |
| GTTGGGACTTTATAATCGTGTGTACACTCAAACAAGTTGTGTCGAATTCGGGTGTTTCACAGGGCCCCCTCAGACACAAATCGGCAATTCGCCTCTTACACGAGGAAGGAGAGACGAATATTATCACTTCCCATCTTTGAAAGTGGCGTAA | >lcl\|ORF3<br>MAPKKKAG<br>WGALRERVI<br>RNFTLSPMA<br>IHCPRLNKL<br>SKIPPLNLP<br>SIIPHVFRPF | Start 207 Stop 989 | >Cre12.g515200.t1.2<br>MAPKKKGGGKKKKKDDGA<br>KLSPPELTLIDLRGSSNLHN<br>TGCNNLSVVKLQCPNLLDS | 405.6 |
| CCACGGCTAAACAAGCTGAGCATAACGGATTGCGCTCAGCTTGAAACAATTATGCTGTGGACGGACGAGCTGACGGAGCTTGACCTGACCGGTGCGTGGAAGGCAGATGGGCAGGTGCTTGAATACATAATTGCTTGCCGAAAGCGC | >lcl\|ORF1<br>MLWTDELTE | Start 52 Stop 198 | N/A | N/A |
| ATGTTCGAGGCGCGGATTACCCAAGGAAGTGTGCTGAAGAAGCTCGTGGAGGCGCTAAAGGAGCTGGTGACCGAGGGCAACTTCGATGTTAGCAACACAGGGCTTTCCCTACAGGCGATGGATTCGTCTCATGTTTGTCTTGTATCGC | >lcl\|ORF1<br>MFEARITQG<br>TLREDGFDH<br>LMFEAPSQD<br>LTSIGDTVLI<br>EPVALTFALF<br>LAPKIDDEDI | Start Begin Stop 807 | >Cre12.g515850.t1.2<br>MFEARITQGAVLKKLVEALK<br>GNEDTITLKAEDNADQLTLM<br>RHNTTPEKPEEQTIIDLKEP | 485.7 |
| CGTCTCTTGTTACTTGTGAAACCGCGTCCTGGCACGCATGCCATGCTGACAGGATGAAAGATGAGCGTGAATGTAAGAAGTTTTAAGCGTCTGTTATGGAGGTGTCAATTGTGCCTCTTGCCCCTCCACGGACTTAGGCCACGGTTACTT | >lcl\|ORF6<br>MYLSSRRM<br>MVAVICSIDF<br>ILAASYSPKA<br>AYPTMQAIIK<br>VMFYVYGAL<br>SNRVFWALN<br>AMAAGTNVS<br>PIGHEAAGA<br>LFGTTNASA<br>LGTSEPQFD | Start 461 Stop 1840 | >Cre12.g515750.t1.1<br>MHSAGGKSRSLAVTDSRGA<br>FNLGHTITNFYGGYLASAYS<br>LWSPLIIASAGVDLMFYVYG<br>VRSSSFLSVLPWVAMACGT<br>ALSTVDTGGPSRLSDLASES<br>ASCAAFLTLGTSEPQFD* | 569.7 |
| CTTTTGTCGCTCATTTCCACTGATGTCTCCTGCTCAGCTGATGAAGAGTGCTTCTCGCGCGGCCGTGCGCCCGGCGCGTTCGGCCGTCCGTGCCTTGGCGCAGCAGCGCACGCAGGCCAAGTCTGCTGTGCGTAAGTCGCTCGCG | >lcl\|ORF2<br>MSPAQLMK<br>LQSQAARA<br>DVAAPIAKQ<br>TGVDLSAVT<br>GAVALYYLTF<br>ASGVPDVP<br>RRIGTSSKV<br>KLQIKPYTAT | Start 23 Stop 1183 | >Cre12.g497300.t2.1<br>MQLANAPRLAARPARSAVK<br>QSGVKILKDGYDVAAPYARI<br>SDPVTLGEFGLGAVALYYLS<br>AQTTALQIASLRRINSGTKVI | 472.2 |
| TTTTGTCGCTCATTTCCACTGATGTCTCCTGCTCAGCTGATGAAGAGTGCTTCTCGCGCGGCCGTGCGCCCGGCGCGTTCGGCCGTCCGTGCCTTGGCGCAGCAGCGCACGCAGGCCAAGTCTGCTGTGCGTAAGTCGCTCGCGA | >lcl\|ORF1<br>MSPAQLMK<br>LQSQAARA<br>DVAAPIAKQ<br>TGVDLSAVT<br>GAVALYYLTF<br>ASGVPDVP<br>RRIGTSSKV<br>KLQIKPYTAT | Start 22 Stop 1182 | >Cre12.g497300.t2.1<br>MQLANAPRLAARPARSAVK<br>QSGVKILKDGYDVAAPYARI<br>SDPVTLGEFGLGAVALYYLS<br>AQTTALQIASLRRINSGTKVI | 472.2 |
| CTAAGAAGTGCGCTAATGACGTCCTGATGTATATATAGCTCGCACGCTGTGCGGCAAACGATGCTTGGGGGTGAACTCATTACTCTCCTGATTGCCTAATTTGCGCAAATAGACTCTGAATATTAAATTGTTGCCCGTCATAATTGTGCTGTG | >lcl\|ORF1<br>MATAIARPAH<br>DDDGRGPTI<br>HHHHSFSGI<br>CVAAAPAPA<br>WIVAYMDIN<br>LTVCGQLGV | Start 304 Stop 1161 | N/A | N/A |
| ACCTGATGCTCTGGCCATGAGGGCACGCGCGGCACCCTTTTGAAACTTTGACACCTGCCCCTGAAGAAGGCGCAGAGAATTGTTCTTATCCTGCCGATGCCGAAAAGCTTTTCGCGGGACCTGTGATCTGATGTACTTGTACATGAAG | >lcl\|ORF4<br>MALKSPGT<br>LNFHETAWF<br>MAWCFVLSI<br>FICAGFYMF<br>PLATPPGAS | Start 845 Stop 1483 | >Cre13.g591851.t1.1<br>MAKASQAFKDALPSSLQKC<br>LMVLTFNIMSCVILIRKSVNF<br>GIMKQTGLYDPRQDEKRKM | 333.2 |
| GAGCATCAATATCGAGCCAAGTATATTACAGTTTAAAAGCCACTAGTTATTTGCGTCACCAAGGTGTTTGTCTGCTGCTAGAGTTCGCTAATTAGTTAAATGGCGTCTGAGCCCACACAAGAGGAGCTTGCGCTTGCACAGCCGGATCCAC<br>GTGGCTTTGGTGGCCGTGGGGGGGATGGTTCGGGGCTGGGTGGTGAAGGCGAGGGTGGCAACGGAGACGGAGGGTTAGGGCTTGGCGGCTGGATTTGCGGCTGAAGGGGGGAAGGACTCGGCGATAATGGATGGGGCGGGCTT | >lcl\|ORF28<br>MKQQAMEK<br>VRVALVVAVF<br>VALVAAVRVA<br>KVAAARVVK<br>VAVRVVRVA<br>GGMVRGW | Start 4035 Stop 4844 | N/A | N/A |
| CTGAGCATCAATATCGAGCCAAGTATATTACAGTTTAAAAGCCACTAGTTATTTGCGTCACCAAGGTGTTTGTCTGCTGCTAGAGTTCGCTAATTAGTTAAATGGCGTCTGAGCCCACACAAGAGGAGCTTGCGCTTGCACAGCCGGATCCA<br>GTGGCTTTGGTGGCGGCGGCGAGGGTGGCCTTGGTGGCGGCGGTGAGGGTGGTAAGGGTGACGGCGGTGAAGGTGGTGAGGGTGGTGGCGGTGAGGGCGGCTTTGGTGGCGGCGGTGATGGTGGCTTTGGTGGCGGCGGCGA | >lcl\|ORF20<br>MKQQAMEK<br>VRVALVVAVF<br>VALVAAVRVA<br>LVAAVKVALV<br>AAATVALVAA<br>ARVALVAAVF<br>VVKVAAVRVA<br>RVAAVRVAL<br>WVVKARVAT | Start 4037 Stop 5278 | N/A | N/A |

| | | | | |
|---|---|---|---|---|
| GTTAGGGAACTTCCACAGTGAACGTGACCGCTACTGTGTTCGTGCTGCTGCCATCACGCGCCAGGCAAGTATGTTGCTTGGCGCTGCTCCTTCCGCCCTTCAACAGTCGGCCCAGCCCTACAGCCAAAGAGCCCAGCGCCCACCCC | >lcl\|ORF1<br>MLLGAAPSA<br>LVVRRRRAP<br>QEAKEEEVE<br>AAYRSDALG<br>AVKVAVEFN | Start 71 Stop 814 | >Cre09.g408750.t1.2<br>MASTTSASGSGSGSEAGA<br>AKFPDVAWETTSMAVVAAS | 158.3 |
| CGGCAGTCTCCGCCTAAGACCAGCGTGCCACGACTTGTCGTACGCCGTCGCCGCGCCGTCGTCGGGGTTAGGAGGGTGGCCCCGGAGCTTTTTAATCGACGGTCGTTGCGGGGCCTTGTGGGCGATGGTGGGTCATTCAGGG | >lcl\|ORF1<br>MVGHSGDP<br>MRLSRASSL<br>VKWEVTSLG<br>PKSARIVFVH | Start 127 Stop 627 | >Cre09.g408750.t1.2<br>MASTTSASGSGSGSEAGA<br>AKFPDVAWETTSMAVVAAS | 159.8 |
| ATACAAATGTTGTTTCGTAATGCGATAATTGGAAGCTGGGACCTTCCCAAGATGCTGCCTCAGCTCGCACTGGGCTAGAAAATCATATCCTTTGACGTCACAGTAATATACATAAAGCCAGTATAGGACTTTTGATGACTCCTGGTTAATCTTA | >lcl\|ORF1<br>MSLYNGPPF<br>VYWYTRDKS<br>ELNELIKGR<br>GVGHTGPG<br>RRAEKRSRN<br>VERGNSLLW<br>YSPSPRTGF<br>GMQAKLYG<br>REVAGAVER<br>MERCRDRH | Start 166 Sop 1599 | >Cre18.g749347.t1.1<br>MSLYNGPPRAGARGGRDQ<br>ALGLKPKTVAKPAPQLDKH<br>GGAGVGGHAGGALPPPPP<br>SGSDSSGDEAAGRERGRE<br>DEGWRGPRGRSRSPRRH* | 189.5 |
| ATACAAATGTTGTTTCGTAATGCGATAATTGGAAGCTGGGACCTTCCCAAGATGCTGCCTCAGCTCGCACTGGGCTAGAAAATCATATCCTTTGACGTCACAGTAATATACATAAAGCCAGTATAGGACTTTTGATGACTCCTGGTTAATCTTA | >lcl\|ORF4<br>MMEVLGLKI<br>MKGLGYVLC<br>GLKSDANVA<br>LKKARREGE<br>RRQCSGPG<br>DRRHVRSRS<br>WDGHGRGN<br>VEGCSPISR | Start 404 Stop 1603 | >Cre18.g749347.t1.1<br>MSLYNGPPRAGARGGRDQ<br>ALGLKPKTVAKPAPQLDKH<br>GGAGVGGHAGGALPPPPP<br>SGSDSSGDEAAGRERGRE<br>DEGWRGPRGRSRSPRRH* | 73.6 |
| TTTCGTCCTTCAGTAGTTCAGTTACGGTACTAACGAACCCAGTTCAAGATAGTTTGGCGTCTGCTCCCTGTTTCTCAATGTCTAGATTTTGCTTAGCCATAGGTTAAAGTTAACGCATTACTTTATTTTGCTCACATTCTATTCGGGCTCATCGC | >lcl\|ORF9<br>MKVDFTLKF<br>AIARCAQCY<br>VLPEVRSEV<br>TAAGEEFLE<br>YVPLLEPQL<br>RALEENVVP<br>AAVPPAGLA<br>YLKVLQSPP<br>QNPSPVKLL<br>LLLFASCMY<br>KNPSDGGS<br>FQYTALVCE<br>AVAGTTTTA<br>VVTLLMHKV<br>DQVMEPCC<br>VPVDLHFSQ<br>LPPAELVRA<br>LIMVYYTSR<br>SELFDQLLL<br>HVSAESALQ | Start 360 Stop 3287 | >Cre09.g392504.t1.1<br>MGPPELGGCRGYTRCSST<br>EPDGHDGGGGGGGASGG | 112.8 |
| GTGGACTTCGAGTTCGCCACGCTCGGAGGATTCGCTGGCGCCGCCACCAGCCTGCCGCCCACTATCCAGGTGGCCTTCCAATACACCGCCCTGGTGTGTGAGAAAACACCACCACCACCACCACCACCACCACCACAGCAGCAGC | >lcl\|ORF1<br>MHKVLRVAE<br>LHFSQAPHL<br>ELVRAVYPVI<br>YYTSRCPPE<br>DQLLLDEPD<br>ESALQLQHE | Start 391 Stop 1206 | >Cre09.g392504.t1.1<br>MGPPELGGCRGYTRCSST<br>EPDGHDGGGGGGGASGG | 112.8 |
| CCCGGGGGCCCGGGGGCCCTGCGTAACTCTGCTGCAGAACTATCCAGTTGTTGCTATGTGTACGGGACCGCCCCTCGACCGTACGTACGTACGTACGAGCACTCCCGGGGCAGTAGATGGAGGTGGGGGAGGGAATGGGGGAGG | >lcl\|ORF2<br>MSAWTNSS<br>SAGVICNRT<br>PGRLPSF | Start 730 Stop 1053 | N/A | N/A |
| TCCTTCTTGCAGCTCTTGGAATTGCTTACCACCTTGTCTGCATTGATCTGTTTTGGGGAGGACATGTACGGCCTGTGACCTTTGTGTACGAATAACCCCCTTCAAATCCGCAGTTGCTGGATTGGCCAAAGCCTTTCGACGGCGGCGTTG | >lcl\|ORF1<br>MPLSTPVIST<br>RAWRTESG<br>LKCLSDQLC<br>FYIIFCY | Start 196 Stop 669 | N/A | N/A |
