## Supplemental Table 3 for "Identification of Cell-Type Specific Alternative Transcripts in the Multicellular *Alga Volvox carteri*"

| sequence | ProteinSeq | StartStop | Chlamy Protein Homolog | Score |
|---|---|---|---|---|
| TCAAAAATTTGGGAGATCGTCTATTGCCAAGTTGAATAAGTTGTCAATTTCGCACGATCCTCTTACTTTTTAAGCGCGTCATTTGCGCAATCAGTACTTCTTCAAGATGGCTCCTCTCGCCCTGAACCGTGCTGCTTCTCGCGTCAGCGCT | >lcl\|ORF3<br>MAPLALNR<br>FVEKDTAG<br>GLGLTALIN<br>AEVVAPPV | Start 107 Stop 649 | >Cre13.g570851.t1.1<br>MYPTVMKPFEAGSASDTV<br>VAAPVAEPVAVEAPAVASE* | 62.4 |
| TCAAAAATTTGGGAGATCGTCTATTGCCAAGTTGAATAAGTTGTCAATTTCGCACGATCCTCTTACTTTTTAAGCGCGTCATTTGCGCAATCAGTACTTCTTCAAGATGGCTCCTCTCGCCCTGAACCGTGCTGCTTCTCGCGTCAGCGCT | >lcl\|ORF3<br>MAPLALNR<br>FVEKDTAG<br>GLGLTALIN<br>AEVVAPPV | Start 107 Stop 604 | >Cre13.g570851.t1.1<br>MYPTVMKPFEAGSASDTV<br>VAAPVAEPVAVEAPAVASE* | 62.4 |
| GTGCAATCTGCAAAATCTTGCTCCTTTCATTTAGTGACAGCTGCCCAACACAACAGCGAAGTCTCCGGGCAACAGAACTCCGATCAGTTCGTTGTAAACTCACTTCCAGTACTAACGCTGGCCCTGTAGATAAACCTGTCCACGTAAGA | >lcl\|ORF1<br>MANHFEEF<br>LPSLLDLPS<br>TFLDLIAEQ<br>FIELMQNM<br>LIDKGIKYL<br>GHLCVRRT<br>ATLERSGG<br>IVVPRSRFA<br>LDDAYYKL<br>EGGSPSPA | Start 172 Stop 1611 | >Cre04.g229700.t1.2<br>MAGDFAAFEAKMKAANLS<br>TSMGLEKAKSLLVVKDGKT<br>PGHGDIYPSLLGSGMLDAL<br>KHKYFNTNNLWVSLEALAA<br>VAVAPALKGAIPLVKLDDGH | 720.3 |
| GTGCAATCTGCAAAATCTTGCTCCTTTCATTTAGTGACAGCTGCCCAACACAACAGCGAAGTCTCCGGGCAACAGAACTCCGATCAGTTCGTTGTAAACTCACTTCCAGTACTAACGCTGGCCCTGTAGATAAACCTGTCCACGTAAGA | >lcl\|ORF1<br>MANHFEEF<br>LPSLLDLPS<br>TFLDLIAEQ<br>FIELMQNM<br>LIDKGIKYL<br>GHLCVRRT<br>ATLERSGG<br>IVVPRSRFA<br>LDDAYYKL<br>EGGSPSPA<br>VVDNQGCW | Start 172 Stop 1716 | >Cre04.g229700.t1.2<br>MAGDFAAFEAKMKAANLS<br>TSMGLEKAKSLLVVKDGKT<br>PGHGDIYPSLLGSGMLDAL<br>KHKYFNTNNLWVSLEALAA<br>VAVAPALKGAIPLVKLDDGH | 717.6 |
| TATAAACGTTATTGCTTAGTATTTCTTTTCACCGGTGGCTTGAATTACCAATTCCTGTTTGCGTCCTTGCGCTCGTTCTATAGTGCTAGGCTTGAGTGGTATCACGCTTGTCATTTTGAAAGAGGCTTGATGCATTAATTATATCGACGATATGT | >lcl\|ORF9<br>MDAQTGN<br>MAVGLSLF<br>VTAEWLCR<br>SDFSLRQS<br>TDGSHAAH<br>KHKINGAG<br>VSTDISNRA<br>I | Start 188 Stop 1243 | >Cre03.g211521.t1.1<br>METATAAPPRLATAVSFTKR<br>MSKNYLSKHLTPSHLKNFY<br>GGITGGMGPAGGPPGTVA<br>PEGIKLGSVSADISTRASCT | 378.6 |
| CGACACATGCCGCTCTTGTGCATGTGCATGTGCATTTTGCCGTCCATCTTGTGAGAACGGATAGGAAGGAAAACAGTTTAAAAGGGGAATGAAGGCGAACAGGTTTCTCAGGATCGCCACACTTTAATAGGAGCTATTGCCGCATTGCG | >lcl\|ORF3<br>MPACTVLG<br>MRLPPSGS<br>AHSPLPRR<br>TPEVAQRC | Start 2227 Stop 2751 | N/A | N/A |
| GTAAAGTTTGCCTAACTCCGACCAGGCTGCGCCGCTAGGCGATGCTTCACCGTTAGACCATCGTTTTCCTCAACGAGCCCTGATTGCGGTGACTTGTTTCTATTACAAGACAGCTATACAGGTGCAACGGCGGACATGGGCAGCGGAC<br>ATAATTGTAGCTAGGGCTGATTTAACAGCGTGAGCGGACATTGTATGGTTGAATAGTATAGGATTGTGTTTGGTAGGACGGATTAGATTTAAGAGTGGGAAGGGGTGTGAGTGGGAGCCTGTTGCCATTAGCATTGGTGCCCCGCTGGAGTG | >lcl\|ORF1<br>MGSGRGLR<br>IKLRGQSVD<br>VAASTEAVP<br>APLVLLDPA<br>PVSAPAPPT<br>AAVALVPTP<br>ILFGSQTGT<br>VISSSTGDG<br>SNYTRFMY<br>ALKAAVSHA<br>AAAGVAAK<br>TAKAAAEAE<br>EVPPPEALP<br>PSGAEDCTS<br>PTAATVGLP<br>PDSSMSYA<br>NFPKREETV<br>GRARGVRE<br>RKVIHLELS<br>LQPAAGAA<br>LLAEHATDA<br>HAPFEALLD<br>RLGVATTWL<br>VMVGPGTG<br>YKEELEAFK<br>HVYVCGDG<br>VWS | Start 136 Stop 4047 | >Cre16.g683550.t1.2<br>MKIGVAVAGGFGALLLLTPL<br>GNAILARLKQQRAAPPPPA<br>VALRKKQEGKPLVGVKFTLL<br>PVAAVSVAAPSASVPAPSTG<br>AAAATGGNAILARLKQQRA<br>NSGAFFVALRKKQEGKPLV<br>VTAGPSGPSRRSVEMMTG<br>PNADSAESLTVAAAAAAPEA<br>GDGDPPENATNFFVALRKK<br>MAAVSIDAPSATAAVKPAAV<br>AAKDAAEKEAAEKETAEKA<br>LAEAEASAAALAAAAQHPE<br>LLSPRETEDGGDDMLLTPR<br>GLLLGLAPVGVDTKGAPAL<br>PGDAIGVLPANHPDLVANL<br>SEKRTLLFLSAKGGKDAYA<br>LASPWTTEGISNPANPVWV<br>LSALHVAFSRAQETKVYVQ | 766.5 |
| CGTCTATACATCCAGGCATTTACCTGACGGAGAAGCGTATCCGAATGGCAATAAACGAGTGTGCAGACTTCCTGCACAAGAAATGCAAACACATACCATGGTGCTAGAATTGTTTACTGTGTTAGCCCATATATACTCATCGCTTCATCGTTG<br>GCCTTGCCTTGCCATTGTGCGCTTTGCCATTGTGCTTTGTTATGGCCTGCGCCCAGGAAGCTGACTAATACATTCTTGGCCTCAGTATGAAAGCTGGGATGTCAGCCTGAGCCCTTGGCGTGGGACTAGACTTCGCTGGCGCAGAGTAT | >lcl\|ORF1<br>MATQNGPF<br>AKASPTTSS<br>PAPEPLKEA<br>LARFRKQRT<br>SEPAAKPPP<br>TCASLNELG<br>GAPPPPPA<br>VEADEVEGV<br>DVAEAAAAK<br>AKAAAEKAA<br>DAARRKDE<br>PPPVEAPPS<br>AAGEPASLL<br>DGETALGD<br>TSSRRRTG<br>VDTKGAPPL<br>SAEKPFWS<br>NHPDLITNL<br>SVGYALSH<br>YAHEISEHQ<br>ARGATHLSV<br>YVPIFLRRA<br>APELLGEAV<br>VYVQDLIKQ<br>LSEQDATAF | Start 172 Stop 3849 | >Cre16.g683550.t1.2<br>MKIGVAVAGGFGALLLLTPL<br>GNAILARLKQQRAAPPPPA<br>VALRKKQEGKPLVGVKFTLL<br>PVAAVSVAAPSASVPAPSTG<br>AAAATGGNAILARLKQQRA<br>NSGAFFVALRKKQEGKPLV<br>VTAGPSGPSRRSVEMMTG<br>PNADSAESLTVAAAAAAPEA<br>GDGDPPENATNFFVALRKK<br>MAAVSIDAPSATAAVKPAAV<br>AAKDAAEKEAAEKETAEKA<br>LAEAEASAAALAAAAQHPE<br>LLSPRETEDGGDDMLLTPR<br>GLLLGLAPVGVDTKGAPAL<br>PGDAIGVLPANHPDLVANL<br>SEKRTLLFLSAKGGKDAYA<br>LASPWTTEGISNPANPVWV<br>LSALHVAFSRAQETKVYVQ | 765.8 |
| TTCACCCGTTCCTAGCGGCAGTCAGTGCGATGGAGACTAGGCGTTGGTTCGCGCAAGCGACAGCACATCGGCAGGAAATCACGCGTCAGGCCCAAGCTATGGACCGGATCATCCGTGCATTTATGTTCTTGGAACACATATGAACCTT | >lcl\|ORF3<br>MQYPVVQS | Start 30 Stop 143 | N/A | N/A |
| CTCTAAAACGTGATTAGGAAAGCGGAACACCTAAAACTTTAACCAGTGCAAGACAAAAGGATGTAAATTAGCGTGCTTTAGAGGCAAAGGTTTGGTCCCCTGCATGGGGTATGACCTGCTGCGCCTTATTTGTTGGCTTGCGCAAAGTTC | >lcl\|ORF1<br>MIPKAATQA<br>ESEEQKLLR<br>GSLASQFSA<br>CVIPVIQVAL<br>MQRMHATE<br>NSVAVETVD<br>IPAKQRIFPA | Start 196 Stop End | >Cre04.g215550.t1.2<br>MTRLHAVPGTNSRVAVDNV<br>GTQAADFAVARGVDLSRLA<br>ERMHATEGGYNLTRLGGLH<br>QDGGAECSPVAIAEIPAVGR<br>VCERLAGHARPFTHSQSEL | 307 |
| CTTGAAGTTAATCACCTTTGTTTGCTCCATTTTTTTACAGTTTCAAATAAAGTTTGTGTTTGTAGGACTCCAACCATGCCGCTGGGCGCTGCCCTTGAATCTCGCGAGGTTCTCAGCTCGTGCCTCAAGGAAGCTCTCGAAGATCCAGATC | >lcl\|ORF1<br>MPLGAALE<br>VDQLKDKLY<br>SVALRVNVQ<br>TPHTSEKKI<br>ATAPKPVITK<br>SPNPAATVA<br>KTTKPGVHL<br>FDASASTSL<br>AKSKETPPA<br>MELAAARE<br>LHNKKMEE<br>TRAEAPVFA<br>VEEYRKTLE<br>SRDDKDGD<br>SALRQSAST<br>SVTELPEEE | Start 76 Stop 2382 | >Cre17.g747847.t1.1<br>MACASPDSRAFLVGCLRSA<br>ANGPVSLRMLGTSSGRVSL<br>RLPTVAEEAVEQVGGVVVV<br>AAHKLRTSMAESPLSPSAL<br>SELRPKSCVLSREDLDLIEA<br>TAPASPPKPAPSPVTKGAA<br>MPDFSAPFRPDPALAKPVT<br>RSPNGAAASPNAANGGLR | 275.8 |
| TCCATGCATCTGGAGACCTTGCTGAGTAGTCCGCAAGGCCAGCGTGTTACGAAGCCACAGCCGTTTACTTTACGCTCGGAGATGCGAGTCAAGAGCACCGTGATGTCTACAGAAGAGATGGAGCTGGCGGCCGCGCGCGAGAAGGC | >lcl\|ORF2<br>MEEKRKKA<br>PVFASEARI<br>KTLEFKARP<br>DGDGIGKG<br>SASTAVVGP<br>PEEEEACSP | Start 355 Stop 1146 | >Cre17.g747847.t1.1<br>MACASPDSRAFLVGCLRSA<br>ANGPVSLRMLGTSSGRVSL<br>RLPTVAEEAVEQVGGVVVV<br>AAHKLRTSMAESPLSPSAL<br>SELRPKSCVLSREDLDLIEA<br>TAPASPPKPAPSPVTKGAA<br>MPDFSAPFRPDPALAKPVT<br>RSPNGAAASPNAANGGLR | 94.7 |
| GTGTTTCCGAGCTACCTAGTGATCTTAAACAGTAGTTGGACCTCGGAGCTCAGCAGCCCTTGCATCGGTTGCGCTACTTTACCGTGTCGCCTAGCGCGCCACAAGCGCAAGCCATGGACCAGGAGGCCCAGGATGCCAAGCTGATTG | >lcl\|ORF10<br>MDQEAQDA<br>QLKLVDVLI<br>LNLIMRHCP<br>PLFPNPAEG<br>ALMESKPAS<br>KMVPLPAAA<br>DFGSTVRTG<br>WKELLIKGG | Start 114 Stop 1316 | >Cre16.g694950.t1.1<br>MADPINRRLKSLDAEVKDR<br>AHIDRPNEAILQLIVRRLSSF<br>SGEASNSAATPNCAAGGP<br>KGFVHMDVKADNIFVDAAG<br>LEHLRQHSKVDEALLASLS | 255.4 |
| TCGGCTTTCAAGTTTTTATTCGGCAGTCATACCTGCATATCGTTTGCTTGGTAGCTGCTTAAAAAAAACAAAAAGAAAGAAGAAAGACAATTAAAAAAGATAAAGATAAACACCAATACAAAGGATAGCTCATGGTGACGTGGGGAGAAAGC | >lcl\|ORF5<br>MRHCPARM<br>NPAEGSASL<br>SKPASVPLV<br>LPAAAVLAG<br>TVRTGSPVT<br>LIKGGCSPP | Start 347 Stop 1237 | >Cre16.g694950.t1.1<br>MADPINRRLKSLDAEVKDR<br>AHIDRPNEAILQLIVRRLSSF<br>SGEASNSAATPNCAAGGP<br>KGFVHMDVKADNIFVDAAG<br>LEHLRQHSKVDEALLASLS | 231.9 |
| TTGAAAGGGCATAGGCTCCGTTTCTCGTTGGGATGCCACGATGATTACAAAGAGGGCCGAGCGACTTAAGCCAGCTAACAACCTACAAGATGTAACAATGATGACATAACAAACTAAACAACAACAACGACTTAAATAGTTATCCAGAGCC<br>TTTTTATTCTTGATCGATGTTCTAAATACATCACCGATGTGTATGTATATGCAGACTGGGCATTGAAATTTATGTCCCTTTTCCATGCTTGGACGACCTCGTTTCGTTGACGGGTATCCTGACAAAAGCGCTTTAGCGCTGCGTGCAATTCAGC | >lcl\|ORF6<br>MGGFGAPD<br>PPALPPTSG<br>DAKLEALVR<br>ASCVGVLAG<br>AERISEFRC<br>CDSTERGE<br>LPYVLEALA<br>AAAAAATTA<br>KMKSVGLD<br>QNKRRSCR<br>HGDNGGGD<br>DGRSAVMP<br>RVSYACRG<br>SLCNMHFL<br>GATGVKCA<br>AGDGSGDN<br>KQLVDSVM | Start 4924 Stop 7419 | >Cre10.g429601.t1.1<br>MGGFGAPDRGGSRESKNS<br>LQRAGNCGVGPILTYTTDD<br>AAIALPLVLYEVRASEALVER<br>PQNIDDVFMREGRNEPSNL<br>PAGAKTPAGKGGKAAAAAT<br>AKKGKGKAGAGKAANAFA<br>RPAPRNFVAEYEAAAAAKG<br>LGEGRLDCPTCAAKLGKW<br>ANFSSFRNKNFGFRLKDQK | 478.8 |
| CGATGACGAAGATGACCAGCGGAGCGAAGACGAAAGCGATGACGACGAAGACGATGAAGACGAAGGCGGAGCGAAGCGTATAGGCCGGATCGTCTAGATCGATACCAACTGAAAGGGCATAGGCTCCGTTTCTCGTTGGGATGCCA<br>GAACTGGAGAGGATGAAAGGGGCCAAGGGGCGGCTGTAGAATGCGCGCGGATTGGGCGATTGGCCGTGACGGTCATTGTTGCACAAACATGTACGGCGTGAGAAATGTTTTTTATTCTTGATCGATGTTCTAAATACATCACCGATGTG | >lcl\|ORF10<br>MRRHGLQT<br>VLVVAIVTLD<br>DRCVEGFD<br>SFTEKDEM<br>VVLISKGLTT<br>VRVGLNPC<br>AATDLTTSP<br>TGSVQELVA<br>GGPVFGGP<br>QQQGGPNL<br>VASAAPAVA<br>AAAEPSAAA<br>KSKSPRGT | Start 2624 Stop 4492 | >Cre10.g429600.t1.2<br>MRRHGLQSPLDPHQLASW<br>CGLCQASVARSSKHCRAC<br>MAMLVAAVVMLGELFFFHV<br>AGAGGAAASAAGGGAAAG<br>GAGGLREPALLGAAAPSGT<br>QLTLGVSDTPPGSSGPTPA<br>VPASPPPASAAQRSLMFPA | 459.1 |
| GGGCGGGCGGGGAGTTGCAGAACGTGGGAGATTGCTTCGCTTCAGGATGAGGGGGCGCTCTGGGAGGCAAGGCACTGTTGAGGCGGTTGAGAGGTAGTCTGTCGGTGGTGGCTGTTAGTTATTATTGCAGGTGGGGAATATGAATG<br>ACGGGGAACGATACATATACGTATCCCCCAAAGCCTTAGGCCACAGAACA | >lcl\|ORF10<br>MDALGPALT<br>PTAGGVIFR<br>LAEHLADQA<br>GPRIDFFLD<br>DSAAKKGG | Start 3286 Stop 3951 | >Cre17.g696850.t1.2<br>MAEQVAELASGFLSFANQV<br>KAIESKDTADDVQWLTYWT<br>IEAAVALGLQAWKSLLAAVK | 236.5 |
