## Supplemental Table 4 for "Identification of Cell-Type Specific Alternative Transcripts in the Multicellular *Alga Volvox carteri*"

| PACid | species | locusName | transcriptName | proteinName |
|---|---|---|---|---|
| PAC:32894653 | V. carteri | Vocar.0054s0007 | Vocar.0054s0007.1 | Vocar.0054s0007.1.p |
| PAC:32890341 | V. carteri | Vocar.0024s0049 | Vocar.0024s0049.1 | Vocar.0024s0049.1.p |
| PAC:32884784 | V. carteri | Vocar.0001s1718 | Vocar.0001s1718.1 | Vocar.0001s1718.1.p |
| PAC:32888047 | V. carteri | Vocar.0007s0295 | Vocar.0007s0295.1 | Vocar.0007s0295.1.p |
| PAC:32894976 | V. carteri | Vocar.0012s0183 | Vocar.0012s0183.1 | Vocar.0012s0183.1.p |
| PAC:32884296 | V. carteri | Vocar.0001s0791 | Vocar.0001s0791.1 | Vocar.0001s0791.1.p |
| PAC:32893870 | V. carteri | Vocar.0006s0453 | Vocar.0006s0453.1 | Vocar.0006s0453.1.p |
| PAC:32888600 | V. carteri | Vocar.0014s0188 | Vocar.0014s0188.1 | Vocar.0014s0188.1.p |
| PAC:32896031 | V. carteri | Vocar.0021s0076 | Vocar.0021s0076.1 | Vocar.0021s0076.1.p |
| PAC:32890182 | V. carteri | Vocar.0031s0077 | Vocar.0031s0077.1 | Vocar.0031s0077.1.p |
| PAC:32894234 | V. carteri | Vocar.0006s0411 | Vocar.0006s0411.1 | Vocar.0006s0411.1.p |
| PAC:32894919 | V. carteri | Vocar.0012s0137 | Vocar.0012s0137.1 | Vocar.0012s0137.1.p |
| PAC:32886039 | V. carteri | Vocar.0001s0835 | Vocar.0001s0835.1 | Vocar.0001s0835.1.p |
| PAC:32884071 | V. carteri | Vocar.0005s0221 | Vocar.0005s0221.1 | Vocar.0005s0221.1.p |
| PAC:32895487 | V. carteri | Vocar.0011s0348 | Vocar.0011s0348.1 | Vocar.0011s0348.1.p |
| PAC:32887568 | V. carteri | Vocar.0029s0046 | Vocar.0029s0046.1 | Vocar.0029s0046.1.p |
| PAC:32889625 | V. carteri | Vocar.0004s0355 | Vocar.0004s0355.1 | Vocar.0004s0355.1.p |
| PAC:32889510 | V. carteri | Vocar.0004s0080 | Vocar.0004s0080.1 | Vocar.0004s0080.1.p |
| PAC:32885242 | V. carteri | Vocar.0001s1052 | Vocar.0001s1052.1 | Vocar.0001s1052.1.p |
| PAC:32891697 | V. carteri | Vocar.0086s0006 | Vocar.0086s0006.1 | Vocar.0086s0006.1.p |
| PAC:32883444 | V. carteri | Vocar.0023s0049 | Vocar.0023s0049.1 | Vocar.0023s0049.1.p |
| PAC:32892054 | V. carteri | Vocar.0042s0009 | Vocar.0042s0009.1 | Vocar.0042s0009.1.p |
| PAC:32896020 | V. carteri | Vocar.0021s0064 | Vocar.0021s0064.1 | Vocar.0021s0064.1.p |
| PAC:32898511 | V. carteri | Vocar.0018s0017 | Vocar.0018s0017.1 | Vocar.0018s0017.1.p |
| PAC:32892754 | V. carteri | Vocar.0019s0108 | Vocar.0019s0108.1 | Vocar.0019s0108.1.p |
| PAC:32898271 | V. carteri | Vocar.0025s0053 | Vocar.0025s0053.1 | Vocar.0025s0053.1.p |
| PAC:32896053 | V. carteri | Vocar.0021s0181 | Vocar.0021s0181.1 | Vocar.0021s0181.1.p |
| PAC:32894168 | V. carteri | Vocar.0006s0010 | Vocar.0006s0010.1 | Vocar.0006s0010.1.p |
| PAC:32895694 | V. carteri | Vocar.0028s0177 | Vocar.0028s0177.1 | Vocar.0028s0177.1.p |
| PAC:32895341 | V. carteri | Vocar.0011s0137 | Vocar.0011s0137.1 | Vocar.0011s0137.1.p |
| PAC:32894017 | V. carteri | Vocar.0006s0283 | Vocar.0006s0283.1 | Vocar.0006s0283.1.p |
| PAC:32887177 | V. carteri | Vocar.0027s0172 | Vocar.0027s0172.1 | Vocar.0027s0172.1.p |
| PAC:32895681 | V. carteri | Vocar.0028s0190 | Vocar.0028s0190.1 | Vocar.0028s0190.1.p |
| PAC:32898512 | V. carteri | Vocar.0018s0208 | Vocar.0018s0208.1 | Vocar.0018s0208.1.p |
| PAC:32888304 | V. carteri | Vocar.0007s0221 | Vocar.0007s0221.1 | Vocar.0007s0221.1.p |
| PAC:32888729 | V. carteri | Vocar.0041s0033 | Vocar.0041s0033.1 | Vocar.0041s0033.1.p |
| PAC:32890335 | V. carteri | Vocar.0024s0032 | Vocar.0024s0032.1 | Vocar.0024s0032.1.p |
| PAC:32895249 | V. carteri | Vocar.0011s0049 | Vocar.0011s0049.1 | Vocar.0011s0049.1.p |
| PAC:32896094 | V. carteri | Vocar.0079s0020 | Vocar.0079s0020.1 | Vocar.0079s0020.1.p |
| PAC:32888058 | V. carteri | Vocar.0007s0126 | Vocar.0007s0126.1 | Vocar.0007s0126.1.p |
| PAC:32889311 | V. carteri | Vocar.0003s0245 | Vocar.0003s0245.1 | Vocar.0003s0245.1.p |
| PAC:32892865 | V. carteri | Vocar.0019s0142 | Vocar.0019s0142.1 | Vocar.0019s0142.1.p |
| PAC:32884331 | V. carteri | Vocar.0001s1049 | Vocar.0001s1049.1 | Vocar.0001s1049.1.p |
| PAC:32884633 | V. carteri | Vocar.0001s1441 | Vocar.0001s1441.1 | Vocar.0001s1441.1.p |

| | | | | |
|---|---|---|---|---|
| PAC:32898664 | V. carteri | Vocar.0018s0074 | Vocar.0018s0074.1 | Vocar.0018s0074.1.p |
| PAC:32887636 | V. carteri | Vocar.0053s0071 | Vocar.0053s0071.1 | Vocar.0053s0071.1.p |
| PAC:32887858 | V. carteri | Vocar.0007s0285 | Vocar.0007s0285.1 | Vocar.0007s0285.1.p |
| PAC:32886730 | V. carteri | Vocar.0020s0164 | Vocar.0020s0164.1 | Vocar.0020s0164.1.p |
| PAC:32887347 | V. carteri | Vocar.0029s0071 | Vocar.0029s0071.1 | Vocar.0029s0071.1.p |
| PAC:32889008 | V. carteri | Vocar.0003s0041 | Vocar.0003s0041.1 | Vocar.0003s0041.1.p |
| PAC:32889974 | V. carteri | Vocar.0004s0045 | Vocar.0004s0045.1 | Vocar.0004s0045.1.p |
| PAC:32892088 | V. carteri | Vocar.0042s0105 | Vocar.0042s0105.1 | Vocar.0042s0105.1.p |
| PAC:32884038 | V. carteri | Vocar.0005s0050 | Vocar.0005s0050.1 | Vocar.0005s0050.1.p |
| PAC:32895947 | V. carteri | Vocar.0021s0177 | Vocar.0021s0177.1 | Vocar.0021s0177.1.p |
| PAC:32893068 | V. carteri | Vocar.0034s0035 | Vocar.0034s0035.1 | Vocar.0034s0035.1.p |
| PAC:32888024 | V. carteri | Vocar.0007s0049 | Vocar.0007s0049.1 | Vocar.0007s0049.1.p |
| PAC:32887754 | V. carteri | Vocar.0045s0043 | Vocar.0045s0043.1 | Vocar.0045s0043.1.p |
| PAC:32884895 | V. carteri | Vocar.0001s1622 | Vocar.0001s1622.1 | Vocar.0001s1622.1.p |
| PAC:32887292 | V. carteri | Vocar.0029s0122 | Vocar.0029s0122.1 | Vocar.0029s0122.1.p |
| PAC:32887918 | V. carteri | Vocar.0007s0376 | Vocar.0007s0376.1 | Vocar.0007s0376.1.p |
| PAC:32895328 | V. carteri | Vocar.0011s0222 | Vocar.0011s0222.1 | Vocar.0011s0222.1.p |
| PAC:32889548 | V. carteri | Vocar.0004s0079 | Vocar.0004s0079.1 | Vocar.0004s0079.1.p |
| PAC:32889685 | V. carteri | Vocar.0004s0121 | Vocar.0004s0121.1 | Vocar.0004s0121.1.p |
| PAC:32894988 | V. carteri | Vocar.0012s0281 | Vocar.0012s0281.1 | Vocar.0012s0281.1.p |
| PAC:32887013 | V. carteri | Vocar.0051s0014 | Vocar.0051s0014.1 | Vocar.0051s0014.1.p |
| PAC:32884498 | V. carteri | Vocar.0001s1417 | Vocar.0001s1417.1 | Vocar.0001s1417.1.p |
| PAC:32889026 | V. carteri | Vocar.0003s0388 | Vocar.0003s0388.1 | Vocar.0003s0388.1.p |
| PAC:32888943 | V. carteri | Vocar.0003s0482 | Vocar.0003s0482.1 | Vocar.0003s0482.1.p |
| PAC:32887953 | V. carteri | Vocar.0007s0146 | Vocar.0007s0146.1 | Vocar.0007s0146.1.p |
| PAC:32897813 | V. carteri | Vocar.0039s0089 | Vocar.0039s0089.1 | Vocar.0039s0089.1.p |
| PAC:32888738 | V. carteri | Vocar.0041s0023 | Vocar.0041s0023.1 | Vocar.0041s0023.1.p |
| PAC:32889007 | V. carteri | Vocar.0003s0174 | Vocar.0003s0174.1 | Vocar.0003s0174.1.p |
| PAC:32888707 | V. carteri | Vocar.0041s0061 | Vocar.0041s0061.1 | Vocar.0041s0061.1.p |
| PAC:32886419 | V. carteri | Vocar.0033s0039 | Vocar.0033s0039.1 | Vocar.0033s0039.1.p |
| PAC:32898381 | V. carteri | Vocar.0025s0150 | Vocar.0025s0150.1 | Vocar.0025s0150.1.p |
| PAC:32894898 | V. carteri | Vocar.0012s0105 | Vocar.0012s0105.1 | Vocar.0012s0105.1.p |
| PAC:32887779 | V. carteri | Vocar.0045s0004 | Vocar.0045s0004.1 | Vocar.0045s0004.1.p |
| PAC:32893340 | V. carteri | Vocar.0009s0027 | Vocar.0009s0027.1 | Vocar.0009s0027.1.p |
| PAC:32888442 | V. carteri | Vocar.0014s0186 | Vocar.0014s0186.1 | Vocar.0014s0186.1.p |
| PAC:32898571 | V. carteri | Vocar.0018s0127 | Vocar.0018s0127.1 | Vocar.0018s0127.1.p |
| PAC:32884154 | V. carteri | Vocar.0001s0508 | Vocar.0001s0508.1 | Vocar.0001s0508.1.p |
| PAC:32886263 | V. carteri | Vocar.0040s0070 | Vocar.0040s0070.1 | Vocar.0040s0070.1.p |
| PAC:32893315 | V. carteri | Vocar.0009s0242 | Vocar.0009s0242.1 | Vocar.0009s0242.1.p |
| PAC:32889623 | V. carteri | Vocar.0004s0225 | Vocar.0004s0225.1 | Vocar.0004s0225.1.p |
| PAC:32885942 | V. carteri | Vocar.0001s0474 | Vocar.0001s0474.1 | Vocar.0001s0474.1.p |
| PAC:32894062 | V. carteri | Vocar.0006s0202 | Vocar.0006s0202.1 | Vocar.0006s0202.1.p |
| PAC:32889209 | V. carteri | Vocar.0003s0491 | Vocar.0003s0491.1 | Vocar.0003s0491.1.p |
| PAC:32889637 | V. carteri | Vocar.0004s0017 | Vocar.0004s0017.1 | Vocar.0004s0017.1.p |
| PAC:32887925 | V. carteri | Vocar.0007s0263 | Vocar.0007s0263.1 | Vocar.0007s0263.1.p |

| | | | | |
|---|---|---|---|---|
| PAC:32898468 | V. carteri | Vocar.0018s0223 | Vocar.0018s0223.1 | Vocar.0018s0223.1.p |
| PAC:32886337 | V. carteri | Vocar.0033s0162 | Vocar.0033s0162.1 | Vocar.0033s0162.1.p |
| PAC:32886100 | V. carteri | Vocar.0001s0944 | Vocar.0001s0944.1 | Vocar.0001s0944.1.p |
| PAC:32888258 | V. carteri | Vocar.0007s0345 | Vocar.0007s0345.1 | Vocar.0007s0345.1.p |
| PAC:32895117 | V. carteri | Vocar.0011s0293 | Vocar.0011s0293.1 | Vocar.0011s0293.1.p |
| PAC:32896111 | V. carteri | Vocar.0079s0013 | Vocar.0079s0013.1 | Vocar.0079s0013.1.p |
| PAC:32894156 | V. carteri | Vocar.0006s0257 | Vocar.0006s0257.1 | Vocar.0006s0257.1.p |
| PAC:32893316 | V. carteri | Vocar.0009s0250 | Vocar.0009s0250.1 | Vocar.0009s0250.1.p |
| PAC:32897420 | V. carteri | Vocar.0008s0294 | Vocar.0008s0294.1 | Vocar.0008s0294.1.p |
| PAC:32896047 | V. carteri | Vocar.0021s0144 | Vocar.0021s0144.1 | Vocar.0021s0144.1.p |
| PAC:32886503 | V. carteri | Vocar.0033s0157 | Vocar.0033s0157.1 | Vocar.0033s0157.1.p |
| PAC:32884304 | V. carteri | Vocar.0001s1220 | Vocar.0001s1220.1 | Vocar.0001s1220.1.p |
| PAC:32887258 | V. carteri | Vocar.0027s0117 | Vocar.0027s0117.1 | Vocar.0027s0117.1.p |
| PAC:32886056 | V. carteri | Vocar.0001s0729 | Vocar.0001s0729.1 | Vocar.0001s0729.1.p |
| PAC:32888855 | V. carteri | Vocar.0003s0318 | Vocar.0003s0318.1 | Vocar.0003s0318.1.p |
| PAC:32894317 | V. carteri | Vocar.0006s0057 | Vocar.0006s0057.1 | Vocar.0006s0057.1.p |
| PAC:32894022 | V. carteri | Vocar.0006s0097 | Vocar.0006s0097.1 | Vocar.0006s0097.1.p |
| PAC:32886094 | V. carteri | Vocar.0001s0806 | Vocar.0001s0806.1 | Vocar.0001s0806.1.p |
| PAC:32884731 | V. carteri | Vocar.0001s1517 | Vocar.0001s1517.1 | Vocar.0001s1517.1.p |
| PAC:32889072 | V. carteri | Vocar.0003s0354 | Vocar.0003s0354.1 | Vocar.0003s0354.1.p |
| PAC:32887720 | V. carteri | Vocar.0038s0031 | Vocar.0038s0031.1 | Vocar.0038s0031.1.p |
| PAC:32894277 | V. carteri | Vocar.0006s0136 | Vocar.0006s0136.1 | Vocar.0006s0136.1.p |
| PAC:32890139 | V. carteri | Vocar.0031s0011 | Vocar.0031s0011.1 | Vocar.0031s0011.1.p |
| PAC:32883636 | V. carteri | Vocar.0005s0199 | Vocar.0005s0199.1 | Vocar.0005s0199.1.p |
| PAC:32895284 | V. carteri | Vocar.0011s0085 | Vocar.0011s0085.1 | Vocar.0011s0085.1.p |
| PAC:32896173 | V. carteri | Vocar.0071s0021 | Vocar.0071s0021.1 | Vocar.0071s0021.1.p |
| PAC:32898524 | V. carteri | Vocar.0018s0171 | Vocar.0018s0171.1 | Vocar.0018s0171.1.p |
| PAC:32886126 | V. carteri | Vocar.0001s0002 | Vocar.0001s0002.1 | Vocar.0001s0002.1.p |
| PAC:32892761 | V. carteri | Vocar.0019s0265 | Vocar.0019s0265.1 | Vocar.0019s0265.1.p |
| PAC:32888772 | V. carteri | Vocar.0041s0030 | Vocar.0041s0030.1 | Vocar.0041s0030.1.p |
| PAC:32886025 | V. carteri | Vocar.0001s1065 | Vocar.0001s1065.1 | Vocar.0001s1065.1.p |
| PAC:32888710 | V. carteri | Vocar.0041s0069 | Vocar.0041s0069.1 | Vocar.0041s0069.1.p |
| PAC:32895245 | V. carteri | Vocar.0011s0226 | Vocar.0011s0226.1 | Vocar.0011s0226.1.p |
| PAC:32885881 | V. carteri | Vocar.0001s1568 | Vocar.0001s1568.1 | Vocar.0001s1568.1.p |
| PAC:32896498 | V. carteri | Vocar.0013s0295 | Vocar.0013s0295.1 | Vocar.0013s0295.1.p |
| PAC:32893201 | V. carteri | Vocar.0009s0344 | Vocar.0009s0344.1 | Vocar.0009s0344.1.p |

| Locus ID[a] | Gonidia1 | Gonidia2 | Somatic1 |
|---|---|---|---|
| Vocar.0054s0007 | 7.8 | 5.82 | 3.98 |
| Vocar.0024s0049 | 6.34 | 5.08 | 2.7 |
| Vocar.0001s1718 | 9.56 | 4.5 | 4.43 |
| Vocar.0007s0295 | 12.87 | 11.34 | 7.11 |
| Vocar.0012s0183 | 4.3 | 2.86 | 2.08 |
| Vocar.0001s0791 | 3.32 | 1.93 | 0.96 |
| Vocar.0006s0453 | 4.17 | 3.1 | 2.12 |
| Vocar.0014s0188 | 3.57 | 3.49 | 2.16 |
| Vocar.0021s0076 | 3.01 | 2.78 | 2.35 |
| Vocar.0031s0077 | 0.66 | 0.21 | 0.33 |
| Vocar.0006s0411 | 3.8 | 3.05 | 2.29 |
| Vocar.0012s0137 | 2.42 | 2.41 | 2.09 |
| Vocar.0001s0835 | 1.47 | 1.37 | 1.11 |
| Vocar.0005s0221 | 18.57 | 17.44 | 12.35 |
| Vocar.0011s0348 | 2.97 | 2.32 | 2.43 |
| Vocar.0029s0046 | 9.07 | 7.9 | 7.18 |
| Vocar.0004s0355 | 33.4 | 28.19 | 33.03 |
| Vocar.0004s0080 | 8.82 | 7.4 | 8.26 |
| Vocar.0001s1052 | 4.32 | 3.28 | 3.76 |
| Vocar.0086s0006 | 6.11 | 6.61 | 5.95 |
| Vocar.0023s0049 | 6.83 | 4.63 | 6.87 |
| Vocar.0042s0009 | 5.45 | 4.77 | 5.69 |
| Vocar.0021s0064 | 2.46 | 1.72 | 2.25 |
| Vocar.0018s0017 | 7.15 | 7.54 | 7.76 |
| Vocar.0019s0108 | 1.87 | 2.08 | 2.76 |
| Vocar.0025s0053 | 0.93 | 0.47 | 0.85 |
| Vocar.0021s0181 | 502 | 481.29 | 652.2 |
| Vocar.0006s0010 | 4.07 | 3.75 | 4.86 |
| Vocar.0028s0177 | 3.06 | 2.51 | 3.72 |
| Vocar.0011s0137 | 4.11 | 3.95 | 5.17 |
| Vocar.0006s0283 | 1.31 | 0.76 | 1.26 |
| Vocar.0027s0172 | 7.66 | 6.01 | 9.92 |
| Vocar.0028s0190 | 3.6 | 3.15 | 4.23 |
| Vocar.0018s0208 | 71.35 | 65.88 | 118.77 |
| Vocar.0007s0221 | 8.76 | 7.14 | 14.27 |
| Vocar.0041s0033 | 6.79 | 5.58 | 1.34 |
| Vocar.0024s0032 | 40.37 | 38.69 | 6.35 |
| Vocar.0011s0049 | 3.93 | 3.29 | 0.51 |
| Vocar.0079s0020 | 3.42 | 2.54 | 0.39 |
| Vocar.0007s0126 | 656.92 | 611.41 | 179.22 |
| Vocar.0003s0245 | 18.42 | 16.35 | 4.13 |
| Vocar.0019s0142 | 6.71 | 5.58 | 1.47 |
| Vocar.0001s1049 | 112.17 | 92.13 | 24.82 |
| Vocar.0001s1441 | 7.44 | 6.23 | 2.03 |

| | | | |
|---|---|---|---|
| Vocar.0018s0074 | 4.68 | 3.2 | 1.05 |
| Vocar.0053s0071 | 17.85 | 15.47 | 6.42 |
| Vocar.0007s0285 | 1.05 | 1.11 | 0 |
| Vocar.0020s0164 | 1.11 | 0.84 | 0 |
| Vocar.0029s0071 | 5.11 | 3.63 | 0.04 |
| Vocar.0003s0041 | 19.44 | 12.26 | 0.3 |
| Vocar.0004s0045 | 1.33 | 1.1 | 0.03 |
| Vocar.0042s0105 | 0.86 | 0.75 | 0.03 |
| Vocar.0005s0050 | 4.68 | 3.83 | 0.15 |
| Vocar.0021s0177 | 0.74 | 0.31 | 0 |
| Vocar.0034s0035 | 2 | 1.99 | 0 |
| Vocar.0007s0049 | 2.1 | 1.77 | 0.03 |
| Vocar.0045s0043 | 2.16 | 1.69 | 0.02 |
| Vocar.0001s1622 | 26.89 | 23.28 | 0.65 |
| Vocar.0029s0122 | 1.92 | 1.58 | 0.05 |
| Vocar.0007s0376 | 5.21 | 4.56 | 0.23 |
| Vocar.0011s0222 | 1.8 | 1.66 | 0.07 |
| Vocar.0004s0079 | 2.48 | 1.65 | 0.09 |
| Vocar.0004s0121 | 11.42 | 8.75 | 0.68 |
| Vocar.0012s0281 | 11.32 | 8.47 | 0.48 |
| Vocar.0051s0014 | 4.11 | 3.62 | 0.28 |
| Vocar.0001s1417 | 23.01 | 17.42 | 0.9 |
| Vocar.0003s0388 | 4.77 | 2.62 | 0.33 |
| Vocar.0003s0482 | 1.4 | 1.11 | 0.12 |
| Vocar.0007s0146 | 6.69 | 4.68 | 0.36 |
| Vocar.0039s0089 | 7.74 | 6.43 | 0.61 |
| Vocar.0041s0023 | 1.62 | 0.95 | 0.11 |
| Vocar.0003s0174 | 1.1 | 0.94 | 0.07 |
| Vocar.0041s0061 | 3.78 | 2.62 | 0.16 |
| Vocar.0033s0039 | 6.14 | 4.5 | 0.47 |
| Vocar.0025s0150 | 5.82 | 4.36 | 0.3 |
| Vocar.0012s0105 | 13.7 | 12.69 | 1.58 |
| Vocar.0045s0004 | 0.74 | 0.44 | 0.05 |
| Vocar.0009s0027 | 1.54 | 1.18 | 0.04 |
| Vocar.0014s0186 | 0.64 | 0.49 | 0.07 |
| Vocar.0018s0127 | 2.76 | 1.97 | 0.28 |
| Vocar.0001s0508 | 2.57 | 2.45 | 0.35 |
| Vocar.0040s0070 | 9.99 | 6.98 | 0.81 |
| Vocar.0009s0242 | 2.6 | 1.79 | 0.26 |
| Vocar.0004s0225 | 2.94 | 2.6 | 0.44 |
| Vocar.0001s0474 | 11.43 | 7.36 | 1.31 |
| Vocar.0006s0202 | 1.26 | 0.72 | 0.15 |
| Vocar.0003s0491 | 0.3 | 0.32 | 0.1 |
| Vocar.0004s0017 | 0.61 | 0.53 | 0.06 |
| Vocar.0007s0263 | 1.15 | 0.69 | 0.2 |

| | | | |
|---|---|---|---|
| Vocar.0018s0223 | 1.86 | 1.23 | 0.37 |
| Vocar.0033s0162 | 0.4 | 0.2 | 0 |
| Vocar.0001s0944 | 1.88 | 1.3 | 0.31 |
| Vocar.0007s0345 | 3.04 | 1.83 | 0.78 |
| Vocar.0011s0293 | 6.42 | 4.18 | 1.51 |
| Vocar.0079s0013 | 6 | 4.83 | 2.17 |
| Vocar.0006s0257 | 173.06 | 146.48 | 45.53 |
| Vocar.0009s0250 | 4.74 | 3.43 | 1.69 |
| Vocar.0008s0294 | 12.02 | 8.59 | 3.58 |
| Vocar.0021s0144 | 7.55 | 6.36 | 2.75 |
| Vocar.0033s0157 | 12.9 | 10.78 | 4.02 |
| Vocar.0001s1220 | 6.7 | 6.19 | 2.44 |
| Vocar.0027s0117 | 0.67 | 0.45 | 0.12 |
| Vocar.0001s0729 | 5.4 | 4.07 | 1.35 |
| Vocar.0003s0318 | 39.32 | 37.32 | 15.33 |
| Vocar.0006s0057 | 3.93 | 3.02 | 1.27 |
| Vocar.0006s0097 | 8.36 | 6.75 | 2.77 |
| Vocar.0001s0806 | 791.84 | 760.41 | 394.04 |
| Vocar.0001s1517 | 11.15 | 9.46 | 4.57 |
| Vocar.0003s0354 | 10.98 | 11.87 | 4.46 |
| Vocar.0038s0031 | 9.85 | 8.13 | 4.38 |
| Vocar.0006s0136 | 6.09 | 5.31 | 2.04 |
| Vocar.0031s0011 | 7.75 | 8.9 | 17.5 |
| Vocar.0005s0199 | 12.9 | 11.52 | 24.58 |
| Vocar.0011s0085 | 11.43 | 10.48 | 29.77 |
| Vocar.0071s0021 | 0.53 | 0.53 | 1.39 |
| Vocar.0018s0171 | 0.51 | 0.37 | 1.26 |
| Vocar.0001s0002 | 0.11 | 0.06 | 0.35 |
| Vocar.0019s0265 | 0.06 | 0.06 | 0.01 |
| Vocar.0041s0030 | 0.08 | 0 | 0 |
| Vocar.0001s1065 | 0.01 | 0.01 | 0.01 |
| Vocar.0041s0069 | 11.67 | 9.81 | 35.29 |
| Vocar.0011s0226 | 1.32 | 1.1 | 6.62 |
| Vocar.0001s1568 | 2.02 | 2.15 | 19.64 |
| Vocar.0013s0295 | 0.34 | 0.11 | 4.58 |
| Vocar.0009s0344 | 0.73 | 0.6 | 19.99 |

| Somatic2 | Gonidia | Somatic | Log2 Expression Ratio |
|---|---|---|---|
| 3.51 | 6.81 | 3.74 | -0.86 |
| 3.82 | 5.71 | 3.26 | -0.81 |
| 3.86 | 7.03 | 4.15 | -0.76 |
| 7.33 | 12.11 | 7.22 | -0.75 |
| 2.17 | 3.58 | 2.12 | -0.75 |
| 2.22 | 2.62 | 1.59 | -0.72 |
| 2.41 | 3.63 | 2.26 | -0.68 |
| 2.61 | 3.53 | 2.38 | -0.57 |
| 1.61 | 2.89 | 1.98 | -0.55 |
| 0.27 | 0.43 | 0.3 | -0.54 |
| 2.59 | 3.42 | 2.44 | -0.49 |
| 1.52 | 2.42 | 1.81 | -0.42 |
| 1.27 | 1.42 | 1.19 | -0.26 |
| 19.55 | 18 | 15.95 | -0.17 |
| 2.33 | 2.64 | 2.38 | -0.15 |
| 8.66 | 8.48 | 7.92 | -0.1 |
| 35.34 | 30.8 | 34.18 | 0.15 |
| 9.95 | 8.11 | 9.1 | 0.17 |
| 4.87 | 3.8 | 4.31 | 0.18 |
| 8.43 | 6.36 | 7.19 | 0.18 |
| 6.72 | 5.73 | 6.79 | 0.24 |
| 6.45 | 5.11 | 6.07 | 0.25 |
| 2.77 | 2.09 | 2.51 | 0.27 |
| 10.91 | 7.34 | 9.33 | 0.35 |
| 2.31 | 1.98 | 2.54 | 0.36 |
| 1.02 | 0.7 | 0.93 | 0.41 |
| 664.47 | 491.64 | 658.33 | 0.42 |
| 5.68 | 3.91 | 5.27 | 0.43 |
| 3.88 | 2.79 | 3.8 | 0.45 |
| 5.94 | 4.03 | 5.55 | 0.46 |
| 1.6 | 1.03 | 1.43 | 0.47 |
| 10.03 | 6.83 | 9.97 | 0.54 |
| 5.88 | 3.37 | 5.06 | 0.58 |
| 166.67 | 68.62 | 142.72 | 1.06 |
| 19.02 | 7.95 | 16.64 | 1.07 |
| 1.51 | 6.18 | 1.43 | -2.11 |
| 12.33 | 39.53 | 9.34 | -2.08 |
| 1.24 | 3.61 | 0.88 | -2.04 |
| 1.14 | 2.98 | 0.77 | -1.95 |
| 160.33 | 634.16 | 169.78 | -1.9 |
| 5.37 | 17.38 | 4.75 | -1.87 |
| 2 | 6.14 | 1.73 | -1.83 |
| 32.99 | 102.15 | 28.91 | -1.82 |
| 1.92 | 6.84 | 1.97 | -1.79 |

| | | | |
|---|---|---|---|
| 1.4 | 3.94 | 1.23 | -1.68 |
| 4.54 | 16.66 | 5.48 | -1.6 |
| 0 | 1.08 | 0 | -9.12 |
| 0 | 0.98 | 0 | -8.93 |
| 0.06 | 4.37 | 0.05 | -6.44 |
| 0.29 | 15.85 | 0.29 | -5.75 |
| 0.02 | 1.21 | 0.02 | -5.68 |
| 0 | 0.81 | 0.02 | -5.62 |
| 0.03 | 4.25 | 0.09 | -5.57 |
| 0.02 | 0.52 | 0.01 | -5.35 |
| 0.11 | 1.99 | 0.05 | -5.21 |
| 0.07 | 1.94 | 0.05 | -5.2 |
| 0.11 | 1.93 | 0.06 | -5 |
| 1.12 | 25.09 | 0.89 | -4.82 |
| 0.08 | 1.75 | 0.06 | -4.79 |
| 0.19 | 4.88 | 0.21 | -4.55 |
| 0.08 | 1.73 | 0.08 | -4.51 |
| 0.13 | 2.07 | 0.11 | -4.21 |
| 0.53 | 10.09 | 0.61 | -4.05 |
| 0.88 | 9.89 | 0.68 | -3.86 |
| 0.27 | 3.87 | 0.28 | -3.8 |
| 2.25 | 20.21 | 1.57 | -3.69 |
| 0.3 | 3.7 | 0.31 | -3.56 |
| 0.1 | 1.26 | 0.11 | -3.53 |
| 0.62 | 5.68 | 0.49 | -3.53 |
| 0.65 | 7.08 | 0.63 | -3.5 |
| 0.14 | 1.28 | 0.12 | -3.38 |
| 0.14 | 1.02 | 0.1 | -3.31 |
| 0.49 | 3.2 | 0.33 | -3.29 |
| 0.7 | 5.32 | 0.58 | -3.19 |
| 0.89 | 5.09 | 0.59 | -3.1 |
| 1.65 | 13.19 | 1.62 | -3.03 |
| 0.1 | 0.59 | 0.07 | -3 |
| 0.31 | 1.36 | 0.18 | -2.94 |
| 0.08 | 0.57 | 0.07 | -2.92 |
| 0.38 | 2.37 | 0.33 | -2.85 |
| 0.38 | 2.51 | 0.36 | -2.79 |
| 1.84 | 8.49 | 1.33 | -2.68 |
| 0.44 | 2.2 | 0.35 | -2.65 |
| 0.46 | 2.77 | 0.45 | -2.62 |
| 1.96 | 9.39 | 1.63 | -2.52 |
| 0.22 | 0.99 | 0.19 | -2.39 |
| 0 | 0.31 | 0.05 | -2.64 |
| 0.14 | 0.57 | 0.1 | -2.5 |
| 0.17 | 0.92 | 0.18 | -2.32 |

| | | | |
|---|---|---|---|
| 0.46 | 1.55 | 0.41 | -1.9 |
| 0.18 | 0.3 | 0.09 | -1.73 |
| 0.68 | 1.59 | 0.49 | -1.69 |
| 0.77 | 2.43 | 0.77 | -1.65 |
| 1.87 | 5.3 | 1.69 | -1.65 |
| 1.46 | 5.42 | 1.82 | -1.58 |
| 77.97 | 159.77 | 61.75 | -1.37 |
| 1.59 | 4.09 | 1.64 | -1.32 |
| 4.77 | 10.31 | 4.17 | -1.3 |
| 2.99 | 6.95 | 2.87 | -1.28 |
| 5.75 | 11.84 | 4.89 | -1.28 |
| 3 | 6.45 | 2.72 | -1.25 |
| 0.34 | 0.56 | 0.23 | -1.25 |
| 2.81 | 4.73 | 2.08 | -1.18 |
| 20.51 | 38.32 | 17.92 | -1.1 |
| 2.21 | 3.48 | 1.74 | -1 |
| 4.89 | 7.55 | 3.83 | -0.98 |
| 431.86 | 776.12 | 412.95 | -0.91 |
| 6.5 | 10.31 | 5.53 | -0.9 |
| 7.89 | 11.43 | 6.18 | -0.89 |
| 5.32 | 8.99 | 4.85 | -0.89 |
| 4.19 | 5.7 | 3.12 | -0.87 |
| 20.25 | 8.33 | 18.87 | 1.18 |
| 31.27 | 12.21 | 27.93 | 1.19 |
| 28.19 | 10.95 | 28.98 | 1.4 |
| 1.75 | 0.53 | 1.57 | 1.57 |
| 1.69 | 0.44 | 1.47 | 1.74 |
| 0.24 | 0.08 | 0.3 | 1.82 |
| 0 | 0.06 | 0 | -3.49 |
| 0.03 | 0.04 | 0.01 | -1.57 |
| 0.01 | 0.01 | 0.01 | -0.03 |
| 41.73 | 10.74 | 38.51 | 1.84 |
| 7.81 | 1.21 | 7.21 | 2.58 |
| 14.84 | 2.08 | 17.24 | 3.05 |
| 5.55 | 0.22 | 5.07 | 4.49 |
| 18.72 | 0.66 | 19.36 | 4.87 |

| **Cell-type expression p** |
| --- |
| Constitutive |
| Constitutive |
| Constitutive |
| Constitutive |
| Constitutive |
| Constitutive |
| Constitutive |
| Constitutive |
| Constitutive |
| Constitutive |
| Constitutive |
| Constitutive |
| Constitutive |
| Constitutive |
| Constitutive |
| Constitutive |
| Constitutive |
| Constitutive |
| Constitutive |
| Constitutive |
| Constitutive |
| Constitutive |
| Constitutive |
| Constitutive |
| Constitutive |
| Constitutive |
| Constitutive |
| Constitutive |
| Constitutive |
| Constitutive |
| Constitutive |
| Constitutive |
| Constitutive |
| Constitutive |
| Constitutive |
| Gonidial biased |
| Gonidial biased |
| Gonidial biased |
| Gonidial biased |
| Gonidial biased |
| Gonidial biased |
| Gonidial biased |
| Gonidial biased |
| Gonidial biased |

Gonidial biased
Gonidial biased
Gonidial specific
Gonidial specific
Gonidial specific
Gonidial specific
Gonidial specific
Gonidial specific
Gonidial specific
Gonidial specific
Gonidial specific
Gonidial specific
Gonidial specific
Gonidial specific
Gonidial specific
Gonidial specific
Gonidial specific
Gonidial specific
Gonidial specific
Gonidial specific
Gonidial specific
Gonidial specific
Gonidial specific
Gonidial specific
Gonidial specific
Gonidial specific
Gonidial specific
Gonidial specific
Gonidial specific
Gonidial specific
Gonidial specific
Gonidial specific
Gonidial specific
Gonidial specific
Gonidial specific
Gonidial specific
Gonidial specific
Gonidial specific
Gonidial specific
Gonidial specific
Gonidial specific
Gonidial specific
Low confidence
Low confidence
Low confidence

Low confidence
Low confidence
Low confidence
Low confidence
Low confidence
Low confidence
Low confidence
Low confidence
Low confidence
Low confidence
Low confidence
Low confidence
Low confidence
Low confidence
Low confidence
Low confidence
Low confidence
Low confidence
Low confidence
Low confidence
Low confidence
Low confidence
Low confidence
Low confidence
Low confidence
Low confidence
Low confidence
Low confidence
Low expression
Low expression
Low expression
Somatic biased
Somatic specific
Somatic specific
Somatic specific
Somatic specific

**defLine**
(1 of 2) PTHR13710:SF81 - ATP-DEPENDENT DNA HELICASE Q-LIKE 3
(1 of 1) K12833 - pre-mRNA branch site protein p14 (SF3B14)
(1 of 4) PTHR13620 - 3-5 EXONUCLEASE
(1 of 12) PF00098 - Zinc knuckle (zf-CCHC)
(1 of 1) K14573 - nucleolar protein 4 (NOP4, RBM28)
(1 of 1) K13093 - HIV Tat-specific factor 1 (HTATSF1)
(1 of 7) PTHR10288 - KH DOMAIN CONTAINING RNA BINDING PROTEIN
(1 of 1) K14807 - ATP-dependent RNA helicase DDX51/DBP6 [EC:3.6.4.13] (DDX51, DBP6)
(1 of 1) K11273 - chromosome transmission fidelity protein 1 [EC:3.6.4.13] (DDX11, CHL1, CTF1)
(1 of 1) K03257 - translation initiation factor 4A (EIF4A)
splicing factor like protein; hnRNP-F-like
(1 of 90) PF00076 - RNA recognition motif. (a.k.a. RRM, RBD, or RNP domain) (RRM_1)
(1 of 1) K11090 - lupus La protein (LA, SSB)
(1 of 1) KOG4307 - RNA binding protein RBM12/SWAN
(1 of 1) 2.7.7.8 - Polyribonucleotide nucleotidyltransferase / Polynucleotide phosphorylase
(1 of 1) K12599 - antiviral helicase SKI2 (SKI2, SKIV2L)
(1 of 13) PF01585 - G-patch domain (G-patch)
(1 of 1) PTHR24012:SF459 - RNA-BINDING PROTEIN 4F
(1 of 1) K12881 - THO complex subunit 4 (THOC4, ALY)
(1 of 1) PF00098//PF06839 - Zinc knuckle (zf-CCHC) // GRF zinc finger (zf-GRF)
(1 of 2) K11294 - nucleolin (NCL, NSR1)
(1 of 2) K14947 - epithelial splicing regulatory protein 1/2 (ESRP1_2)
(1 of 6) PF00570 - HRDC domain (HRDC)
(1 of 1) K16911 - ATP-dependent RNA helicase DDX21 [EC:3.6.4.13] (DDX21)
(1 of 90) PF00076 - RNA recognition motif. (a.k.a. RRM, RBD, or RNP domain) (RRM_1)
(1 of 90) PF00076 - RNA recognition motif. (a.k.a. RRM, RBD, or RNP domain) (RRM_1)
(1 of 90) PF00076 - RNA recognition motif. (a.k.a. RRM, RBD, or RNP domain) (RRM_1)
(1 of 1) PTHR23139:SF72 - RNA-BINDING PROTEIN 8A
(1 of 1) PF00076//PF04564//PF14259 - RNA recognition motif. (a.k.a. RRM, RBD, or RNP domain)
(1 of 1) K13161 - heterogeneous nuclear ribonucleoprotein R (HNRNPR)
(1 of 1) K02349 - DNA polymerase theta [EC:2.7.7.7] (POLQ)
(1 of 1) PTHR12183:SF11 - C3HC4 TYPE RING-FINGER PROTEIN-RELATED
(1 of 2) K17679 - ATP-dependent RNA helicase MSS116, mitochondrial [EC:3.6.4.13] (MSS116)
(1 of 2) K17679 - ATP-dependent RNA helicase MSS116, mitochondrial [EC:3.6.4.13] (MSS116)
(1 of 2) K14411 - RNA-binding protein Musashi (MSI)
(1 of 1) K12835 - ATP-dependent RNA helicase DDX42 (DDX42, SF3B125)
(1 of 1) PTHR24031:SF0 - ATP-DEPENDENT RNA HELICASE DDX28-RELATED
(1 of 1) KOG0144//KOG0145 - RNA-binding protein CUGBP1/BRUNO (RRM superfamily) // RNA-b
(1 of 90) PF00076 - RNA recognition motif. (a.k.a. RRM, RBD, or RNP domain) (RRM_1)
(1 of 2) PTHR23139//PTHR23139:SF58 - RNA-BINDING PROTEIN // SUBFAMILY NOT NAMED
(1 of 1) K14805 - ATP-dependent RNA helicase DDX24/MAK5 [EC:3.6.4.13] (DDX24, MAK5)
(1 of 1) K11884 - RNA-binding protein PNO1 (PNO1, DIM2)
(1 of 2) K13095 - splicing factor 1 (SF1)
(1 of 1) PTHR23139:SF8 - CLEAVAGE STIMULATION FACTOR 64 KILODALTON SUBUNIT

(1 of 1) PTHR11390:SF21 - DNA TOPOISOMERASE 3-ALPHA
(1 of 1) PTHR24031:SF303 - DEAD-BOX ATP-DEPENDENT RNA HELICASE 47, MITOCHONDRIAL
(1 of 90) PF00076 - RNA recognition motif. (a.k.a. RRM, RBD, or RNP domain) (RRM_1)
(1 of 1) PTHR15672 - CAMP-REGULATED PHOSPHOPROTEIN 21 RELATED R3H DOMAIN CONTAININ
(1 of 2) PTHR23149 - G PATCH DOMAIN CONTAINING PROTEIN
(1 of 3) K12900 - FUS-interacting serine-arginine-rich protein 1 (FUSIP1)
(1 of 1) K18663 - activating signal cointegrator complex subunit 3 [EC:3.6.4.12] (ASCC3)
(1 of 1) K13126 - polyadenylate-binding protein (PABPC)
(1 of 2) PTHR10501 - U1 SMALL NUCLEAR RIBONUCLEOPROTEIN A/U2 SMALL NUCLEAR RIBONUC
(1 of 1) KOG0947 - Cytoplasmic exosomal RNA helicase SKI2, DEAD-box superfamily
(1 of 1) PTHR24012:SF318 - RNA-BINDING (RRM/RBD/RNP MOTIFS) FAMILY PROTEIN
(1 of 1) PF00013//PF00226 - KH domain (KH_1) // DnaJ domain (DnaJ)
(1 of 12) PF00098 - Zinc knuckle (zf-CCHC)
(1 of 1) PTHR10887:SF366 - DNA-BINDING PROTEIN-RELATED
(1 of 5) K12823 - ATP-dependent RNA helicase DDX5/DBP2 (DDX5, DBP2)
(1 of 1) PTHR23003:SF3 - FI21236P1
(1 of 1) K11094 - U2 small nuclear ribonucleoprotein B'' (SNRPB2)
(1 of 90) PF00076 - RNA recognition motif. (a.k.a. RRM, RBD, or RNP domain) (RRM_1)
(1 of 16) PF00013 - KH domain (KH_1)
(1 of 2) PTHR10288:SF145 - P-ELEMENT SOMATIC INHIBITOR, ISOFORM C
(1 of 1) PTHR14074:SF16 - PROTEIN DRH-3
(1 of 1) K13103 - tuftelin-interacting protein 11 (TFIP11)
putative mitochondrial nuclease
(1 of 1) K12898 - heterogeneous nuclear ribonucleoprotein F/H (HNRNPF_H)
(1 of 1) PTHR24012:SF388 - SPLICING FACTOR, CC1-LIKE PROTEIN
(1 of 1) PF00098//PF00569//PF07910 - Zinc knuckle (zf-CCHC) // Zinc finger, ZZ type (ZZ) // Pepti
(1 of 16) PF00013 - KH domain (KH_1)
(1 of 1) K11093 - U1 small nuclear ribonucleoprotein 70kDa (SNRP70)
(1 of 2) K14837 - nucleolar protein 12 (NOP12)
(1 of 2) PTHR13710:SF81 - ATP-DEPENDENT DNA HELICASE Q-LIKE 3
(1 of 90) PF00076 - RNA recognition motif. (a.k.a. RRM, RBD, or RNP domain) (RRM_1)
(1 of 1) PTHR23273:SF12 - REPLICATION PROTEIN A 70 KDA DNA-BINDING SUBUNIT C-RELATED
(1 of 1) K10844 - DNA excision repair protein ERCC-2 (ERCC2, XPD)
(1 of 3) KOG4207 - Predicted splicing factor, SR protein superfamily
(1 of 1) K11135 - Pin2-interacting protein X1 (PINX1)
(1 of 2) K13207 - CUG-BP- and ETR3-like factor (CUGBP, BRUNOL, CELF)
(1 of 1) PTHR24012:SF492 - HETEROGENEOUS NUCLEAR RIBONUCLEOPROTEIN A0
(1 of 1) K13116 - ATP-dependent RNA helicase DDX41 [EC:3.6.4.13] (DDX41, ABS)
(1 of 1) K14401 - cleavage and polyadenylation specificity factor subunit 1 (CPSF1, CFT1)
(1 of 2) KOG1457 - RNA binding protein (contains RRM repeats)
(1 of 1) K10899 - ATP-dependent DNA helicase Q1 [EC:3.6.4.12] (RECQL)
(1 of 1) PTHR14089:SF2 - PRE-MRNA-SPLICING FACTOR CWC2
(1 of 2) 6.1.1.6 - Lysine--tRNA ligase / Lysyl-tRNA synthetase
(1 of 1) K02357 - elongation factor Ts (tsf, TSFM)
(1 of 14) K08857 - NIMA (never in mitosis gene a)-related kinase [EC:2.7.11.1] (NEK)

(1 of 1) PTHR23002 - ZINC FINGER CCHC DOMAIN CONTAINING PROTEIN
(1 of 2) K13095 - splicing factor 1 (SF1)
(1 of 2) K12598 - ATP-dependent RNA helicase DOB1 (MTR4, SKIV2L2)
(1 of 1) KOG0117 - Heterogeneous nuclear ribonucleoprotein R (RRM superfamily)
(1 of 1) K13117 - ATP-dependent RNA helicase DDX35 [EC:3.6.4.13] (DHX35)
(1 of 1) K03015 - DNA-directed RNA polymerase II subunit RPB7 (RPB7, POLR2G)
(1 of 1) K14806 - ATP-dependent RNA helicase DDX31/DBP7 [EC:3.6.4.13] (DDX31, DBP7)
(1 of 1) K13177 - ATP-dependent RNA helicase DDX1 [EC:3.6.4.13] (DDX1)
(1 of 1) KOG0326//KOG0327//KOG0328//KOG0331//KOG0335//KOG0340 - ATP-dependent RNA h
(1 of 90) PF00076 - RNA recognition motif. (a.k.a. RRM, RBD, or RNP domain) (RRM_1)
(1 of 1) PTHR13989 - REPLICATION PROTEIN A-RELATED
(1 of 1) KOG1994 - Predicted RNA binding protein, contains G-patch and Zn-finger domains
(1 of 1) PF05918 - Apoptosis inhibitory protein 5 (API5) (API5)
(1 of 77) 3.6.4.13 - RNA helicase
(1 of 2) PTHR23139:SF9 - SPLICING FACTOR U2AF 65 KDA SUBUNIT
(1 of 3) K12591 - exosome complex exonuclease RRP6 (RRP6, EXOSC10)
(1 of 3) K12591 - exosome complex exonuclease RRP6 (RRP6, EXOSC10)
(1 of 1) K12893 - splicing factor, arginine/serine-rich 4/5/6 (SFRS4_5_6)
(1 of 1) K12811 - ATP-dependent RNA helicase DDX46/PRP5 (DDX46, PRP5)
(1 of 1) K12619 - 5'-3' exoribonuclease 2 (XRN2, RAT1)
(1 of 1) PTHR23329//PTHR23329:SF9 - TUFTELIN-INTERACTING PROTEIN 11-RELATED // SUBFAM
(1 of 2) K14944 - RNA-binding protein Nova (NOVA)
(1 of 1) PTHR18806 - RBM25 PROTEIN
(1 of 1) K14787 - multiple RNA-binding domain-containing protein 1 (MRD1, RBM19)
(1 of 4) PTHR13620 - 3-5 EXONUCLEASE
(1 of 2) PTHR24031:SF249 - ATP-DEPENDENT DNA HELICASE RECG
(1 of 1) PTHR23329:SF2 - ZINC FINGER CCCH-TYPE WITH G PATCH DOMAIN-CONTAINING PROTEIN
(1 of 2) K14944 - RNA-binding protein Nova (NOVA)
(1 of 1) K14779 - ATP-dependent RNA helicase DDX52/ROK1 [EC:3.6.4.13] (DDX52, ROK1)
(1 of 1) PTHR17614 - ZINC FINGER-CONTAINING
(1 of 90) PF00076 - RNA recognition motif. (a.k.a. RRM, RBD, or RNP domain) (RRM_1)
(1 of 7) PTHR10288 - KH DOMAIN CONTAINING RNA BINDING PROTEIN
(1 of 7) PTHR10288 - KH DOMAIN CONTAINING RNA BINDING PROTEIN
(1 of 1) K18666 - activating signal cointegrator complex subunit 1 (ASCC1)
(1 of 2) PTHR14304 - P30 DBC PROTEIN
subunit of circadian RNA-binding protein

PFAM
PF00270; PF00271
PF00076
PF01612
PF00098
PF00076; PF14259
PF00076; PF14237
PF00013
PF00270; PF00271
PF13307; PF06733
PF00270; PF00271; PF04851
PF14259
PF00076
PF05383; PF14259; PF08777
PF14259
PF03726; PF03725; PF00013; PF00575; PF01138
PF08148; PF00270; PF00271; PF13234
PF01585
PF00076
PF00076
PF06839; PF00098
PF00076
PF14259
PF00570
PF08152; PF00270; PF00271
PF00076
PF00076
PF00076
PF00076
PF00076; PF04564; PF14259
PF00076
PF00270; PF00271; PF00476
PF00013
PF00270; PF00271
PF00270; PF00271
PF00076
PF00270; PF00271
PF00270; PF00271
PF00076
PF00076
PF00076
PF00270; PF00271
PF00013
PF16275; PF00076; PF00013
PF00076; PF14304; PF14327

PF06839; PF00098; PF01131; PF01751
PF00270; PF00271
PF00076
PF01424
PF01585
PF00076
PF02889; PF00270; PF00271
PF00658; PF00076
PF00076
PF08148; PF00270
PF00076
PF00226; PF00013
PF00098
PF13245; PF01424
PF00270; PF00271
PF00076
PF00076; PF13893
PF00076
PF00013
PF00013
PF00270; PF10258
PF12457; PF01585; PF07842
PF01223
PF14259
PF15519; PF00076; PF13893
PF00569; PF00098; PF07910
PF00013
PF12220; PF00076
PF00076; PF14259
PF00270; PF00271
PF00076
PF04057; PF08646; PF01336; PF16900
PF13307; PF04851; PF06777
PF00076
PF01585
PF00076
PF00076
PF00270; PF00271
PF10433; PF03178
PF00076; PF14259
PF00270; PF00271; PF00570
PF16131; PF00076
PF00152; PF01336
PF00889; PF00575
PF00076

PF00098
PF16275; PF00076; PF00013
PF08148; PF00270; PF00271; PF13234
PF00076
PF04408; PF07717; PF00270; PF00271
PF03876; PF00575
PF13959; PF00270; PF00271
PF00622; PF00270; PF00271
PF00270; PF00271
PF00076
PF08784; PF01336
PF01585; PF13821
PF05918; PF00076; PF07744
PF00270; PF00271
PF00076; PF14259
PF01612; PF00570
PF01612; PF00570
PF00076
PF00270; PF00271
PF03159
PF01585
PF00013
PF01480; PF00076
PF16651; PF00076; PF14259
PF01612
PF00270; PF00271; PF00929
PF01585
PF00013
PF00270; PF00271
PF01585; PF12171
PF00076
PF00013
PF00013
PF00013; PF10469
PF01585; PF14443; PF14444
PF00397; PF00013

PFAM_DEF
PF00271 - Helicase conserved C-terminal domain; PF00270 - DEAD/DEAH box helicase
PF00076 - RNA recognition motif. (a.k.a. RRM, RBD, or RNP domain)
PF01612 - 3'-5' exonuclease
PF00098 - Zinc knuckle
PF00076 - RNA recognition motif. (a.k.a. RRM, RBD, or RNP domain); PF14259 - RNA recognition r
PF00076 - RNA recognition motif. (a.k.a. RRM, RBD, or RNP domain); PF14237 - Domain of unknov
PF00013 - KH domain
PF00271 - Helicase conserved C-terminal domain; PF00270 - DEAD/DEAH box helicase
PF06733 - DEAD_2; PF13307 - Helicase C-terminal domain
PF00271 - Helicase conserved C-terminal domain; PF00270 - DEAD/DEAH box helicase; PF04851 - T
PF14259 - RNA recognition motif (a.k.a. RRM, RBD, or RNP domain)
PF00076 - RNA recognition motif. (a.k.a. RRM, RBD, or RNP domain)
PF14259 - RNA recognition motif (a.k.a. RRM, RBD, or RNP domain); PF08777 - RNA binding motif
PF14259 - RNA recognition motif (a.k.a. RRM, RBD, or RNP domain)
PF00575 - S1 RNA binding domain; PF01138 - 3' exoribonuclease family, domain 1; PF03725 - 3' e
PF13234 - rRNA-processing arch domain; PF00271 - Helicase conserved C-terminal domain; PF081
PF01585 - G-patch domain
PF00076 - RNA recognition motif. (a.k.a. RRM, RBD, or RNP domain)
PF00076 - RNA recognition motif. (a.k.a. RRM, RBD, or RNP domain)
PF00098 - Zinc knuckle; PF06839 - GRF zinc finger
PF00076 - RNA recognition motif. (a.k.a. RRM, RBD, or RNP domain)
PF14259 - RNA recognition motif (a.k.a. RRM, RBD, or RNP domain)
PF00570 - HRDC domain
PF08152 - GUCT (NUC152) domain; PF00271 - Helicase conserved C-terminal domain; PF00270 - D
PF00076 - RNA recognition motif. (a.k.a. RRM, RBD, or RNP domain)
PF00076 - RNA recognition motif. (a.k.a. RRM, RBD, or RNP domain)
PF00076 - RNA recognition motif. (a.k.a. RRM, RBD, or RNP domain)
PF00076 - RNA recognition motif. (a.k.a. RRM, RBD, or RNP domain)
PF00076 - RNA recognition motif. (a.k.a. RRM, RBD, or RNP domain); PF14259 - RNA recognition r
PF00076 - RNA recognition motif. (a.k.a. RRM, RBD, or RNP domain)
PF00476 - DNA polymerase family A; PF00271 - Helicase conserved C-terminal domain; PF00270 -
PF00013 - KH domain
PF00271 - Helicase conserved C-terminal domain; PF00270 - DEAD/DEAH box helicase
PF00271 - Helicase conserved C-terminal domain; PF00270 - DEAD/DEAH box helicase
PF00076 - RNA recognition motif. (a.k.a. RRM, RBD, or RNP domain)
PF00271 - Helicase conserved C-terminal domain; PF00270 - DEAD/DEAH box helicase
PF00271 - Helicase conserved C-terminal domain; PF00270 - DEAD/DEAH box helicase
PF00076 - RNA recognition motif. (a.k.a. RRM, RBD, or RNP domain)
PF00076 - RNA recognition motif. (a.k.a. RRM, RBD, or RNP domain)
PF00076 - RNA recognition motif. (a.k.a. RRM, RBD, or RNP domain)
PF00271 - Helicase conserved C-terminal domain; PF00270 - DEAD/DEAH box helicase
PF00013 - KH domain
PF00076 - RNA recognition motif. (a.k.a. RRM, RBD, or RNP domain); PF16275 - Splicing factor 1 h
PF00076 - RNA recognition motif. (a.k.a. RRM, RBD, or RNP domain); PF14327 - Hinge domain of c

PF00098 - Zinc knuckle; PF06839 - GRF zinc finger; PF01751 - Toprim domain; PF01131 - DNA topo
PF00271 - Helicase conserved C-terminal domain; PF00270 - DEAD/DEAH box helicase
PF00076 - RNA recognition motif. (a.k.a. RRM, RBD, or RNP domain)
PF01424 - R3H domain
PF01585 - G-patch domain
PF00076 - RNA recognition motif. (a.k.a. RRM, RBD, or RNP domain)
PF00271 - Helicase conserved C-terminal domain; PF02889 - Sec63 Brl domain; PF00270 - DEAD/D
PF00076 - RNA recognition motif. (a.k.a. RRM, RBD, or RNP domain); PF00658 - Poly-adenylate bi
PF00076 - RNA recognition motif. (a.k.a. RRM, RBD, or RNP domain)
PF08148 - DSHCT (NUC185) domain; PF00270 - DEAD/DEAH box helicase
PF00076 - RNA recognition motif. (a.k.a. RRM, RBD, or RNP domain)
PF00013 - KH domain; PF00226 - DnaJ domain
PF00098 - Zinc knuckle
PF01424 - R3H domain; PF13245 - Part of AAA domain
PF00271 - Helicase conserved C-terminal domain; PF00270 - DEAD/DEAH box helicase
PF00076 - RNA recognition motif. (a.k.a. RRM, RBD, or RNP domain)
PF00076 - RNA recognition motif. (a.k.a. RRM, RBD, or RNP domain); PF13893 - RNA recognition r
PF00076 - RNA recognition motif. (a.k.a. RRM, RBD, or RNP domain)
PF00013 - KH domain
PF00013 - KH domain
PF10258 - PHAX RNA-binding domain; PF00270 - DEAD/DEAH box helicase
PF07842 - GC-rich sequence DNA-binding factor-like protein; PF01585 - G-patch domain; PF12457
PF01223 - DNA/RNA non-specific endonuclease
PF14259 - RNA recognition motif (a.k.a. RRM, RBD, or RNP domain)
PF00076 - RNA recognition motif. (a.k.a. RRM, RBD, or RNP domain); PF15519 - linker between RF
PF07910 - Peptidase family C78; PF00098 - Zinc knuckle; PF00569 - Zinc finger, ZZ type
PF00013 - KH domain
PF00076 - RNA recognition motif. (a.k.a. RRM, RBD, or RNP domain); PF12220 - U1 small nuclear
PF00076 - RNA recognition motif. (a.k.a. RRM, RBD, or RNP domain); PF14259 - RNA recognition r
PF00271 - Helicase conserved C-terminal domain; PF00270 - DEAD/DEAH box helicase
PF00076 - RNA recognition motif. (a.k.a. RRM, RBD, or RNP domain)
PF16900 - Replication protein A OB domain; PF01336 - OB-fold nucleic acid binding domain; PF08
PF06777 - Helical and beta-bridge domain; PF13307 - Helicase C-terminal domain; PF04851 - Type
PF00076 - RNA recognition motif. (a.k.a. RRM, RBD, or RNP domain)
PF01585 - G-patch domain
PF00076 - RNA recognition motif. (a.k.a. RRM, RBD, or RNP domain)
PF00076 - RNA recognition motif. (a.k.a. RRM, RBD, or RNP domain)
PF00271 - Helicase conserved C-terminal domain; PF00270 - DEAD/DEAH box helicase
PF03178 - CPSF A subunit region; PF10433 - Mono-functional DNA-alkylating methyl methanesulfo
PF00076 - RNA recognition motif. (a.k.a. RRM, RBD, or RNP domain); PF14259 - RNA recognition r
PF00570 - HRDC domain; PF00271 - Helicase conserved C-terminal domain; PF00270 - DEAD/DEAH
PF00076 - RNA recognition motif. (a.k.a. RRM, RBD, or RNP domain); PF16131 - Torus domain
PF01336 - OB-fold nucleic acid binding domain; PF00152 - tRNA synthetases class II (D, K and N)
PF00889 - Elongation factor TS; PF00575 - S1 RNA binding domain
PF00076 - RNA recognition motif. (a.k.a. RRM, RBD, or RNP domain)

PF00098 - Zinc knuckle
PF00076 - RNA recognition motif. (a.k.a. RRM, RBD, or RNP domain); PF16275 - Splicing factor 1 h
PF13234 - rRNA-processing arch domain; PF00271 - Helicase conserved C-terminal domain; PF081
PF00076 - RNA recognition motif. (a.k.a. RRM, RBD, or RNP domain)
PF04408 - Helicase associated domain (HA2); PF00271 - Helicase conserved C-terminal domain; PF
PF00575 - S1 RNA binding domain; PF03876 - SHS2 domain found in N terminus of Rpb7p/Rpc25p,
PF13959 - Domain of unknown function (DUF4217); PF00271 - Helicase conserved C-terminal dom
PF00622 - SPRY domain; PF00271 - Helicase conserved C-terminal domain; PF00270 - DEAD/DEAH
PF00271 - Helicase conserved C-terminal domain; PF00270 - DEAD/DEAH box helicase
PF00076 - RNA recognition motif. (a.k.a. RRM, RBD, or RNP domain)
PF08784 - Replication protein A C terminal; PF01336 - OB-fold nucleic acid binding domain
PF01585 - G-patch domain; PF13821 - Domain of unknown function (DUF4187)
PF00076 - RNA recognition motif. (a.k.a. RRM, RBD, or RNP domain); PF07744 - SPOC domain; PF(
PF00271 - Helicase conserved C-terminal domain; PF00270 - DEAD/DEAH box helicase
PF00076 - RNA recognition motif. (a.k.a. RRM, RBD, or RNP domain); PF14259 - RNA recognition r
PF00570 - HRDC domain; PF01612 - 3'-5' exonuclease
PF00570 - HRDC domain; PF01612 - 3'-5' exonuclease
PF00076 - RNA recognition motif. (a.k.a. RRM, RBD, or RNP domain)
PF00271 - Helicase conserved C-terminal domain; PF00270 - DEAD/DEAH box helicase
PF03159 - XRN 5'-3' exonuclease N-terminus
PF01585 - G-patch domain
PF00013 - KH domain
PF00076 - RNA recognition motif. (a.k.a. RRM, RBD, or RNP domain); PF01480 - PWI domain
PF00076 - RNA recognition motif. (a.k.a. RRM, RBD, or RNP domain); PF14259 - RNA recognition r
PF01612 - 3'-5' exonuclease
PF00929 - Exonuclease; PF00271 - Helicase conserved C-terminal domain; PF00270 - DEAD/DEAH b
PF01585 - G-patch domain
PF00013 - KH domain
PF00271 - Helicase conserved C-terminal domain; PF00270 - DEAD/DEAH box helicase
PF01585 - G-patch domain; PF12171 - Zinc-finger double-stranded RNA-binding
PF00076 - RNA recognition motif. (a.k.a. RRM, RBD, or RNP domain)
PF00013 - KH domain
PF00013 - KH domain
PF00013 - KH domain; PF10469 - AKAP7 2'5' RNA ligase-like domain
PF01585 - G-patch domain; PF14444 - S1-like; PF14443 - DBC1
PF00397 - WW domain; PF00013 - KH domain

PANTHER
PTHR13710:SF81; PTHR13710
PTHR12785
PTHR13620; PTHR13620:SF0

PTHR24012
PTHR15608
PTHR10288
PTHR24031; PTHR24031:SF68
PTHR11472:SF41; PTHR11472
PTHR24031; PTHR24031:SF308
PTHR13976

PTHR22792:SF64; PTHR22792
PTHR13976
PTHR11252:SF0; PTHR11252
48 - DSHCT (NUC185) domain; PF
PTHR23106
PTHR24012:SF459; PTHR24012
PTHR19965

PTHR24012:SF374; PTHR24012
PTHR13976

PTHR24031; PTHR24031:SF330

PTHR23139:SF72; PTHR23139
PTHR22849
PTHR24012:SF374; PTHR24012
PTHR10133
PTHR12183; PTHR12183:SF11
PTHR24031; PTHR24031:SF232
PTHR24031; PTHR24031:SF232
PTHR24012:SF332; PTHR24012
PTHR24031; PTHR24031:SF125
PTHR24031; PTHR24031:SF0
PTHR24012
PTHR24012
PTHR23139; PTHR23139:SF58
PTHR24031; PTHR24031:SF91
PTHR12826
PTHR11208; PTHR11208:SF45
PTHR23139; PTHR23139:SF8

PTHR11390; PTHR11390:SF21
PTHR24031; PTHR24031:SF303

PTHR15672; PTHR15672:SF8
PTHR23149:SF9; PTHR23149
PTHR23147
PTHR24031; PTHR24031:SF249
PTHR24012; PTHR24012:SF491
PTHR10501

PTHR24012:SF318; PTHR24012
PTHR24078

PTHR10887; PTHR10887:SF366
PTHR24031
PTHR23003:SF3; PTHR23003
PTHR10501

PTHR10288:SF145; PTHR10288
PTHR14074; PTHR14074:SF16
PTHR23329:SF1; PTHR23329
PTHR13966
PTHR13976
PTHR24012:SF388; PTHR24012

PTHR13952:SF5; PTHR13952
PTHR24012
PTHR13710:SF81; PTHR13710
PTHR24012
PTHR23273:SF12; PTHR23273
PTHR11472:SF1; PTHR11472
PTHR24012
PTHR23149:SF9; PTHR23149
PTHR24012:SF359; PTHR24012
PTHR24012; PTHR24012:SF492
PTHR24031; PTHR24031:SF20
PTHR10644:SF2; PTHR10644
PTHR24012; PTHR24012:SF393
PTHR13710:SF72; PTHR13710
PTHR14089:SF2; PTHR14089
PTHR22594; PTHR22594:SF20
PTHR11741:SF0; PTHR11741

PTHR23002:SF69; PTHR23002
PTHR11208; PTHR11208:SF45
48 - DSHCT (NUC185) domain; PF
PTHR24012
PTHR18934
PTHR12709; PTHR12709:SF4
PTHR24031; PTHR24031:SF282
PTHR24031
PTHR24031

PTHR13989; PTHR13989:SF16
PTHR21032
PTHR24012

PTHR23139:SF9; PTHR23139
PTHR12124:SF41; PTHR12124
PTHR12124:SF41; PTHR12124
PTHR24012
PTHR24031; PTHR24031:SF25
PTHR12341; PTHR12341:SF30
PTHR23329; PTHR23329:SF9
PTHR10288; PTHR10288:SF142
PTHR18806
PTHR24012:SF477; PTHR24012
PTHR13620; PTHR13620:SF0
PTHR24031; PTHR24031:SF249
PTHR23329; PTHR23329:SF2
PTHR10288; PTHR10288:SF142
PTHR24031; PTHR24031:SF336
PTHR17614
PTHR24012
PTHR10288
PTHR10288
PTHR13360; PTHR13360:SF1
PTHR14304:SF11; PTHR14304
PTHR10288:SF145; PTHR10288

PANTHER_DEF
PTHR13710 - DNA HELICASE RECQ FAMILY MEMBER; PTHR13710:SF81 - ATP-DEPENDENT DNA HEI
PTHR12785 - FAMILY NOT NAMED
PTHR13620:SF0 - EXONUCLEASE 3'-5' DOMAIN-CONTAINING PROTEIN 2; PTHR13620 - 3-5 EXONU

PTHR24012 - FAMILY NOT NAMED
PTHR15608 - SPLICING FACTOR U2AF-ASSOCIATED PROTEIN 2
PTHR10288 - KH DOMAIN CONTAINING RNA BINDING PROTEIN
PTHR24031 - RNA HELICASE; PTHR24031:SF68 - ATP-DEPENDENT RNA HELICASE DDX51
PTHR11472:SF41 - ATP-DEPENDENT RNA HELICASE DDX11-RELATED; PTHR11472 - DNA REPAIR DE
PTHR24031:SF308 - SUBFAMILY NOT NAMED; PTHR24031 - RNA HELICASE
PTHR13976 - HETEROGENEOUS NUCLEAR RIBONUCLEOPROTEIN-RELATED

PTHR22792 - LUPUS LA PROTEIN-RELATED; PTHR22792:SF64 - LA PROTEIN 1-RELATED
PTHR13976 - HETEROGENEOUS NUCLEAR RIBONUCLEOPROTEIN-RELATED
PTHR11252 - POLYRIBONUCLEOTIDE NUCLEOTIDYLTRANSFERASE; PTHR11252:SF0 - POLYRIBONU
00270 - DEAD/DEAH box helicase
PTHR23106 - FAMILY NOT NAMED
PTHR24012 - FAMILY NOT NAMED; PTHR24012:SF459 - RNA-BINDING PROTEIN 4F
PTHR19965 - RNA AND EXPORT FACTOR BINDING PROTEIN

PTHR24012 - FAMILY NOT NAMED; PTHR24012:SF374 - SQUAMOUS CELL CARCINOMA ANTIGEN F
PTHR13976 - HETEROGENEOUS NUCLEAR RIBONUCLEOPROTEIN-RELATED

PTHR24031 - RNA HELICASE; PTHR24031:SF330 - DEAD-BOX ATP-DEPENDENT RNA HELICASE 7

PTHR23139 - RNA-BINDING PROTEIN; PTHR23139:SF72 - RNA-BINDING PROTEIN 8A
PTHR22849 - WDSAM1 PROTEIN
PTHR24012 - FAMILY NOT NAMED; PTHR24012:SF374 - SQUAMOUS CELL CARCINOMA ANTIGEN F
PTHR10133 - DNA POLYMERASE I
PTHR12183 - UNCHARACTERIZED RING ZINC FINGER-CONTAINING PROTEIN; PTHR12183:SF11 - C
PTHR24031:SF232 - ATP-DEPENDENT RNA HELICASE TDRD12-RELATED; PTHR24031 - RNA HELICAS
PTHR24031:SF232 - ATP-DEPENDENT RNA HELICASE TDRD12-RELATED; PTHR24031 - RNA HELICAS
PTHR24012 - FAMILY NOT NAMED; PTHR24012:SF332 - SUBFAMILY NOT NAMED
PTHR24031:SF125 - ATP-DEPENDENT RNA HELICASE DDX42; PTHR24031 - RNA HELICASE
PTHR24031:SF0 - ATP-DEPENDENT RNA HELICASE DDX28-RELATED; PTHR24031 - RNA HELICASE
PTHR24012 - FAMILY NOT NAMED
PTHR24012 - FAMILY NOT NAMED
PTHR23139 - RNA-BINDING PROTEIN; PTHR23139:SF58 - SUBFAMILY NOT NAMED
PTHR24031:SF91 - ATP-DEPENDENT RNA HELICASE DDX24; PTHR24031 - RNA HELICASE
PTHR12826 - FAMILY NOT NAMED
PTHR11208:SF45 - SPLICING FACTOR 1; PTHR11208 - RNA-BINDING PROTEIN RELATED
PTHR23139:SF8 - CLEAVAGE STIMULATION FACTOR 64 KILODALTON SUBUNIT; PTHR23139 - RNA-

PTHR11390:SF21 - DNA TOPOISOMERASE 3-ALPHA; PTHR11390 - PROKARYOTIC DNA TOPOISOME
PTHR24031:SF303 - DEAD-BOX ATP-DEPENDENT RNA HELICASE 47, MITOCHONDRIAL; PTHR24031

PTHR15672:SF8 - PROTEIN ENCORE; PTHR15672 - CAMP-REGULATED PHOSPHOPROTEIN 21 RELAT
PTHR23149:SF9 - PIN2/TERF1-INTERACTING TELOMERASE INHIBITOR 1; PTHR23149 - G PATCH DC
PTHR23147 - SERINE/ARGININE RICH SPLICING FACTOR
PTHR24031 - RNA HELICASE; PTHR24031:SF249 - ATP-DEPENDENT DNA HELICASE RECG
PTHR24012 - FAMILY NOT NAMED; PTHR24012:SF491 - FOUND IN NEURONS, ISOFORM A-RELATE
PTHR10501 - U1 SMALL NUCLEAR RIBONUCLEOPROTEIN A/U2 SMALL NUCLEAR RIBONUCLEOPRC

PTHR24012 - FAMILY NOT NAMED; PTHR24012:SF318 - RNA-BINDING (RRM/RBD/RNP MOTIFS) F
PTHR24078 - DNAJ HOMOLOG SUBFAMILY C MEMBER

PTHR10887:SF366 - DNA-BINDING PROTEIN-RELATED; PTHR10887 - DNA2/NAM7 HELICASE FAMIL
PTHR24031 - RNA HELICASE
PTHR23003 - RNA RECOGNITION MOTIF RRM DOMAIN CONTAINING PROTEIN; PTHR23003:SF3 -
PTHR10501 - U1 SMALL NUCLEAR RIBONUCLEOPROTEIN A/U2 SMALL NUCLEAR RIBONUCLEOPRC

PTHR10288 - KH DOMAIN CONTAINING RNA BINDING PROTEIN; PTHR10288:SF145 - P-ELEMENT S
PTHR14074:SF16 - PROTEIN DRH-3; PTHR14074 - HELICASE WITH DEATH DOMAIN-RELATED
PTHR23329:SF1 - TUFTELIN-INTERACTING PROTEIN 11; PTHR23329 - TUFTELIN-INTERACTING PRC
PTHR13966 - ENDONUCLEASE RELATED
PTHR13976 - HETEROGENEOUS NUCLEAR RIBONUCLEOPROTEIN-RELATED
PTHR24012 - FAMILY NOT NAMED; PTHR24012:SF388 - SPLICING FACTOR, CC1-LIKE PROTEIN

PTHR13952 - U1 SMALL NUCLEAR RIBONUCLEOPROTEIN 70 KD; PTHR13952:SF5 - U1 SMALL NUCL
PTHR24012 - FAMILY NOT NAMED
PTHR13710 - DNA HELICASE RECQ FAMILY MEMBER; PTHR13710:SF81 - ATP-DEPENDENT DNA HEI
PTHR24012 - FAMILY NOT NAMED
PTHR23273:SF12 - REPLICATION PROTEIN A 70 KDA DNA-BINDING SUBUNIT C-RELATED; PTHR232
PTHR11472:SF1 - TFIIH BASAL TRANSCRIPTION FACTOR COMPLEX HELICASE XPD SUBUNIT; PTHR1
PTHR24012 - FAMILY NOT NAMED
PTHR23149:SF9 - PIN2/TERF1-INTERACTING TELOMERASE INHIBITOR 1; PTHR23149 - G PATCH DC
PTHR24012:SF359 - SUBFAMILY NOT NAMED; PTHR24012 - FAMILY NOT NAMED
PTHR24012 - FAMILY NOT NAMED; PTHR24012:SF492 - HETEROGENEOUS NUCLEAR RIBONUCLEO
PTHR24031:SF20 - ATP-DEPENDENT RNA HELICASE DDX41-RELATED; PTHR24031 - RNA HELICASE
PTHR10644 - DNA REPAIR/RNA PROCESSING CPSF FAMILY; PTHR10644:SF2 - CLEAVAGE AND POL
PTHR24012 - FAMILY NOT NAMED; PTHR24012:SF393 - SUBFAMILY NOT NAMED
PTHR13710 - DNA HELICASE RECQ FAMILY MEMBER; PTHR13710:SF72 - ATP-DEPENDENT DNA HEL
PTHR14089 - CELL CYCLE CONTROL PROTEIN; PTHR14089:SF2 - PRE-MRNA-SPLICING FACTOR CWC
PTHR22594:SF20 - LYSINE--TRNA LIGASE; PTHR22594 - ASPARTYL/LYSYL-TRNA SYNTHETASE
PTHR11741 - ELONGATION FACTOR TS; PTHR11741:SF0 - ELONGATION FACTOR TS, MITOCHONDI

PTHR23002:SF69 - SUBFAMILY NOT NAMED; PTHR23002 - ZINC FINGER CCHC DOMAIN CONTAININ
PTHR11208:SF45 - SPLICING FACTOR 1; PTHR11208 - RNA-BINDING PROTEIN RELATED
00270 - DEAD/DEAH box helicase
PTHR24012 - FAMILY NOT NAMED
PTHR18934 - ATP-DEPENDENT RNA HELICASE
PTHR12709:SF4 - DNA-DIRECTED RNA POLYMERASE II SUBUNIT RPB7; PTHR12709 - DNA-DIRECTE
PTHR24031 - RNA HELICASE; PTHR24031:SF282 - ATP-DEPENDENT RNA HELICASE DBP7
PTHR24031 - RNA HELICASE
PTHR24031 - RNA HELICASE

PTHR13989:SF16 - RPA2 PROTEIN; PTHR13989 - REPLICATION PROTEIN A-RELATED
PTHR21032 - UNCHARACTERIZED
PTHR24012 - FAMILY NOT NAMED

PTHR23139:SF9 - SPLICING FACTOR U2AF 65 KDA SUBUNIT; PTHR23139 - RNA-BINDING PROTEIN
PTHR12124 - POLYMYOSITIS/SCLERODERMA AUTOANTIGEN-RELATED; PTHR12124:SF41 - POLYNU
PTHR12124 - POLYMYOSITIS/SCLERODERMA AUTOANTIGEN-RELATED; PTHR12124:SF41 - POLYNU
PTHR24012 - FAMILY NOT NAMED
PTHR24031:SF25 - PROTEIN F53H1.1, ISOFORM A; PTHR24031 - RNA HELICASE
PTHR12341 - 5'->3' EXORIBONUCLEASE; PTHR12341:SF30 - 5'-3' EXORIBONUCLEASE 2-RELATED
PTHR23329 - TUFTELIN-INTERACTING PROTEIN 11-RELATED; PTHR23329:SF9 - SUBFAMILY NOT N
PTHR10288:SF142 - PASILLA, ISOFORM D; PTHR10288 - KH DOMAIN CONTAINING RNA BINDING P
PTHR18806 - RBM25 PROTEIN
PTHR24012 - FAMILY NOT NAMED; PTHR24012:SF477 - PROTEIN RBD-1
PTHR13620:SF0 - EXONUCLEASE 3'-5' DOMAIN-CONTAINING PROTEIN 2; PTHR13620 - 3-5 EXONU
PTHR24031 - RNA HELICASE; PTHR24031:SF249 - ATP-DEPENDENT DNA HELICASE RECG
PTHR23329:SF2 - ZINC FINGER CCCH-TYPE WITH G PATCH DOMAIN-CONTAINING PROTEIN; PTHR2
PTHR10288:SF142 - PASILLA, ISOFORM D; PTHR10288 - KH DOMAIN CONTAINING RNA BINDING P
PTHR24031 - RNA HELICASE; PTHR24031:SF336 - ATP-DEPENDENT RNA HELICASE DDX52-RELATED
PTHR17614 - ZINC FINGER-CONTAINING
PTHR24012 - FAMILY NOT NAMED
PTHR10288 - KH DOMAIN CONTAINING RNA BINDING PROTEIN
PTHR10288 - KH DOMAIN CONTAINING RNA BINDING PROTEIN
PTHR13360:SF1 - ACTIVATING SIGNAL COINTEGRATOR 1 COMPLEX SUBUNIT 1; PTHR13360 - FAM
PTHR14304:SF11 - PROTEIN LST-3, ISOFORM A; PTHR14304 - P30 DBC PROTEIN
PTHR10288 - KH DOMAIN CONTAINING RNA BINDING PROTEIN; PTHR10288:SF145 - P-ELEMENT S

KOG
KOG0352
KOG0114
CLEASE

KOG0131

KOG0350
KOG1133

KOG4211

KOG4213
KOG4307
KOG1067

KOG4212; KOG0114; KOG0105; KOG0106
KOG0533

KOG0130; KOG0131
KOG4211

KOG0331

KOG0131
KOG0131
KOG0950
3HC4 TYPE RING-FINGER PROTEIN-RELATED
KOG0342
KOG0342
KOG0144

KOG0335
KOG0145; KOG0144

KOG0347
KOG3273

BINDING PROTEIN

RASE
KOG0330; KOG0340; KOG0331; KOG0327

TED R3H DOMAIN CONTAINING PROTEIN
OMAIN CONTAINING PROTEIN

KOG0123
KOG0114
KOG0947
KOG0131

LY
KOG0331
KOG0131; KOG0123; KOG0145; KOG0144
KOG4206

OMATIC INHIBITOR, ISOFORM C

KOG2184
KOG3721

KOG4206; KOG0114; KOG0124

KOG0113

KOG0352

73 - REPLICATION FACTOR A 1, RFA1
1472 - DNA REPAIR DEAD HELICASE RAD3/XP-D SUBFAMILY MEM
KOG4207
OMAIN CONTAINING PROTEIN
KOG0130
PROTEIN A0

KOG1896
KOG1457
LICASE Q1
C2

KOG1071

NG PROTEIN
KOG0119

KOG0131; KOG0117

KOG3298
KOG0348

KOG0340; KOG0331; KOG0335; KOG0327; KOG0326; KOG0328

KOG1994

JCLEOTIDYL TRANSFERASE, RIBONUCLEASE H FOLD PROTEIN WIT
JCLEOTIDYL TRANSFERASE, RIBONUCLEASE H FOLD PROTEIN WIT
KOG0106
KOG0331

AMED
ROTEIN
KOG2253
KOG0110
CLEASE

.3329 - TUFTELIN-INTERACTING PROTEIN 11-RELATED
ROTEIN
KOG0344

KOG2814

KOG2190

KOG_DEF
KOG0352 - ATP-dependent DNA helicase
KOG0114 - Predicted RNA-binding protein (RRM superfamily)

KOG0131 - Splicing factor 3b, subunit 4

KOG0350 - DEAD-box ATP-dependent RNA helicase
KOG1133 - Helicase of the DEAD superfamily

KOG4211 - Splicing factor hnRNP-F and related RNA-binding proteins

KOG4213 - RNA-binding protein La
KOG4307 - RNA binding protein RBM12/SWAN
KOG1067 - Predicted RNA-binding polyribonucleotide nucleotidyltransferase

KOG0106 - Alternative splicing factor SRp55/B52/SRp75 (RRM superfamily); KOG4212 - RNA-bindi
KOG0533 - RRM motif-containing protein

KOG0130 - RNA-binding protein RBM8/Tsunagi (RRM superfamily); KOG0131 - Splicing factor 3b,
KOG4211 - Splicing factor hnRNP-F and related RNA-binding proteins

KOG0331 - ATP-dependent RNA helicase

KOG0131 - Splicing factor 3b, subunit 4
KOG0131 - Splicing factor 3b, subunit 4
KOG0950 - DNA polymerase theta/eta, DEAD-box superfamily

KOG0342 - ATP-dependent RNA helicase pitchoune
KOG0342 - ATP-dependent RNA helicase pitchoune
KOG0144 - RNA-binding protein CUGBP1/BRUNO (RRM superfamily)

KOG0335 - ATP-dependent RNA helicase
KOG0145 - RNA-binding protein ELAV/HU (RRM superfamily); KOG0144 - RNA-binding protein CU

KOG0347 - RNA helicase
KOG3273 - Predicted RNA-binding protein Pno1p interacting with Nob1p and involved in 26S prote

KOG0330 - ATP-dependent RNA helicase; KOG0327 - Translation initiation factor 4F, helicase subu

KOG0123 - Polyadenylate-binding protein (RRM superfamily)
KOG0114 - Predicted RNA-binding protein (RRM superfamily)
KOG0947 - Cytoplasmic exosomal RNA helicase SKI2, DEAD-box superfamily
KOG0131 - Splicing factor 3b, subunit 4

KOG0331 - ATP-dependent RNA helicase
KOG0123 - Polyadenylate-binding protein (RRM superfamily); KOG0145 - RNA-binding protein ELA'
KOG4206 - Spliceosomal protein snRNP-U1A/U2B

KOG2184 - Tuftelin-interacting protein TIP39, contains G-patch domain
KOG3721 - Mitochondrial endonuclease

KOG0124 - Polypyrimidine tract-binding protein PUF60 (RRM superfamily); KOG4206 - Spliceosom

KOG0113 - U1 small nuclear ribonucleoprotein (RRM superfamily)

KOG0352 - ATP-dependent DNA helicase

BER
KOG4207 - Predicted splicing factor, SR protein superfamily

KOG0130 - RNA-binding protein RBM8/Tsunagi (RRM superfamily)

KOG1896 - mRNA cleavage and polyadenylation factor II complex, subunit CFT1 (CPSF subunit)
KOG1457 - RNA binding protein (contains RRM repeats)

KOG1071 - Mitochondrial translation elongation factor EF-Tsmt, catalyzes nucleotide exchange on

KOG0119 - Splicing factor 1/branch point binding protein (RRM superfamily)

KOG0117 - Heterogeneous nuclear ribonucleoprotein R (RRM superfamily); KOG0131 - Splicing fac

KOG3298 - DNA-directed RNA polymerase subunit E'
KOG0348 - ATP-dependent RNA helicase

KOG0335 - ATP-dependent RNA helicase; KOG0328 - Predicted ATP-dependent RNA helicase FAL1,

KOG1994 - Predicted RNA binding protein, contains G-patch and Zn-finger domains

'H HRDC DOMAIN
'H HRDC DOMAIN
KOG0106 - Alternative splicing factor SRp55/B52/SRp75 (RRM superfamily)
KOG0331 - ATP-dependent RNA helicase

KOG2253 - U1 snRNP complex, subunit SNU71 and related PWI-motif proteins
KOG0110 - RNA-binding protein (RRM superfamily)

KOG0344 - ATP-dependent RNA helicase

KOG2814 - Transcription coactivator complex, P50 component (LigT RNA ligase/phosphodiesterase

KOG2190 - PolyC-binding proteins alphaCP-1 and related KH domain proteins

| EC | EC_DEF | KEGG |
|---|---|---|
| 3.6.4.12 | 3.6.4.12 - DNA helicase | |
| | | K12833 |
| 3.6.4.12 | 3.6.4.12 - DNA helicase | |
| | | K14573 |
| | | K13093 |
| 3.6.4.13 | 3.6.4.13 - RNA helicase | K14807 |
| 3.6.4.13 | 3.6.4.13 - RNA helicase | K11273 |
| 3.6.4.13 | 3.6.4.13 - RNA helicase | K03257 |
| | | K14947 |
| | | K11090 |
| 2.7.7.8 | 2.7.7.8 - Polyribonucleotide nucleotidyltransferase | K00962 |
| 3.6.4.13 | 3.6.4.13 - RNA helicase | K12599 |
| ing protein hnRNP-M; KOG0105 - Alternative splicing factor ASF/SF2 (RRM supe | | |
| | | K12881 |
| 5.99.1.2 | 5.99.1.2 - DNA topoisomerase | |
| subunit 4 | | K11294 |
| | | K14947 |
| 3.6.4.12 | 3.6.4.12 - DNA helicase | |
| 3.6.4.13 | 3.6.4.13 - RNA helicase | K16911 |
| | | K13161 |
| 2.7.7.7 | 2.7.7.7 - DNA-directed DNA polymerase | K02349 |
| 3.6.4.13 | 3.6.4.13 - RNA helicase | K17679 |
| 3.6.4.13 | 3.6.4.13 - RNA helicase | K17679 |
| | | K14411 |
| 3.6.4.13 | 3.6.4.13 - RNA helicase | K12835 |
| 3.6.4.13 | 3.6.4.13 - RNA helicase | K12823 |
| GBP1/BRUNO (RRM superfamily) | | |
| | | K12837 |
| 3.6.4.13 | 3.6.4.13 - RNA helicase | K14805 |
| asome assembly | | K11884 |
| | | K13095 |

| | | |
|---|---|---|
| 5.99.1.2 | 5.99.1.2 - DNA topoisomerase | K03165 |
| 3.6.4.13 | 3.6.4.13 - RNA helicase | |
| | | K12900 |
| 3.6.4.13 | 3.6.4.13 - RNA helicase | K18663 |
| | | K13126 |
| 3.6.4.13 | 3.6.4.13 - RNA helicase | |
| | | K12741 |
| 3.6.4.12; 3.6.4.13 | 3.6.4.13 - RNA helicase; 3.6.4.12 - DNA helicase | |
| 3.6.4.13 | 3.6.4.13 - RNA helicase | K12823 |
| V/HU (RRM superfamily); KOG0144 - RNA-binding protein CUGBP1/BRUNO (RR | | |
| | | K11094 |
| 3.6.4.13 | 3.6.4.13 - RNA helicase | |
| | | K13103 |
| | | K01173 |
| | | K12898 |
| al protein snRNP-U1A/U2B; KOG0114 - Predicted RNA-binding protein ( | | K13091 |
| | | K11093 |
| | | K14837 |
| 3.6.4.12 | 3.6.4.12 - DNA helicase | |
| | | K07466 |
| 3.6.4.12 | 3.6.4.12 - DNA helicase | K10844 |
| | | K11135 |
| | | K13207 |
| | | K12741 |
| 3.6.4.13 | 3.6.4.13 - RNA helicase | K13116 |
| | | K14401 |
| 3.6.4.12 | 3.6.4.12 - DNA helicase | K10899 |
| 6.1.1.6 | 6.1.1.6 - Lysine--tRNA ligase | K04567 |
| EF-Tumt | | K02357 |
| | | K08857 |

| | | | |
|---|---|---|---|
| | | | K13095 |
| 3.6.4.13 | 3.6.4.13 - RNA helicase | | K12598 |
| tor 3b, subunit 4 | | | |
| 3.6.4.13 | 3.6.4.13 - RNA helicase | | K13117 |
| 2.7.7.6 | 2.7.7.6 - DNA-directed RNA polymerase | | K03015 |
| 3.6.4.13 | 3.6.4.13 - RNA helicase | | K14806 |
| 3.6.4.13 | 3.6.4.13 - RNA helicase | | K13177 |
| 3.6.4.13 | 3.6.4.13 - RNA helicase | | |
| 3.6.4.13 | 3.6.4.13 - RNA helicase | | |
| | | | K12837 |
| 3.1.13.5 | 3.1.13.5 - Ribonuclease D | | K12591 |
| 3.1.13.5 | 3.1.13.5 - Ribonuclease D | | K12591 |
| | | | K12893 |
| 3.6.4.13 | 3.6.4.13 - RNA helicase | | K12811 |
| | | | K12619 |
| | | | K14944 |
| | | | K14787 |
| 3.6.4.12 | 3.6.4.12 - DNA helicase | | |
| | | | K14944 |
| 3.6.4.13 | 3.6.4.13 - RNA helicase | | K14779 |
| e family) | | | K18666 |
| | | | K13210 |

KEGG_DEF

K12833 - pre-mRNA branch site protein p14

K14573 - nucleolar protein 4
K13093 - ""

K14807 - ""
K11273 - ""
K03257 - translation initiation factor 4A
K14947 - ""

K11090 - lupus La protein

K00962 - polyribonucleotide nucleotidyltransferase
K12599 - antiviral helicase SKI2

rfamily); KOG0114 - Predicted RNA-binding protein (RRM superfamil
K12881 - THO complex subunit 4

K11294 - nucleolin
K14947 - ""

K16911 - ""

K13161 - ""
K02349 - ""

K17679 - ""
K17679 - ""
K14411 - RNA-binding protein Musashi
K12835 - ATP-dependent RNA helicase DDX42
K12823 - ATP-dependent RNA helicase DDX5/DBP2

K12837 - splicing factor U2AF 65 kDa subunit
K14805 - ""
K11884 - ""
K13095 - ""

K03165 - DNA topoisomerase III

K12900 - FUS-interacting serine-arginine-rich protein 1
K18663 - ""
K13126 - polyadenylate-binding protein

K12741 - heterogeneous nuclear ribonucleoprotein A1/A3

K12823 - ATP-dependent RNA helicase DDX5/DBP2
M superfamily); KOG0131 - Splicing factor 3b, subunit 4
K11094 - U2 small nuclear ribonucleoprotein B''

K13103 - ""
K01173 - endonuclease G, mitochondrial
K12898 - ""
K13091 - ""

K11093 - U1 small nuclear ribonucleoprotein 70kDa
K14837 - ""

K07466 - replication factor A1
K10844 - DNA excision repair protein ERCC-2

K11135 - ""
K13207 - ""
K12741 - heterogeneous nuclear ribonucleoprotein A1/A3
K13116 - ""
K14401 - cleavage and polyadenylation specificity factor subunit 1

K10899 - ""

K04567 - lysyl-tRNA synthetase, class II
K02357 - ""
K08857 - ""

K13095 - ""
K12598 - ATP-dependent RNA helicase DOB1

K13117 - ""
K03015 - DNA-directed RNA polymerase II subunit RPB7
K14806 - ""
K13177 - ""

K12837 - splicing factor U2AF 65 kDa subunit
K12591 - exosome complex exonuclease RRP6
K12591 - exosome complex exonuclease RRP6
K12893 - splicing factor, arginine/serine-rich 4/5/6
K12811 - ATP-dependent RNA helicase DDX46/PRP5
K12619 - 5'-3' exoribonuclease 2

K14944 - ""

K14787 - ""

K14944 - ""
K14779 - ""

K18666 - ""

K13210 - ""

GO
GO:0003676; GO:0005524; GO:0006310; GO:0008026
GO:0000166; GO:0003676
GO:0003676; GO:0008408; GO:0006139
GO:0003676; GO:0008270
GO:0000166; GO:0003676
GO:0000166; GO:0003676
GO:0003676; GO:0003723
GO:0003676; GO:0005524
GO:0003676; GO:0003677; GO:0005524; GO:0016818; GO:0008026; GO:0004003; GO:0006139
GO:0003676; GO:0003677; GO:0005524; GO:0016787
GO:0000166; GO:0003676
GO:0000166; GO:0003676
GO:0000166; GO:0003676; GO:0005634; GO:0003723; GO:0030529; GO:0006396
GO:0000166; GO:0003676; GO:0042309; GO:0050825; GO:0050826
GO:0003676; GO:0000175; GO:0003723; GO:0006396; GO:0006402; GO:0004654
GO:0003676; GO:0005524
GO:0003676
GO:0000166; GO:0003676
GO:0000166; GO:0003676
GO:0003676; GO:0008270
GO:0000166; GO:0003676
GO:0000166; GO:0003676
GO:0000166; GO:0003676; GO:0003824; GO:0044237; GO:0005622
GO:0003676; GO:0005524; GO:0005634; GO:0003723; GO:0004386
GO:0000166; GO:0003676; GO:0005488
GO:0000166; GO:0003676
GO:0000166; GO:0003676; GO:0042309; GO:0050825; GO:0050826
GO:0000166; GO:0003676
GO:0003676; GO:0004842; GO:0016567
GO:0000166; GO:0003676
GO:0003676; GO:0003677; GO:0005524; GO:0003887; GO:0006260
GO:0003676; GO:0003723
GO:0003676; GO:0005524
GO:0003676; GO:0005524
GO:0000166; GO:0003676
GO:0003676; GO:0005524
GO:0003676; GO:0005524
GO:0000166; GO:0003676
GO:0000166; GO:0003676
GO:0000166; GO:0003676
GO:0003676; GO:0005524
GO:0003676; GO:0003723
GO:0000166; GO:0003676; GO:0003723; GO:0045131; GO:0000398
GO:0000166; GO:0003676; GO:0031124

GO:0003676; GO:0003677; GO:0008270; GO:0003916; GO:0003917; GO:0006265
GO:0003676; GO:0005524
GO:0000166; GO:0003676
GO:0003676
GO:0003676
GO:0000166; GO:0003676
GO:0003676; GO:0005524
GO:0003676; GO:0003723
GO:0000166; GO:0003676; GO:0017069; GO:0000398
GO:0003676; GO:0005524
GO:0000166; GO:0003676
GO:0003676; GO:0003723
GO:0003676; GO:0008270
GO:0003676
GO:0003676; GO:0005524
GO:0000166; GO:0003676
GO:0000166; GO:0003676; GO:0017069; GO:0000398
GO:0000166; GO:0003676
GO:0003676; GO:0003723
GO:0003676; GO:0003723
GO:0003676; GO:0005524
GO:0003676; GO:0003677; GO:0005634; GO:0006355
GO:0003676; GO:0046872; GO:0016787
GO:0000166; GO:0003676
GO:0000166; GO:0003676
GO:0003676; GO:0008270
GO:0003723
GO:0000166; GO:0003676
GO:0000166; GO:0003676
GO:0003676; GO:0003677; GO:0005524; GO:0005634; GO:0006351; GO:0006310; GO:0008026
GO:0003676
GO:0003676; GO:0003677; GO:0005634; GO:0006281; GO:0006260; GO:0006310
GO:0003676; GO:0003677; GO:0005524; GO:0005634; GO:0016818; GO:0008026; GO:0006289; GC
GO:0000166; GO:0003676
GO:0003676
GO:0000166; GO:0003676
GO:0000166; GO:0003676
GO:0003676; GO:0005524; GO:0008270
GO:0003676; GO:0005634
GO:0000166; GO:0003676
GO:0000166; GO:0003676; GO:0005524; GO:0003824; GO:0006310; GO:0008026; GO:0044237; GC
GO:0000166; GO:0003676; GO:0046872
GO:0000166; GO:0003676; GO:0005524; GO:0006418; GO:0004812; GO:0005737; GO:0004824; GC
GO:0003676; GO:0005515; GO:0003723; GO:0003735; GO:0003746; GO:0005840; GO:0006414; GC
GO:0000166; GO:0003676

GO:0003676; GO:0008270
GO:0000166; GO:0003676; GO:0008270; GO:0003723; GO:0045131; GO:0000398
GO:0003676; GO:0005524
GO:0000166; GO:0003676
GO:0003676; GO:0005524; GO:0004386
GO:0003676; GO:0003899; GO:0006351
GO:0003676; GO:0005524
GO:0003676; GO:0005524; GO:0005515
GO:0003676; GO:0005524
GO:0003676
GO:0003676; GO:0003677; GO:0005634; GO:0006281; GO:0006260; GO:0006310
GO:0003676
GO:0003676
GO:0003676; GO:0005524
GO:0000166; GO:0003676
GO:0000166; GO:0003676; GO:0008408; GO:0003824; GO:0044237; GO:0006139; GO:0005622
GO:0000166; GO:0003676; GO:0008408; GO:0003824; GO:0044237; GO:0006139; GO:0005622
GO:0000166; GO:0003676
GO:0003676; GO:0005524; GO:0003723
GO:0003676; GO:0004527
GO:0003676
GO:0003676; GO:0003723
GO:0003676; GO:0006397
GO:0000166; GO:0003676
GO:0003676; GO:0008408; GO:0006139
GO:0003676; GO:0005524
GO:0003676
GO:0003676; GO:0003723
GO:0003676; GO:0005524
GO:0003676; GO:0046872
GO:0000166; GO:0003676; GO:0005515
GO:0003676; GO:0003723
GO:0003676; GO:0003723
GO:0003676; GO:0003723
GO:0003676; GO:0042309; GO:0050825; GO:0006355; GO:0050826
GO:0005515; GO:0003723

| GO_DEF | PATHWAY_DEF |
|---|---|
| GO:0003676 - Interacting selectively and non-covalently with any nucleic acid.; | |
| GO:0003676 - Interacting selectively and non-covalently with any nucleic acid.; | |
| GO:0006139 - Any cellular metabolic process involving nucleobases, nucleoside | |
| GO:0003676 - Interacting selectively and non-covalently with any nucleic acid.; | |
| GO:0003676 - Interacting selectively and non-covalently with any nucleic acid.; | |
| GO:0003676 - Interacting selectively and non-covalently with any nucleic acid.; | |
| GO:0003723 - Interacting selectively and non-covalently with an RNA molecule | |
| GO:0003676 - Interacting selectively and non-covalently with any nucleic acid.; | |
| GO:0016818 - Catalysis of the hydrolysis of any acid anhydride which contains p | |
| GO:0016787 - Catalysis of the hydrolysis of various bonds, e.g. C-O, C-N, C-C, ph | |
| GO:0003676 - Interacting selectively and non-covalently with any nucleic acid.; | |
| GO:0003676 - Interacting selectively and non-covalently with any nucleic acid.; | |
| GO:0003723 - Interacting selectively and non-covalently with an RNA molecule | |
| GO:0003676 - Interacting selectively and non-covalently with any nucleic acid.; | |
| GO:0006402 - The chemical reactions and pathways resulting in the breakdown | |
| GO:0003676 - Interacting selectively and non-covalently with any nucleic acid.; | |
| GO:0003676 - Interacting selectively and non-covalently with any nucleic acid. | |
| GO:0003676 - Interacting selectively and non-covalently with any nucleic acid.; | |
| GO:0003676 - Interacting selectively and non-covalently with any nucleic acid.; | |
| GO:0003676 - Interacting selectively and non-covalently with any nucleic acid.; | |
| GO:0003676 - Interacting selectively and non-covalently with any nucleic acid.; | |
| GO:0003676 - Interacting selectively and non-covalently with any nucleic acid.; | |
| GO:0003676 - Interacting selectively and non-covalently with any nucleic acid.; | |
| GO:0004386 - Catalysis of the reaction: NTP + H2O = NDP + phosphate, to drive | |
| GO:0003676 - Interacting selectively and non-covalently with any nucleic acid.; | |
| GO:0003676 - Interacting selectively and non-covalently with any nucleic acid.; | |
| GO:0003676 - Interacting selectively and non-covalently with any nucleic acid.; | |
| GO:0003676 - Interacting selectively and non-covalently with any nucleic acid.; | |
| GO:0016567 - The process in which one or more ubiquitin groups are added to | |
| GO:0003676 - Interacting selectively and non-covalently with any nucleic acid.; | |
| GO:0003887 - Catalysis of the reaction: deoxynucleoside triphosphate + DNA(n) | |
| GO:0003723 - Interacting selectively and non-covalently with an RNA molecule | |
| GO:0003676 - Interacting selectively and non-covalently with any nucleic acid.; | |
| GO:0003676 - Interacting selectively and non-covalently with any nucleic acid.; | |
| GO:0003676 - Interacting selectively and non-covalently with any nucleic acid.; | |
| GO:0003676 - Interacting selectively and non-covalently with any nucleic acid.; | |
| GO:0003676 - Interacting selectively and non-covalently with any nucleic acid.; | |
| GO:0003676 - Interacting selectively and non-covalently with any nucleic acid.; | |
| GO:0003676 - Interacting selectively and non-covalently with any nucleic acid.; | |
| GO:0003676 - Interacting selectively and non-covalently with any nucleic acid.; | |
| GO:0003676 - Interacting selectively and non-covalently with any nucleic acid.; | |
| GO:0003723 - Interacting selectively and non-covalently with an RNA molecule | |
| GO:0003723 - Interacting selectively and non-covalently with an RNA molecule | |
| GO:0003676 - Interacting selectively and non-covalently with any nucleic acid.; | |

GO:0003916 - Catalysis of the transient cleavage and passage of individual DN/
GO:0003676 - Interacting selectively and non-covalently with any nucleic acid.;
GO:0003676 - Interacting selectively and non-covalently with any nucleic acid.;
GO:0003676 - Interacting selectively and non-covalently with any nucleic acid.
GO:0003676 - Interacting selectively and non-covalently with any nucleic acid.
GO:0003676 - Interacting selectively and non-covalently with any nucleic acid.;
GO:0003676 - Interacting selectively and non-covalently with any nucleic acid.;
GO:0003723 - Interacting selectively and non-covalently with an RNA molecule
GO:0017069 - Interacting selectively and non-covalently with a small nuclear R
GO:0003676 - Interacting selectively and non-covalently with any nucleic acid.;
GO:0003676 - Interacting selectively and non-covalently with any nucleic acid.;
GO:0003723 - Interacting selectively and non-covalently with an RNA molecule
GO:0003676 - Interacting selectively and non-covalently with any nucleic acid.;
GO:0003676 - Interacting selectively and non-covalently with any nucleic acid.
GO:0003676 - Interacting selectively and non-covalently with any nucleic acid.;
GO:0003676 - Interacting selectively and non-covalently with any nucleic acid.;
GO:0017069 - Interacting selectively and non-covalently with a small nuclear R
GO:0003676 - Interacting selectively and non-covalently with any nucleic acid.;
GO:0003723 - Interacting selectively and non-covalently with an RNA molecule
GO:0003723 - Interacting selectively and non-covalently with an RNA molecule
GO:0003676 - Interacting selectively and non-covalently with any nucleic acid.;
GO:0003676 - Interacting selectively and non-covalently with any nucleic acid.;
GO:0016787 - Catalysis of the hydrolysis of various bonds, e.g. C-O, C-N, C-C, ph
GO:0003676 - Interacting selectively and non-covalently with any nucleic acid.;
GO:0003676 - Interacting selectively and non-covalently with any nucleic acid.;
GO:0003676 - Interacting selectively and non-covalently with any nucleic acid.;
GO:0003723 - Interacting selectively and non-covalently with an RNA molecule
GO:0003676 - Interacting selectively and non-covalently with any nucleic acid.;
GO:0003676 - Interacting selectively and non-covalently with any nucleic acid.;
GO:0003676 - Interacting selectively and non-covalently with any nucleic acid.;
GO:0003676 - Interacting selectively and non-covalently with any nucleic acid.
GO:0006260 - The cellular metabolic process in which a cell duplicates one or r
GO:0016787 - Catalysis of the hydrolysis of various bonds, e.g. C-O, C-N, C-C, ph
GO:0003676 - Interacting selectively and non-covalently with any nucleic acid.;
GO:0003676 - Interacting selectively and non-covalently with any nucleic acid.
GO:0003676 - Interacting selectively and non-covalently with any nucleic acid.;
GO:0003676 - Interacting selectively and non-covalently with any nucleic acid.;
GO:0003676 - Interacting selectively and non-covalently with any nucleic acid.;
GO:0003676 - Interacting selectively and non-covalently with any nucleic acid.;
GO:0003676 - Interacting selectively and non-covalently with any nucleic acid.;
GO:0003676 - Interacting selectively and non-covalently with any nucleic acid.;
GO:0003676 - Interacting selectively and non-covalently with any nucleic acid.;
GO:0004824 - Catalysis of the reaction: ATP + L-lysine + t TRNA-CHARGING-PW
GO:0006412 - The cellular metabolic process in which a protein is formed, usin
GO:0003676 - Interacting selectively and non-covalently with any nucleic acid.;

GO:0003676 - Interacting selectively and non-covalently with any nucleic acid.;
GO:0003723 - Interacting selectively and non-covalently with an RNA molecule
GO:0003676 - Interacting selectively and non-covalently with any nucleic acid.;
GO:0003676 - Interacting selectively and non-covalently with any nucleic acid.;
GO:0004386 - Catalysis of the reaction: NTP + H2O = NDP + phosphate, to drive
GO:0003676 - Interacting selectively and non-covalently with any nucleic acid.;
GO:0003676 - Interacting selectively and non-covalently with any nucleic acid.;
GO:0003676 - Interacting selectively and non-covalently with any nucleic acid.;
GO:0003676 - Interacting selectively and non-covalently with any nucleic acid.;
GO:0003676 - Interacting selectively and non-covalently with any nucleic acid.
GO:0006260 - The cellular metabolic process in which a cell duplicates one or r
GO:0003676 - Interacting selectively and non-covalently with any nucleic acid.
GO:0003676 - Interacting selectively and non-covalently with any nucleic acid.
GO:0003676 - Interacting selectively and non-covalently with any nucleic acid.;
GO:0003676 - Interacting selectively and non-covalently with any nucleic acid.;
GO:0006139 - Any cellular metabolic process involving nucleobases, nucleoside
GO:0006139 - Any cellular metabolic process involving nucleobases, nucleoside
GO:0003676 - Interacting selectively and non-covalently with any nucleic acid.;
GO:0003723 - Interacting selectively and non-covalently with an RNA molecule
GO:0004527 - Catalysis of the hydrolysis of ester linkages within nucleic acids b
GO:0003676 - Interacting selectively and non-covalently with any nucleic acid.
GO:0003723 - Interacting selectively and non-covalently with an RNA molecule
GO:0003676 - Interacting selectively and non-covalently with any nucleic acid.;
GO:0003676 - Interacting selectively and non-covalently with any nucleic acid.;
GO:0006139 - Any cellular metabolic process involving nucleobases, nucleoside
GO:0003676 - Interacting selectively and non-covalently with any nucleic acid.;
GO:0003676 - Interacting selectively and non-covalently with any nucleic acid.
GO:0003723 - Interacting selectively and non-covalently with an RNA molecule
GO:0003676 - Interacting selectively and non-covalently with any nucleic acid.;
GO:0003676 - Interacting selectively and non-covalently with any nucleic acid.;
GO:0003676 - Interacting selectively and non-covalently with any nucleic acid.;
GO:0003723 - Interacting selectively and non-covalently with an RNA molecule
GO:0003723 - Interacting selectively and non-covalently with an RNA molecule
GO:0003723 - Interacting selectively and non-covalently with an RNA molecule
GO:0003676 - Interacting selectively and non-covalently with any nucleic acid.;
GO:0003723 - Interacting selectively and non-covalently with an RNA molecule

alias
GO:0008026 - Catalysis of the reaction: ATP + H2O = ADP + phosphate, to drive the unwinding o
GO:0000166 - Interacting selectively and non-covalently with a nucleotide, any compound consi
s, nucleotides and nucleic acids.; GO:0003676 - Interacting selectively and non-covalently with a
GO:0008270 - Interacting selectively and non-covalently with zinc (Zn) ions.
GO:0000166 - Interacting selectively and non-covalently with a nucleotide, any compound consi
GO:0000166 - Interacting selectively and non-covalently with a nucleotide, any compound consi
or a portion thereof.; GO:0003676 - Interacting selectively and non-covalently with any nucleic a
GO:0005524 - Interacting selectively and non-covalently with ATP, adenosine 5'-triphosphate, a
hosphorus.; GO:0006139 - Any cellular metabolic process involving nucleobases, nucleosides, nu
osphoric anhydride bonds, etc. Hydrolase is the systematic name for any enzyme of EC class 3.;
GO:0000166 - Interacting selectively and non-covalently with a nucleotide, any compound consi
GO:0000166 - Interacting selectively and non-covalently with a nucleotide, any compound consi
or a portion thereof.; GO:0030529 - An intracellular macromolecular complex containing both p
GO:0000166 - Interacting selectively and non-covalently with a nucleotide, any compound consi
of mRNA, messenger RNA, which is responsible for carrying the coded genetic 'message', tran
GO:0005524 - Interacting selectively and non-covalently with ATP, adenosine 5'-triphosphate, a

GO:0000166 - Interacting selectively and non-covalently with a nucleotide, any compound consi
GO:0000166 - Interacting selectively and non-covalently with a nucleotide, any compound consi
GO:0008270 - Interacting selectively and non-covalently with zinc (Zn) ions.
GO:0000166 - Interacting selectively and non-covalently with a nucleotide, any compound consi
GO:0000166 - Interacting selectively and non-covalently with a nucleotide, any compound consi
GO:0000166 - Interacting selectively and non-covalently with a nucleotide, any compound consi
the unwinding of a DNA or RNA helix.; GO:0003723 - Interacting selectively and non-covalently
GO:0000166 - Interacting selectively and non-covalently with a nucleotide, any compound consi
GO:0000166 - Interacting selectively and non-covalently with a nucleotide, any compound consi
GO:0000166 - Interacting selectively and non-covalently with a nucleotide, any compound consi
GO:0000166 - Interacting selectively and non-covalently with a nucleotide, any compound consi
a protein.; GO:0003676 - Interacting selectively and non-covalently with any nucleic acid.; GO:00
GO:0000166 - Interacting selectively and non-covalently with a nucleotide, any compound consi
= diphosphate + DNA(n+1); the synthesis of DNA from deoxyribonucleotide triphosphates in the
or a portion thereof.; GO:0003676 - Interacting selectively and non-covalently with any nucleic a
GO:0005524 - Interacting selectively and non-covalently with ATP, adenosine 5'-triphosphate, a
GO:0005524 - Interacting selectively and non-covalently with ATP, adenosine 5'-triphosphate, a
GO:0000166 - Interacting selectively and non-covalently with a nucleotide, any compound consi
GO:0005524 - Interacting selectively and non-covalently with ATP, adenosine 5'-triphosphate, a
GO:0005524 - Interacting selectively and non-covalently with ATP, adenosine 5'-triphosphate, a
GO:0000166 - Interacting selectively and non-covalently with a nucleotide, any compound consi
GO:0000166 - Interacting selectively and non-covalently with a nucleotide, any compound consi
GO:0000166 - Interacting selectively and non-covalently with a nucleotide, any compound consi
GO:0005524 - Interacting selectively and non-covalently with ATP, adenosine 5'-triphosphate, a
or a portion thereof.; GO:0003676 - Interacting selectively and non-covalently with any nucleic a
or a portion thereof.; GO:0045131 - Interacting selectively and non-covalently with a pre-mRNA
GO:0031124 - Any process involved in forming the mature 3' end of an mRNA molecule.; GO:00

\ strands or double helices through one another, resulting a topological transformation in doubl
GO:0005524 - Interacting selectively and non-covalently with ATP, adenosine 5'-triphosphate, a
GO:0000166 - Interacting selectively and non-covalently with a nucleotide, any compound consi:

GO:0000166 - Interacting selectively and non-covalently with a nucleotide, any compound consi:
GO:0005524 - Interacting selectively and non-covalently with ATP, adenosine 5'-triphosphate, a
or a portion thereof.; GO:0003676 - Interacting selectively and non-covalently with any nucleic a
NA (snRNA).; GO:0003676 - Interacting selectively and non-covalently with any nucleic acid.; GC
GO:0005524 - Interacting selectively and non-covalently with ATP, adenosine 5'-triphosphate, a
GO:0000166 - Interacting selectively and non-covalently with a nucleotide, any compound consi:
or a portion thereof.; GO:0003676 - Interacting selectively and non-covalently with any nucleic a
GO:0008270 - Interacting selectively and non-covalently with zinc (Zn) ions.

GO:0005524 - Interacting selectively and non-covalently with ATP, adenosine 5'-triphosphate, a
GO:0000166 - Interacting selectively and non-covalently with a nucleotide, any compound consi:
NA (snRNA).; GO:0003676 - Interacting selectively and non-covalently with any nucleic acid.; GC
GO:0000166 - Interacting selectively and non-covalently with a nucleotide, any compound consi:
or a portion thereof.; GO:0003676 - Interacting selectively and non-covalently with any nucleic a
or a portion thereof.; GO:0003676 - Interacting selectively and non-covalently with any nucleic a
GO:0005524 - Interacting selectively and non-covalently with ATP, adenosine 5'-triphosphate, a
GO:0005634 - A membrane-bounded organelle of eukaryotic cells in which chromosomes are ho
nosphoric anhydride bonds, etc. Hydrolase is the systematic name for any enzyme of EC class 3.;
GO:0000166 - Interacting selectively and non-covalently with a nucleotide, any compound consi:
GO:0000166 - Interacting selectively and non-covalently with a nucleotide, any compound consi:
GO:0008270 - Interacting selectively and non-covalently with zinc (Zn) ions.
or a portion thereof.
GO:0000166 - Interacting selectively and non-covalently with a nucleotide, any compound consi:
GO:0000166 - Interacting selectively and non-covalently with a nucleotide, any compound consi:
GO:0005634 - A membrane-bounded organelle of eukaryotic cells in which chromosomes are ho

nore molecules of DNA. DNA replication begins when specific sequences, known as origins of re
nosphoric anhydride bonds, etc. Hydrolase is the systematic name for any enzyme of EC class 3.;
GO:0000166 - Interacting selectively and non-covalently with a nucleotide, any compound consi:

GO:0000166 - Interacting selectively and non-covalently with a nucleotide, any compound consi:
GO:0000166 - Interacting selectively and non-covalently with a nucleotide, any compound consi:
GO:0005524 - Interacting selectively and non-covalently with ATP, adenosine 5'-triphosphate, a
GO:0005634 - A membrane-bounded organelle of eukaryotic cells in which chromosomes are ho
GO:0000166 - Interacting selectively and non-covalently with a nucleotide, any compound consi:
GO:0000166 - Interacting selectively and non-covalently with a nucleotide, any compound consi:
GO:0000166 - Interacting selectively and non-covalently with a nucleotide, any compound consi:
Y - ""
g the sequence of a mature mRNA molecule to specify the sequence of amino acids in a polype
GO:0000166 - Interacting selectively and non-covalently with a nucleotide, any compound consi:

GO:0008270 - Interacting selectively and non-covalently with zinc (Zn) ions.
or a portion thereof.; GO:0045131 - Interacting selectively and non-covalently with a pre-mRNA
GO:0005524 - Interacting selectively and non-covalently with ATP, adenosine 5'-triphosphate, a
GO:0000166 - Interacting selectively and non-covalently with a nucleotide, any compound consi:
the unwinding of a DNA or RNA helix.; GO:0003676 - Interacting selectively and non-covalently
GO:0006351 - The cellular synthesis of RNA on a template of DNA.; GO:0003899 - Catalysis of t
GO:0005524 - Interacting selectively and non-covalently with ATP, adenosine 5'-triphosphate, a
GO:0005515 - Interacting selectively and non-covalently with any protein or protein complex (a
GO:0005524 - Interacting selectively and non-covalently with ATP, adenosine 5'-triphosphate, a

nore molecules of DNA. DNA replication begins when specific sequences, known as origins of re

GO:0005524 - Interacting selectively and non-covalently with ATP, adenosine 5'-triphosphate, a
GO:0000166 - Interacting selectively and non-covalently with a nucleotide, any compound consi:
s, nucleotides and nucleic acids.; GO:0003676 - Interacting selectively and non-covalently with a
s, nucleotides and nucleic acids.; GO:0003676 - Interacting selectively and non-covalently with a
GO:0000166 - Interacting selectively and non-covalently with a nucleotide, any compound consi:
or a portion thereof.; GO:0003676 - Interacting selectively and non-covalently with any nucleic a
by removing nucleotide residues from the 3' or 5' end.; GO:0003676 - Interacting selectively and

or a portion thereof.; GO:0003676 - Interacting selectively and non-covalently with any nucleic a
GO:0006397 - Any process involved in the conversion of a primary mRNA transcript into one or r
GO:0000166 - Interacting selectively and non-covalently with a nucleotide, any compound consi:
s, nucleotides and nucleic acids.; GO:0003676 - Interacting selectively and non-covalently with a
GO:0005524 - Interacting selectively and non-covalently with ATP, adenosine 5'-triphosphate, a

or a portion thereof.; GO:0003676 - Interacting selectively and non-covalently with any nucleic a
GO:0005524 - Interacting selectively and non-covalently with ATP, adenosine 5'-triphosphate, a
GO:0046872 - Interacting selectively and non-covalently with any metal ion.
GO:0005515 - Interacting selectively and non-covalently with any protein or protein complex (a
or a portion thereof.; GO:0003676 - Interacting selectively and non-covalently with any nucleic a
or a portion thereof.; GO:0003676 - Interacting selectively and non-covalently with any nucleic a
or a portion thereof.; GO:0003676 - Interacting selectively and non-covalently with any nucleic a
GO:0050825 - Interacting selectively and non-covalently with ice, water reduced to the solid sta
or a portion thereof.; GO:0005515 - Interacting selectively and non-covalently with any protein

f a DNA or RNA helix.; GO:0005524 - Interacting selectively and non-covalently with ATP, ade
sting of a nucleoside that is esterified with (ortho)phosphate or an oligophosphate at any hyd
ıny nucleic acid.; GO:0008408 - Catalysis of the hydrolysis of ester linkages within nucleic acic

sting of a nucleoside that is esterified with (ortho)phosphate or an oligophosphate at any hyd
sting of a nucleoside that is esterified with (ortho)phosphate or an oligophosphate at any hyd
acid.
universally important coenzyme and enzyme regulator.
ıcleotides and nucleic acids.; GO:0003676 - Interacting selectively and non-covalently with an
GO:0003676 - Interacting selectively and non-covalently with any nucleic acid.; GO:0005524 -
sting of a nucleoside that is esterified with (ortho)phosphate or an oligophosphate at any hyd
sting of a nucleoside that is esterified with (ortho)phosphate or an oligophosphate at any hyd
rotein and RNA molecules.; GO:0003676 - Interacting selectively and non-covalently with any
sting of a nucleoside that is esterified with (ortho)phosphate or an oligophosphate at any hyd
ıscribed from DNA, to sites of protein assembly at the ribosomes.; GO:0003723 - Interacting s
universally important coenzyme and enzyme regulator.

sting of a nucleoside that is esterified with (ortho)phosphate or an oligophosphate at any hyd
sting of a nucleoside that is esterified with (ortho)phosphate or an oligophosphate at any hyd

sting of a nucleoside that is esterified with (ortho)phosphate or an oligophosphate at any hyd
sting of a nucleoside that is esterified with (ortho)phosphate or an oligophosphate at any hyd
sting of a nucleoside that is esterified with (ortho)phosphate or an oligophosphate at any hyd
with an RNA molecule or a portion thereof.; GO:0003676 - Interacting selectively and non-co
sting of a nucleoside that is esterified with (ortho)phosphate or an oligophosphate at any hyd
sting of a nucleoside that is esterified with (ortho)phosphate or an oligophosphate at any hyd
sting of a nucleoside that is esterified with (ortho)phosphate or an oligophosphate at any hyd
sting of a nucleoside that is esterified with (ortho)phosphate or an oligophosphate at any hyd
)04842 - Catalysis of the transfer of ubiquitin from one protein to another via the reaction X-U
sting of a nucleoside that is esterified with (ortho)phosphate or an oligophosphate at any hyd
presence of a DNA template and a 3'hydroxyl group.; GO:0006260 - The cellular metabolic p
acid.
universally important coenzyme and enzyme regulator.
universally important coenzyme and enzyme regulator.
sting of a nucleoside that is esterified with (ortho)phosphate or an oligophosphate at any hyd
universally important coenzyme and enzyme regulator.
universally important coenzyme and enzyme regulator.
sting of a nucleoside that is esterified with (ortho)phosphate or an oligophosphate at any hyd
sting of a nucleoside that is esterified with (ortho)phosphate or an oligophosphate at any hyd
sting of a nucleoside that is esterified with (ortho)phosphate or an oligophosphate at any hyd
universally important coenzyme and enzyme regulator.
acid.
branch point sequence, located upstream of the 3' splice site.; GO:0003676 - Interacting sel
)00166 - Interacting selectively and non-covalently with a nucleotide, any compound consistin

e-stranded DNA.; GO:0003676 - Interacting selectively and non-covalently with any nucleic ac
universally important coenzyme and enzyme regulator.
sting of a nucleoside that is esterified with (ortho)phosphate or an oligophosphate at any hyd

sting of a nucleoside that is esterified with (ortho)phosphate or an oligophosphate at any hyd
universally important coenzyme and enzyme regulator.
acid.
):0000166 - Interacting selectively and non-covalently with a nucleotide, any compound consi
universally important coenzyme and enzyme regulator.
sting of a nucleoside that is esterified with (ortho)phosphate or an oligophosphate at any hyd
acid.

universally important coenzyme and enzyme regulator.
sting of a nucleoside that is esterified with (ortho)phosphate or an oligophosphate at any hyd
):0000166 - Interacting selectively and non-covalently with a nucleotide, any compound consi
sting of a nucleoside that is esterified with (ortho)phosphate or an oligophosphate at any hyd
acid.
acid.
universally important coenzyme and enzyme regulator.
oused and replicated. In most cells, the nucleus contains all of the cell's chromosomes except
GO:0003676 - Interacting selectively and non-covalently with any nucleic acid.; GO:0046872 -
sting of a nucleoside that is esterified with (ortho)phosphate or an oligophosphate at any hyd
sting of a nucleoside that is esterified with (ortho)phosphate or an oligophosphate at any hyd

sting of a nucleoside that is esterified with (ortho)phosphate or an oligophosphate at any hyd
sting of a nucleoside that is esterified with (ortho)phosphate or an oligophosphate at any hyd
oused and replicated. In most cells, the nucleus contains all of the cell's chromosomes except

eplication, are recognized and bound by initiation proteins, and ends when the original DNA m
GO:0016818 - Catalysis of the hydrolysis of any acid anhydride which contains phosphorus.; G
sting of a nucleoside that is esterified with (ortho)phosphate or an oligophosphate at any hyd

sting of a nucleoside that is esterified with (ortho)phosphate or an oligophosphate at any hyd
sting of a nucleoside that is esterified with (ortho)phosphate or an oligophosphate at any hyd
universally important coenzyme and enzyme regulator.; GO:0008270 - Interacting selectively
oused and replicated. In most cells, the nucleus contains all of the cell's chromosomes except
sting of a nucleoside that is esterified with (ortho)phosphate or an oligophosphate at any hyd
sting of a nucleoside that is esterified with (ortho)phosphate or an oligophosphate at any hyd
sting of a nucleoside that is esterified with (ortho)phosphate or an oligophosphate at any hyd

ptide chain. Translation is mediated by the ribosome, and begins with the formation of a terr
sting of a nucleoside that is esterified with (ortho)phosphate or an oligophosphate at any hyd

\ branch point sequence, located upstream of the 3' splice site.; GO:0003676 - Interacting sele
universally important coenzyme and enzyme regulator.
sting of a nucleoside that is esterified with (ortho)phosphate or an oligophosphate at any hyd
with any nucleic acid.; GO:0005524 - Interacting selectively and non-covalently with ATP, ade
he reaction: nucleoside triphosphate + RNA(n) = diphosphate + RNA(n+1). Utilizes a DNA tem
universally important coenzyme and enzyme regulator.
complex of two or more proteins that may include other nonprotein molecules).; GO:000552
universally important coenzyme and enzyme regulator.

plication, are recognized and bound by initiation proteins, and ends when the original DNA m

universally important coenzyme and enzyme regulator.
sting of a nucleoside that is esterified with (ortho)phosphate or an oligophosphate at any hyd
ny nucleic acid.; GO:0000166 - Interacting selectively and non-covalently with a nucleotide, a
ny nucleic acid.; GO:0000166 - Interacting selectively and non-covalently with a nucleotide, a
sting of a nucleoside that is esterified with (ortho)phosphate or an oligophosphate at any hyd
acid.; GO:0005524 - Interacting selectively and non-covalently with ATP, adenosine 5'-triphos
non-covalently with any nucleic acid.

acid.
more mature mRNA(s) prior to translation into polypeptide.
sting of a nucleoside that is esterified with (ortho)phosphate or an oligophosphate at any hyd
ny nucleic acid.; GO:0008408 - Catalysis of the hydrolysis of ester linkages within nucleic acid
universally important coenzyme and enzyme regulator.

acid.
universally important coenzyme and enzyme regulator.

complex of two or more proteins that may include other nonprotein molecules).; GO:000016
acid.
acid.
acid.
te by cold temperature. It is a white or transparent colorless substance, crystalline, brittle, ar
or protein complex (a complex of two or more proteins that may include other nonprotein mo

nosine 5'-triphosphate, a universally important coenzyme and enzyme regulator.; GO:000631
lroxyl group on the ribose or deoxyribose.
ls by removing nucleotide residues from the 3' end.

lroxyl group on the ribose or deoxyribose.
lroxyl group on the ribose or deoxyribose.

ıy nucleic acid.; GO:0008026 - Catalysis of the reaction: ATP + H2O = ADP + phosphate, to driv
· Interacting selectively and non-covalently with ATP, adenosine 5'-triphosphate, a universally
lroxyl group on the ribose or deoxyribose.
lroxyl group on the ribose or deoxyribose.
ʼ nucleic acid.; GO:0000166 - Interacting selectively and non-covalently with a nucleotide, any
lroxyl group on the ribose or deoxyribose.; GO:0050825 - Interacting selectively and non-coval
selectively and non-covalently with an RNA molecule or a portion thereof.; GO:0004654 - Cata

lroxyl group on the ribose or deoxyribose.
lroxyl group on the ribose or deoxyribose.

lroxyl group on the ribose or deoxyribose.
lroxyl group on the ribose or deoxyribose.
lroxyl group on the ribose or deoxyribose.; GO:0003824 - Catalysis of a biochemical reaction a
valently with any nucleic acid.; GO:0005634 - A membrane-bounded organelle of eukaryotic c
lroxyl group on the ribose or deoxyribose.; GO:0005488 - The selective, non-covalent, often st
lroxyl group on the ribose or deoxyribose.
lroxyl group on the ribose or deoxyribose.; GO:0050825 - Interacting selectively and non-coval
lroxyl group on the ribose or deoxyribose.
Jb + Y --> Y-Ub + X, where both X-Ub and Y-Ub are covalent linkages.
lroxyl group on the ribose or deoxyribose.
process in which a cell duplicates one or more molecules of DNA. DNA replication begins wher

lroxyl group on the ribose or deoxyribose.

lroxyl group on the ribose or deoxyribose.
lroxyl group on the ribose or deoxyribose.
lroxyl group on the ribose or deoxyribose.

ectively and non-covalently with any nucleic acid.; GO:0000166 - Interacting selectively and nc
g of a nucleoside that is esterified with (ortho)phosphate or an oligophosphate at any hydrox

id.; GO:0006265 - The process in which a transformation is induced in the topological structu

lroxyl group on the ribose or deoxyribose.

lroxyl group on the ribose or deoxyribose.

sting of a nucleoside that is esterified with (ortho)phosphate or an oligophosphate at any hyd

lroxyl group on the ribose or deoxyribose.

lroxyl group on the ribose or deoxyribose.
sting of a nucleoside that is esterified with (ortho)phosphate or an oligophosphate at any hyd
lroxyl group on the ribose or deoxyribose.

: the organellar chromosomes, and is the site of RNA synthesis and processing. In some speci
- Interacting selectively and non-covalently with any metal ion.
lroxyl group on the ribose or deoxyribose.
lroxyl group on the ribose or deoxyribose.

lroxyl group on the ribose or deoxyribose.
lroxyl group on the ribose or deoxyribose.
: the organellar chromosomes, and is the site of RNA synthesis and processing. In some speci

iolecule has been completely duplicated and the copies topologically separated. The unit of re
iO:0006139 - Any cellular metabolic process involving nucleobases, nucleosides, nucleotides a
lroxyl group on the ribose or deoxyribose.

lroxyl group on the ribose or deoxyribose.
lroxyl group on the ribose or deoxyribose.
and non-covalently with zinc (Zn) ions.
: the organellar chromosomes, and is the site of RNA synthesis and processing. In some speci
lroxyl group on the ribose or deoxyribose.
lroxyl group on the ribose or deoxyribose.; GO:0008026 - Catalysis of the reaction: ATP + H2O
lroxyl group on the ribose or deoxyribose.; GO:0046872 - Interacting selectively and non-coval

iary complex between aminoacylated initiator methionine tRNA, GTP, and initiation factor 2,
lroxyl group on the ribose or deoxyribose.

ectively and non-covalently with any nucleic acid.; GO:0000166 - Interacting selectively and nc

lroxyl group on the ribose or deoxyribose.
enosine 5'-triphosphate, a universally important coenzyme and enzyme regulator.
nplate, i.e. the catalysis of DNA-template-directed extension of the 3'-end of an RNA strand b

4 - Interacting selectively and non-covalently with ATP, adenosine 5'-triphosphate, a universal

nolecule has been completely duplicated and the copies topologically separated. The unit of re

lroxyl group on the ribose or deoxyribose.
ny compound consisting of a nucleoside that is esterified with (ortho)phosphate or an oligoph
ny compound consisting of a nucleoside that is esterified with (ortho)phosphate or an oligoph
lroxyl group on the ribose or deoxyribose.
phate, a universally important coenzyme and enzyme regulator.

lroxyl group on the ribose or deoxyribose.
ls by removing nucleotide residues from the 3' end.

5 - Interacting selectively and non-covalently with a nucleotide, any compound consisting of a

nd viscoidal.; GO:0006355 - Any process that modulates the frequency, rate or extent of cellul
olecules).

0 - Any process in which a new genotype is formed by reassortment of genes resulting in gen

e the unwinding of a DNA or RNA helix.; GO:0004003 - Catalysis of the reaction: ATP + H2O =
 important coenzyme and enzyme regulator.; GO:0003677 - Any molecular function by which

 compound consisting of a nucleoside that is esterified with (ortho)phosphate or an oligophos
ently with ice, water reduced to the solid state by cold temperature. It is a white or transpare
ılysis of the reaction: ATP + RNA(n) = diphosphate + RNA(n+1).; GO:0003676 - Interacting sele

t physiological temperatures. In biologically catalyzed reactions, the reactants are known as s
:ells in which chromosomes are housed and replicated. In most cells, the nucleus contains all
ɔichiometric, interaction of a molecule with one or more specific sites on another molecule.

ently with ice, water reduced to the solid state by cold temperature. It is a white or transpare

n specific sequences, known as origins of replication, are recognized and bound by initiation p

ɔn-covalently with a nucleotide, any compound consisting of a nucleoside that is esterified wi
yl group on the ribose or deoxyribose.

re of a double-stranded DNA helix, resulting in a change in linking number.; GO:0003917 - Cat

roxyl group on the ribose or deoxyribose.; GO:0000398 - The joining together of exons from o

roxyl group on the ribose or deoxyribose.; GO:0000398 - The joining together of exons from o

es, or in specialized cell types, RNA metabolism or DNA replication may be absent.; GO:0003

es, or in specialized cell types, RNA metabolism or DNA replication may be absent.; GO:0008

eplication usually corresponds to the genome of the cell, an organelle, or a virus. The templat
and nucleic acids.; GO:0003676 - Interacting selectively and non-covalently with any nucleic ac

es, or in specialized cell types, RNA metabolism or DNA replication may be absent.

= ADP + phosphate, to drive the unwinding of a DNA or RNA helix.; GO:0003824 - Catalysis of
ently with any metal ion.

which subsequently associates with the small subunit of the ribosome and an mRNA. Transl on-covalently with a nucleotide, any compound consisting of a nucleoside that is esterified wi

y one nucleotide at a time. Can initiate a chain 'de novo'.

ly important coenzyme and enzyme regulator.

eplication usually corresponds to the genome of the cell, an organelle, or a virus. The templat

hosphate at any hydroxyl group on the ribose or deoxyribose.; GO:0008408 - Catalysis of the hy

hosphate at any hydroxyl group on the ribose or deoxyribose.; GO:0008408 - Catalysis of the hy

nucleoside that is esterified with (ortho)phosphate or an oligophosphate at any hydroxyl grou

ar DNA-templated transcription.; GO:0042309 - Any homoeostatic process in which an organi ıe combinations different from those that were present in the parents. In eukaryotes genetic ı

ADP + phosphate; this reaction drives the unwinding of the DNA helix.; GO:0005524 - Interac
a gene product interacts selectively and non-covalently with DNA (deoxyribonucleic acid).

;phate at any hydroxyl group on the ribose or deoxyribose.; GO:0005634 - A membrane-bound
ent colorless substance, crystalline, brittle, and viscoidal.; GO:0042309 - Any homoeostatic pro
ectively and non-covalently with any nucleic acid.; GO:0000175 - Catalysis of the sequential cl

substrates, and the catalysts are naturally occurring macromolecular substances known as en
of the cell's chromosomes except the organellar chromosomes, and is the site of RNA synthe

ent colorless substance, crystalline, brittle, and viscoidal.; GO:0042309 - Any homoeostatic pro

proteins, and ends when the original DNA molecule has been completely duplicated and the c

th (ortho)phosphate or an oligophosphate at any hydroxyl group on the ribose or deoxyribose.

talysis of a DNA topological transformation by transiently cleaving one DNA strand at a time t

ne or more primary transcripts of messenger RNA (mRNA) and the excision of intron sequenc

ne or more primary transcripts of messenger RNA (mRNA) and the excision of intron sequenc

677 - Any molecular function by which a gene product interacts selectively and non-covalently

026 - Catalysis of the reaction: ATP + H2O = ADP + phosphate, to drive the unwinding of a DN

e for replication can either be an existing DNA molecule or RNA.; GO:0003676 - Interacting se
cid.; GO:0005634 - A membrane-bounded organelle of eukaryotic cells in which chromosomes

a biochemical reaction at physiological temperatures. In biologically catalyzed reactions, the

ation ends with the release of a polypeptide chain from the ribosome.; GO:0006414 - The suc th (ortho)phosphate or an oligophosphate at any hydroxyl group on the ribose or deoxyribose.

e for replication can either be an existing DNA molecule or RNA.; GO:0003676 - Interacting s[illegible]

ydrolysis of ester linkages within nucleic acids by removing nucleotide residues from the 3' er

ydrolysis of ester linkages within nucleic acids by removing nucleotide residues from the 3' er

ıp on the ribose or deoxyribose.

sm maintains its internal body temperature at a relatively constant value. This is achieved by recombination can occur by chromosome assortment, intrachromosomal recombination, or n

ting selectively and non-covalently with ATP, adenosine 5'-triphosphate, a universally importa

led organelle of eukaryotic cells in which chromosomes are housed and replicated. In most ce
ocess in which an organism maintains its internal body temperature at a relatively constant va
eavage of mononucleotides from a free 3' terminus of an RNA molecule.; GO:0006396 - Any

zymes. Enzymes possess specific binding sites for substrates, and are usually composed whol
esis and processing. In some species, or in specialized cell types, RNA metabolism or DNA rep

ocess in which an organism maintains its internal body temperature at a relatively constant va

opies topologically separated. The unit of replication usually corresponds to the genome of th

; GO:0000398 - The joining together of exons from one or more primary transcripts of messe to allow passage of another strand; changes the linking number by +1 per catalytic cycle.; GO

ces, via a spliceosomal mechanism, so that mRNA consisting only of the joined exons is produ

ces, via a spliceosomal mechanism, so that mRNA consisting only of the joined exons is produ

/ with DNA (deoxyribonucleic acid).; GO:0006355 - Any process that modulates the frequency,

A or RNA helix.; GO:0006351 - The cellular synthesis of RNA on a template of DNA.; GO:0005

electively and non-covalently with any nucleic acid.; GO:0005634 - A membrane-bounded orga
s are housed and replicated. In most cells, the nucleus contains all of the cell's chromosomes

reactants are known as substrates, and the catalysts are naturally occurring macromolecular

cessive addition of amino acid residues to a nascent polypeptide chain during protein biosynt

; GO:0000398 - The joining together of exons from one or more primary transcripts of messe

electively and non-covalently with any nucleic acid.; GO:0005634 - A membrane-bounded orga

nd.; GO:0003824 - Catalysis of a biochemical reaction at physiological temperatures. In biolog
nd.; GO:0003824 - Catalysis of a biochemical reaction at physiological temperatures. In biolog

using metabolic processes to counteract fluctuations in the temperature of the environment onreciprocal interchromosomal recombination. Intrachromosomal recombination occurs by cr

ınt coenzyme and enzyme regulator.; GO:0003677 - Any molecular function by which a gene p

ells, the nucleus contains all of the cell's chromosomes except the organellar chromosomes, a
alue. This is achieved by using metabolic processes to counteract fluctuations in the tempera
process involved in the conversion of one or more primary RNA transcripts into one or more r

ly or largely of protein, but RNA that has catalytic activity (ribozyme) is often also regarded a
lication may be absent.; GO:0005524 - Interacting selectively and non-covalently with ATP, ad

alue. This is achieved by using metabolic processes to counteract fluctuations in the tempera

e cell, an organelle, or a virus. The template for replication can either be an existing DNA mo

nger RNA (mRNA) and the excision of intron sequences, via a spliceosomal mechanism, so th

:0003677 - Any molecular function by which a gene product interacts selectively and non-cova

ıced.

ıced.

, rate or extent of cellular DNA-templated transcription.

524 - Interacting selectively and non-covalently with ATP, adenosine 5'-triphosphate, a univer

anelle of eukaryotic cells in which chromosomes are housed and replicated. In most cells, the
except the organellar chromosomes, and is the site of RNA synthesis and processing. In som

substances known as enzymes. Enzymes possess specific binding sites for substrates, and ar

hesis.; GO:0003723 - Interacting selectively and non-covalently with an RNA molecule or a po nger RNA (mRNA) and the excision of intron sequences, via a spliceosomal mechanism, so th

anelle of eukaryotic cells in which chromosomes are housed and replicated. In most cells, the

;ically catalyzed reactions, the reactants are known as substrates, and the catalysts are natur

;ically catalyzed reactions, the reactants are known as substrates, and the catalysts are natur

.; GO:0050826 - Any process that results in a change in state or activity of a cell or an organis rossing over. In bacteria it may occur by genetic transformation, conjugation, transduction, or

product interacts selectively and non-covalently with DNA (deoxyribonucleic acid).

nd is the site of RNA synthesis and processing. In some species, or in specialized cell types, R
ture of the environment.; GO:0050826 - Any process that results in a change in state or activi
nature RNA molecules.

s enzymatic.; GO:0005622 - The living contents of a cell; the matter contained within (but no
denosine 5'-triphosphate, a universally important coenzyme and enzyme regulator.

ture of the environment.; GO:0050826 - Any process that results in a change in state or activi

lecule or RNA.; GO:0003676 - Interacting selectively and non-covalently with any nucleic acid

at mRNA consisting only of the joined exons is produced.

ılently with DNA (deoxyribonucleic acid).; GO:0008270 - Interacting selectively and non-covale

rsally important coenzyme and enzyme regulator.; GO:0006310 - Any process in which a new

nucleus contains all of the cell's chromosomes except the organellar chromosomes, and is th
e species, or in specialized cell types, RNA metabolism or DNA replication may be absent.; G(

re usually composed wholly or largely of protein, but RNA that has catalytic activity (ribozyme

rtion thereof.; GO:0003746 - Functions in chain elongation during polypeptide synthesis at th ıat mRNA consisting only of the joined exons is produced.; GO:0008270 - Interacting selective

nucleus contains all of the cell's chromosomes except the organellar chromosomes, and is tl

ally occurring macromolecular substances known as enzymes. Enzymes possess specific bindi
ally occurring macromolecular substances known as enzymes. Enzymes possess specific bindi

m (in terms of movement, secretion, enzyme production, gene expression, etc.) as a result o

F-duction.

RNA metabolism or DNA replication may be absent.; GO:0006396 - Any process involved in the
ty of a cell or an organism (in terms of movement, secretion, enzyme production, gene expres

t including) the plasma membrane, usually taken to exclude large vacuoles and masses of se

ty of a cell or an organism (in terms of movement, secretion, enzyme production, gene expres

.; GO:0005524 - Interacting selectively and non-covalently with ATP, adenosine 5'-triphosphat ently with zinc (Zn) ions.

genotype is formed by reassortment of genes resulting in gene combinations different from

ne site of RNA synthesis and processing. In some species, or in specialized cell types, RNA me
O:0006289 - A DNA repair process in which a small region of the strand surrounding the dama

) is often also regarded as enzymatic.; GO:0005524 - Interacting selectively and non-covalent

e ribosome.; GO:0003676 - Interacting selectively and non-covalently with any nucleic acid.; G ly and non-covalently with zinc (Zn) ions.

ne site of RNA synthesis and processing. In some species, or in specialized cell types, RNA me

ng sites for substrates, and are usually composed wholly or largely of protein, but RNA that h
ng sites for substrates, and are usually composed wholly or largely of protein, but RNA that h

f a freezing stimulus, temperatures below 0 degrees Celsius.

e conversion of one or more primary RNA transcripts into one or more mature RNA molecules
ssion, etc.) as a result of a freezing stimulus, temperatures below 0 degrees Celsius.

cretory or ingested material. In eukaryotes it includes the nucleus and cytoplasm.; GO:00442[illegible]

ssion, etc.) as a result of a freezing stimulus, temperatures below 0 degrees Celsius.

te, a universally important coenzyme and enzyme regulator.; GO:0003677 - Any molecular fur those that were present in the parents. In eukaryotes genetic recombination can occur by chro

etabolism or DNA replication may be absent.; GO:0006310 - Any process in which a new geno
age is removed from the DNA helix as an oligonucleotide. The small gap left in the DNA helix

tly with ATP, adenosine 5'-triphosphate, a universally important coenzyme and enzyme regula

GO:0005515 - Interacting selectively and non-covalently with any protein or protein complex (a etabolism or DNA replication may be absent.; GO:0006310 - Any process in which a new geno

as catalytic activity (ribozyme) is often also regarded as enzymatic.; GO:0005622 - The living
as catalytic activity (ribozyme) is often also regarded as enzymatic.; GO:0005622 - The living

;.

37 - The chemica

nction by which a gene prod

omosome assortment, intrachromosom

type is formed by reassortm
is filled in by the sequ

ator.; GO:0006310 -

a complex of two or more pro

type is formed by reassortm

contents of
contents of
